# Misleading Success: Genomes Reveal Critical Risks to European Gray Wolves

**DOI:** 10.64898/2026.03.20.713253

**Authors:** Sara Ravagni, Daniele Battilani, Isabel Salado, Diana Lobo, Carlos Sarabia, Carlos Leiva, Romolo Caniglia, Elena Fabbri, Paolo Ciucci, Matteo Girardi, Francisco Iglesias-Santos, Josip Kusak, Federica Mattucci, Morteza Naderi, Carsten Nowak, Çağan H. Şekercioğlu, Tomasz Skrbinsek, Edoardo Velli, Astrid Vik Stronen, Carles Vila, Raquel Godinho, Jennifer A. Leonard, Cristiano Vernesi

## Abstract

Have European gray wolves recovered? Despite an increase to ∼21,000 wolves (*Canis lupus*), our genomic analyses reveal significant risks to their long-term viability. We analyzed over 200 whole-genomes spanning five major European populations. Rather than a single recovering population, European wolves form a mosaic of isolated, independently evolving lineages, mostly diverging in the late Pleistocene. All lineages have contemporary effective population sizes below the threshold for long-term viability (Ne ≥ 500) and show extensive inbreeding. Runs of homozygosity reveal population-specific inbreeding histories spanning recent to deep timeframes. Most lineages exhibit higher realized than masked genetic load, indicating emerging inbreeding depression. These findings challenge claims that downlisting European wolves is biologically warranted: none of these populations currently meets thresholds associated with favorable conservation status.

## Main Text

Persecution and habitat loss in the 19th and 20th centuries decimated Europe’s large carnivores, with the gray wolf (*Canis lupus*) a key example of range contraction (*1*). Unlike wolves in North America, where recovery has been actively supported (*2*), wolf recolonization across Europe over the last ∼50 years has been driven by natural dispersal from a handful of remnant peninsular and eastern source populations (*1*). The refugial populations – principally in the Italian Peninsula, Iberian Peninsula and the Dinaric-Balkan region – now constitute the genetic backbone of central and western Europe’s wolves, but carry the genomic scars of historical bottlenecks and prolonged isolation. In the current scenario of global genetic diversity loss (*3*), maintaining adequate genetic variation within populations has been recognized as one of the four overarching goals of the Convention on Biological Diversity’s Global Biodiversity Framework to ensure long-term species viability (*4*). In Europe, this perspective is being integrated into law: the EU Nature Restoration Law explicitly links ecological recovery to measurable outcomes, and under the Habitats Directive (92/43/EEC) there is growing recognition that Favourable Conservation Status (FCS) must incorporate population-genetic criteria such as effective population size (Ne), not just census size and range (*5*). For long-term evolutionary resilience, a widely used benchmark is Ne ≥ 500, with Ne ≥ 50 considered a minimum to avoid severe short-term inbreeding depression and increased risk of extinction (*6*, *7*).

The gray wolf is emblematic in this debate. Legal protections over recent decades have allowed European wolves to rebound demographically and geographically, with current estimates exceeding 20,000 individuals and an important recovery in their historical range (*8*). This apparent success, together with increasing human-wildlife conflicts including attacks on livestock, has contributed to political pressure to reduce protection levels. In 2025 wolves were downgraded from “strictly protected” to “protected” status at the EU level, granting member states broader latitude for lethal control (*9*). Current controversies around wolf management and culling limits (*10*, *11*) highlight a critical uncertainty: do increased numbers and range recovery reflect a genuinely secure conservation status, or do they mask underlying genetic fragility?

Earlier genetic studies using divergent genetic and genomic markers – mtDNA, microsatellites and SNPs – suggested structure among European wolves (*12*). Distinct demographic histories were reported for the Scandinavian, Italian, Dinaric-Balkan and NW Iberian (Iberian hereafter) populations (*13*), and low genetic diversity and inbreeding (*14–16*) as well as inbreeding depression (*17*, *18*) were locally documented. The now-extinct Sierra Morena lineage in southern Spain exemplifies how long-term isolation, genetic erosion and hybridization with dogs can drive local extinction (*19*). However, these studies lacked the resolution to quantify genome-wide diversity, inbreeding, and genetic load across populations, or to determine whether European wolves function as a single recovering metapopulation or as multiple independently evolving lineages of differing conservation concern.

Here we use whole-genome data to clarify the demographic and evolutionary histories of European wolves and assess their genetic status in the context of current policy. We generated whole-genome sequences for 89 European and 16 Turkish wolves. These data were analyzed together with 176 publicly available genomes from European, Asian and North American wolves, as well as domestic dogs and other canids, yielding one of the most comprehensive European wolf genomic datasets compiled to date (Tables S1, S2). These analyses focus on five previously identified European populations: Dinaric-Balkan, Italian Peninsula, Karelian, Iberian and Scandinavian (*1*, *8*). After depth and quality filtering, we removed wolves with >5% dog ancestry (see supplementary materials, Tables S3-S5), duplicates, and first-degree relatives (Table S6). The resulting dataset included 16,497,458 SNPs for 123 European wolves (Dinaric-Balkan, n = 27; Italian Peninsula, n = 35; Karelian, n = 27; Iberian, n = 17; Scandinavian, n = 17), 51 Asian wolves (Western Asia, n = 6; Türkiye, n = 13; Central Asia, n = 12; Eastern Asia, n = 12; Indian, n = 1; Tibetan, n = 7 – wolves classified into populations following reference *20*) and 27 North American wolves (Fig. 1A, Table S7), complemented with 197 domestic dogs and three canid outgroups.

**Fig. 1.**
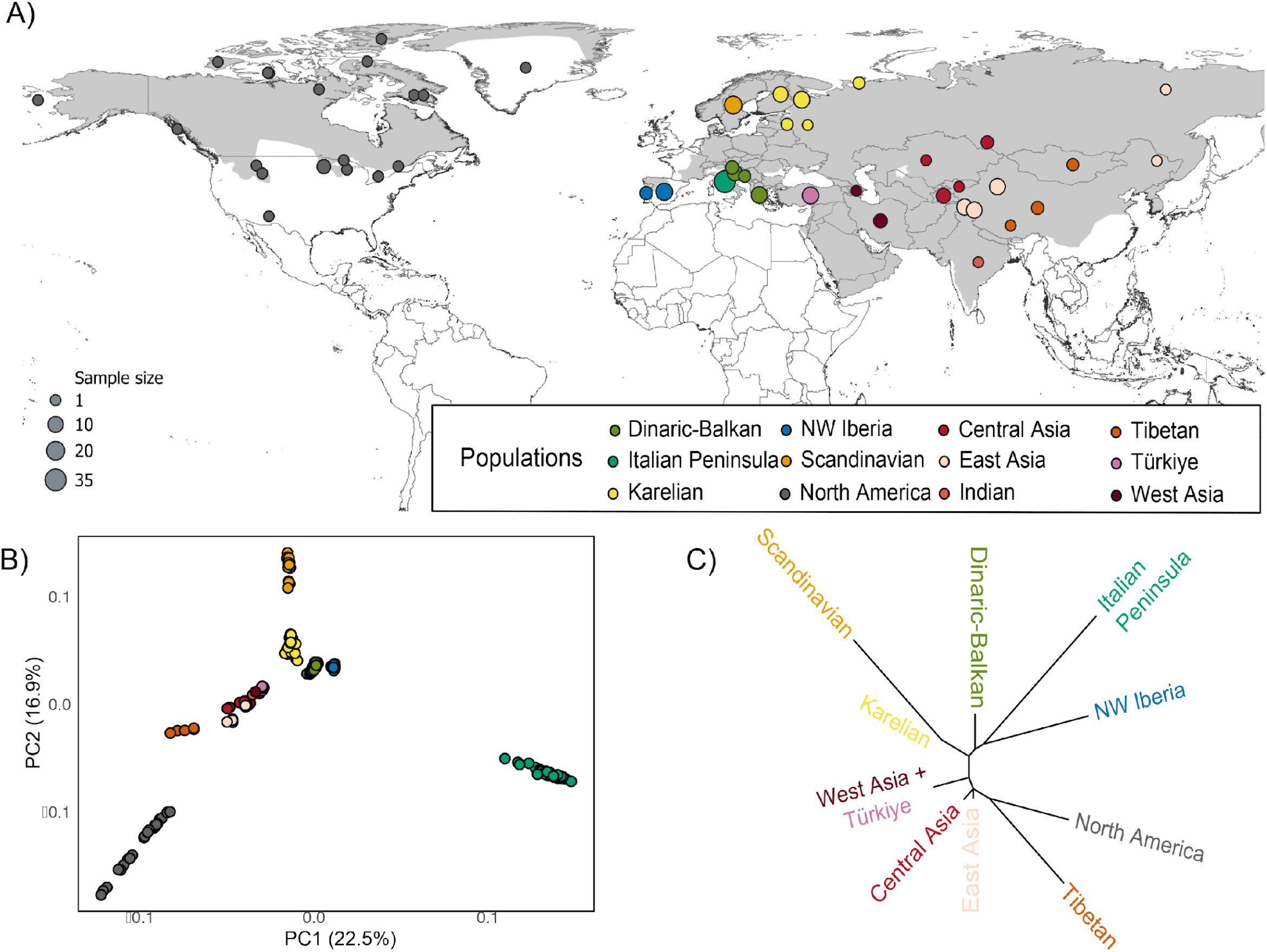
Gray wolf populations differentiation. (A) Global distribution of whole-genome data used. Circle size indicates sample size by locality, colored by population assignment. Gray shadow area corresponds to the species distribution (*38*). When no detailed locality information was available for samples, the centroid per country was used. (B) Principal Component Analysis (PCA) of wolves based on 9,068,899 genome-wide SNPs. (C) Maximum-likelihood tree of the different populations.

### Five distinct lineages across Europe

Principal component analysis cleanly separated wolves by continent (Fig. 1B). Within Europe, wolves formed five distinct clusters corresponding to populations. Italian Peninsula wolves were the most divergent along PC1, while Dinaric-Balkan, Karelian, Iberian and Scandinavian wolves occupied separate positions along PC1 and PC2. ADMIXTURE analyses were consistent with these results, with all five European wolf populations showing unique genetic ancestry components (Figs. S1-S2, Tables S8-S9) and minimal evidence of recent gene flow among southern populations. Two Scandinavian wolves displayed substantial Karelian ancestry, consistent with known recent immigration (*21*). Phylogenomic trees supported the same population structure with Karelian and Scandinavian forming a well-supported clade, and southern European populations appearing as highly divergent (Figs. 1C, S3-S5). These combined lines of evidence indicate that differentiation may result from long-term isolation in addition to recent anthropogenic bottlenecks.

### Deep divergences among European wolf populations

Dated phylogenomic analyses using complementary methods – MiSTI (22) and the TT-method (*23*) – placed the divergence of most European populations in the late Pleistocene (Table S10; see also *13*), coinciding with population bottlenecks between 10,000-20,000 years ago (Figs. S6-S7). Divergence time estimates were generally concordant between methods (e.g., for Italian Peninsula vs. Dinaric-Balkan wolves: MiSTI ∼12,869 years, TT ∼12,147 years). The extreme genetic drift in the Scandinavian population, founded by just three immigrants from Karelia in the 1980s (*24*), has greatly accelerated its differentiation, yielding an older estimate in MiSTI than the historically documented founding date captured by TT (Fig. S8, Tables S10, S11). In all other cases, divergence substantially predated 19^th^-20th century persecution, indicating that present-day structure primarily reflects ancient post-glacial biogeographic histories rather than recent fragmentation. These populations have therefore been isolated for millennia with minimal evidence of gene flow among them (Figs. 1B, S1-S2), except for the Scandinavian population which receives occasional immigrants from Karelia (Fig. S1-S2). The southern populations (i.e., Dinaric-Balkan, Italian Peninsula, Iberian) cannot currently be considered a metapopulation for conservation purposes.

### Genetic diversity and demographic declines vary across European populations

Genome-wide heterozygosity showed pronounced differences among populations. Asian wolves exhibited the highest overall levels, with Tibetan wolves somewhat lower than other Asian groups (Fig. S9). Within Europe, Karelian and Dinaric-Balkan wolves had the highest mean heterozygosity (including per-site heterozygosity and mean π; Fig. 2A, Tables S12, S13). Italian Peninsula wolves presented uniformly low diversity, while Iberian and Scandinavian wolves showed reduced heterozygosity and elevated inter-individual variance, consistent with historical bottlenecks, recent founder events in Scandinavia (*24*), and/or cryptic fragmentation in Iberia (*14*, *25*). Demographic reconstructions indicated strong recent bottlenecks in all five European populations, with current Ne values well below conservation thresholds (Fig. 3). In the Late Medieval period, Karelian and Dinaric-Balkan populations exceeded Ne = 4,000 and the Iberian and Italian Peninsula populations were around Ne = 2,000 in the early part of the modeled timeframe. Italy was the first to fall below Ne = 500, approximately two centuries ago; the other European populations did not decline to these levels until nearly the 20th century (Fig. 3). The Scandinavian population trajectory was distinctive, with Ne near zero until ∼20-30 years ago, reflecting its recent founding from very few individuals.

**Fig. 2.**
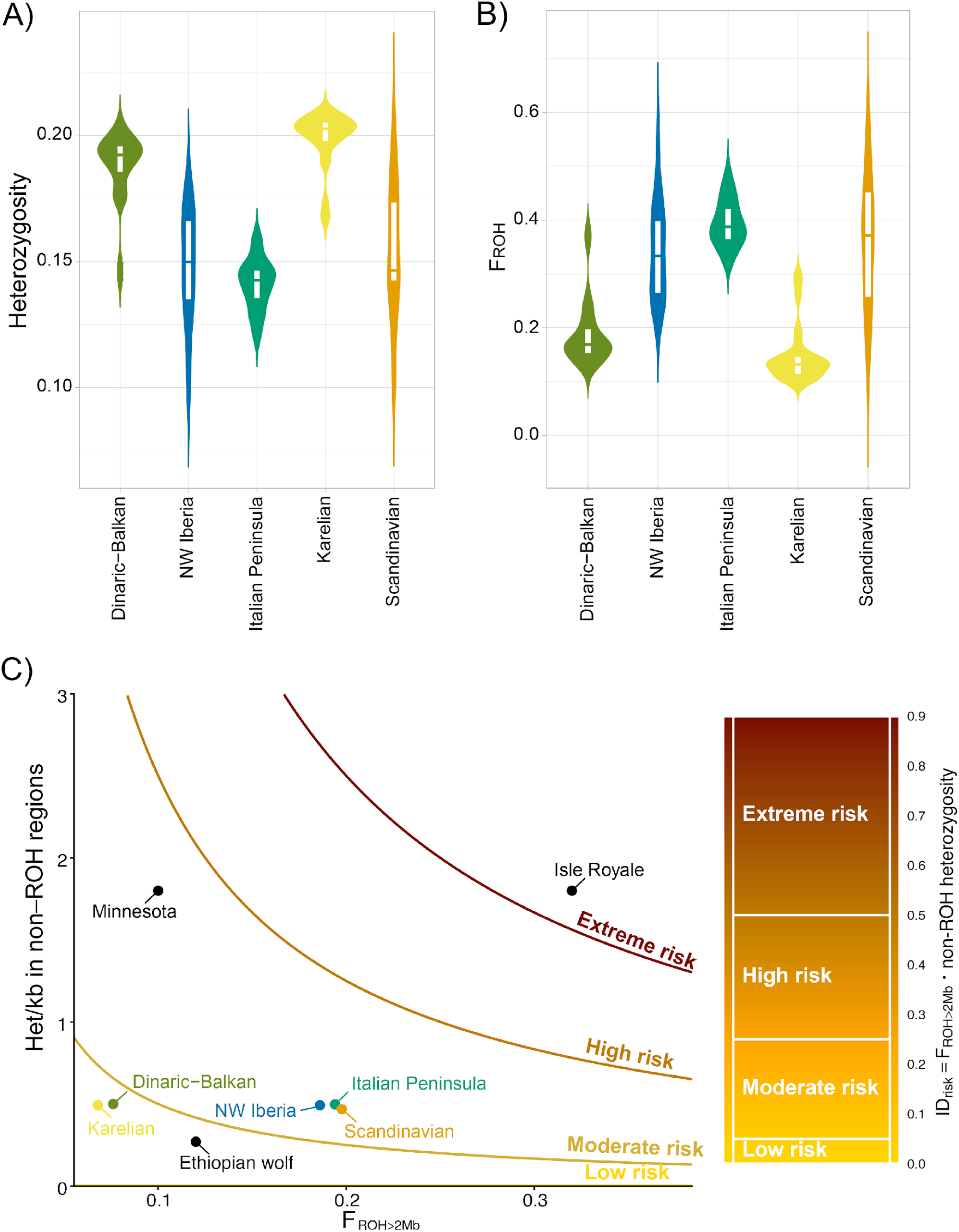
Homozygosity and genetic load in European gray wolf populations. (A) Variance in genome-wide heterozygosity per population. (B) Inbreeding coefficient based on runs of homozygosity (FROH) per population. (C) Inbreeding depression risk assessment (IDrisk), based on heterozygosity and FROH, a genomically based prediction of risk that a population suffers from inbreeding depression (*39*). Highly inbred Isle Royale and Minnesota wolves as well as the endangered Ethiopian wolves (C. simensis) are included for comparison.

**Fig. 3.**
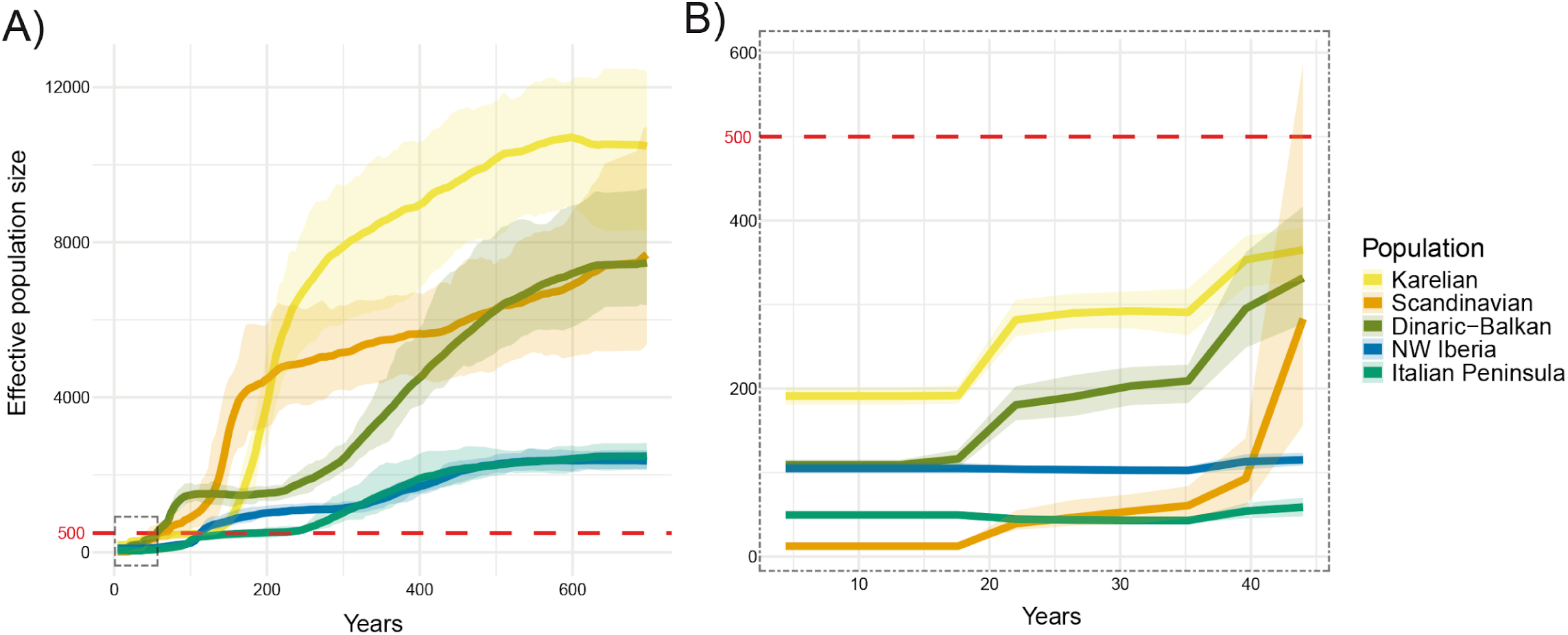
Demographic inference of gray wolf populations. (A) Effective population size (Ne) in the last 600 years (corresponding to about 150 generations) estimated by GONE2 and (B) zooming in on the last 45 years (dotted box in panel A). Lines represent geometric means and shaded ribbons denote 95% confidence intervals across 20 independent replicates. The red dashed line shows the Ne=500 threshold. All populations show a strong reduction in effective population size in the last 100-300 years. More ancient demographic reconstructions were made with Stairway plots, SMC++ and PSMC (Figs. S6, S7, S14).

### No European wolf population meets long-term viability thresholds, Ne > 500

Contemporary Ne estimates based on a linkage disequilibrium-based method, GONE2 (*26*), converged on very low values, all far below the Ne = 500 threshold for long-term adaptive potential. The lowest Ne was estimated for Scandinavia at 12.4 (95% CI: 11.7-13.1); Italy fell below the Ne = 50 threshold associated with severe short-term risk of inbreeding depression at 49.5 (45.1-54.2). The remaining populations exceeded Ne = 50 but were well below 500: Iberian 105.0 (99.3-111.6), Dinaric-Balkan 109.3 (101.3-115.2), and Karelia 191.0 (180.9-201.3).

Unaccounted within-population substructure may bias these estimates. Analyses correcting for this effect revealed a modest increase in Ne over recent generations, consistent with ongoing demographic recovery (Fig. S10). Estimates of subpopulation number varied widely and should be interpreted cautiously, as they may reflect pack-level genealogical structure rather than discrete demographic units (Table S14). Critically, all populations remained clearly below the Ne = 500 threshold regardless of analytical approach (Fig. S10; Table S14). Although census numbers have increased in all regions, these recoveries have not yet translated into effective population sizes compatible with favorable conservation status.

### High inbreeding and significant genetic load

Genomic inbreeding coefficients derived from runs of homozygosity (F_ROH_) mirrored heterozygosity and demographic patterns. The populations with the highest inbreeding – Italy (0.394 ± 0.04), Scandinavia (0.359 ± 0.12) and Iberia (0.343 ± 0.09) – are precisely those with the lowest Ne. The Scandinavian and Iberian populations showed particularly high inter-individual variance in both metrics (Fig. 2A, 2B), suggesting that these populations are not panmictic; in Scandinavia this likely reflects recent gene flow, while in Iberia it can be attributed to previously documented population structure (*25*). The length distribution of ROH revealed distinct inbreeding histories (Table S15, Fig. S11, S12). Italian Peninsula wolves carried the largest total length of short and intermediate ROH segments (mean ≈ 0.288 Gb and 0.153 Gb, respectively), consistent with a long-term bottleneck and isolation spanning 38-372 generations. Scandinavian wolves, in contrast, carried the greatest burden of long ROH (mean ≈ 0.435 Gb), a hallmark of intense, very recent inbreeding over fewer than ∼18 generations. Italian Peninsula and Iberian wolves also exhibited substantial long ROH reflecting both historical isolation and recent demographic decline (Italian Peninsula mean ≈ 0.428 Gb; Iberian mean ≈ 0.409 Gb).

Average heterozygosity outside ROH was relatively similar across populations, indicating that most standing diversity is maintained in non-inbred genomic regions. The proportion of the genome in long ROH (>2 Mb) differed sharply across populations and, as elevated F_ROH_ is precisely what drives the shift of genetic load from masked to realized (*27*), largely determined inbreeding depression risk categories (IDrisk, Fig. 2C). Italian Peninsula, Scandinavian and Iberian wolves fell in the “moderate” inbreeding depression risk category – notable given that inbreeding depression has already been documented in the one population where data exist to test for it (Scandinavia; *17*, *18*). Total genetic load (masked + realized) was greatest in Karelian wolves, yet the critical difference among populations lay in the ratio of masked to realized load (Fig. S13, Table S16, S17). In large, historically connected populations, deleterious variants are mostly masked and exert limited fitness effects. In Italy, Iberia and Dinaric-Balkan, prolonged isolation and recent bottlenecks have shifted much of this load into the realized category (Fig. 4), exposing recessive deleterious alleles and increasing risk of inbreeding depression. Scandinavian wolves have been shown to suffer inbreeding depression (17,18), which was acute in individuals with higher inbreeding coefficients (F_ROH_; Fig. 2B). The estimated realized genetic load of Iberian and Italian Peninsula wolves exceeded that of Scandinavian wolves, suggesting inbreeding depression is likely worse in those populations, though no empirical data currently exist to evaluate this prediction.

**Fig. 4.**
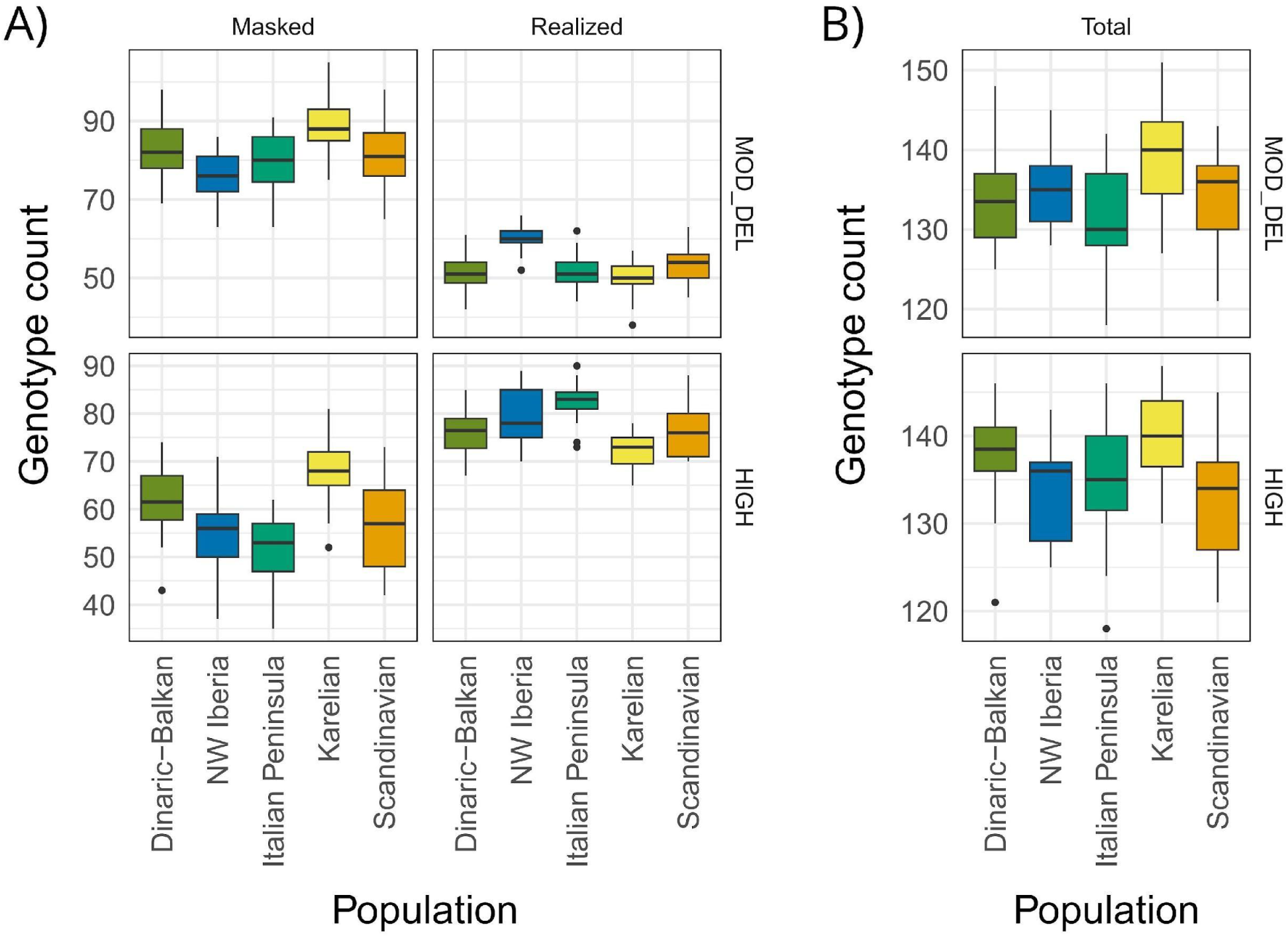
Genetic load estimates in European gray wolf populations across functionally deleterious classes (i.e., MOD_DEL = moderate deleteriousness; HIGH = high deleteriousness). (A) Masked and realized genetic loads across populations. (B) Total genetic load, sum of masked and realized loads. Genetic load estimates are based on the number of homozygous (realized load) and heterozygous (masked) loci carrying deleterious variants.

### Census counts mask the genetic fragility of recovering wolves

Our results challenge the legal framework which considers European wolves as a single unit, where numbers and recovery are typically reported at the continental scale (*1*, *8*). We show that the five analyzed European populations are highly differentiated, independently evolving lineages whose divergence dates to the late Pleistocene, except for the recently founded Scandinavian population. All five exhibit clear signatures of historical and recent bottlenecks - low genetic diversity relative to Asian wolves, steep declines in Ne over the past 100-200 generations, and extensive ROH indicative of persistent inbreeding. Despite demographic increase in census size and range, genomic recovery has not followed: genetic health in these populations remains compromised. Effective population sizes remain far below Ne = 500, and two populations hover near or below the Ne = 50 boundary associated with severe short-term inbreeding risks. Karelia is the least compromised population but is still well below Ne = 500. By the genetic criteria increasingly advocated for Favorable Conservation Status under EU policy (*5*), none of them can be considered secure in the long-term.

### Legal protection can drive natural recolonization

Wolves have comparable histories of range contraction and demographic decline to other widespread European large carnivores (*28–30*). Unlike brown bears (*30*), wildcats (*31*) and Eurasian lynx (*32*), however, the recent range recovery of wolves in Europe has been largely natural, facilitated by legal protection and ecological versatility rather than intensive reintroduction and conservation programs (*33*). Widespread natural recolonization demonstrates the power of legal protection, but this recovery has not yet progressed sufficiently to achieve genomic health. Wolves are continuing to recolonize where legal protection holds and appear to have the potential to meet conservation goals without the expensive interventions required for some other carnivores. Legal protection of wolves may therefore be sufficient to secure at least some European populations - but only if it is not withdrawn before populations reach healthy sizes and connectivity.

### Perceived overabundance masks genomic vulnerability

Growing human-wolf conflicts in recently recolonized areas, where wolves come into contact with farming communities with limited recent experience of coexisting with wolves, may have led to a biased general perception of overabundance of wolves, partly motivating the recent EU decision to relax protection. Our genomic data reveal that mere numeric abundance masks substantial genetic vulnerability. Policies that encourage increased lethal control or otherwise reduce census sizes risk further depressing already low Ne, exacerbating inbreeding, eroding adaptive potential and increasing realized genetic load – especially in already compromised populations. The recent extinction of the Sierra Morena population in Spain underscores how isolated, genetically eroded lineages can rapidly spiral toward extinction when conservation measures are relaxed or delayed (*19*, *34*).

### Genomic data can better inform conservation strategies

Conservation and management must therefore acknowledge that European wolves are not a single homogeneous unit but a set of distinct evolutionary lineages with different histories and vulnerabilities. At minimum, the five populations identified here – Karelian, Scandinavian, Dinaric-Balkan, Italian Peninsula and Iberian – should be treated as separate management units. For each, strategies should aim to increase Ne through demographic growth and to facilitate connectivity where feasible in order to reduce inbreeding and increase gene flow. Encouragingly, recent long-distance dispersal events (*35–37*) indicate that legal protection is already facilitating the natural re-establishment of connectivity among previously isolated lineages. Maintaining and strengthening these protections is essential if genetic recovery is to eventually follow the demographic gains documented so far. More broadly, the European wolf exemplifies a critical conservation pivot: demographic recovery is not synonymous with genetic recovery. As global and regional frameworks move toward integrating genetic indicators into conservation targets to ensure long-term adaptability and evolutionary potential, increasingly available whole-genome data provide the necessary resolution to detect hidden risks beneath positive demographic trends. The evolutionary future of European wolves – and the ecological functions they provide – will depend on policies informed not only by trends in counts and ranges, but by the genomic evidence now available. The message in the data is unambiguous: European wolves have returned, but they are not yet safe.

## Acknowledgments

The authors thank all the people who helped with the fieldwork and allowed the collection of the samples sequenced and analysed in this study. We are grateful to the regional governments of Asturias, Cantabria, Galicia, Castilla-La Mancha in Spain for providing samples. Slovenian samples were collected by the Slovenian Forest Service within the scope of national monitoring of large carnivores. We particularly thank Matej Bartol and Rok Černe for their diligent collection of genetic samples of wolves in Slovenia. For Turkish samples, the animal study was approved by Türkiye’s Department of Nature Conservation and National Parks and the Ministry of Agriculture and Forestry for granting the permit for this research (No. 72784983-488.04-114100 and E-21264211-288.04-1602322). All animal procedures followed the Kafkas University local ethical committee for animal experimentation (KAÜ HADYEK) guidelines under KAÜ HADYEK/ 2018-050 and KAÜ-HADYEK/ 2021-083 permission numbers. The study was conducted in accordance with the local legislation and institutional requirements. We thank the laboratory facilities: Conservation Genomics Research Unit of the Fondazione Edmund Mach (FEM), Molecular Ecology Lab of the Doñana Biological Station (LEM-EBD) and Sakarya University Molecular Biology and Genetic Laboratory. We are grateful to the IT team at the University of Ljubljana for providing access to their computing server and for their technical support, with special thanks to Elena Pazhenkova for her assistance throughout this project. CS gratefully acknowledges Prof. Bridgett vonHoldt and the High-Performance Computing (HPC) Research Center at Princeton University, USA, for providing support and computational resources for analyses.

## Author contributions

Conceptualization: SR, CV, DB, JAL, IS, CVA, PC, RC, EF

Resources: PC, RG, RC, EF, TS, AVS, CV, CVA, MN, JAL, CN, IS, DL, FM, EV, MG, JK, CHS

Data curation: SR, DL

Formal analysis: SR, DB, IS, CS, CL, FIS

Funding acquisition: PC, RG, RC, EF, TS, AVS, CV, CVA, MN, JAL, CN Visualization: SR, DB, IS

Writing – original draft: SR, CV, JAL, CVA, DB, IS Writing – review & editing: all authors

## Data, code, and materials availability

Raw sequencing reads for 78 wolves (including all individuals used for the analyses presented in this paper) will be deposited in ENA under this accession number: PRJEB110322. Additional 27 genomes from wolf-dog admixed individuals and 8 village dogs, which were used solely for initial quality filtering and are not part of the final dataset, will be made available with subsequent publications within the Biodiversa+ WOLFNESS project and can be requested directly to the authors if needed.

## Supplementary Materials

Materials and Methods

Figs. S1 to S14

Tables S1 to S17

References (*40–78*)

