## Supplementary Materials for "Misleading Success: Genomes Reveal Critical Risks to European Gray Wolves"

**The PDF file includes:**

Materials and Methods
Figs. S1 to S14
References (40-78)

**Other Supplementary Materials for this manuscript include the following:**

Tables S1 to S17

### Materials and Methods

Sampling and molecular methods

Wolf tissue samples were collected between 2002-2023 in different localities (Table S1). All samples derived from wolves dead or captured for reasons other than this study and were stored by regional or national governments or academic institutions. Sample selection was conducted within the framework of the Biodiversa+ WOLFNESS project, where potentially admixed individuals were specifically targeted to assess the genomic consequences of hybridization between wolves and dogs. Non-admixed wolves were also sequenced to be used as reference. Muscular tissue was stored in 50 ml of 95% ethanol at −20 °C and blood was collected and preserved in EDTA solution at − 20 °C until processing in the lab. DNA extraction from tissue fragments or blood (~25 mg) was done using the DNeasy Blood and Tissue Kit (Qiagen) with an overnight digestion at 56 °C. Elution was carried out at the different laboratories (Conservation Genomics Research Unit of the Fondazione Edmund Mach FEM, Italy; Molecular Ecology Lab of the Doñana Biological Station LEM-EBD, Spain; Sakarya University Molecular Biology and Genetic Laboratory, Türkiye; CIBIO molecular facilities CIBIO-BIOPOLIS, Portugal; Dpt. of Biology, Biotechnical Faculty, University of Ljubljana, Slovenia; Senckenberg Research Institute, Centre for Wildlife Genetics, Germany) using two sequential washes of 50 µL AE buffer, each preceded by a 10-minute incubation at 37 °C. Prior to elution, samples were stored at −20 °C in the DNeasy Mini spin columns. Following dilution to a concentration of 100 ng/ul, DNA extracts were shipped to Novogene facilities (Cambridge, UK), where they were stored at -20ºC until processing. Sample quality control was checked through an electrophoresis 1% agarose gel and using Qubit 2.0 to detect potential contamination or degradation of DNA and accurately quantify the concentration of double-stranded DNA (Life Technologies, CA, USA). Libraries were constructed by Novogene using a PCR-free preparation protocol. Summarising, the extracted DNA was fragmented to an average length of 350 bp using the Covaris LE220 Plus focused-ultrasonicator (Covaris, Massachusetts, USA), fragments ends were blunted using enzymes with exonuclease/polymerase activity. Following an adenylation of the 3' ends of the DNA fragments and the ligation of adapters, products were purified using the AMPure XP system (Beckman Coulter, Beverly, USA). Libraries concentrations were quantified using Qubit and quantitative PCR (qPCR), and fragment size was checked using the NGSK3 High Sensitivity DNA Assay with the Perkin Elmer LabChip instrument. Finally, libraries were paired-end sequenced for a 150 bp fragment-length with Illumina's NovaSeq X Plus technology at Novogene facilities.

In addition to our 105 newly sequenced wolves, eight Italian village dogs, and three Turkish dogs, we incorporated publicly available whole-genome data for 176 wolves from Europe (93), Asia (50) and North America (33), 144 purebred dogs belonging to 74 medium and large sized breeds, 42 village dogs from Asia, Africa and Europe, and three canid outgroups (African wild dog Lycaon pictus, dhole Cuon alpinus and culpeo Lycalopex culpaeus). Sample metadata—including GenBank accession numbers and sampling localities—are provided in Table S1 and S2.

Alignment, variant calling and filtering

Raw sequencing data quality was initially assessed using FASTQC v0.12.1 (40) and the results were summarized with MultiQC v1.22.1 (41). Adapter removal and quality trimming were performed using FASTP v0.23.4 (42). Processed reads were then aligned to the complete CanFam3.1 dog reference genome (43; RefSeq assembly accession: GCF_000002285.3) using BWA-MEM v0.7.17-r1188 (44). We performed read group tagging and duplicate marking with PICARD v3.2.0 (Broad Institute; https://broadinstitute.github.io/picard/).

We generated gVCF files using GATK v4.5.0.0 (45) by running HaplotypeCaller, GenomicsDBImport, and GenotypeGVCFs for each chromosome, and then concatenated the results using GATK's GatherVcfs function. Hard filtering was applied following the GATK Best Practice Workflow (alternative protocol 2) with the following thresholds: QD < 2.0, FS > 60.0, MQ < 40.0, MQRankSum < –12.5, SOR > 3.0, and ReadPosRankSum < –8.0. These filters were implemented using VariantFiltration to mark variants and SelectVariant to apply the filters, retaining only SNPs. Subsequently, we used VCFTOOLS v0.1.16 (46) to further filter variants by excluding those with a minor allele frequency below 1%, missingness above 10%, a quality score less than 30, and a depth lower than 5×. We excluded samples with a mean coverage lower than 10X and more than 10% missing data.

Linkage disequilibrium (LD) pruning was performed in PLINK v2.00a5.12LM (47) using a 10 kb sliding window, 5 bp step size, and an $r^2$ threshold of 0.5. To estimate the global proportion of the genome derived from dog ancestry, we conducted admixture analyses using ADMIXTURE v1.3.0 (48) and assuming 2 different populations (K = 2). We ran these analyses separately for European, Asian and North American wolves to take into account their genetic differences, which could bias the results focused only on the introgression with dogs. We ran 15 independent iterations using default parameters and combined the results from the different runs using the CLUMPAK (Cluster Markov Packager Across K) server with default settings (49). We retained only wolf samples that showed a global dog ancestry lower than 5%.

To avoid bias from close kinship, we estimated pairwise relatedness in wolves with NgsRelate2 (50) and excluded individuals exceeding KING-robust kinship ≥ 0.20, $R0 \leq 0.10$, and $R1 \geq 0.50$ (51).

Population structure

We explored population structure by performing a principal components analysis (PCA) on autosomal SNPs in PLINK and visualized the first two principal components in R v4.4.2 using "ggplot2" (52). We assessed genetic structure across gray wolves with ADMIXTURE at K values from 2 to 10, with 15 independent replicates per K. We ran ADMIXTURE twice: first on the full dataset including all wolf populations, and then on European wolf populations only. In both cases, the optimal K was selected by minimizing cross-validation error, and results across replicates were summarized using CLUMPAK under default settings (49) and visualized with pong (53).

Phylogenetic analyses

We reconstructed phylogenetic relationships among wolves using both population-level and individual-level approaches. All analyses included North American, Asian, and European gray wolves, with African wild dog, dhole, and culpeo as outgroups.

At the population level, we inferred a maximum-likelihood (ML) tree using TreeMix v1.13 (54), which models relationships among populations based on allele-frequency covariance. We first converted our VCF to PLINK format and then generated stratified allele-frequency files per population. These were reformatted for TreeMix using a custom Python script (adapted from Nelson's plink2treemix.py; https://github.com/thomnelson/tools). TreeMix v1.13 was run with African wild dog as outgroup, grouping SNPs into blocks of 1,000 and performing 100 bootstrap replicates. We explored models with zero to five migration edges to evaluate the presence and directionality of gene flow among populations. No migration edges among European wolf populations were supported, so we present the tree without migration.

At the individual level, we used two complementary approaches. First, we built a neighbour-joining (NJ) tree based on pairwise identity-by-state (IBS) distances computed with PLINK v1.9 (commands "--distance square ibs" and "allele-ct"). Using the resulting distance matrix as input, we built and plotted the NJ tree in R, using African wild dog, dhole, and culpeo as outgroups, with the packages "ape" (55) and "ggtree" (56). To ensure that using different outgroups did not bias the topology, we repeated the NJ analysis separately with each outgroup in turn; topologies were concordant, so we retained the full three-outgroup tree.

Second, we generated a NJ tree using fastreeR (57), which computes pairwise distances and builds the tree directly from a VCF. We used as input the VCF excluding related and admixed individuals and performed 100 bootstrap replicates to assess tree support. We plotted the resulting Newick tree in R, rooting it on the African wild dog, with the packages “ape” (55) and “ggtree” (56).

Demographic analyses

To reconstruct recent and ancient demographic trajectories in European wolf populations, we applied different complementary methods. To reconstruct recent changes in effective population size (Ne), we applied the linkage disequilibrium-based method implemented in GONE2 (26), which produces reliable Ne estimates over the last ~150 generations. We used the unpruned SNP dataset to retain linkage information, subsampling up to 50,000 high-quality SNPs per autosome. All GONE2 parameters were set to their defaults except for the mean recombination rate, which was set to 1.3459 cM/Mb (58). We ran 20 independent replicates per population to generate empirical confidence intervals and assumed a generation time of 4.4 years (59). To assess the potential influence of within-population substructure on Ne estimates, analyses were also run both with default settings and with the -x flag, which models the sample as a random draw from a metapopulation composed of subpopulations of equal size (26). Simulations showed that accounting for potential substructure may yield more reliable Ne estimates for very recent generations following a population size reduction, but can fail to reconstruct the reduction itself, which may instead appear as a gradual linear decrease (26).

To reconstruct older changes in Ne, we applied the coalescence-based method implemented in StairwayPlot2 (60), which infers demographic histories from site frequency spectra (SFS). We used the SNP dataset without minor allele filtering and separately for each population

(Dinaric-Balkan, Iberian, Italian, Karelian, and Scandinavian) including only non-related individuals. Subsequently, a folded SFS was obtained for each population using the vcf2sfs.py python script (https://github.com/marqueda/SFS-scripts/blob/master/foldSFS.py). We prepared blueprints for StairwayPlot2 using population-specific nseq, SFS and nrand parameters, and the following shared parameters: L = 2,500,000,000; whether_folded = true; ninput = 200; mu = 4.5e-9; year_per_generation = 4.4. StairwayPlot2 results were plotted together with GONE2 results (default settings) using the ggplot2 R package.

For more ancient demographic inference, we employed the Pairwise Sequentially Markovian Coalescent model (PSMC; 61). From each population we selected 4–5 individuals with mean genomic coverage > 20×, sampling across their geographic range. For each selected BAM, we generated autosomal consensus sequences by running samtools mpileup against the CanFam3.1 reference, excluding reads with excessive mismatches (-C 50), then called variants with bcftools call -c. We converted VCF to FASTQ with vcfutils.pl vcf2fq (62) while removing sites with depth < 5 or > 100. We then ran PSMC v0.6.5-r67 under three time‑interval schemes —"4+25*2+4+6", "2+2+25*2+4+6", and "1+1+1+1+25*2+4+6"— following previous studies on gray wolf and recent recommendations (63). After inspecting trajectories for spurious "false peaks," we selected the "2+2+25*2+4+6" time interval for our final analyses. We used a mutation rate of $4.5\times10^{-9}$ per site per generation (64) and the same 4.4 year generation time used in GONE2.

To provide an additional coalescence-based reconstruction of demographic history, we applied SMC++ (65), which extends the Sequentially Markovian Coalescent model by incorporating information from the full sample frequency spectrum of unphased genomes, enabling higher resolution in the recent past compared to PSMC. We used the filtered SNP dataset across all 38 autosomes. For each population (Dinaric, n = 27; Iberian, n = 17; Italian, n = 35; Karelian, n = 27; Scandinavian, n = 17), we converted the VCF into SMC++ input format using vcf2smc. Following the author's recommendation of using between 2 and 10 individuals per run (Terhorst, https://github.com/popgenmethods/smcpp), we selected six lineages per run and generated n − 5 composite likelihood replicates per population by sliding this window across the full sample, which were treated as bootstrap replicates to assess uncertainty in the estimated trajectories. We then ran SMC++ estimate on each replicate independently with the same per-generation mutation rate as above and default values for the thinning and timepoints parameters. Results were plotted using the ggplot2 R package assuming a generation time of 4.4 years, consistent with all other demographic analyses.

<u>Divergence time estimates</u>

Divergence times among European wolf populations were inferred using two complementary approaches: MiSTI (Migration-aware Split Time Inference; 22) and the Two-Two (TT) method (23, 66). While MiSTI estimates split times and post-divergence migration by fitting independent population size trajectories inferred from PSMC to a model of population separation with possible asymmetric gene flow, the TT method derives divergence parameters directly from the two-dimensional site frequency spectrum (2DSFS) computed between pairs of genomes, estimating branch-specific split times ($T_1$, $T_2$) and ancestral effective size ($N_a$) under a simple split model without migration.

For MiSTI, pairwise 2DSFS files were generated with ANGSD (67) and converted to MiSTI format using the ANGSDSFS.py utility. Joint pairwise time steps were identified with calc_time.py, and migration intervals were defined in different time bins according to Marine Isotope Stages (MIS) following (68) and (69), subdividing the Holocene into MIS1 (14–11.7 ka) and MIS0 (11.7–0 ka); adjacent time bins with fewer than two time steps were merged. MiSTI commands were parallelized with GNU parallel (70), and log-likelihood profiles across time steps were fitted with ninth-degree polynomials (retained if $R^2 > 0.97$), with the polynomial maximum taken as the most likely divergence time per individual pair.

The TT method was implemented following (71). Given the high sequencing coverage and high genotyping quality, genomic positions absent from the 2DSFS were assumed to represent homozygous reference genotypes (0/0_0/0) and the monomorphic site count was corrected accordingly; genotypes were otherwise categorized into the nine possible diploid combination bins (0/0_0/0, 0/0_0/1, … 1/1_1/1). These two methods provide complementary perspectives on population divergence: MiSTI integrates demographic history and migration, whereas the TT approach captures the signal of sequence differentiation from site-frequency information alone. Applying both allows for cross-validation of split-time estimates derived from distinct underlying assumptions and data structures. Both methods were applied to autosomal data. For each population pair, average divergence times were calculated across all individual comparisons, excluding negative branch estimates. For the TT method, negative values of $T_1$ and $T_2$ were excluded from the final calculation of divergence time, and when both metrics were positive their mean was used. Population-level estimates were summarized with 95% confidence intervals across all valid individual-level comparisons. Statistical comparisons between MiSTI and TT results were performed using Mann-Whitney U and Student's t-tests to assess potential differences in split-time distributions.

Intra-population genomic diversity

To compare the patterns of genomic variation among the five European wolf populations, we estimated the observed heterozygosity (Ho) and the nucleotide diversity ($\pi$). As the coverage is relatively high (>10X, mean 22.7X) for all 123 samples, we calculated both statistics using VCFTOOLS v. 0.1.16 (42). For nucleotide diversity, we applied a sliding window approach with a window size of 100 kbp and calculated the mean value across the genome. We estimated the mean weighted FST among all pairs of European populations using VCFTOOLS.

Inbreeding and runs of homozygosity

Runs of homozygosity (ROH), which represent putative identity-by-descent segments, were inferred only for European populations using PLINK with 100 kb sliding windows (16). We required each ROH to contain at least 100 SNPs at a density of ≥ 1 SNP per 50 kb. To accommodate genotyping errors, up to one heterozygous site and five missing calls were permitted per 1 Mb window within a ROH (72). Consecutive ROHs were defined as segments separated by at least 1 Mb. For each individual, we calculated the total length of all ROHs (SROH) and the total number of ROHs (NROH). We further partitioned SROH into three tract-length classes—short (100 kb–1 Mb), intermediate (1–2 Mb), and long (> 2 Mb)—to examine their respective contributions to genomic inbreeding. The inbreeding coefficient based on ROH (FROH) was then estimated as SROH divided by the total autosomal genome length covered by SNPs (2,201,821,400 bp), and plotted by population. We calculated ROHs coalescence time aiming to define the time at which the ROHs were likely formed. To do so, we used the formula L = 100/2t cM (73) where L is the length of the ROH, cM is the recombination rate, and t is the time of coalescence in generations.

To quantify the genomic consequences of inbreeding, we estimated the inbreeding depression statistic (ID), as defined by (39). This index combines the genomic burden of long runs of homozygosity (longROH) with residual heterozygosity outside ROH regions (nonROH het/kb), capturing both recent inbreeding and the remaining adaptive potential of the genome. For each individual, we computed: ID=longROH (kbp) × nonROH (het/kbp)

Individual ID values were averaged within each population using arithmetic means, as in (39), to obtain population-level estimates. We then compared these values against the quantitative extinction-risk thresholds defined by (39). To contextualize our results, we included the three reference Canis populations analyzed in that study — the Ethiopian wolf (Canis simensis), Minnesota gray wolf (Canis lupus), and Isle Royale wolf (Canis lupus) — representing low, intermediate, and high extinction-risk categories, respectively.

Genetic load

We used publicly available short read data from three outgroups, Lycaon pictus (African wild dog, SRA accession: SAMN09924608), Canis latrans (Coyote, SRA accession: SRR7107770), and Canis aureus (Golden Jackal, SRA accession: SRR7976426), and mapped the sequences to the dog reference genome using the same procedure as described above, keeping only sites covered with at least five reads per outgroup. To avoid ascertainment bias towards the dog reference allele, we did not use called genotypes for the outgroups but instead pseudohaploidized the genomes by randomly drawing one allele for each species using the read coverage as weight. This was done for each filtered variant site from above using custom python scripts adapted to the pipeline of (74). The ancestral state of polymorphisms segregating in wolves was inferred for all sites where the outgroups agreed on one of the two alleles present in the wolf data set. The ancestral alleles were added as AA tags to the INFO field of the vcf file using a custom bash script. For all sites considered to contain deleterious variation, the derived allele was assumed to be the deleterious allele. All analyses were based on polarized mutations only.

We used the domestic dog genome annotation gtf, cDS and protein files (CanFam3.1; Ensembl release 104 https://may2021.archive.ensembl.org/Canis_lupus_familiaris/Info/Index) to build custom snpEff v.4.3.183 (75) and SIFT4G v.6.084 (76) databases with default settings. We then annotated and predicted the effects of variants using the aforementioned tools. We classified putatively deleterious variants into three categories: 1) Low impact (LOW) variants likely to be not deleterious (i.e. synonymous), 2) Moderate impact (MODERATE) variants likely to modify protein effectiveness (i.e. nonsynonymous), and 3) High impact (HIGH) variants likely to disrupt protein function (i.e. loss of function LoF, stop codons, splice donor variant and splice acceptor, or start codon lost) (77). We used SIFT (Sorting Intolerant From Tolerant) score to discriminate MODERATE in tolerated nonsynonymous (SIFT score ≥ 0.05) (MOD_TOL) and putatively deleterious nonsynonymous (SIFT score < 0.05) (MOD_DEL) variants. Then we used a custom script to estimate individual allele frequencies and genotype counts of LOW, MOD_TOL, MOD_DEL, and HIGH-impact variant-derived alleles, for each wolf in the dataset. In particular, the count of heterozygous genotypes represented the "masked" load, which quantifies the potential loss of fitness due to (partially) recessive deleterious mutations that may become expressed in future generations. The count of homozygous genotypes for the derived alleles represented the “realized” load, which reduces fitness in the current generation; the sum of “masked” and “realized” load represented the “total” load (78). Statistical comparisons between genetic load counts among European wolf population pairs were performed using the Wilcoxon test to assess potential differences at deleterious variants (i.e., MOD_DEL, HIGH).

A)

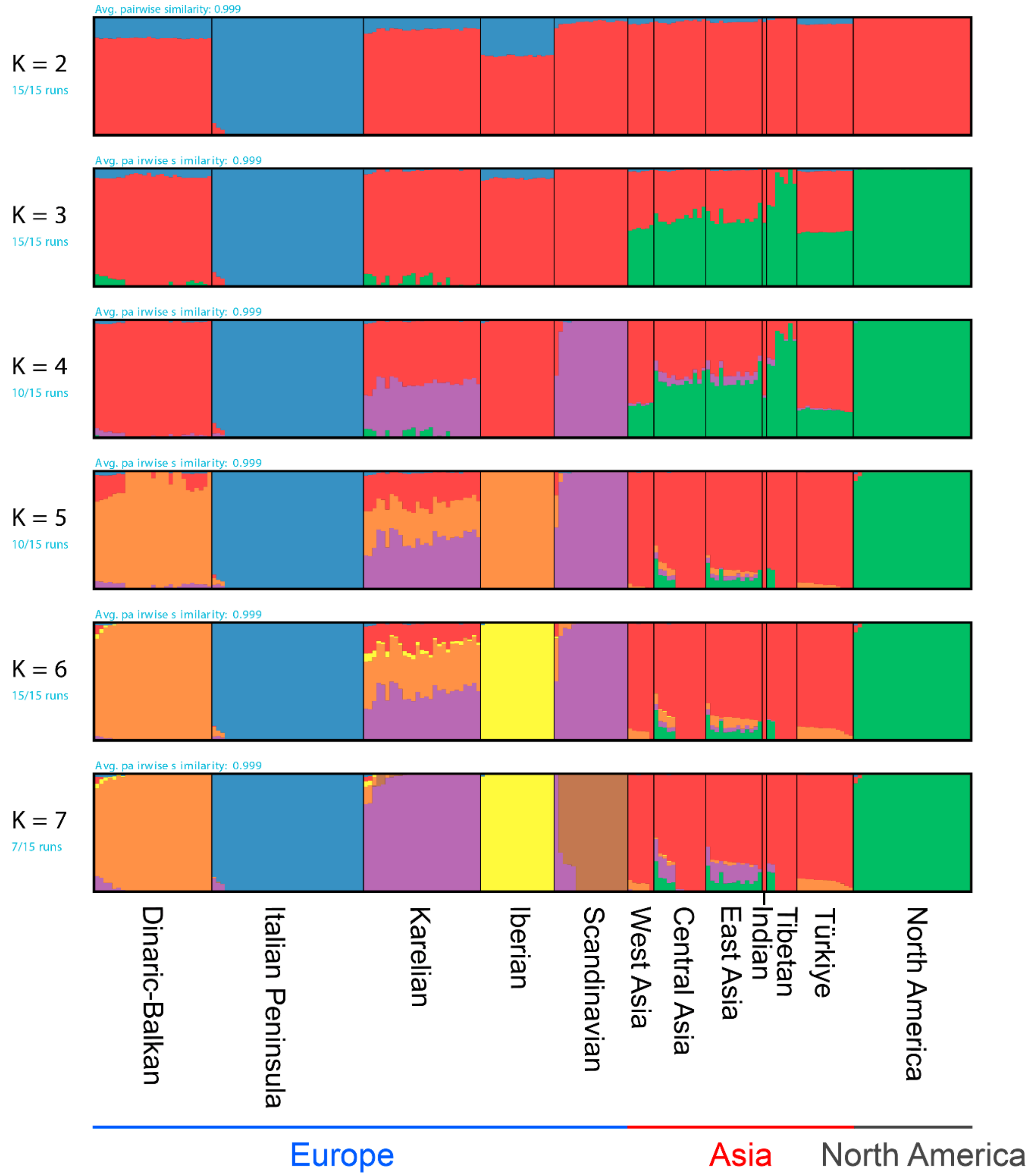

Avg. pairwise similarity: 0.999
K = 2
15/15 runs
Avg. pa irwise s imilarity: 0.999
K = 3
15/15 runs
Avg. pa irwise s imilarity: 0.999
K = 4
10/15 runs
Avg. pa irwise s imilarity: 0.999
K = 5
10/15 runs
Avg. pa irwise s imilarity: 0.999
K = 6
15/15 runs
Avg. pa irwise s imilarity: 0.999
K = 7
7/15 runs
Dinaric-Balkan
Italian Peninsula
Karelian
Iberian
Scandinavian
West Asia
Central Asia
East Asia
Indian
Tibetan
Türkiye
North America
Europe
Asia
North America


B)

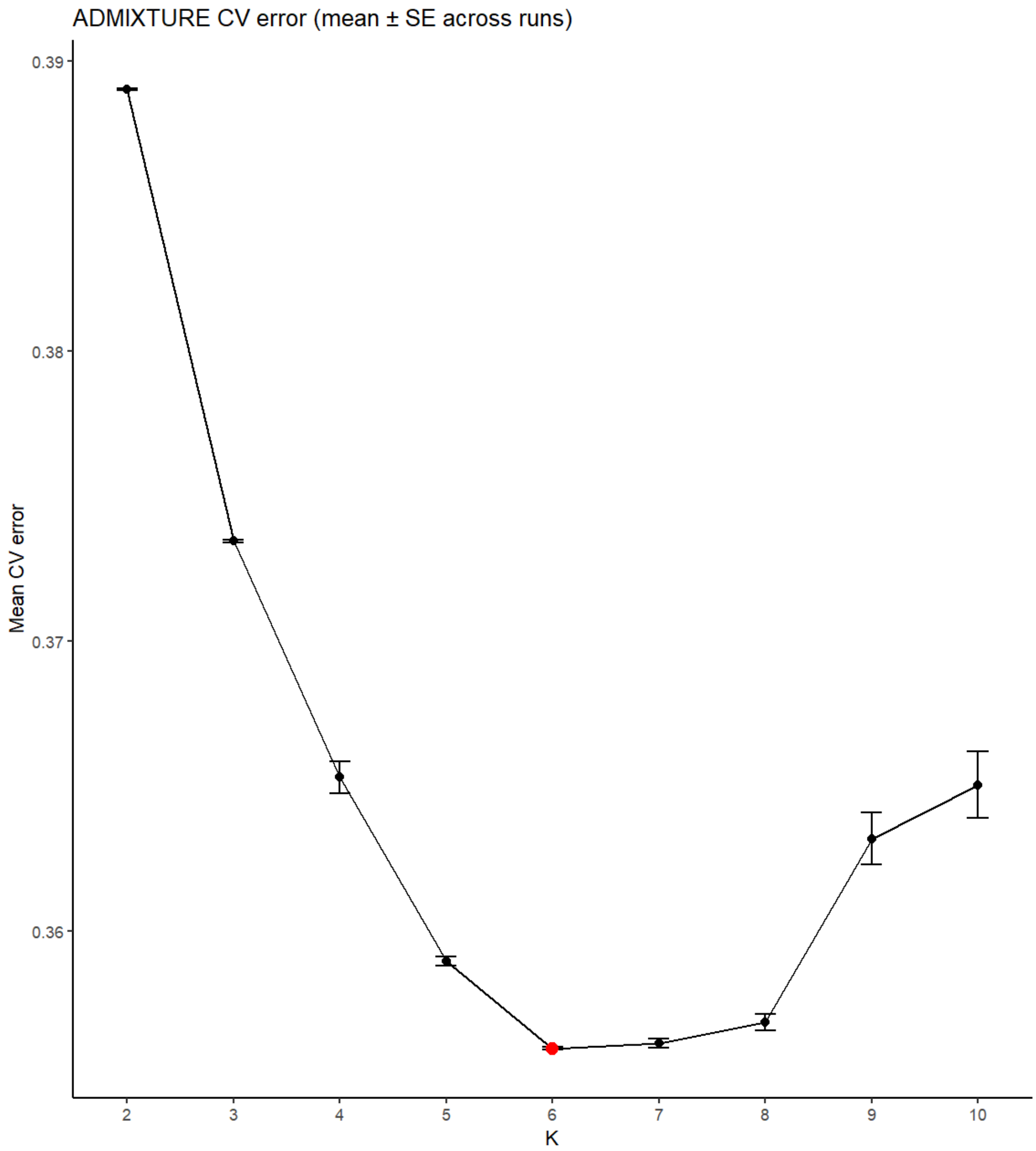


**Fig. S1.**

Population structure inferred by ADMIXTURE for wolf populations across continents. (A) Ancestry proportions for all wolf populations at K = 2-7. Each vertical bar represents an individual, colored by estimated membership in K ancestral clusters. K = 6 is the optimal number of clusters. Most European populations appear well differentiated, except for the Karelian population, which shows mixed ancestry from Dinaric-Balkan, Scandinavian, and Asian wolves. (B) Cross-validation error across K values from 2 to 10 (mean and standard errors across 15 independent replicates), identifying K = 6 as the value that best explains the data.

**A)**

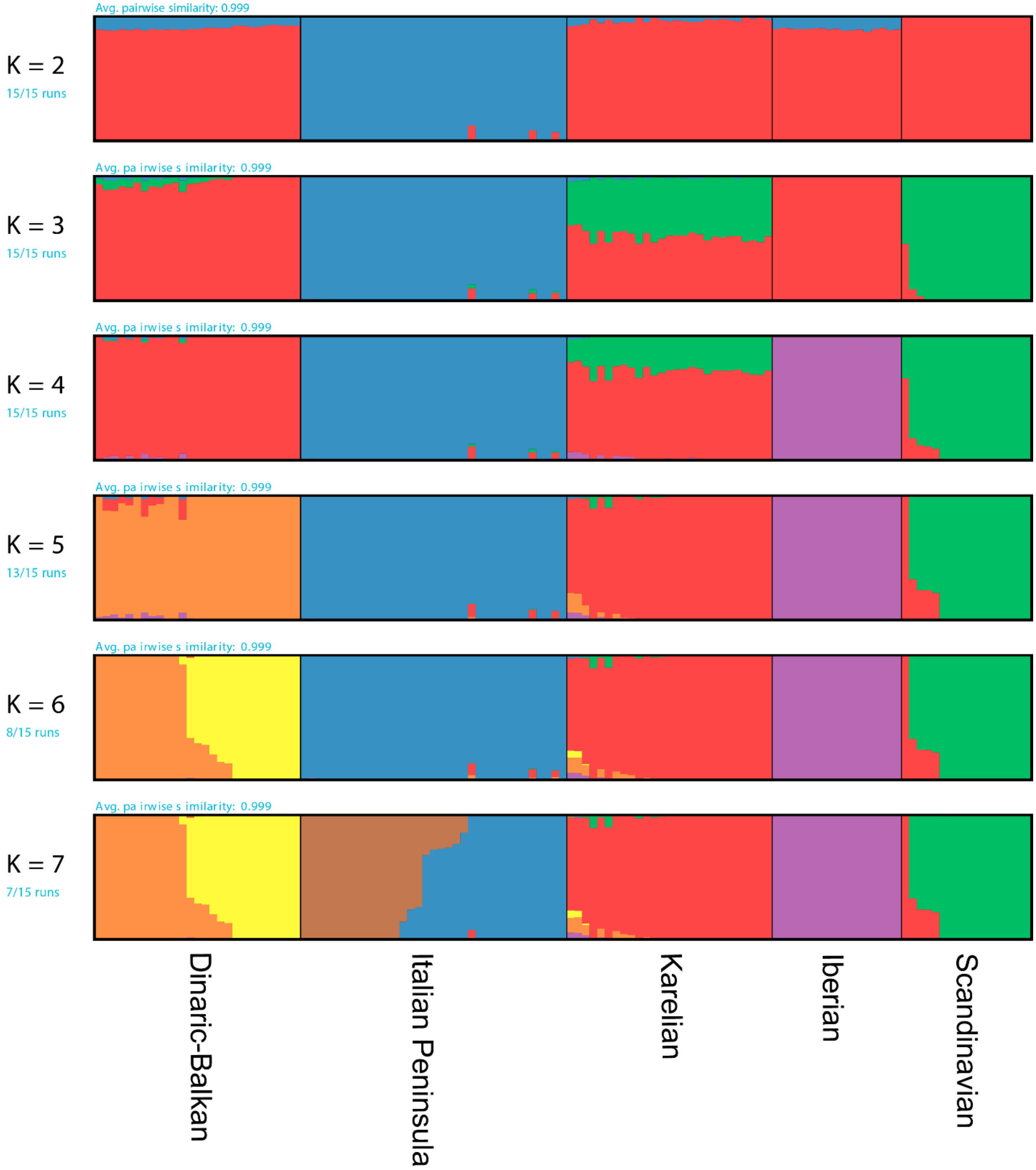
Avg. pairwise similarity: 0.999
K = 2
15/15 runs
Avg. pa irwise s imilarity: 0.999
K = 3
15/15 runs
Avg. pa irwise s imilarity: 0.999
K = 4
15/15 runs
Avg. pa irwise s imilarity: 0.999
K = 5
13/15 runs
Avg. pa irwise s imilarity: 0.999
K = 6
8/15 runs
Avg. pa irwise s imilarity: 0.999
K = 7
7/15 runs
Dinaric-Balkan
Italian Peninsula
Karelian
Iberian
Scandinavian

**B)**

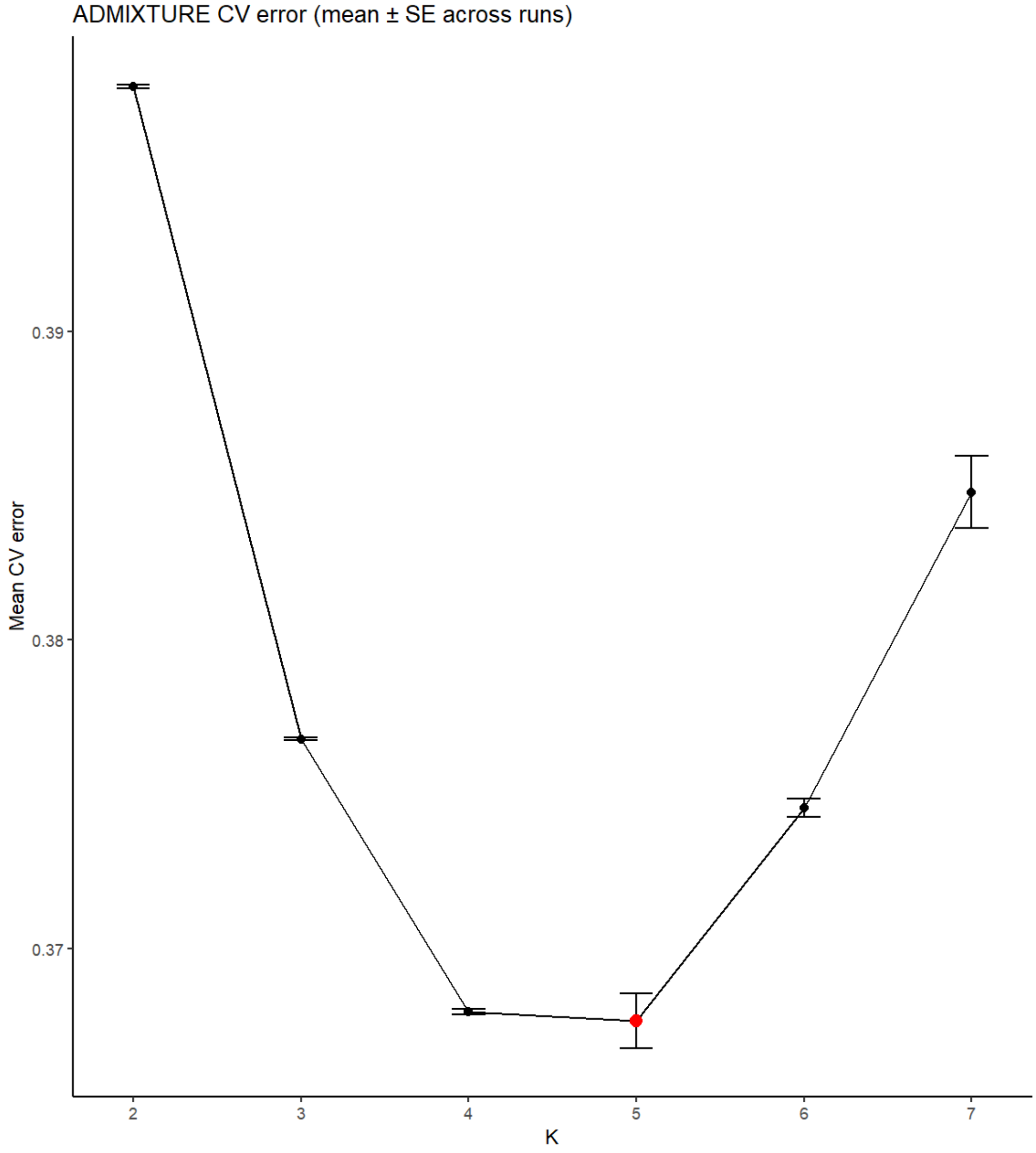


**Fig. S2.**

Population structure inferred by ADMIXTURE only for European wolf populations. (A) Ancestry proportions for European wolf populations at K = 2-7. Each vertical bar represents an individual, colored by estimated membership in K ancestral clusters. K = 5 is the optimal number of ancestral components. (B) Cross-validation error across K values from 2 to 7 (mean and standard errors across 15 independent replicates), identifying K = 5 as the value that best explains the data.

**A)**

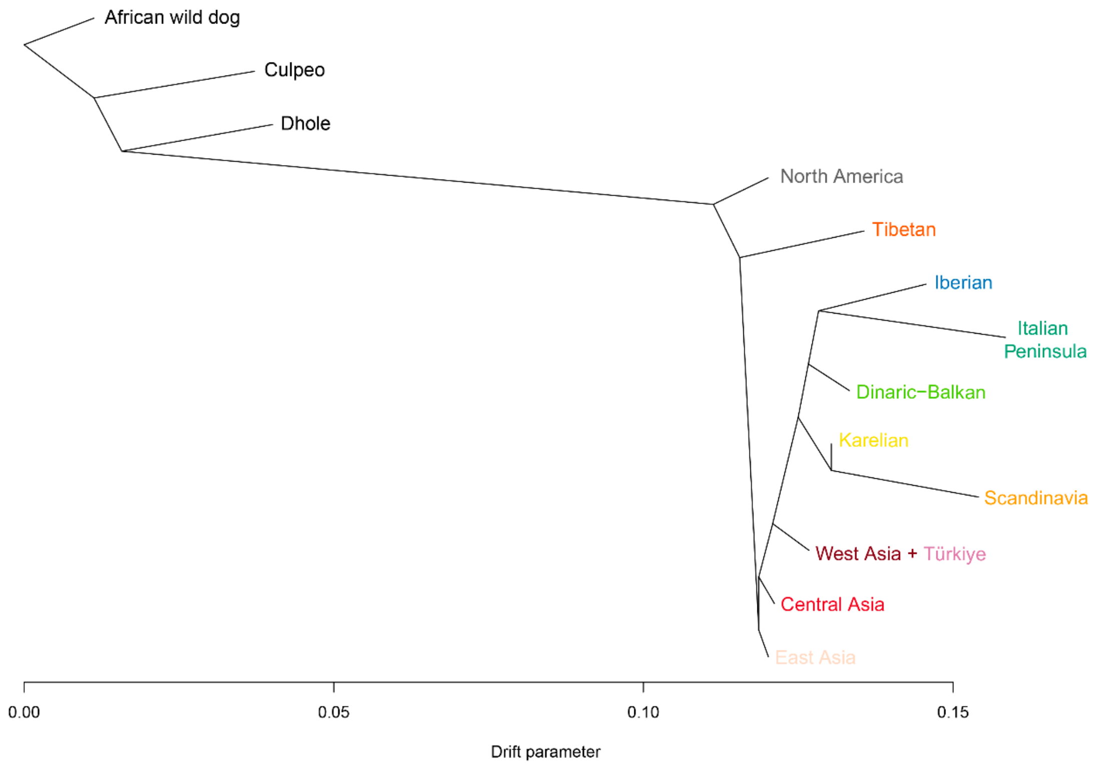

African wild dog
Culpeo
Dhole
North America
Tibetan
Iberian
Italian Peninsula
Dinaric–Balkan
Karelian
Scandinavia
West Asia + Türkiye
Central Asia
East Asia
0.00
0.05
0.10
0.15
Drift parameter

**B)**

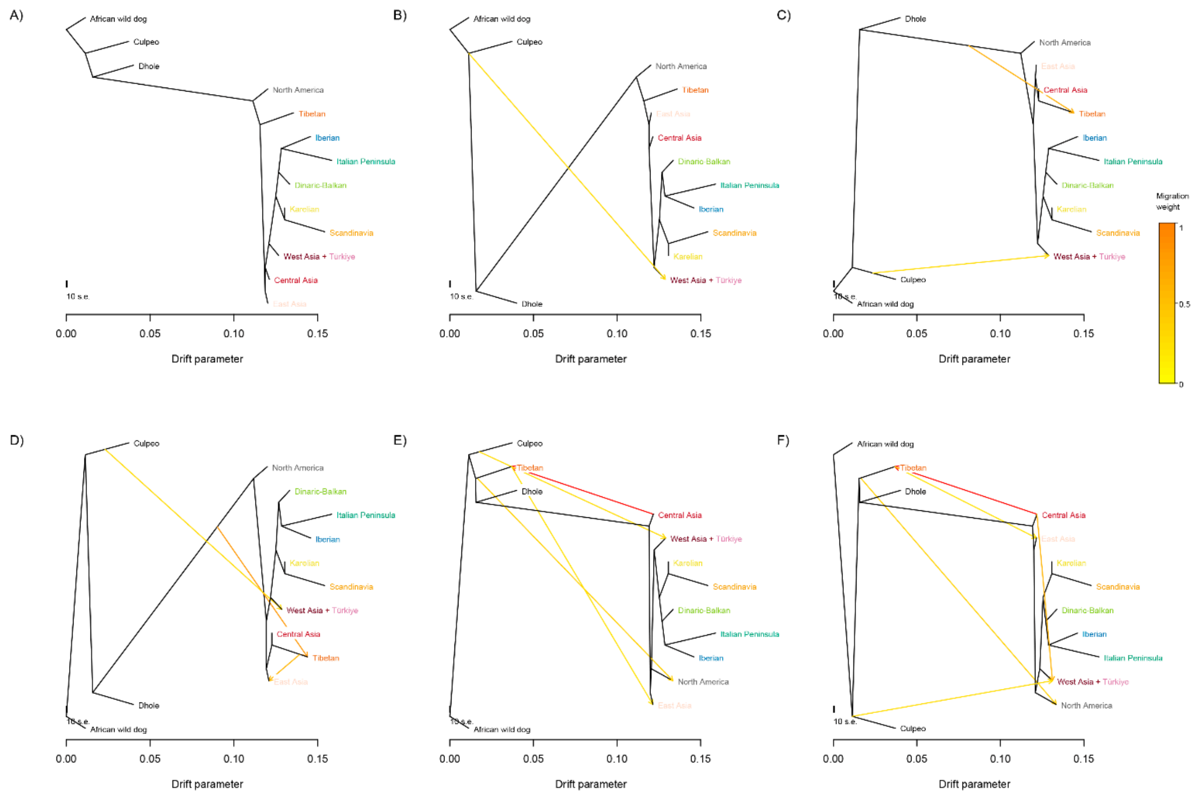


**Fig. S3.**

Maximum-likelihood population tree inferred by TreeMix. The tree includes European, Asian and North American wolf populations, with African wild dog, culpeo, and dhole as outgroups. The tree is rooted on the African wild dog. Branch lengths are proportional to the drift parameter. (A) Tree without migration edges. (B) Trees allowing zero to five migration edges (subpanels, left to right, top to bottom). Migration arrows are colored by weight, from yellow (low) to orange (high). All inferred migration edges involve non-European populations; no gene flow was detected among European wolf populations.

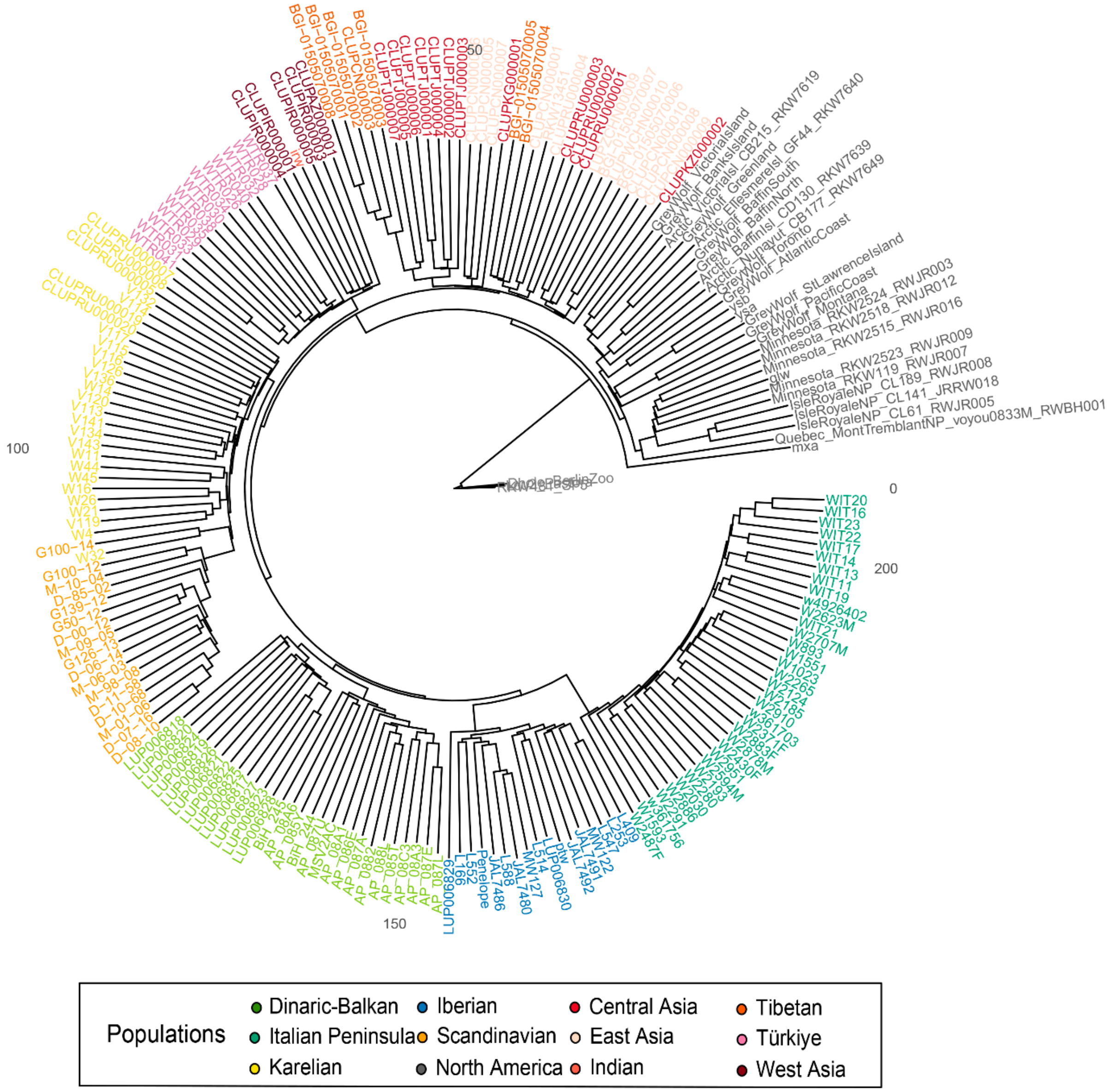


**Fig. S4.**

Neighbour-joining tree based on identity-by-state (IBS) distances. The tree includes European, Asian and North American wolf populations, rooted on the African wild dog, with culpeo and dhole as additional outgroups. Tip labels are colored by population according to the legend. European wolf populations form distinct, well-supported clusters, consistent with pronounced genetic differentiation among them. The topology was concordant when the analysis was repeated with each outgroup individually.

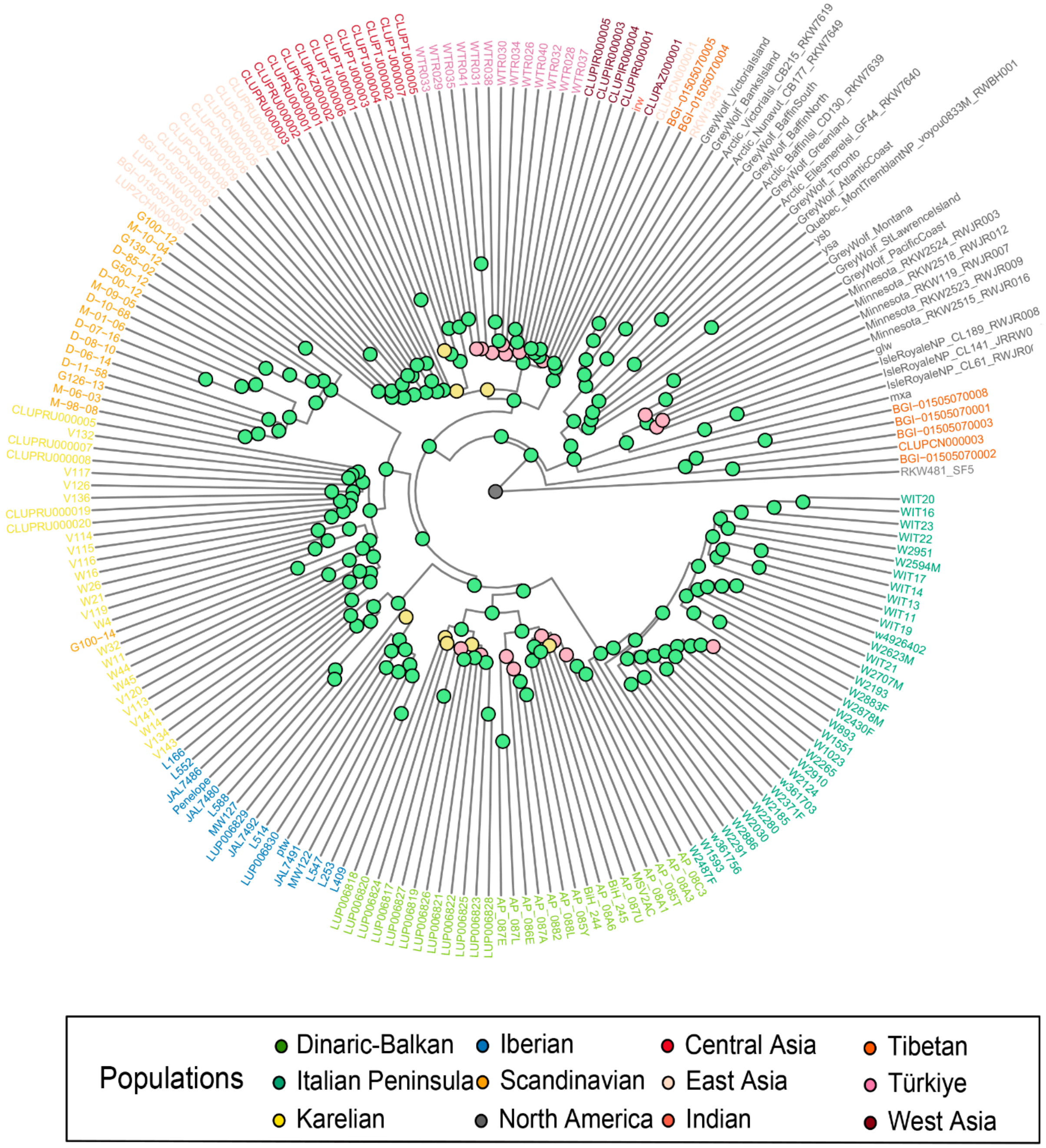


**Fig. S5.**

Phylogenomic tree with bootstrap support values using fastreeR. Node bootstrap values are labelled as coloured circles as strong (higher than 90%, green), moderate (between 70-90, yellow) or weak (lower than 70%, pink) support. Bootstrap values were estimated from 100 replicates. The tree was rooted using the African wild dog as the outgroup.

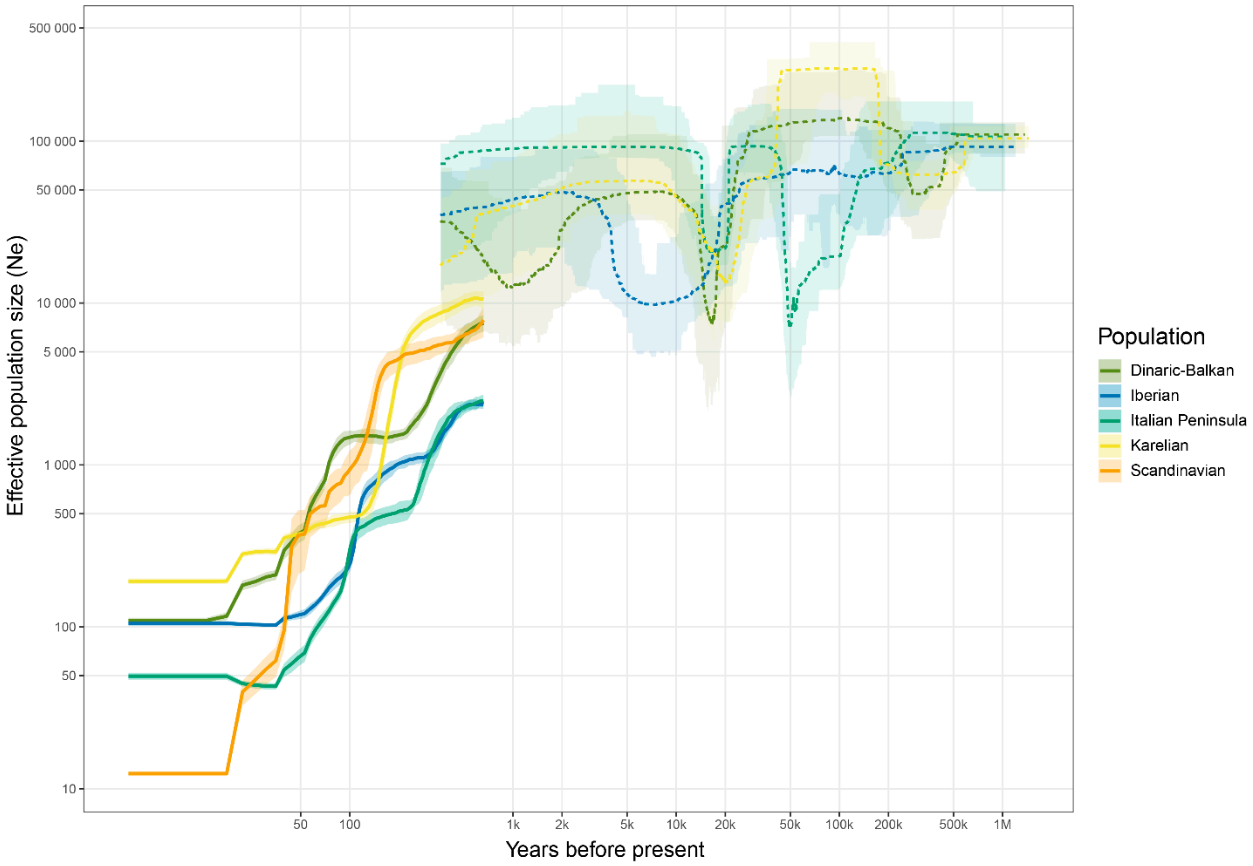


**Fig. S6.**

Stairway plots combined with results from GONE2 (default settings) across European wolf populations. Scandinavian wolves were excluded from Stairway plots as this population was founded in the last decades.

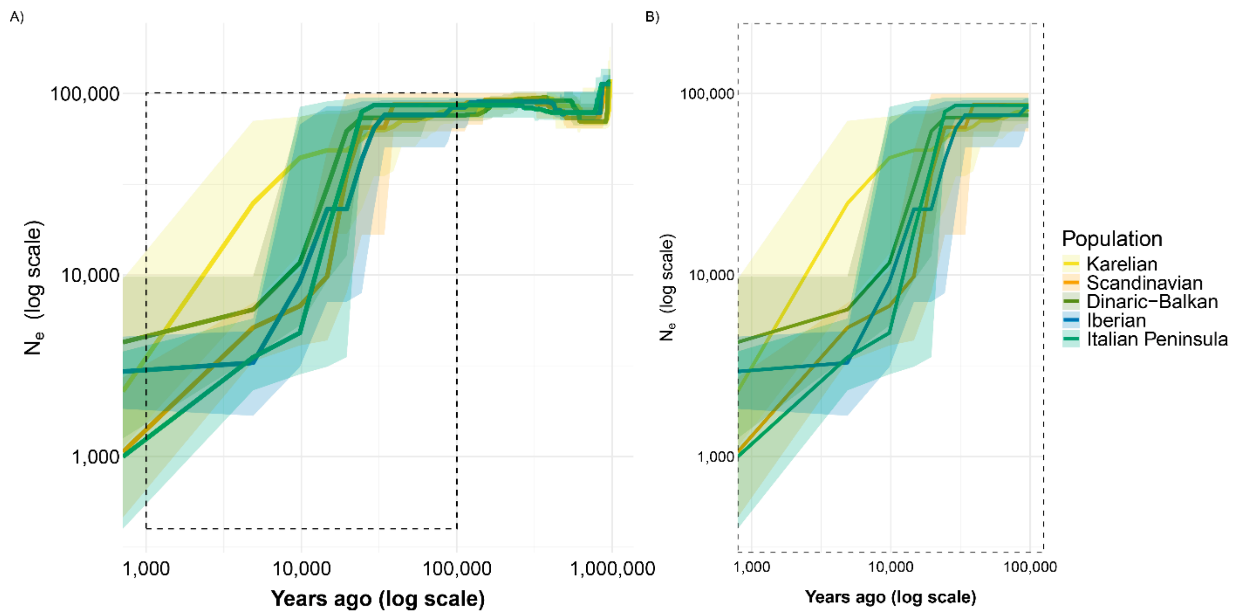


**Fig. S7.**

Effective population size (Ne) trajectories inferred by SMC++ for five European wolf populations. (A) Full demographic history on a log-log scale. (B) Zoomed view of the 1,000–100,000 years ago window, where SMC++ has the greatest inferential power. Lines represent geometric means and shaded ribbons denote 95% confidence intervals across bootstrap chunks.

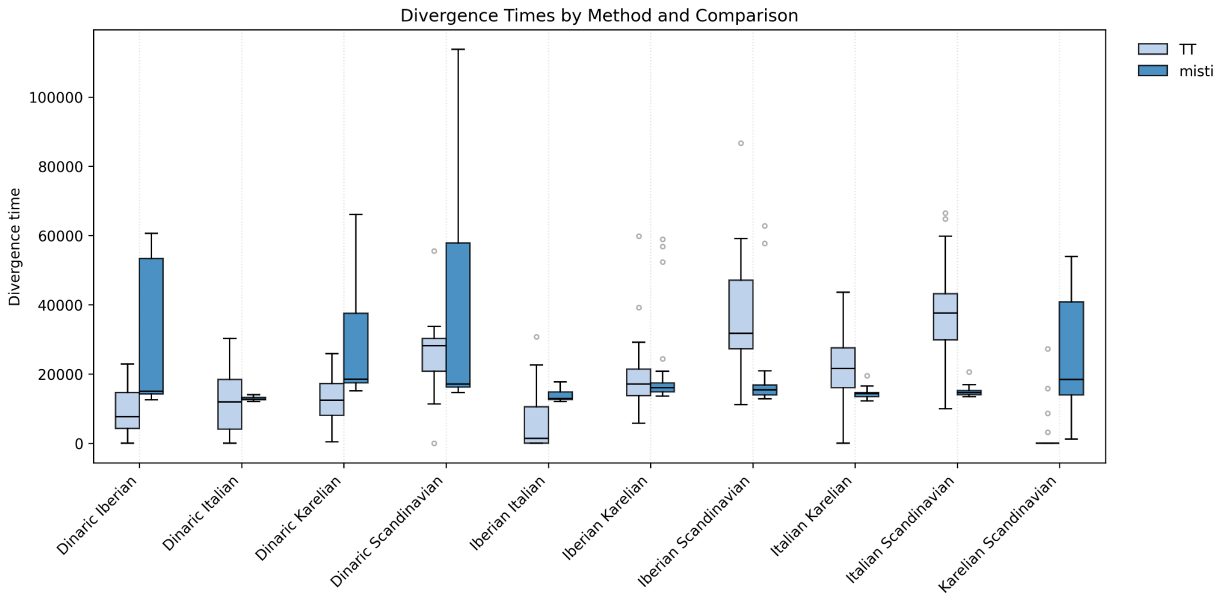


**Fig. S8.**

Average divergence times and confidence intervals across comparisons of the different European wolf populations using TT-method and MiSTI. Only autosomes were used.

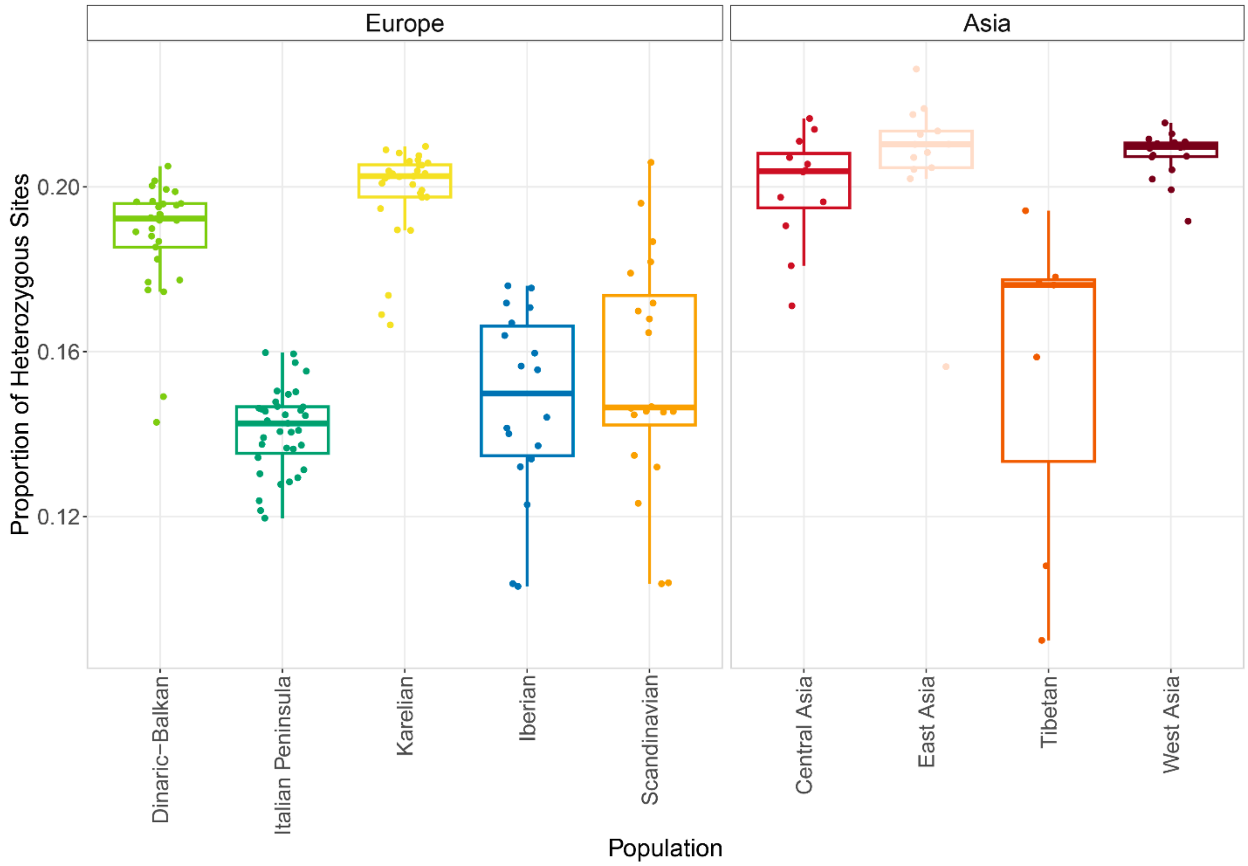


**Fig. S9.**

Heterozygosity in European and Asian wolves. The largest populations show higher heterozygosity compared to smaller and more isolated ones.

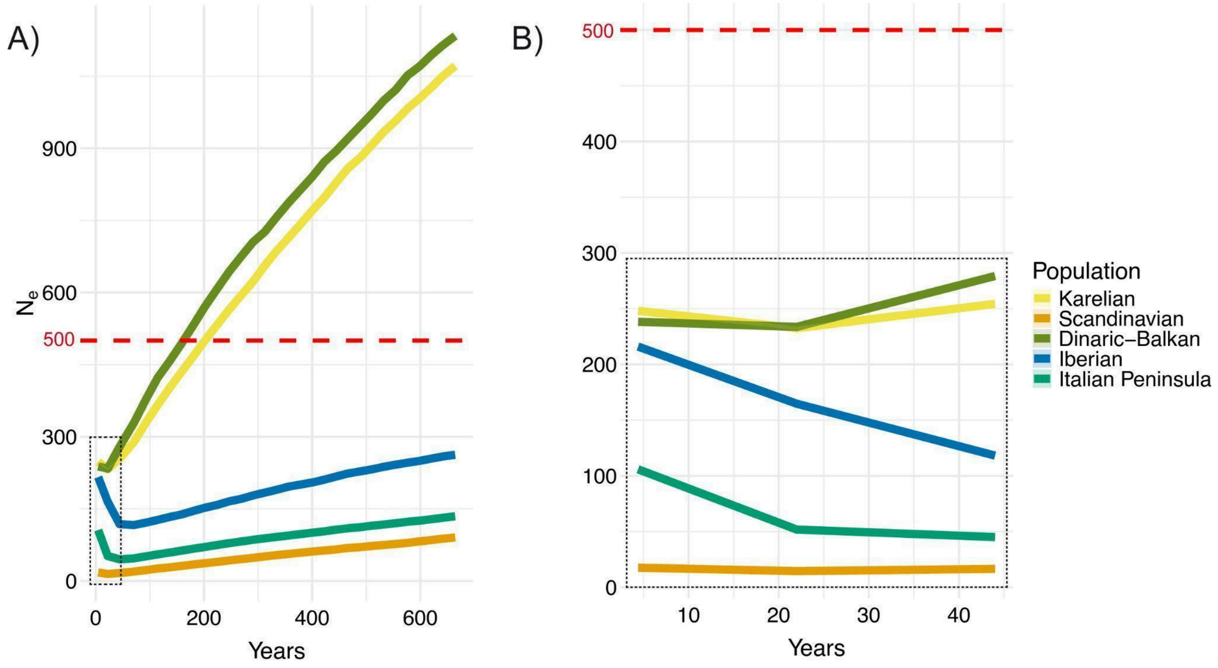


**Fig. S10.**

Demographic inference of gray wolf populations. (A) Effective population size (Ne) in the last 600 years (corresponding to about 150 generations) estimated by GONE2 accounting for structure within each population (option -x) and (B) zooming in on the last 45 years (dotted box in panel A). Lines represent geometric means and shaded ribbons denote 95% confidence intervals across 20 independent replicates (not visible in the graph due to narrow intervals). The red dashed line shows the Ne=500 threshold.

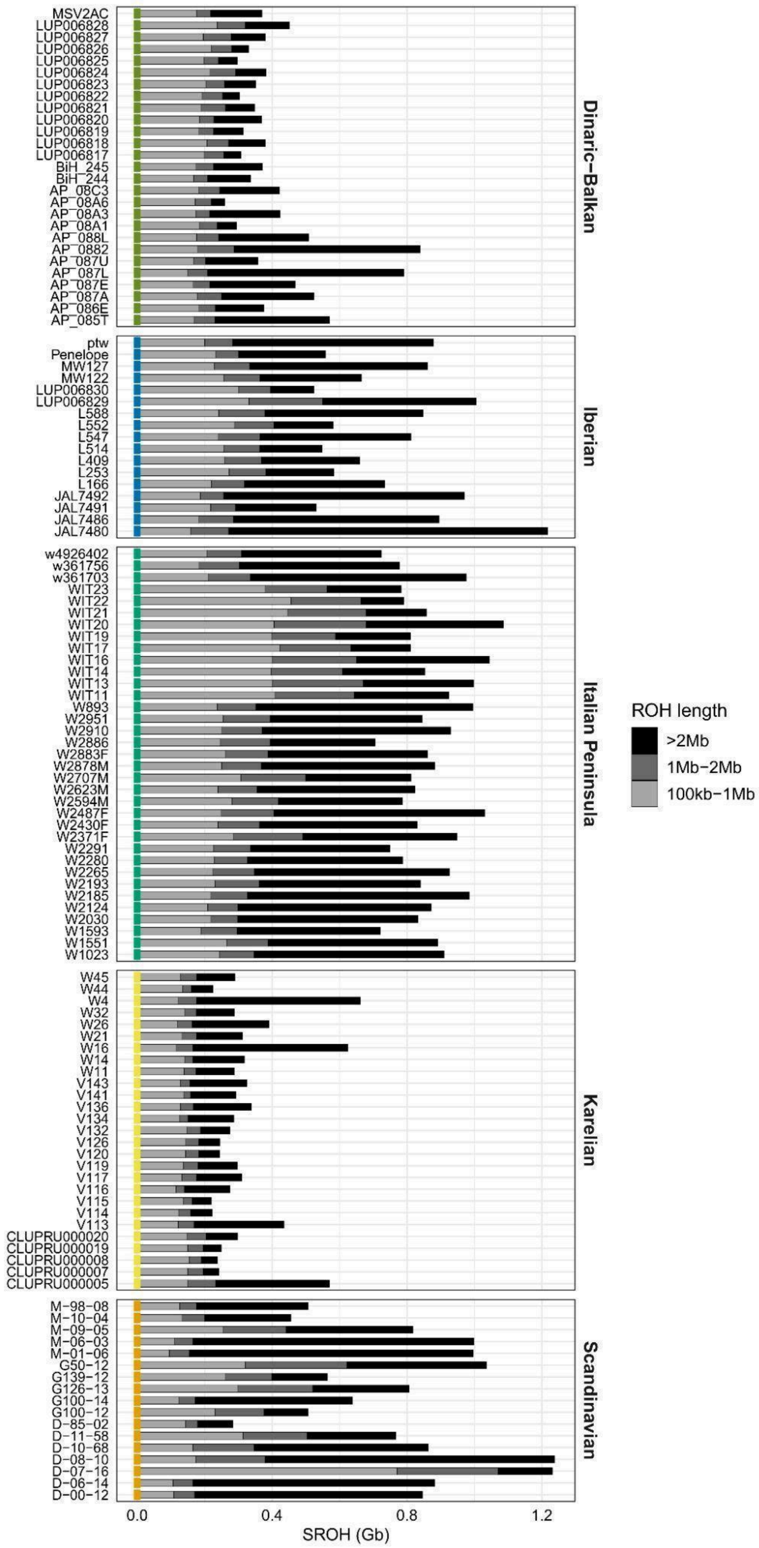


**Fig. S11.**

Sum of runs of homozygosity (ROHs) for different ROH sizes. A grey to black palette indicates short (100 kb–1 Mb), intermediate (1–2 Mb), and long (> 2 Mb) ROHs.

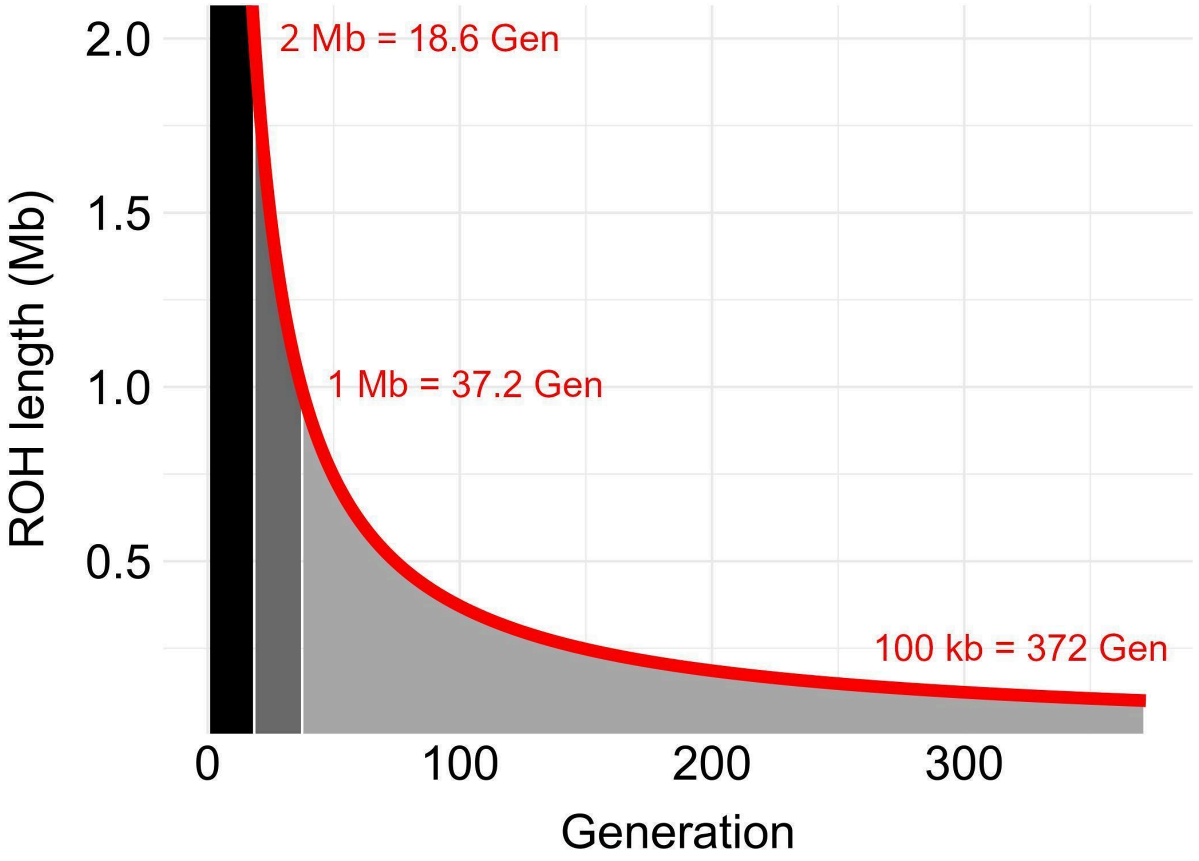


**Fig. S12.**

Coalescence time estimation for runs of homozygosity (ROHs) using the formula of Thompson (2013). Shaded regions (black, dark grey, and grey) indicate the inferred coalescence times (in generations) associated with ROHs of 2 Mb and 1 Mb, which define thresholds for long (>2 Mb), intermediate (1-2 Mb), and short (100 kb-1 Mb) ROHs. The corresponding generation estimates are indicated in red on the figure. The 100-kb threshold is shown as the lower bound for short ROHs.

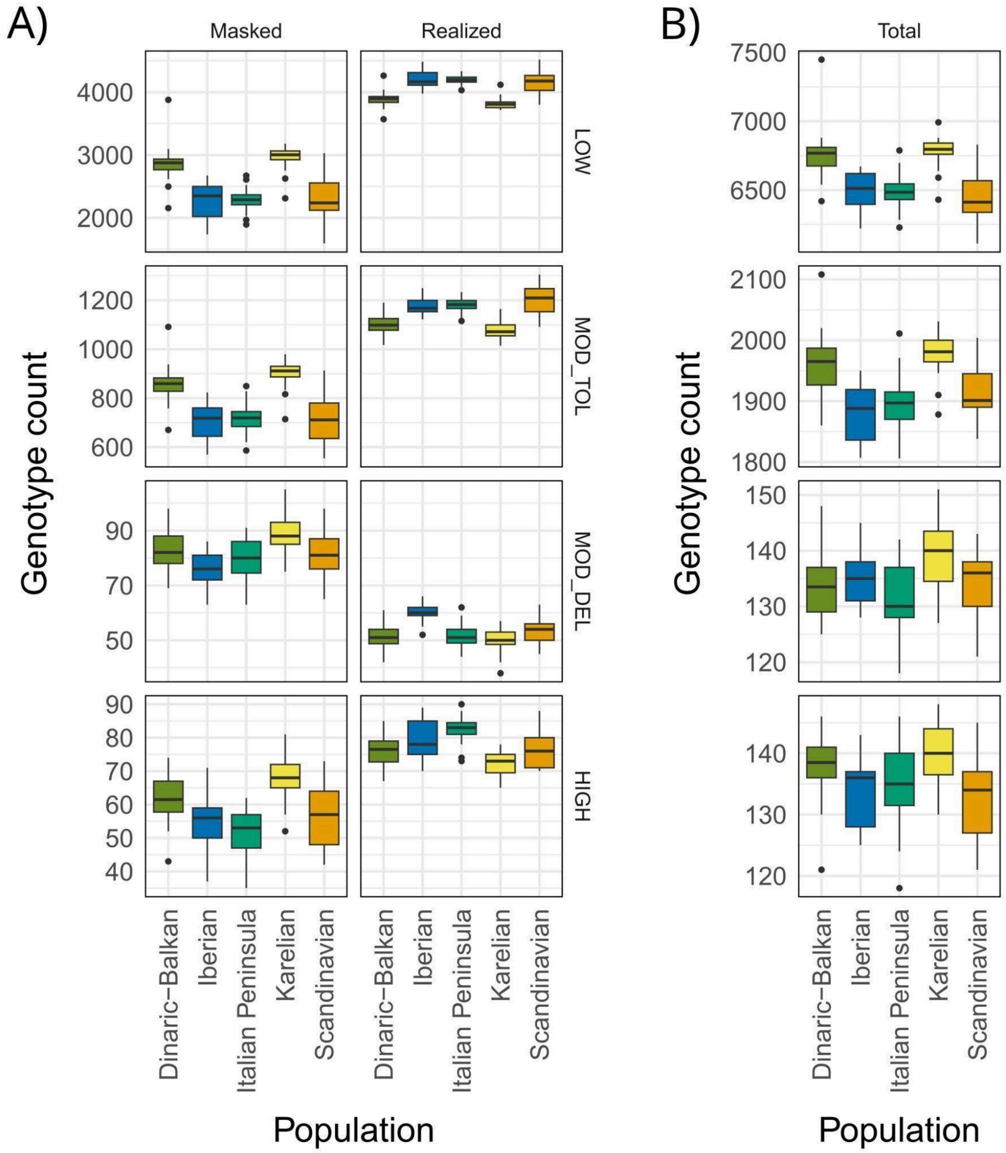


**Fig. S13.**

Genetic load estimates in European gray wolf populations across functional classes (i.e., LOW = low deleteriousness; MOD_TOL = moderate and tolerated deleteriousness; MOD_DEL = moderate deleteriousness; HIGH = high deleteriousness). A) Masked and realized genetic loads are calculated from variants in heterozygous and homozygous states, respectively. B) Total genetic load represents the sum of masked and realized load. Genetic load estimates are based on the number of genotyped loci carrying deleterious variants.

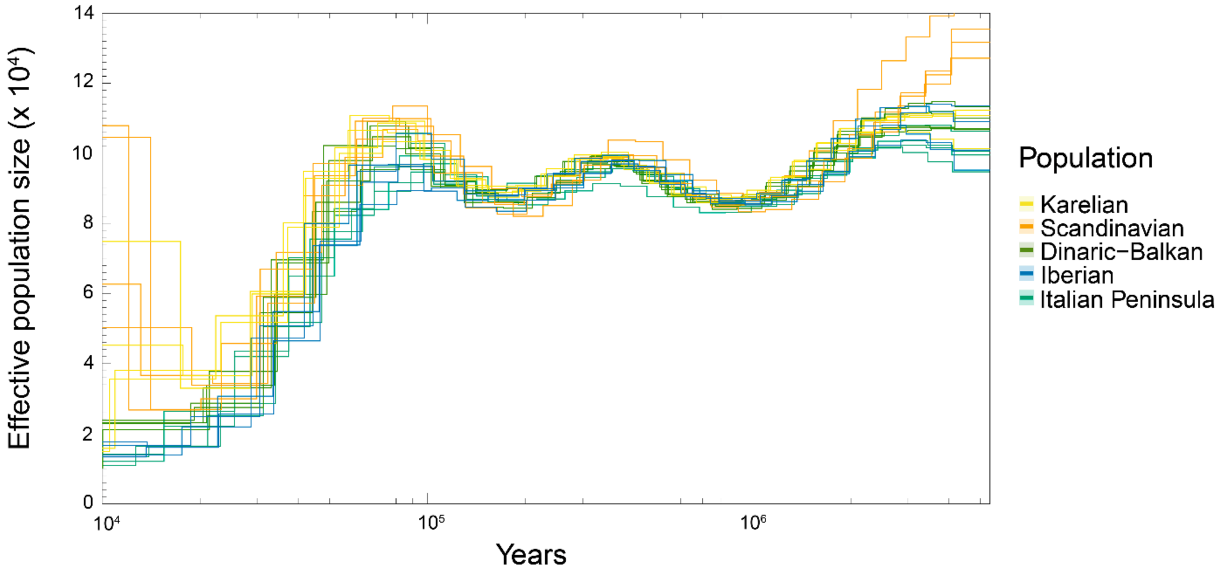


**Fig. S14.**

Pairwise sequentially Markovian coalescent (PSMC) analysis of the demographic histories of 5 wolves representing the different European populations, showing a large population decline starting about 100,000 years ago.
