## Supplementary Tables for "Misleading Success: Genomes Reveal Critical Risks to European Gray Wolves"

**Table S1.** Sample information and pre-processing results of resequenced wolves from this study.

| VCF_ID | BioProject | Accession number | Species | Country |
|---|---|---|---|---|
| AP_07M5 | PRJEB110322 | pending | *Canis lupus* | Croatia |
| AP_082A | not available (only used for initial filters) | not available (only used fo1 | *Canis lupus* | Croatia |
| AP_0844 | not available (only used for initial filters) | not available (only used fo1 | *Canis lupus* | Croatia |
| AP_085T | PRJEB110322 | pending | *Canis lupus* | Slovenia |
| AP_085Y | not available (only used for initial filters) | not available (only used fo1 | *Canis lupus* | Croatia |
| AP_086E | PRJEB110322 | pending | *Canis lupus* | Slovenia |
| AP_086F | PRJEB110322 | pending | *Canis lupus* | Slovenia |
| AP_086Y | not available (only used for initial filters) | not available (only used fo1 | *Canis lupus* | Croatia |
| AP_0870 | not available (only used for initial filters) | not available (only used fo1 | *Canis lupus* | Croatia |
| AP_0872 | PRJEB110322 | pending | *Canis lupus* | Croatia |
| AP_0878 | PRJEB110322 | pending | *Canis lupus* | Croatia |
| AP_087A | PRJEB110322 | pending | *Canis lupus* | Croatia |
| AP_087E | PRJEB110322 | pending | *Canis lupus* | Croatia |
| AP_087F | not available (only used for initial filters) | not available (only used fo1 | *Canis lupus* | Croatia |
| AP_087L | PRJEB110322 | pending | *Canis lupus* | Croatia |
| AP_087U | PRJEB110322 | pending | *Canis lupus* | Croatia |
| AP_087Y | not available (only used for initial filters) | not available (only used fo1 | *Canis lupus* | Croatia |
| AP_0882 | PRJEB110322 | pending | *Canis lupus* | Croatia |
| AP_0886 | not available (only used for initial filters) | not available (only used fo1 | *Canis lupus* | Croatia |
| AP_088F | PRJEB110322 | pending | *Canis lupus* | Bosnia & H |
| AP_088L | PRJEB110322 | pending | *Canis lupus* | Croatia |
| AP_088P | PRJEB110322 | pending | *Canis lupus* | Croatia |
| AP_08A1 | PRJEB110322 | pending | *Canis lupus* | Croatia |
| AP_08A3 | PRJEB110322 | pending | *Canis lupus* | Croatia |
| AP_08A6 | PRJEB110322 | pending | *Canis lupus* | Croatia |
| AP_08AK | not available (only used for initial filters) | not available (only used fo1 | *Canis lupus* | Croatia |
| AP_08C3 | PRJEB110322 | pending | *Canis lupus* | Slovenia |
| AP_08C5 | PRJEB110322 | pending | *Canis lupus* | Slovenia |
| BiH_244 | PRJEB110322 | pending | *Canis lupus* | Bosnia & H |
| BiH_245 | PRJEB110322 | pending | *Canis lupus* | Bosnia & H |
| CH_0LLU | not available (only used for initial filters) | not available (only used fo1 | *Canis lupus* | Croatia |
| CH_0LLY | not available (only used for initial filters) | not available (only used fo1 | *Canis lupus* | Croatia |
| CH_0LM3 | not available (only used for initial filters) | not available (only used fo1 | *Canis lupus* | Croatia |

| | | | | |
|---|---|---|---|---|
| CK_0PET | not available (only used for initial filters) | not available (only used foı | *Canis lupus* | Slovenia |
| DTR001 | not available (only used for initial filters) | not available (only used foı | *Canis familiari* | Turkey |
| DTR002 | not available (only used for initial filters) | not available (only used foı | *Canis familiari* | Turkey |
| DTR003 | not available (only used for initial filters) | not available (only used foı | *Canis familiari* | Turkey |
| JAL7480 | PRJEB110322 | pending | *Canis lupus* | Spain |
| JAL7485 | not available (only used for initial filters) | not available (only used foı | *Canis lupus* | Spain |
| JAL7486 | PRJEB110322 | pending | *Canis lupus* | Spain |
| JAL7491 | PRJEB110322 | pending | *Canis lupus* | Spain |
| JAL7492 | PRJEB110322 | pending | *Canis lupus* | Spain |
| L058 | not available (only used for initial filters) | not available (only used foı | *Canis lupus* | Spain |
| L166 | PRJEB110322 | pending | *Canis lupus* | Spain |
| L186 | PRJEB110322 | pending | *Canis lupus* | Spain |
| L352 | not available (only used for initial filters) | not available (only used foı | *Canis lupus* | Spain |
| L781 | PRJEB110322 | pending | *Canis lupus* | Spain |
| L800 | PRJEB110322 | pending | *Canis lupus* | Spain |
| M2C3H | PRJEB110322 | pending | *Canis lupus* | Croatia |
| M2CHL | not available (only used for initial filters) | not available (only used foı | *Canis lupus* | Croatia |
| M2E71 | not available (only used for initial filters) | not available (only used foı | *Canis lupus* | Slovenia |
| MSV2AC | PRJEB110322 | pending | *Canis lupus* | Slovenia |
| W1433 | not available (only used for initial filters) | not available (only used foı | *Canis lupus* | Italy |
| W1551 | PRJEB110322 | pending | *Canis lupus* | Italy |
| W1593 | PRJEB110322 | pending | *Canis lupus* | Italy |
| W1869 | PRJEB110322 | pending | *Canis lupus* | Italy |
| W191598 | not available (only used for initial filters) | not available (only used foı | *Canis lupus* | Germany |
| W200546 | not available (only used for initial filters) | not available (only used foı | *Canis lupus* | Germany |
| W2030 | PRJEB110322 | pending | *Canis lupus* | Italy |
| W2124 | PRJEB110322 | pending | *Canis lupus* | Italy |
| W2185 | PRJEB110322 | pending | *Canis lupus* | Italy |
| W2192 | not available (only used for initial filters) | not available (only used foı | *Canis lupus* | Italy |
| W2193 | PRJEB110322 | pending | *Canis lupus* | Italy |
| W224957 | not available (only used for initial filters) | not available (only used foı | *Canis lupus* | Germany |
| W2265 | PRJEB110322 | pending | *Canis lupus* | Italy |
| W226855 | not available (only used for initial filters) | not available (only used foı | *Canis lupus* | Germany |
| W2280 | PRJEB110322 | pending | *Canis lupus* | Italy |
| W2291 | PRJEB110322 | pending | *Canis lupus* | Italy |

| | | | | |
|---|---|---|---|---|
| W2371F | PRJEB110322 | pending | *Canis lupus* | Italy |
| W2419 | not available (only used for initial filters) | not available (only used for | *Canis lupus* | Italy |
| W2430F | PRJEB110322 | pending | *Canis lupus* | Italy |
| W2470 | PRJEB110322 | pending | *Canis lupus* | Italy |
| W2487F | PRJEB110322 | pending | *Canis lupus* | Italy |
| W2499M | PRJEB110322 | pending | *Canis lupus* | Italy |
| W2500M | PRJEB110322 | pending | *Canis lupus* | Italy |
| W2501F | PRJEB110322 | pending | *Canis lupus* | Italy |
| W2594M | PRJEB110322 | pending | *Canis lupus* | Italy |
| W2623M | PRJEB110322 | pending | *Canis lupus* | Italy |
| W2685F | PRJEB110322 | pending | *Canis lupus* | Italy |
| W2707M | PRJEB110322 | pending | *Canis lupus* | Italy |
| W2725 | not available (only used for initial filters) | not available (only used for | *Canis lupus* | Italy |
| W2728 | PRJEB110322 | pending | *Canis lupus* | Italy |
| W2878M | PRJEB110322 | pending | *Canis lupus* | Italy |
| W2883F | PRJEB110322 | pending | *Canis lupus* | Italy |
| W2886 | PRJEB110322 | pending | *Canis lupus* | Italy |
| W2910 | PRJEB110322 | pending | *Canis lupus* | Italy |
| W2951 | PRJEB110322 | pending | *Canis lupus* | Italy |
| w361703 | PRJEB110322 | pending | *Canis lupus* | Italy |
| w361756 | PRJEB110322 | pending | *Canis lupus* | Italy |
| w484049 | not available (only used for initial filters) | not available (only used for | *Canis lupus* | Italy |
| w4902419 | PRJEB110322 | pending | *Canis lupus* | Italy |
| w4926402 | PRJEB110322 | pending | *Canis lupus* | Italy |
| WTR025 | PRJEB110322 | pending | *Canis lupus* | Turkey |
| WTR026 | PRJEB110322 | pending | *Canis lupus* | Turkey |
| WTR027 | PRJEB110322 | pending | *Canis lupus* | Turkey |
| WTR028 | PRJEB110322 | pending | *Canis lupus* | Turkey |
| WTR029 | PRJEB110322 | pending | *Canis lupus* | Turkey |
| WTR030 | PRJEB110322 | pending | *Canis lupus* | Turkey |
| WTR031 | PRJEB110322 | pending | *Canis lupus* | Turkey |
| WTR032 | PRJEB110322 | pending | *Canis lupus* | Turkey |
| WTR033 | PRJEB110322 | pending | *Canis lupus* | Turkey |
| WTR034 | PRJEB110322 | pending | *Canis lupus* | Turkey |
| WTR035 | PRJEB110322 | pending | *Canis lupus* | Turkey |

| | | | | |
|---|---|---|---|---|
| WTR037 | PRJEB110322 | pending | *Canis lupus* | Turkey |
| WTR038 | PRJEB110322 | pending | *Canis lupus* | Turkey |
| WTR039 | PRJEB110322 | pending | *Canis lupus* | Turkey |
| WTR040 | PRJEB110322 | pending | *Canis lupus* | Turkey |
| WTR041 | PRJEB110322 | pending | *Canis lupus* | Turkey |
| W1129 | not available (only used for initial filters) | not available (only used fo1 | *Canis familiari* | Italy |
| W1208 | not available (only used for initial filters) | not available (only used fo1 | *Canis familiari* | Italy |
| W1297 | not available (only used for initial filters) | not available (only used fo1 | *Canis familiari* | Italy |
| W1431 | not available (only used for initial filters) | not available (only used fo1 | *Canis familiari* | Italy |
| W1432 | not available (only used for initial filters) | not available (only used fo1 | *Canis familiari* | Italy |
| W1449 | not available (only used for initial filters) | not available (only used fo1 | *Canis familiari* | Italy |
| W1564 | not available (only used for initial filters) | not available (only used fo1 | *Canis familiari* | Italy |
| W1578 | not available (only used for initial filters) | not available (only used fo1 | *Canis familiari* | Italy |

**Table S1.** Sample i

| VCF_ID | Locality | Population | Year | # raw reads | # filtered reads | # mapped Reads | % Mapped Rea |
|---|---|---|---|---|---|---|---|
| AP_07M5 | Dalmatia (HRV) | Dinaric-Balkan | 2006 | 471.228.464 | 466.941.293 | 464.983.880 | 99,60% |
| AP_082A | Dalmatia (HRV) | Dinaric-Balkan | 2006 | 515.908.318 | 512.492.464 | 491.451.379 | 95,90% |
| AP_0844 | Dalmatia (HRV) | Dinaric-Balkan | 2009 | 523.241.256 | 519.863.366 | 516.859.378 | 99,40% |
| AP_085T | | Dinaric-Balkan | 2005 | 555.188.888 | 551.008.658 | 549.762.147 | 99,80% |
| AP_085Y | | Dinaric-Balkan | 2012 | 458.725.088 | 454.720.318 | 452.718.315 | 99,60% |
| AP_086E | | Dinaric-Balkan | 2012 | 677.428.020 | 672.297.920 | 667.525.686 | 99,30% |
| AP_086F | | Dinaric-Balkan | 2012 | 451.511.008 | 446.696.200 | 444.519.419 | 99,50% |
| AP_086Y | Dalmatia (HRV) | Dinaric-Balkan | 2010 | 401.667.050 | 398.164.054 | 396.455.318 | 99,60% |
| AP_0870 | Dalmatia (HRV) | Dinaric-Balkan | 2010 | 489.855.632 | 485.421.904 | 483.765.036 | 99,70% |
| AP_0872 | Dalmatia (HRV) | Dinaric-Balkan | 2010 | 234.693.878 | 232.840.878 | 231.988.351 | 99,60% |
| AP_0878 | Dalmatia (HRV) | Dinaric-Balkan | 2010 | 443.116.494 | 439.238.455 | 437.246.140 | 99,50% |
| AP_087A | Lika (HRV) | Dinaric-Balkan | 2011 | 431.598.048 | 427.556.285 | 425.796.643 | 99,60% |
| AP_087E | Gorski kotar (HRV) | Dinaric-Balkan | 2010 | 400.744.260 | 397.690.459 | 396.567.241 | 99,70% |
| AP_087F | Dalmatia (HRV) | Dinaric-Balkan | 2010 | 487.956.452 | 483.802.615 | 481.923.127 | 99,60% |
| AP_087L | Gorski kotar (HRV) | Dinaric-Balkan | 2010 | 544.082.224 | 539.306.240 | 537.491.276 | 99,70% |
| AP_087U | Lika (HRV) | Dinaric-Balkan | 2010 | 596.142.444 | 591.435.173 | 589.188.819 | 99,60% |
| AP_087Y | Dalmatia (HRV) | Dinaric-Balkan | 2008 | 512.185.882 | 507.701.086 | 505.729.203 | 99,60% |
| AP_0882 | Gorski kotar (HRV) | Dinaric-Balkan | 2006 | 498.832.356 | 496.149.202 | 491.367.621 | 99,00% |
| AP_0886 | Dalmatia (HRV) | Dinaric-Balkan | 2011 | 546.902.390 | 542.429.459 | 540.289.005 | 99,60% |
| AP_088F | Dalmatia (HRV) | Dinaric-Balkan | 2010 | 456.091.820 | 452.679.914 | 450.114.438 | 99,40% |
| AP_088L | Gorski kotar (HRV) | Dinaric-Balkan | 2011 | 547.015.570 | 542.740.210 | 540.896.108 | 99,70% |
| AP_088P | Dalmatia (HRV) | Dinaric-Balkan | 2010 | 530.579.130 | 525.772.526 | 523.608.129 | 99,60% |
| AP_08A1 | Lika (HRV) | Dinaric-Balkan | 2011 | 509.636.600 | 505.236.405 | 503.226.173 | 99,60% |
| AP_08A3 | Gorski kotar (HRV) | Dinaric-Balkan | 2012 | 405.622.500 | 402.707.420 | 400.597.291 | 99,50% |
| AP_08A6 | Dalmatia (HRV) | Dinaric-Balkan | 2011 | 483.462.750 | 479.829.727 | 478.308.623 | 99,70% |
| AP_08AK | Dalmatia (HRV) | Dinaric-Balkan | 2011 | 512.343.170 | 508.250.652 | 506.802.854 | 99,70% |
| AP_08C3 | | Dinaric-Balkan | 2011 | 466.244.480 | 463.080.255 | 461.786.377 | 99,70% |
| AP_08C5 | | Dinaric-Balkan | 2012 | 538.491.724 | 534.192.980 | 532.877.820 | 99,80% |
| BiH_244 | BIH | Dinaric-Balkan | 2016 | 457.106.244 | 454.246.907 | 452.723.390 | 99,70% |
| BiH_245 | BIH | Dinaric-Balkan | 2016 | 413.555.796 | 409.853.793 | 408.251.042 | 99,60% |
| CH_0LLU | Dalmatia (HRV) | Dinaric-Balkan | 2006 | 418.731.080 | 415.468.149 | 414.248.565 | 99,70% |
| CH_0LLY | Dalmatia (HRV) | Dinaric-Balkan | 2013 | 473.866.808 | 469.962.621 | 453.742.921 | 96,50% |
| CH_0LM3 | Lika (HRV) | Dinaric-Balkan | 2013 | 481.922.482 | 478.612.459 | 475.984.597 | 99,50% |

| | | | | | | | |
|---|---|---|---|---|---|---|---|
| CK_0PET | | Dinaric-Balkan | 2023 | 669.952.896 | 665.058.574 | 662.792.690 | 99,70% |
| DTR001 | Sarıkamış | Village dog Turkey | 2023 | 449.817.038 | 446.140.012 | 444.521.643 | 99,60% |
| DTR002 | Sarıkamış | Village dog Turkey | 2023 | 443.288.712 | 438.853.000 | 437.398.101 | 99,70% |
| DTR003 | Sarıkamış | Village dog Turkey | 2023 | 494.913.436 | 489.643.426 | 487.815.689 | 99,60% |
| JAL7480 | Guadalajara | NW Iberia | 2020 | 323.789.916 | 320.586.300 | 319.819.769 | 99,80% |
| JAL7485 | Asturias | NW Iberia | 2020 | 514.526.632 | 508.605.404 | 507.454.587 | 99,80% |
| JAL7486 | Zamora | NW Iberia | 2020 | 434.749.010 | 430.623.325 | 429.606.034 | 99,80% |
| JAL7491 | Lugo | NW Iberia | 2020 | 545.742.748 | 539.385.847 | 537.819.048 | 99,70% |
| JAL7492 | A Coruña | NW Iberia | 2020 | 414.171.942 | 409.573.617 | 408.247.062 | 99,70% |
| L058 | Asturias | NW Iberia | 2002 | 403.287.426 | 400.657.487 | 393.813.634 | 98,30% |
| L166 | Asturias | NW Iberia | 2004 | 506.520.744 | 501.796.423 | 499.262.380 | 99,50% |
| L186 | Asturias | NW Iberia | 2006 | 400.929.988 | 398.175.927 | 397.143.576 | 99,70% |
| L352 | Asturias | NW Iberia | 2007 | 529.210.864 | 524.875.954 | 523.327.383 | 99,70% |
| L781 | Galicia | NW Iberia | 2015 | 392.280.552 | 388.692.860 | 387.452.745 | 99,70% |
| L800 | Galicia | NW Iberia | 2015 | 452.785.986 | 448.963.956 | 447.576.174 | 99,70% |
| M2C3H | Dalmatia (HRV) | Dinaric-Balkan | 2016 | 580.788.398 | 576.013.941 | 573.579.849 | 99,60% |
| M2CHL | Dalmatia (HRV) | Dinaric-Balkan | Likely 2010 | 400.503.956 | 397.330.490 | 395.722.556 | 99,60% |
| M2E71 | | Dinaric-Balkan | 2023 | 452.356.276 | 448.234.548 | 446.204.119 | 99,50% |
| MSV2AC | | Dinaric-Balkan | 2023 | 558.506.026 | 553.163.274 | 549.677.122 | 99,40% |
| W1433 | Campania | Italian Peninsula | 2012 | 460.426.038 | 454.874.416 | 449.744.304 | 98,90% |
| W1551 | Umbria | Italian Peninsula | 2010 | 315.695.126 | 311.773.695 | 310.046.038 | 99,40% |
| W1593 | Toscana | Italian Peninsula | 2013 | 407.978.194 | 402.305.324 | 399.247.066 | 99,20% |
| W1869 | Emilia Romagna | Italian Peninsula | 2015 | 358.384.456 | 355.027.121 | 353.237.265 | 99,50% |
| W191598 | | Central European | | 434.029.904 | 429.814.715 | 428.225.944 | 99,60% |
| W200546 | | Central European | | 450.039.210 | 445.919.729 | 442.274.057 | 99,20% |
| W2030 | Emilia-Romagna | Italian Peninsula | 2016 | 404.574.356 | 400.433.894 | 398.439.955 | 99,50% |
| W2124 | Piemonte | Italian Peninsula | 2017 | 434.549.600 | 430.547.460 | 427.927.321 | 99,40% |
| W2185 | Emilia Romagna | Italian Peninsula | 2017 | 403.484.664 | 400.024.176 | 398.334.133 | 99,60% |
| W2192 | Molise | Italian Peninsula | 2017 | 404.929.434 | 401.088.820 | 382.355.024 | 95,30% |
| W2193 | Molise | Italian Peninsula | 2017 | 503.068.672 | 498.705.743 | 489.623.652 | 98,20% |
| W224957 | | Central European | | 395.492.220 | 391.861.161 | 390.322.695 | 99,60% |
| W2265 | Emilia Romagna | Italian Peninsula | 2018 | 422.238.960 | 418.848.098 | 417.347.660 | 99,60% |
| W226855 | | Central European | | 555.874.462 | 549.736.148 | 546.825.316 | 99,50% |
| W2280 | Emilia Romagna | Italian Peninsula | 2018 | 541.078.960 | 535.696.724 | 533.439.873 | 99,60% |
| W2291 | Piemonte | Italian Peninsula | 2018 | 404.836.988 | 401.893.705 | 400.396.433 | 99,60% |

| | | | | | | | |
|---|---|---|---|---|---|---|---|
| W2371F | Toscana | Italian Peninsula | 2019 | 296.600.960 | 295.630.035 | 293.639.880 | 99,30% |
| W2419 | Molise | Italian Peninsula | 2019 | 374.212.434 | 372.521.151 | 358.427.351 | 96,20% |
| W2430F | Abruzzo | Italian Peninsula | 2019 | 285.992.880 | 279.645.140 | 278.420.136 | 99,60% |
| W2470 | Abruzzo | Italian Peninsula | 2020 | 257.829.128 | 254.009.978 | 251.551.056 | 99,00% |
| W2487F | Calabria | Italian Peninsula | 2019 | 323.320.086 | 320.137.502 | 316.295.991 | 98,80% |
| W2499M | Calabria | Italian Peninsula | 2020 | 44.899.488 | 42.961.518 | 42.787.480 | 99,60% |
| W2500M | Calabria | Italian Peninsula | 2020 | 526.467.274 | 534.827.186 | 531.793.333 | 99,40% |
| W2501F | Calabria | Italian Peninsula | 2020 | 31.491.644 | 32.596.586 | 26.258.806 | 80,60% |
| W2594M | Abruzzo | Italian Peninsula | 2021 | 406.015.170 | 402.493.801 | 400.988.992 | 99,60% |
| W2623M | Umbria | Italian Peninsula | 2021 | 463.449.176 | 459.001.926 | 456.283.270 | 99,40% |
| W2685F | Marche | Italian Peninsula | 2021 | 488.129.170 | 495.886.269 | 460.030.876 | 92,80% |
| W2707M | Emilia Romagna | Italian Peninsula | 2021 | 372.043.214 | 369.642.296 | 367.946.745 | 99,50% |
| W2725 | Lazio | Italian Peninsula | 2020 | 399.671.734 | 396.600.377 | 394.969.117 | 99,60% |
| W2728 | Lazio | Italian Peninsula | 2021 | 359.029.676 | 365.385.582 | 253.804.587 | 69,50% |
| W2878M | Molise | Italian Peninsula | 2022 | 451.368.208 | 447.610.398 | 445.012.244 | 99,40% |
| W2883F | Molise | Italian Peninsula | 2023 | 463.605.880 | 459.808.345 | 454.519.420 | 98,80% |
| W2886 | Piemonte | Italian Peninsula | 2022 | 559.913.990 | 561.091.007 | 521.764.536 | 93,00% |
| W2910 | Piemonte | Italian Peninsula | 2023 | 429.666.290 | 426.341.475 | 424.684.928 | 99,60% |
| W2951 | Abruzzo | Italian Peninsula | 2023 | 419.022.572 | 416.020.998 | 414.544.684 | 99,60% |
| w361703 | Arcidosso (GR) | Italian Peninsula | 2022 | 482.124.794 | 477.980.721 | 476.394.429 | 99,70% |
| w361756 | Arcidosso (GR) | Italian Peninsula | 2022 | 526.229.858 | 521.358.733 | 519.620.149 | 99,70% |
| w484049 | Arcidosso (GR) | Italian Peninsula | 2022 | 426.841.212 | 423.366.236 | 421.949.656 | 99,70% |
| w4902419 | Arcidosso (GR) | Italian Peninsula | 2022 | 605.050.720 | 599.664.067 | 597.682.350 | 99,70% |
| w4926402 | Arcidosso (GR) | Italian Peninsula | 2022 | 388.922.622 | 385.888.929 | 384.533.370 | 99,60% |
| WTR025 | Sarıkamış | Turkey | 2020 | 404.558.918 | 400.630.597 | 399.086.839 | 99,60% |
| WTR026 | Sarıkamış | Turkey | 2020 | 464.778.602 | 459.429.380 | 458.073.368 | 99,70% |
| WTR027 | Sarıkamış | Turkey | 2020 | 402.347.848 | 398.424.168 | 396.925.894 | 99,60% |
| WTR028 | Sarıkamış | Turkey | 2020 | 384.185.150 | 381.082.646 | 379.711.988 | 99,60% |
| WTR029 | Sarıkamış | Turkey | 2020 | 615.725.672 | 609.143.421 | 607.219.586 | 99,70% |
| WTR030 | Sarıkamış | Turkey | 2021 | 490.860.456 | 485.735.422 | 484.193.862 | 99,70% |
| WTR031 | Sarıkamış | Turkey | 2021 | 482.341.500 | 477.332.073 | 475.867.692 | 99,70% |
| WTR032 | Sarıkamış | Turkey | 2021 | 456.202.068 | 451.981.664 | 450.429.324 | 99,70% |
| WTR033 | Sarıkamış | Turkey | 2021 | 473.650.436 | 468.912.962 | 467.291.882 | 99,70% |
| WTR034 | Sarıkamış | Turkey | 2021 | 530.261.780 | 525.169.298 | 523.482.384 | 99,70% |
| WTR035 | Sarıkamış | Turkey | 2022 | 517.449.940 | 512.711.221 | 511.110.460 | 99,70% |

| | | | | | | | |
|---|---|---|---|---|---|---|---|
| WTR037 | Sarıkamış | Turkey | 2022 | 559.107.134 | 553.668.912 | 551.776.424 | 99,70% |
| WTR038 | Sarıkamış | Turkey | 2022 | 458.163.056 | 454.307.199 | 452.971.329 | 99,70% |
| WTR039 | Sarıkamış | Turkey | 2022 | 504.742.664 | 500.519.511 | 498.751.555 | 99,60% |
| WTR040 | Sarıkamış | Turkey | 2023 | 503.187.418 | 499.127.494 | 497.431.928 | 99,70% |
| WTR041 | Sarıkamış | Turkey | 2023 | 504.221.050 | 500.171.207 | 498.439.896 | 99,70% |
| W1129 | | Village dog Italy | | 407.228.402 | 407.037.131 | 405.860.993 | 99,71% |
| W1208 | | Village dog Italy | | 653.688.548 | 651.278.443 | 649.123.694 | 99,67% |
| W1297 | | Village dog Italy | | 539.741.094 | 539.892.456 | 529.209.619 | 98,02% |
| W1431 | | Village dog Italy | | 550.572.934 | 550.261.653 | 548.780.023 | 99,73% |
| W1432 | | Village dog Italy | | 422.184.798 | 422.199.340 | 420.874.845 | 99,69% |
| W1449 | | Village dog Italy | | 462.109.232 | 461.832.176 | 446.684.334 | 96,72% |
| W1564 | | Village dog Italy | | 466.923.294 | 466.714.150 | 459.454.212 | 99,01% |
| W1578 | | Village dog Italy | | 416.648.216 | 417.787.054 | 386.566.372 | 92,53% |

**Table S1.** Sample i

| VCF_ID | % Properly Pai | % Duplicates | MeanDepth | Passed quality 1 | Passed dog | Passed relatedness filters |
|---|---|---|---|---|---|---|
| AP_07M5 | 97,20% | 13,80% | 22,62 | yes | no | NA |
| AP_082A | 93,70% | 11,10% | 21,44 | yes | no | NA |
| AP_0844 | 97,30% | 16,70% | 23,99 | yes | no | NA |
| AP_085T | 97,70% | 13,10% | 26,68 | yes | yes | yes |
| AP_085Y | 97,20% | 13,00% | 22,41 | yes | no | NA |
| AP_086E | 97,00% | 17,70% | 30,59 | yes | yes | yes |
| AP_086F | 97,10% | 14,30% | 22,12 | yes | yes | no |
| AP_086Y | 97,20% | 14,00% | 19,91 | yes | no | NA |
| AP_0870 | 97,30% | 17,30% | 23,59 | yes | no | NA |
| AP_0872 | 96,60% | 20,70% | 10,82 | yes | no | NA |
| AP_0878 | 97,00% | 16,90% | 21,22 | yes | no | NA |
| AP_087A | 96,80% | 16,70% | 20,38 | yes | yes | yes |
| AP_087E | 97,50% | 16,60% | 19,57 | yes | yes | yes |
| AP_087F | 97,30% | 18,20% | 23,00 | yes | no | NA |
| AP_087L | 97,20% | 18,10% | 26,16 | yes | yes | yes |
| AP_087U | 97,50% | 19,10% | 27,72 | yes | yes | yes |
| AP_087Y | 97,20% | 18,60% | 24,56 | yes | no | NA |
| AP_0882 | 96,20% | 13,30% | 19,46 | yes | yes | yes |
| AP_0886 | 97,10% | 18,50% | 25,94 | yes | no | NA |
| AP_088F | 97,30% | 13,40% | 20,48 | yes | no | NA |
| AP_088L | 97,40% | 16,20% | 26,55 | yes | yes | yes |
| AP_088P | 97,20% | 19,30% | 24,74 | yes | no | NA |
| AP_08A1 | 96,80% | 18,80% | 23,86 | yes | yes | yes |
| AP_08A3 | 97,10% | 20,10% | 18,82 | yes | yes | yes |
| AP_08A6 | 97,30% | 18,90% | 22,62 | yes | yes | yes |
| AP_08AK | 97,50% | 18,70% | 24,48 | yes | no | NA |
| AP_08C3 | 97,20% | 15,30% | 20,16 | yes | yes | yes |
| AP_08C5 | 97,60% | 16,80% | 24,08 | yes | yes | no |
| BiH_244 | 97,30% | 17,70% | 21,40 | yes | yes | yes |
| BiH_245 | 96,70% | 24,60% | 18,08 | yes | yes | yes |
| CH_0LLU | 97,50% | 17,00% | 20,52 | yes | no | NA |
| CH_0LLY | 94,40% | 17,60% | 21,38 | yes | no | NA |
| CH_0LM3 | 97,40% | 16,20% | 22,58 | yes | no | NA |

| | | | | | | |
|---|---|---|---|---|---|---|
| CK_0PET | 97,60% | 15,60% | 30,60 | yes | no | NA |
| DTR001 | 97,40% | 14,60% | 21,79 | yes | no | NA |
| DTR002 | 97,30% | 15,70% | 21,30 | yes | no | NA |
| DTR003 | 96,90% | 15,90% | 24,09 | yes | no | NA |
| JAL7480 | 97,60% | 12,10% | 15,62 | yes | yes | yes |
| JAL7485 | 96,90% | 10,70% | 24,96 | yes | no | NA |
| JAL7486 | 97,60% | 10,70% | 21,96 | yes | yes | yes |
| JAL7491 | 97,60% | 12,60% | 27,77 | yes | yes | yes |
| JAL7492 | 97,60% | 11,70% | 21,31 | yes | yes | yes |
| L058 | 96,20% | 14,60% | 18,47 | yes | no | NA |
| L166 | 95,70% | 14,20% | 23,01 | yes | yes | yes |
| L186 | 97,50% | 12,40% | 18,73 | yes | no | NA |
| L352 | 97,30% | 16,70% | 25,11 | yes | no | NA |
| L781 | 97,20% | 16,60% | 19,34 | yes | no | NA |
| L800 | 96,90% | 13,50% | 20,47 | yes | no | NA |
| M2C3H | 97,60% | 14,10% | 26,41 | yes | no | NA |
| M2CHL | 97,50% | 18,80% | 18,85 | yes | no | NA |
| M2E71 | 97,30% | 15,80% | 21,12 | yes | no | NA |
| MSV2AC | 97,00% | 17,40% | 25,59 | yes | yes | yes |
| W1433 | 94,60% | 22,00% | 20,37 | yes | no | NA |
| W1551 | 91,60% | 23,00% | 13,05 | yes | yes | yes |
| W1593 | 90,80% | 22,10% | 17,69 | yes | yes | yes |
| W1869 | 95,40% | 23,20% | 15,76 | yes | no | NA |
| W191598 | 96,80% | 24,80% | 19,19 | yes | no | NA |
| W200546 | 96,10% | 20,40% | 19,50 | yes | no | NA |
| W2030 | 95,70% | 22,70% | 18,07 | yes | yes | yes |
| W2124 | 95,40% | 23,30% | 18,92 | yes | yes | yes |
| W2185 | 96,50% | 22,60% | 18,12 | yes | yes | yes |
| W2192 | 92,40% | 22,20% | 17,05 | yes | no | NA |
| W2193 | 95,40% | 21,10% | 21,84 | yes | yes | yes |
| W224957 | 97,00% | 19,30% | 17,45 | yes | no | NA |
| W2265 | 97,00% | 21,30% | 19,15 | yes | yes | yes |
| W226855 | 95,20% | 25,30% | 24,18 | yes | no | NA |
| W2280 | 96,80% | 24,40% | 23,33 | yes | yes | yes |
| W2291 | 96,90% | 21,90% | 18,10 | yes | yes | yes |

| | | | | | | |
|---|---|---|---|---|---|---|
| W2371F | 88,70% | 18,00% | 12,45 | yes | yes | yes |
| W2419 | 87,30% | 10,30% | 16,71 | yes | no | yes |
| W2430F | 93,20% | 12,90% | 13,76 | yes | yes | yes |
| W2470 | 86,20% | 11,70% | 11,51 | no | NA | NA |
| W2487F | 92,70% | 11,90% | 15,32 | yes | yes | yes |
| W2499M | 93,10% | 9,40% | 2,18 | no | NA | NA |
| W2500M | 88,80% | 12,70% | 19,50 | no | NA | NA |
| W2501F | 61,00% | 12,10% | 0,76 | no | NA | NA |
| W2594M | 96,70% | 23,00% | 17,22 | yes | yes | yes |
| W2623M | 96,70% | 23,00% | 20,33 | yes | yes | yes |
| W2685F | 83,30% | 18,60% | 14,76 | no | NA | NA |
| W2707M | 90,30% | 22,20% | 14,85 | yes | yes | yes |
| W2725 | 96,50% | 22,30% | 17,63 | yes | no | NA |
| W2728 | 61,50% | 24,00% | 6,32 | no | NA | NA |
| W2878M | 96,50% | 18,10% | 20,37 | yes | yes | yes |
| W2883F | 96,20% | 18,60% | 20,62 | yes | yes | yes |
| W2886 | 80,80% | 17,60% | 21,75 | yes | yes | yes |
| W2910 | 96,90% | 20,40% | 19,73 | yes | yes | yes |
| W2951 | 97,30% | 20,30% | 19,45 | yes | yes | yes |
| w361703 | 97,20% | 21,40% | 22,29 | yes | yes | yes |
| w361756 | 97,30% | 20,90% | 24,11 | yes | yes | yes |
| w484049 | 96,70% | 21,00% | 19,72 | yes | no | NA |
| w4902419 | 97,10% | 22,60% | 27,50 | yes | no | NA |
| w4926402 | 97,00% | 20,70% | 18,07 | yes | yes | yes |
| WTR025 | 97,20% | 17,60% | 19,39 | yes | yes | no |
| WTR026 | 97,40% | 11,60% | 23,93 | yes | yes | yes |
| WTR027 | 97,30% | 15,70% | 19,76 | yes | yes | no |
| WTR028 | 97,50% | 16,00% | 18,91 | yes | yes | yes |
| WTR029 | 97,30% | 16,00% | 29,84 | yes | yes | yes |
| WTR030 | 97,40% | 17,20% | 23,84 | yes | yes | yes |
| WTR031 | 97,20% | 16,30% | 23,73 | yes | yes | yes |
| WTR032 | 97,20% | 14,60% | 22,94 | yes | yes | yes |
| WTR033 | 97,10% | 14,40% | 23,77 | yes | yes | yes |
| WTR034 | 97,50% | 16,60% | 25,79 | yes | yes | yes |
| WTR035 | 97,30% | 14,70% | 25,71 | yes | yes | yes |

| | | | | | | |
|---|---|---|---|---|---|---|
| WTR037 | 97,10% | 15,40% | 27,74 | yes | yes | yes |
| WTR038 | 97,40% | 14,30% | 23,01 | yes | yes | yes |
| WTR039 | 97,30% | 17,00% | 24,65 | yes | yes | no |
| WTR040 | 97,30% | 17,10% | 24,60 | yes | yes | yes |
| WTR041 | 97,20% | 16,80% | 24,68 | yes | yes | yes |
| W1129 | 97,70% | 25,01% | 17,46 | yes | no | NA |
| W1208 | 97,28% | 32,73% | 25,68 | yes | no | NA |
| W1297 | 95,91% | 25,52% | 21,68 | yes | no | NA |
| W1431 | 98,12% | 28,52% | 22,43 | yes | no | NA |
| W1432 | 97,80% | 24,08% | 18,16 | yes | no | NA |
| W1449 | 95,01% | 28,33% | 18,44 | yes | no | NA |
| W1564 | 97,33% | 26,70% | 19,41 | yes | no | NA |
| W1578 | 89,93% | 14,95% | 14,13 | yes | no | NA |

**Table S2.** Information about previously published genomes analyzed in this study.

| **VCF_ID** | **BioProject** | **BioSample** | **Species** |
|---|---|---|---|
| 140447_S11 | PRJNA448733 | SAMN08873033 | *Canis familiaris* |
| 149323_S6 | PRJNA448733 | SAMN08873141 | *Canis familiaris* |
| 165414_S20 | PRJNA448733 | SAMN08873243 | *Canis familiaris* |
| 171515_S19 | PRJNA448733 | SAMN08873245 | *Canis familiaris* |
| 173006_S10 | PRJNA448733 | SAMN08873036 | *Canis familiaris* |
| 173486_S3 | PRJNA448733 | SAMN08873146 | *Canis familiaris* |
| AKBH000001 | PRJNA648123 | SAMN29845569 | *Canis familiaris* |
| AKBH000003 | PRJNA648123 | SAMN29845571 | *Canis familiaris* |
| AMAL000001 | PRJNA648123 | SAMN15625708 | *Canis familiaris* |
| AMAL000002 | PRJNA648123 | SAMN19522718 | *Canis familiaris* |
| AMST000001 | PRJNA648123 | SAMN15625709 | *Canis familiaris* |
| AMST000006 | PRJNA648123 | SAMN20151360 | *Canis familiaris* |
| ANAT000003 | PRJNA188158 | SAMN21036208 | *Canis familiaris* |
| ANAT000004 | PRJNA648123 | SAMN20151362 | *Canis familiaris* |
| Arctic_BaffinIsl_CD130_RKW7639 | PRJNA512209 | SAMN10662574 | *Canis lupus* |
| Arctic_EllesmereIsl_GF44_RKW7640 | PRJNA512209 | SAMN10662575 | *Canis lupus* |
| Arctic_Nunavut_CB177_RKW7649 | PRJNA512209 | SAMN10662576 | *Canis lupus* |
| Arctic_VictoriaIsl_CB215_RKW7619 | PRJNA512209 | SAMN10662573 | *Canis lupus* |
| AUSS000001 | PRJNA648123 | SAMN15625719 | *Canis familiaris* |
| AUSS000003 | PRJNA648123 | SAMN20151355 | *Canis familiaris* |
| BEAU000004 | PRJNA648123 | SAMN20151377 | *Canis familiaris* |
| BEAU000007 | PRJNA648123 | SAMN19522750 | *Canis familiaris* |
| BELS000001 | PRJNA648123 | SAMN15625732 | *Canis familiaris* |
| BELS000003 | PRJNA648123 | SAMN20151397 | *Canis familiaris* |
| BGI-01505070001 | PRJNA448733 | SAMN03653003 | *Canis lupus* |
| BGI-01505070002 | PRJNA448733 | SAMN03652998 | *Canis lupus* |
| BGI-01505070003 | PRJNA448733 | SAMN03653001 | *Canis lupus* |
| BGI-01505070004 | PRJNA448733 | SAMN03653004 | *Canis lupus* |
| BGI-01505070005 | PRJNA448733 | SAMN03652999 | *Canis lupus* |
| BGI-01505070006 | PRJNA448733 | SAMN03653002 | *Canis lupus* |
| BGI-01505070007 | PRJNA448733 | SAMN03653000 | *Canis lupus* |
| BGI-01505070008 | PRJNA448733 | SAMN03652997 | *Canis lupus* |
| BLDH000002 | PRJNA648123 | SAMN21035065 | *Canis familiaris* |

| | | | |
|---|---|---|---|
| BLDH000003 | PRJNA648123 | SAMN19522761 | *Canis familiaris* |
| BMAL000001 | PRJNA648123 | SAMN15625737 | *Canis familiaris* |
| BMAL000002 | PRJNA648123 | SAMN15625738 | *Canis familiaris* |
| BMSH000005 | PRJNA648123 | SAMN21035071 | *Canis familiaris* |
| BMSH000006 | PRJNA648123 | SAMN21035072 | *Canis familiaris* |
| BOER000002 | PRJNA648123 | SAMN15625740 | *Canis familiaris* |
| BOER000003 | PRJNA648123 | SAMN29845199 | *Canis familiaris* |
| BOXR000001 | PRJNA648123 | SAMN15625743 | *Canis familiaris* |
| BOXR000005 | PRJNA648123 | SAMN20151370 | *Canis familiaris* |
| BRAC000001 | PRJNA648123 | SAMN20151407 | *Canis familiaris* |
| BRAC000004 | PRJNA648123 | SAMN20151410 | *Canis familiaris* |
| BRIA000005 | PRJNA648123 | SAMN20151403 | *Canis familiaris* |
| BRIA000007 | PRJNA648123 | SAMN19522776 | *Canis familiaris* |
| BRMD000001 | PRJNA648123 | SAMN15625755 | *Canis familiaris* |
| BRMD000004 | PRJNA648123 | SAMN19522786 | *Canis familiaris* |
| BRTR000002 | PRJNA648123 | SAMN21035084 | *Canis familiaris* |
| BRTR000004 | PRJNA648123 | SAMN21035086 | *Canis familiaris* |
| CAUC000001 | PRJNA648123 | SAMN15625779 | *Canis familiaris* |
| CCRT000003 | PRJNA648123 | SAMN21035098 | *Canis familiaris* |
| CCRT000004 | PRJNA648123 | SAMN21035099 | *Canis familiaris* |
| CHBR000001 | PRJNA648123 | SAMN15625780 | *Canis familiaris* |
| CHBR000002 | PRJNA648123 | SAMN15625781 | *Canis familiaris* |
| CLUPAZ000001 | PRJNA648123 | SAMN29845311 | *Canis lupus* |
| CLUPCN000001 | PRJNA648123 | SAMN29845312 | *Canis lupus* |
| CLUPCN000002 | PRJNA648123 | SAMN29845313 | *Canis lupus* |
| CLUPCN000003 | PRJNA648123 | SAMN29845314 | *Canis lupus* |
| CLUPCN000004 | PRJNA648123 | SAMN29845315 | *Canis lupus* |
| CLUPCN000005 | PRJNA648123 | SAMN29845316 | *Canis lupus* |
| CLUPCN000006 | PRJNA648123 | SAMN29845317 | *Canis lupus* |
| CLUPCN000007 | PRJNA648123 | SAMN29845318 | *Canis lupus* |
| CLUPCN000008 | PRJNA648123 | SAMN29845319 | *Canis lupus* |
| CLUPCN000009 | PRJNA648123 | SAMN29845320 | *Canis lupus* |
| CLUPCN000010 | PRJNA648123 | SAMN29845321 | *Canis lupus* |
| CLUPIR000001 | PRJNA648123 | SAMN29845324 | *Canis lupus* |
| CLUPIR000002 | PRJNA648123 | SAMN29845325 | *Canis lupus* |

| | | | |
|---|---|---|---|
| CLUPIR000003 | PRJNA648123 | SAMN29845326 | *Canis lupus* |
| CLUPIR000004 | PRJNA648123 | SAMN29845327 | *Canis lupus* |
| CLUPIR000005 | PRJNA648123 | SAMN29845328 | *Canis lupus* |
| CLUPIR000006 | PRJNA648123 | SAMN29845329 | *Canis lupus* |
| CLUPKG000001 | PRJNA648123 | SAMN29845331 | *Canis lupus* |
| CLUPKZ000002 | PRJNA648123 | SAMN29845332 | *Canis lupus* |
| CLUPRU000001 | PRJNA648123 | SAMN29845333 | *Canis lupus* |
| CLUPRU000002 | PRJNA648123 | SAMN29845334 | *Canis lupus* |
| CLUPRU000003 | PRJNA648123 | SAMN29845335 | *Canis lupus* |
| CLUPRU000004 | PRJNA648123 | SAMN29845336 | *Canis lupus* |
| CLUPRU000005 | PRJNA648123 | SAMN29845337 | *Canis lupus* |
| CLUPRU000006 | PRJNA648123 | SAMN29845338 | *Canis lupus* |
| CLUPRU000007 | PRJNA648123 | SAMN29845339 | *Canis lupus* |
| CLUPRU000008 | PRJNA648123 | SAMN29845340 | *Canis lupus* |
| CLUPRU000018 | PRJNA648123 | SAMN29845347 | *Canis lupus* |
| CLUPRU000019 | PRJNA648123 | SAMN29845348 | *Canis lupus* |
| CLUPRU000020 | PRJNA648123 | SAMN29845349 | *Canis lupus* |
| CLUPTJ000001 | PRJNA648123 | SAMN29845350 | *Canis lupus* |
| CLUPTJ000002 | PRJNA648123 | SAMN29845351 | *Canis lupus* |
| CLUPTJ000003 | PRJNA648123 | SAMN29845352 | *Canis lupus* |
| CLUPTJ000004 | PRJNA648123 | SAMN29845353 | *Canis lupus* |
| CLUPTJ000005 | PRJNA648123 | SAMN29845354 | *Canis lupus* |
| CLUPTJ000006 | PRJNA648123 | SAMN29845355 | *Canis lupus* |
| CLUPTJ000007 | PRJNA648123 | SAMN29845356 | *Canis lupus* |
| CNCS000003 | PRJNA648123 | SAMN20151428 | *Canis familiaris* |
| CNCS000004 | PRJNA648123 | SAMN19522798 | *Canis familiaris* |
| COLL000007 | PRJNA648123 | SAMN20151440 | *Canis familiaris* |
| COLL000013 | PRJNA648123 | SAMN19522799 | *Canis familiaris* |
| COOK000002 | PRJNA648123 | SAMN15625802 | *Canis familiaris* |
| COOK000007 | PRJNA648123 | SAMN15625807 | *Canis familiaris* |
| CZEC000003 | PRJNA648123 | SAMN29949170 | *Canis familiaris* |
| CZWO000001 | PRJNA648123 | SAMN20151458 | *Canis familiaris* |
| D-00-12 | PRJEB20635 | SAMEA104090975 | *Canis lupus* |
| D-06-14 | PRJEB20635 | SAMEA104090980 | *Canis lupus* |
| D-07-16 | PRJEB20635 | SAMEA104090983 | *Canis lupus* |

| | | | |
|---|---|---|---|
| D-08-10 | PRJEB20635 | SAMEA104090988 | *Canis lupus* |
| D-10-68 | PRJEB20635 | SAMEA104091000 | *Canis lupus* |
| D-11-58 | PRJEB20635 | SAMEA104091003 | *Canis lupus* |
| D-85-02 | PRJEB20635 | SAMEA104091008 | *Canis lupus* |
| DALM000002 | PRJNA648123 | SAMN15625820 | *Canis familiaris* |
| Dalmatian01 | PRJNA448733 | SAMN08872976 | *Canis familiaris* |
| DEER000003 | PRJNA648123 | SAMN21035120 | *Canis familiaris* |
| DEER000004 | PRJNA648123 | SAMN19522813 | *Canis familiaris* |
| Dhole_BerlinZoo | PRJNA494815 | SAMN10180424 | *Cuon alpinus* |
| EFXH000002 | PRJNA648123 | SAMN15625831 | *Canis familiaris* |
| EFXH000003 | PRJNA648123 | SAMN15625832 | *Canis familiaris* |
| ENTB000001 | PRJNA648123 | SAMN15625833 | *Canis familiaris* |
| ENTB000002 | PRJNA648123 | SAMN15625834 | *Canis familiaris* |
| ESMD000002 | PRJNA648123 | SAMN20151483 | *Canis familiaris* |
| ESMD000003 | PRJNA648123 | SAMN20151484 | *Canis familiaris* |
| FlatcoatedRetriever03 | PRJNA448733 | SAMN08873011 | *Canis familiaris* |
| FLCR000002 | PRJNA648123 | SAMN15625853 | *Canis familiaris* |
| G100-12 | PRJEB20635 | SAMEA104091023 | *Canis lupus* |
| G100-14 | PRJEB20635 | SAMEA104091024 | *Canis lupus* |
| G109-11 | PRJEB20635 | SAMEA104091026 | *Canis lupus* |
| G126-13 | PRJEB20635 | SAMEA104091029 | *Canis lupus* |
| G139-12 | PRJEB20635 | SAMEA104091030 | *Canis lupus* |
| G31-13 | PRJEB20635 | SAMEA104091035 | *Canis lupus* |
| G37-10 | PRJEB20635 | SAMEA104091039 | *Canis lupus* |
| G50-12 | PRJEB20635 | SAMEA104091041 | *Canis lupus* |
| GALG000002 | PRJNA648123 | SAMN21035164 | *Canis familiaris* |
| GALG000003 | PRJNA648123 | SAMN21035165 | *Canis familiaris* |
| glw | PRJNA255370 | SAMN02921310 | *Canis lupus* |
| GORD000002 | PRJNA648123 | SAMN21035174 | *Canis familiaris* |
| GPYR000003 | PRJNA648123 | SAMN21035177 | *Canis familiaris* |
| GPYR000004 | PRJNA648123 | SAMN21035178 | *Canis familiaris* |
| GreaterSwissMountainDog01 | PRJNA448733 | SAMN08873062 | *Canis familiaris* |
| GreyWolf_AtlanticCoast | PRJNA496590 | SAMN10246086 | *Canis lupus* |
| GreyWolf_BaffinNorth | PRJNA496590 | SAMN10246087 | *Canis lupus* |
| GreyWolf_BaffinSouth | PRJNA496590 | SAMN10246088 | *Canis lupus* |

| | | | |
|---|---|---|---|
| GreyWolf_BanksIsland | PRJNA496590 | SAMN10246089 | *Canis lupus* |
| GreyWolf_EllesmereIsland | PRJNA494815 | SAMN10180431 | *Canis lupus* |
| GreyWolf_Greenland | PRJNA494815 | SAMN10180430 | *Canis lupus* |
| GreyWolf_Montana | PRJEB31639 | SAMEA7091403 | *Canis lupus* |
| GreyWolf_PacificCoast | PRJNA496590 | SAMN10246091 | *Canis lupus* |
| GreyWolf_StLawrenceIsland | PRJNA496590 | SAMN10246094 | *Canis lupus* |
| GreyWolf_Toronto | PRJNA496590 | SAMN10246095 | *Canis lupus* |
| GreyWolf_VictoriaIsland | PRJNA496590 | SAMN10246096 | *Canis lupus* |
| GRSD000002 | PRJNA648123 | SAMN15625866 | *Canis familiaris* |
| GRSD000003 | PRJNA648123 | SAMN15625867 | *Canis familiaris* |
| GSMD000004 | PRJNA648123 | SAMN19522863 | *Canis familiaris* |
| HOVA000003 | PRJNA648123 | SAMN21035194 | *Canis familiaris* |
| HOVA000004 | PRJNA648123 | SAMN19522869 | *Canis familiaris* |
| HWHP000001 | PRJNA648123 | SAMN21035204 | *Canis familiaris* |
| IBIZ000010 | PRJNA648123 | SAMN21035207 | *Canis familiaris* |
| IBIZ000011 | PRJNA648123 | SAMN21035208 | *Canis familiaris* |
| inw | PRJNA255370 | SAMN02921311 | *Canis lupus* |
| Iran_wolf | PRJNA494719 | SAMN10174939 | *Canis lupus* |
| IRSE000001 | PRJNA648123 | SAMN15625886 | *Canis familiaris* |
| IRSE000003 | PRJNA648123 | SAMN21035215 | *Canis familiaris* |
| irw | PRJNA255370 | SAMN02921312 | *Canis lupus* |
| IsleRoyaleNP_CL141_JRRW018 | PRJNA512209 | SAMN10662572 | *Canis lupus* |
| IsleRoyaleNP_CL189_RWJR008 | PRJNA512209 | SAMN10662584 | *Canis lupus* |
| IsleRoyaleNP_CL61_RWJR005 | PRJNA512209 | SAMN10662582 | *Canis lupus* |
| IWSP000001 | PRJNA648123 | SAMN15625887 | *Canis familiaris* |
| IWSP000002 | PRJNA648123 | SAMN15625888 | *Canis familiaris* |
| KANG000001 | PRJNA648123 | SAMN29845575 | *Canis familiaris* |
| KANG000003 | PRJNA648123 | SAMN29845577 | *Canis familiaris* |
| KARS000005 | PRJNA648123 | SAMN29845582 | *Canis familiaris* |
| KARS000006 | PRJNA648123 | SAMN29845583 | *Canis familiaris* |
| KUVZ000002 | PRJNA648123 | SAMN15626060 | *Canis familiaris* |
| KUVZ000007 | PRJNA648123 | SAMN21035243 | *Canis familiaris* |
| L253 | PRJNA1078274 | SAMN39993575 | *Canis lupus* |
| L409 | PRJNA1078274 | SAMN39993577 | *Canis lupus* |
| L514 | PRJNA1078274 | SAMN39993578 | *Canis lupus* |

| | | | |
|---|---|---|---|
| L547 | PRJNA1078274 | SAMN39993579 | *Canis lupus* |
| L552 | PRJNA1078274 | SAMN39993580 | *Canis lupus* |
| L588 | PRJNA1078274 | SAMN39993581 | *Canis lupus* |
| L844 | PRJNA1078274 | SAMN39993582 | *Canis lupus* |
| LAGO000001 | PRJNA648123 | SAMN15625893 | *Canis familiaris* |
| LAGO000004 | PRJNA648123 | SAMN15625896 | *Canis familiaris* |
| Lcu2_Pastora | PRJNA448733 | SAMN02487034 | *Lycalopex culpaeus* |
| LEOP000003 | PRJNA648123 | SAMN15625900 | *Canis familiaris* |
| LEOP000008 | PRJNA648123 | SAMN15625904 | *Canis familiaris* |
| LUP006817 | PRJNA648123 | SAMN17023496 | *Canis lupus* |
| LUP006818 | PRJNA648123 | SAMN17023497 | *Canis lupus* |
| LUP006819 | PRJNA648123 | SAMN17023498 | *Canis lupus* |
| LUP006820 | PRJNA648123 | SAMN17023499 | *Canis lupus* |
| LUP006821 | PRJNA648123 | SAMN17023500 | *Canis lupus* |
| LUP006822 | PRJNA648123 | SAMN17023501 | *Canis lupus* |
| LUP006823 | PRJNA648123 | SAMN17023502 | *Canis lupus* |
| LUP006824 | PRJNA648123 | SAMN17023503 | *Canis lupus* |
| LUP006825 | PRJNA648123 | SAMN17023504 | *Canis lupus* |
| LUP006826 | PRJNA648123 | SAMN17023505 | *Canis lupus* |
| LUP006827 | PRJNA648123 | SAMN17023506 | *Canis lupus* |
| LUP006828 | PRJNA648123 | SAMN17023507 | *Canis lupus* |
| LUP006829 | PRJNA648123 | SAMN17023514 | *Canis lupus* |
| LUP006830 | PRJNA648123 | SAMN17023515 | *Canis lupus* |
| LUPWCHN00003 | PRJNA448733 | SAMN03168394 | *Canis lupus* |
| LUPWCHN00010 | PRJNA448733 | SAMN03168399 | *Canis lupus* |
| LUPZCHN00006 | PRJNA448733 | SAMN03168396 | *Canis lupus* |
| LUPZCHN00009 | PRJNA448733 | SAMN03168398 | *Canis lupus* |
| LUPZCHN00013 | PRJNA448733 | SAMN03168400 | *Canis lupus* |
| M-01-06 | PRJEB20635 | SAMEA104091051 | *Canis lupus* |
| M-06-03 | PRJEB20635 | SAMEA104091058 | *Canis lupus* |
| M-09-05 | PRJEB20635 | SAMEA104091063 | *Canis lupus* |
| M-10-04 | PRJEB20635 | SAMEA104091065 | *Canis lupus* |
| M-98-08 | PRJEB20635 | SAMEA104091071 | *Canis lupus* |
| Maremma01 | PRJNA685036 | SAMN17192085 | *Canis familiaris* |
| MARM000006 | PRJNA648123 | SAMN21035278 | *Canis familiaris* |

| | | | |
|---|---|---|---|
| MastinoAbruzzese01 | PRJNA685036 | SAMN17192086 | *Canis familiaris* |
| Mexican_wolf | PRJNA494719 | SAMN10174942 | *Canis lupus* |
| Minnesota_RKW119_RWJR007 | PRJNA512209 | SAMN10662583 | *Canis lupus* |
| Minnesota_RKW2515_RWJR016 | PRJNA512209 | SAMN10662591 | *Canis lupus* |
| Minnesota_RKW2518_RWJR012 | PRJNA512209 | SAMN10662588 | *Canis lupus* |
| Minnesota_RKW2523_RWJR009 | PRJNA512209 | SAMN10662585 | *Canis lupus* |
| Minnesota_RKW2524_RWJR003 | PRJNA512209 | SAMN10662580 | *Canis lupus* |
| Minnesota_wolf | PRJNA494719 | SAMN10174943 | *Canis lupus* |
| Mongolian_wolf | PRJNA494719 | SAMN10174944 | *Canis lupus* |
| MW122 | PRJEB57290 | SAMEA117560979 | *Canis lupus* |
| MW127 | PRJEB57290 | SAMEA117560980 | *Canis lupus* |
| mxa | PRJNA255370 | SAMN02921314 | *Canis lupus* |
| NEAP000004 | PRJNA648123 | SAMN19522885 | *Canis familiaris* |
| NEAP000005 | PRJNA648123 | SAMN19522886 | *Canis familiaris* |
| NELK000002 | PRJNA648123 | SAMN15625929 | *Canis familiaris* |
| NEWF000002 | PRJNA648123 | SAMN15625932 | *Canis familiaris* |
| NEWF000006 | PRJNA648123 | SAMN19522887 | *Canis familiaris* |
| NorwegianElkhound02 | PRJNA448733 | SAMN08873199 | *Canis familiaris* |
| Penelope | PRJNA476541 | SAMN04851099 | *Canis lupus* |
| PITB000001 | PRJNA648123 | SAMN21035342 | *Canis familiaris* |
| PITB000002 | PRJNA648123 | SAMN21035343 | *Canis familiaris* |
| POPT000001 | PRJNA648123 | SAMN21035354 | *Canis familiaris* |
| PortugueseWaterDog09 | PRJNA448733 | SAMN08873227 | *Canis familiaris* |
| POSD000005 | PRJNA648123 | SAMN21035357 | *Canis familiaris* |
| PPOD000004 | PRJNA648123 | SAMN21035363 | *Canis familiaris* |
| PPOP000001 | PRJNA648123 | SAMN15625954 | *Canis familiaris* |
| PT61 | PRJNA448733 | SAMN02485604 | *Canis familiaris* |
| ptw | PRJNA255370 | SAMN02921316 | *Canis lupus* |
| PTWD000008 | PRJNA648123 | SAMN21035373 | *Canis familiaris* |
| PYMF000005 | PRJNA648123 | SAMN21035387 | *Canis familiaris* |
| PYMF000006 | PRJNA648123 | SAMN21035388 | *Canis familiaris* |
| Qinghai_wolf | PRJNA494719 | SAMN10174946 | *Canis lupus* |
| Quebec_MontTremblantNP_voyou0833M_RWBH001 | PRJNA512209 | SAMN10662577 | *Canis lupus* |
| RHOD000001 | PRJNA648123 | SAMN15625962 | *Canis familiaris* |
| RKW13451 | PRJNA274504 | SAMN03366711 | *Canis lupus* |

| | | | |
|---|---|---|---|
| RKW481_SF5 | PRJNA488093 | SAMN09924608 | *Lycaon pictus* |
| SAAR000002 | PRJNA648123 | SAMN21035401 | *Canis familiaris* |
| SAAR000005 | PRJNA648123 | SAMN19522962 | *Canis familiaris* |
| SALU000001 | PRJNA648123 | SAMN29845357 | *Canis familiaris* |
| SALU000002 | PRJNA648123 | SAMN29845584 | *Canis familiaris* |
| SAMO000002 | PRJNA648123 | SAMN15625967 | *Canis familiaris* |
| Samoyed01 | PRJNA448733 | SAMN08873258 | *Canis familiaris* |
| SGGO000002 | PRJNA648123 | SAMN21035410 | *Canis familiaris* |
| SGGO000004 | PRJNA648123 | SAMN21035412 | *Canis familiaris* |
| SLOU000001 | PRJNA648123 | SAMN29845586 | *Canis familiaris* |
| SLOU000003 | PRJNA648123 | SAMN21035430 | *Canis familiaris* |
| SPIN000001 | PRJNA648123 | SAMN15625998 | *Canis familiaris* |
| SPIN000004 | PRJNA648123 | SAMN15626001 | *Canis familiaris* |
| SPMT000002 | PRJNA648123 | SAMN21035437 | *Canis familiaris* |
| SPMT000005 | PRJNA648123 | SAMN19522932 | *Canis familiaris* |
| spw | PRJNA255370 | SAMN02921319 | *Canis lupus* |
| SPWD000002 | PRJNA648123 | SAMN20151489 | *Canis familiaris* |
| SPWD000003 | PRJNA648123 | SAMN19522934 | *Canis familiaris* |
| SRAB000001 | PRJNA648123 | SAMN29845358 | *Canis familiaris* |
| SRAB000002 | PRJNA648123 | SAMN29845359 | *Canis familiaris* |
| TERV000005 | PRJNA648123 | SAMN20151402 | *Canis familiaris* |
| TOSA000002 | PRJNA648123 | SAMN15626025 | *Canis familiaris* |
| TOSA000003 | PRJNA648123 | SAMN29845226 | *Canis familiaris* |
| TURM000002 | PRJNA648123 | SAMN29845591 | *Canis familiaris* |
| TURM000004 | PRJNA648123 | SAMN29845593 | *Canis familiaris* |
| TURV000001 | PRJNA648123 | SAMN15626026 | *Canis familiaris* |
| TWCH000001 | PRJNA648123 | SAMN15626028 | *Canis familiaris* |
| TWCH000002 | PRJNA648123 | SAMN15626029 | *Canis familiaris* |
| V113 | PRJEB39198 | SAMEA7038708 | *Canis lupus* |
| V114 | PRJEB39198 | SAMEA7038709 | *Canis lupus* |
| V115 | PRJEB39198 | SAMEA7038710 | *Canis lupus* |
| V116 | PRJEB39198 | SAMEA7038711 | *Canis lupus* |
| V117 | PRJEB39198 | SAMEA7038712 | *Canis lupus* |
| V119 | PRJEB39198 | SAMEA7038713 | *Canis lupus* |
| V120 | PRJEB39198 | SAMEA7038714 | *Canis lupus* |

| | | | |
|---|---|---|---|
| V126 | PRJEB39198 | SAMEA7038715 | *Canis lupus* |
| V132 | PRJEB39198 | SAMEA7038716 | *Canis lupus* |
| V134 | PRJEB39198 | SAMEA7038717 | *Canis lupus* |
| V136 | PRJEB39198 | SAMEA7038718 | *Canis lupus* |
| V141 | PRJEB39198 | SAMEA7038720 | *Canis lupus* |
| V143 | PRJEB39198 | SAMEA7038722 | *Canis lupus* |
| V3064 | PRJEB39198 | ERS4802915 | *Canis lupus* |
| V3065 | PRJEB39198 | ERS4802916 | *Canis lupus* |
| V3069 | PRJEB39198 | ERS4802917 | *Canis lupus* |
| VILLAF000001 | PRJNA648123 | SAMN29845227 | *Canis familiaris* |
| VILLAF000002 | PRJNA648123 | SAMN29845228 | *Canis familiaris* |
| VILLAF000003 | PRJNA648123 | SAMN29845229 | *Canis familiaris* |
| VILLAZ000002 | PRJNA648123 | SAMN29845596 | *Canis familiaris* |
| VILLAZ000003 | PRJNA648123 | SAMN29845597 | *Canis familiaris* |
| VILLAZ000004 | PRJNA648123 | SAMN29845598 | *Canis familiaris* |
| VILLAZ000005 | PRJNA648123 | SAMN29845599 | *Canis familiaris* |
| VILLAZ000007 | PRJNA648123 | SAMN29845601 | *Canis familiaris* |
| VILLBG000001 | PRJNA648123 | SAMN29845233 | *Canis familiaris* |
| VILLBG000002 | PRJNA648123 | SAMN29845234 | *Canis familiaris* |
| VILLCG000001 | PRJNA648123 | SAMN29845239 | *Canis familiaris* |
| VILLCG000004 | PRJNA648123 | SAMN29845242 | *Canis familiaris* |
| VILLCG000006 | PRJNA648123 | SAMN29845244 | *Canis familiaris* |
| VILLCG000011 | PRJNA648123 | SAMN29845603 | *Canis familiaris* |
| VILLCG000013 | PRJNA648123 | SAMN29845605 | *Canis familiaris* |
| VILLCN000005 | PRJNA648123 | SAMN29845364 | *Canis familiaris* |
| VILLCN000013 | PRJNA648123 | SAMN29845372 | *Canis familiaris* |
| VILLCN000088 | PRJNA648123 | SAMN29845447 | *Canis familiaris* |
| VILLCN000145 | PRJNA648123 | SAMN29845504 | *Canis familiaris* |
| VILLCN000153 | PRJNA648123 | SAMN29845512 | *Canis familiaris* |
| VILLIR000021 | PRJNA648123 | SAMN29845527 | *Canis familiaris* |
| VILLIR000023 | PRJNA648123 | SAMN29845529 | *Canis familiaris* |
| VILLIR000024 | PRJNA648123 | SAMN29845530 | *Canis familiaris* |
| VILLIR000026 | PRJNA648123 | SAMN29845532 | *Canis familiaris* |
| VILLIR000027 | PRJNA648123 | SAMN29845533 | *Canis familiaris* |
| VILLKE000002 | PRJNA648123 | SAMN29845538 | *Canis familiaris* |

| | | | |
|---|---|---|---|
| VILLKE000009 | PRJNA648123 | SAMN29845545 | *Canis familiaris* |
| VILLKE000010 | PRJNA648123 | SAMN29845546 | *Canis familiaris* |
| VILLKE000011 | PRJNA648123 | SAMN29845547 | *Canis familiaris* |
| VILLKE000019 | PRJNA648123 | SAMN29845555 | *Canis familiaris* |
| VILLLR000005 | PRJNA648123 | SAMN29845272 | *Canis familiaris* |
| VILLLR000011 | PRJNA648123 | SAMN29845278 | *Canis familiaris* |
| VILLLR000014 | PRJNA648123 | SAMN29845281 | *Canis familiaris* |
| VILLLR000016 | PRJNA648123 | SAMN29845283 | *Canis familiaris* |
| VILLLR000017 | PRJNA648123 | SAMN29845284 | *Canis familiaris* |
| VILLTJ000001 | PRJNA648123 | SAMN29845567 | *Canis familiaris* |
| VILLUZ000002 | PRJNA648123 | SAMN29845627 | *Canis familiaris* |
| VILLUZ000003 | PRJNA648123 | SAMN29845628 | *Canis familiaris* |
| VILLUZ000004 | PRJNA648123 | SAMN29845629 | *Canis familiaris* |
| VILLUZ000005 | PRJNA648123 | SAMN29845630 | *Canis familiaris* |
| VILLUZ000006 | PRJNA648123 | SAMN29845631 | *Canis familiaris* |
| VIZS000007 | PRJNA648123 | SAMN19522941 | *Canis familiaris* |
| W1023 | | | *Canis lupus* |
| W11 | PRJEB28342 | SAMEA4853019 | *Canis lupus* |
| W14 | PRJEB28342 | SAMEA4853022 | *Canis lupus* |
| W1456 | | | *Canis lupus* |
| W16 | PRJEB39198 | SAMEA7038685 | *Canis lupus* |
| W21 | PRJEB28342 | SAMEA4853027 | *Canis lupus* |
| W26 | PRJEB28342 | SAMEA4853031 | *Canis lupus* |
| W29 | PRJEB39198 | SAMEA7038689 | *Canis lupus* |
| W32 | PRJEB39198 | SAMEA7038692 | *Canis lupus* |
| W4 | PRJEB39198 | SAMEA7038695 | *Canis lupus* |
| W44 | PRJEB28342 | SAMEA4853039 | *Canis lupus* |
| W45 | PRJEB39198 | SAMEA7038699 | *Canis lupus* |
| W893 | | | *Canis lupus* |
| WEIM000002 | PRJNA648123 | SAMN15626040 | *Canis familiaris* |
| WEIM000005 | PRJNA648123 | SAMN19522942 | *Canis familiaris* |
| WHPG000001 | PRJNA648123 | SAMN15626046 | *Canis familiaris* |
| WHPG000002 | PRJNA648123 | SAMN15626047 | *Canis familiaris* |
| WIT11 | PRJEB80121 | SAMEA116045426 | *Canis lupus* |
| WIT13 | PRJEB80121 | SAMEA116045428 | *Canis lupus* |

| | | | |
|---|---|---|---|
| WIT14 | PRJEB80121 | SAMEA116045429 | *Canis lupus* |
| WIT16 | PRJEB80121 | SAMEA116045431 | *Canis lupus* |
| WIT17 | PRJEB80121 | SAMEA116045432 | *Canis lupus* |
| WIT18 | PRJEB80121 | SAMEA116045433 | *Canis lupus* |
| WIT19 | PRJEB80121 | SAMEA116045434 | *Canis lupus* |
| WIT20 | PRJEB80121 | SAMEA116045435 | *Canis lupus* |
| WIT21 | PRJEB80121 | SAMEA116045437 | *Canis lupus* |
| WIT22 | PRJEB80121 | SAMEA116045438 | *Canis lupus* |
| WIT23 | PRJEB80121 | SAMEA116045439 | *Canis lupus* |
| wSierraMorena | PRJNA482523 | SAMN09708541 | *Canis lupus* |
| Yellowstone1_wolf | PRJNA494719 | SAMN10174953 | *Canis lupus* |
| Yellowstone2_wolf | PRJNA494719 | SAMN10174954 | *Canis lupus* |
| Yellowstone3_wolf | PRJNA494719 | SAMN10174955 | *Canis lupus* |
| ysa | PRJNA255370 | SAMN02921320 | *Canis lupus* |
| ysb | PRJNA255370 | SAMN02921321 | *Canis lupus* |
| ZERD000002 | PRJNA648123 | SAMN29845641 | *Canis familiaris* |
| ZERD000003 | PRJNA648123 | SAMN29845642 | *Canis familiaris* |
| Kenya_GoldenJackal | PRJNA494719 | SAMN10174941 | *Canis aureus* |
| cac | PRJNA448733 | SAMN02921301 | *Canis latrans* |

**Table S2.** Information about previously published genomes

| VCF_ID | Breed/Country | Country | Continent |
|---|---|---|---|
| 140447_S11 | Golden Retriever | dog | not applicable |
| 149323_S6 | Labrador Retriever | dog | not applicable |
| 165414_S20 | Rottweiler | dog | not applicable |
| 171515_S19 | Rottweiler | dog | not applicable |
| 173006_S10 | Golden Retriever | dog | not applicable |
| 173486_S3 | Labrador Retriever | dog | not applicable |
| AKBH000001 | Akbash | dog | not applicable |
| AKBH000003 | Akbash | dog | not applicable |
| AMAL000001 | Alaskan Malamute | dog | not applicable |
| AMAL000002 | Alaskan Malamute | dog | not applicable |
| AMST000001 | American Staffordshire Terrier | dog | not applicable |
| AMST000006 | American Staffordshire Terrier | dog | not applicable |
| ANAT000003 | Anatolian Shepherd Dog | dog | not applicable |
| ANAT000004 | Anatolian Shepherd Dog | dog | not applicable |
| Arctic_BaffinIsl_CD130_RKW7639 | Canada | Canada | North America |
| Arctic_EllesmereIsl_GF44_RKW7640 | Canada | Canada | North America |
| Arctic_Nunavut_CB177_RKW7649 | Canada | Canada | North America |
| Arctic_VictoriaIsl_CB215_RKW7619 | Canada | Canada | North America |
| AUSS000001 | Australian Shepherd | dog | not applicable |
| AUSS000003 | Australian Shepherd | dog | not applicable |
| BEAU000004 | Beauceron | dog | not applicable |
| BEAU000007 | Beauceron | dog | not applicable |
| BELS000001 | Belgian Sheepdog | dog | not applicable |
| BELS000003 | Belgian Sheepdog | dog | not applicable |
| BGI-01505070001 | Mongolia | Mongolia | Asia |
| BGI-01505070002 | China | China | Asia |
| BGI-01505070003 | China | China | Asia |
| BGI-01505070004 | China | China | Asia |
| BGI-01505070005 | China | China | Asia |
| BGI-01505070006 | China | China | Asia |
| BGI-01505070007 | China | China | Asia |
| BGI-01505070008 | Mongolia | Mongolia | Asia |
| BLDH000002 | Bloodhound | dog | not applicable |

| | | | |
|---|---|---|---|
| BLDH000003 | Bloodhound | dog | not applicable |
| BMAL000001 | Belgian Malinois | dog | not applicable |
| BMAL000002 | Belgian Malinois | dog | not applicable |
| BMSH000005 | Bavarian Mountain Scent Hound | dog | not applicable |
| BMSH000006 | Bavarian Mountain Scent Hound | dog | not applicable |
| BOER000002 | Boerboel | dog | not applicable |
| BOER000003 | Boerboel | dog | not applicable |
| BOXR000001 | Boxer | dog | not applicable |
| BOXR000005 | Boxer | dog | not applicable |
| BRAC000001 | Bracco Italiano | dog | not applicable |
| BRAC000004 | Bracco Italiano | dog | not applicable |
| BRIA000005 | Briard | dog | not applicable |
| BRIA000007 | Briard | dog | not applicable |
| BRMD000001 | Bernese Mountain Dog | dog | not applicable |
| BRMD000004 | Bernese Mountain Dog | dog | not applicable |
| BRTR000002 | Black Russian Terrier | dog | not applicable |
| BRTR000004 | Black Russian Terrier | dog | not applicable |
| CAUC000001 | Caucasian Shepherd Dog | dog | not applicable |
| CCRT000003 | Curly-coated Retriever | dog | not applicable |
| CCRT000004 | Curly-coated Retriever | dog | not applicable |
| CHBR000001 | Chesapeake Bay Retriever | dog | not applicable |
| CHBR000002 | Chesapeake Bay Retriever | dog | not applicable |
| CLUPAZ000001 | Azerbaijan | Azerbaijan | Asia |
| CLUPCN000001 | China | China | Asia |
| CLUPCN000002 | China | China | Asia |
| CLUPCN000003 | China | China | Asia |
| CLUPCN000004 | China | China | Asia |
| CLUPCN000005 | China | China | Asia |
| CLUPCN000006 | China | China | Asia |
| CLUPCN000007 | China | China | Asia |
| CLUPCN000008 | China | China | Asia |
| CLUPCN000009 | China | China | Asia |
| CLUPCN000010 | China | China | Asia |
| CLUPIR000001 | Iran | Iran | Asia |
| CLUPIR000002 | Iran | Iran | Asia |

| | | | |
|---|---|---|---|
| CLUPIR000003 | Iran | Iran | Asia |
| CLUPIR000004 | Iran | Iran | Asia |
| CLUPIR000005 | Iran | Iran | Asia |
| CLUPIR000006 | Iran | Iran | Asia |
| CLUPKG000001 | Kyrgyzstan | Kyrgyzstan | Asia |
| CLUPKZ000002 | Kazakhstan | Kazakhstan | Asia |
| CLUPRU000001 | Asian Russia | Asian Russia | Asia |
| CLUPRU000002 | Asian Russia | Asian Russia | Asia |
| CLUPRU000003 | Asian Russia | Asian Russia | Asia |
| CLUPRU000004 | Asian Russia | Asian Russia | Asia |
| CLUPRU000005 | European Russia | European Russia | Europe |
| CLUPRU000006 | European Russia | European Russia | Europe |
| CLUPRU000007 | European Russia | European Russia | Europe |
| CLUPRU000008 | European Russia | European Russia | Europe |
| CLUPRU000018 | Asian Russia | Asian Russia | Asia |
| CLUPRU000019 | European Russia | European Russia | Europe |
| CLUPRU000020 | European Russia | European Russia | Europe |
| CLUPTJ000001 | Tajikistan | Tajikistan | Asia |
| CLUPTJ000002 | Tajikistan | Tajikistan | Asia |
| CLUPTJ000003 | Tajikistan | Tajikistan | Asia |
| CLUPTJ000004 | Tajikistan | Tajikistan | Asia |
| CLUPTJ000005 | Tajikistan | Tajikistan | Asia |
| CLUPTJ000006 | Tajikistan | Tajikistan | Asia |
| CLUPTJ000007 | Tajikistan | Tajikistan | Asia |
| CNCS000003 | Cane Corso | dog | not applicable |
| CNCS000004 | Cane Corso | dog | not applicable |
| COLL000007 | Collie | dog | not applicable |
| COLL000013 | Collie | dog | not applicable |
| COOK000002 | Chinook | dog | not applicable |
| COOK000007 | Chinook | dog | not applicable |
| CZEC000003 | Czechoslovakian Wolfdog | dog | not applicable |
| CZWO000001 | Czechoslovakian Wolfdog | dog | not applicable |
| D-00-12 | Scandinavia | Scandinavia | Europe |
| D-06-14 | Scandinavia | Scandinavia | Europe |
| D-07-16 | Scandinavia | Scandinavia | Europe |

| | | | |
|---|---|---|---|
| D-08-10 | Scandinavia | Scandinavia | Europe |
| D-10-68 | Scandinavia | Scandinavia | Europe |
| D-11-58 | Scandinavia | Scandinavia | Europe |
| D-85-02 | Scandinavia | Scandinavia | Europe |
| DALM000002 | Dalmatian | dog | not applicable |
| Dalmatian01 | Dalmatian | dog | not applicable |
| DEER000003 | Scottish Deerhound | dog | not applicable |
| DEER000004 | Scottish Deerhound | dog | not applicable |
| Dhole_BerlinZoo | Dhole | outgroup - dhole | not applicable |
| EFXH000002 | English Foxhound | dog | not applicable |
| EFXH000003 | English Foxhound | dog | not applicable |
| ENTB000001 | Entlebucher Mountain Dog | dog | not applicable |
| ENTB000002 | Entlebucher Mountain Dog | dog | not applicable |
| ESMD000002 | Estrela Mountain Dog | dog | not applicable |
| ESMD000003 | Estrela Mountain Dog | dog | not applicable |
| FlatcoatedRetriever03 | Flat-Coated Retriever | dog | not applicable |
| FLCR000002 | Flat-Coated Retriever | dog | not applicable |
| G100-12 | Scandinavia | Scandinavia | Europe |
| G100-14 | Scandinavia | Scandinavia | Europe |
| G109-11 | Scandinavia | Scandinavia | Europe |
| G126-13 | Scandinavia | Scandinavia | Europe |
| G139-12 | Scandinavia | Scandinavia | Europe |
| G31-13 | Scandinavia | Scandinavia | Europe |
| G37-10 | Scandinavia | Scandinavia | Europe |
| G50-12 | Scandinavia | Scandinavia | Europe |
| GALG000002 | Galgo Espanol | dog | not applicable |
| GALG000003 | Galgo Espanol | dog | not applicable |
| glw | US | US | North America |
| GORD000002 | Gordon Setter | dog | not applicable |
| GPYR000003 | Great Pyrenees | dog | not applicable |
| GPYR000004 | Great Pyrenees | dog | not applicable |
| GreaterSwissMountainDog01 | Greater Swiss Mountain Dog | dog | not applicable |
| GreyWolf_AtlanticCoast | Canada | Canada | North America |
| GreyWolf_BaffinNorth | Canada | Canada | North America |
| GreyWolf_BaffinSouth | Canada | Canada | North America |

| | | | |
|---|---|---|---|
| GreyWolf_BanksIsland | Canada | Canada | North America |
| GreyWolf_EllesmereIsland | Canada | Canada | North America |
| GreyWolf_Greenland | Greenland | Greenland | North America |
| GreyWolf_Montana | US | US | North America |
| GreyWolf_PacificCoast | US | US | North America |
| GreyWolf_StLawrenceIsland | US | US | North America |
| GreyWolf_Toronto | Canada | Canada | North America |
| GreyWolf_VictoriaIsland | Canada | Canada | North America |
| GRSD000002 | German Shepherd | dog | not applicable |
| GRSD000003 | German Shepherd | dog | not applicable |
| GSMD000004 | Greater Swiss Mountain Dog | dog | not applicable |
| HOVA000003 | Hovawart | dog | not applicable |
| HOVA000004 | Hovawart | dog | not applicable |
| HWHP000001 | Vizsla | dog | not applicable |
| IBIZ000010 | Ibizan Hound | dog | not applicable |
| IBIZ000011 | Ibizan Hound | dog | not applicable |
| inw | India | India | Asia |
| Iran_wolf | Iran | Iran | Asia |
| IRSE000001 | Irish Setter | dog | not applicable |
| IRSE000003 | Irish Setter | dog | not applicable |
| irw | Iran | Iran | Asia |
| IsleRoyaleNP_CL141_JRRW018 | US | US | North America |
| IsleRoyaleNP_CL189_RWJR008 | US | US | North America |
| IsleRoyaleNP_CL61_RWJR005 | US | US | North America |
| IWSP000001 | Irish Water Spaniel | dog | not applicable |
| IWSP000002 | Irish Water Spaniel | dog | not applicable |
| KANG000001 | Kangal | dog | not applicable |
| KANG000003 | Kangal | dog | not applicable |
| KARS000005 | Kars | dog | not applicable |
| KARS000006 | Kars | dog | not applicable |
| KUVZ000002 | Kuvasz | dog | not applicable |
| KUVZ000007 | Kuvasz | dog | not applicable |
| L253 | Spain | Spain | Europe |
| L409 | Spain | Spain | Europe |
| L514 | Spain | Spain | Europe |

| | | | |
|---|---|---|---|
| L547 | Spain | Spain | Europe |
| L552 | Spain | Spain | Europe |
| L588 | Spain | Spain | Europe |
| L844 | Spain | Spain | Europe |
| LAGO000001 | Lagotto Romagnolo | dog | not applicable |
| LAGO000004 | Lagotto Romagnolo | dog | not applicable |
| Lcu2_Pastora | culpeo | outgroup - culpeo | not applicable |
| LEOP000003 | Catahoula Leopard Dog | dog | not applicable |
| LEOP000008 | Catahoula Leopard Dog | dog | not applicable |
| LUP006817 | Greece | Greece | Europe |
| LUP006818 | Greece | Greece | Europe |
| LUP006819 | Greece | Greece | Europe |
| LUP006820 | Greece | Greece | Europe |
| LUP006821 | Greece | Greece | Europe |
| LUP006822 | Greece | Greece | Europe |
| LUP006823 | Greece | Greece | Europe |
| LUP006824 | Greece | Greece | Europe |
| LUP006825 | Greece | Greece | Europe |
| LUP006826 | Greece | Greece | Europe |
| LUP006827 | Greece | Greece | Europe |
| LUP006828 | Greece | Greece | Europe |
| LUP006829 | Portugal | Portugal | Europe |
| LUP006830 | Portugal | Portugal | Europe |
| LUPWCHN00003 | China | China | Asia |
| LUPWCHN00010 | China | China | Asia |
| LUPZCHN00006 | China | China | Asia |
| LUPZCHN00009 | China | China | Asia |
| LUPZCHN00013 | China | China | Asia |
| M-01-06 | Scandinavia | Scandinavia | Europe |
| M-06-03 | Scandinavia | Scandinavia | Europe |
| M-09-05 | Scandinavia | Scandinavia | Europe |
| M-10-04 | Scandinavia | Scandinavia | Europe |
| M-98-08 | Scandinavia | Scandinavia | Europe |
| Maremma01 | Maremma Sheepdog | dog | not applicable |
| MARM000006 | Maremma Sheepdog | dog | not applicable |

| | | | |
|---|---|---|---|
| MastinoAbruzzese01 | Mastino Abruzzese | dog | not applicable |
| Mexican_wolf | US | US | North America |
| Minnesota_RKW119_RWJR007 | US | US | North America |
| Minnesota_RKW2515_RWJR016 | US | US | North America |
| Minnesota_RKW2518_RWJR012 | US | US | North America |
| Minnesota_RKW2523_RWJR009 | US | US | North America |
| Minnesota_RKW2524_RWJR003 | US | US | North America |
| Minnesota_wolf | US | US | North America |
| Mongolian_wolf | Mongolia | Mongolia | Asia |
| MW122 | Spain | Spain | Europe |
| MW127 | Spain | Spain | Europe |
| mxa | US | US | North America |
| NEAP000004 | Neapolitan Mastiff | dog | not applicable |
| NEAP000005 | Neapolitan Mastiff | dog | not applicable |
| NELK000002 | Norwegian Elkhound | dog | not applicable |
| NEWF000002 | Newfoundland | dog | not applicable |
| NEWF000006 | Newfoundland | dog | not applicable |
| NorwegianElkhound02 | Norwegian Elkhound | dog | not applicable |
| Penelope | Spain | Spain | Europe |
| PITB000001 | Pit Bull Terrier | dog | not applicable |
| PITB000002 | Pit Bull Terrier | dog | not applicable |
| POPT000001 | Portuguese Sheepdog | dog | not applicable |
| PortugueseWaterDog09 | Portuguese Water Dog | dog | not applicable |
| POSD000005 | Portuguese Sheepdog | dog | not applicable |
| PPOD000004 | Portuguese Podengo | dog | not applicable |
| PPOP000001 | Portuguese Podengo | dog | not applicable |
| PT61 | Village dog Portugal | dog | not applicable |
| ptw | Portugal | Portugal | Europe |
| PTWD000008 | Portuguese Water Dog | dog | not applicable |
| PYMF000005 | Pyrenean Mastiff | dog | not applicable |
| PYMF000006 | Pyrenean Mastiff | dog | not applicable |
| Qinghai_wolf | China | China | Asia |
| Quebec_MontTremblantNP_voyou0833M_RWBH001 | Canada | Canada | North America |
| RHOD000001 | Rhodesian Ridgeback | dog | not applicable |
| RKW13451 | China | China | Asia |

| | | | |
|---|---|---|---|
| RKW481_SF5 | African wild dog | outgroup - African wild dog | not applicable |
| SAAR000002 | Saarloos Wolfdog | dog | not applicable |
| SAAR000005 | Saarloos Wolfdog | dog | not applicable |
| SALU000001 | Saluki | dog | not applicable |
| SALU000002 | Saluki | dog | not applicable |
| SAMO000002 | Samoyed | dog | not applicable |
| Samoyed01 | Samoyed | dog | not applicable |
| SGGO000002 | Segugio Italiano | dog | not applicable |
| SGGO000004 | Segugio Italiano | dog | not applicable |
| SLOU000001 | Sloughi | dog | not applicable |
| SLOU000003 | Sloughi | dog | not applicable |
| SPIN000001 | Spinone Italiano | dog | not applicable |
| SPIN000004 | Spinone Italiano | dog | not applicable |
| SPMT000002 | Spanish Mastiff | dog | not applicable |
| SPMT000005 | Spanish Mastiff | dog | not applicable |
| spw | Spain | Spain | Europe |
| SPWD000002 | Spanish Water Dog | dog | not applicable |
| SPWD000003 | Spanish Water Dog | dog | not applicable |
| SRAB000001 | Sarabi | dog | not applicable |
| SRAB000002 | Sarabi | dog | not applicable |
| TERV000005 | Belgian Tervuren | dog | not applicable |
| TOSA000002 | Tosa Inu | dog | not applicable |
| TOSA000003 | Tosa Inu | dog | not applicable |
| TURM000002 | Turkish Mastiff | dog | not applicable |
| TURM000004 | Turkish Mastiff | dog | not applicable |
| TURV000001 | Belgian Tervuren | dog | not applicable |
| TWCH000001 | Treeing Walker Coonhound | dog | not applicable |
| TWCH000002 | Treeing Walker Coonhound | dog | not applicable |
| V113 | European Russia | European Russia | Europe |
| V114 | European Russia | European Russia | Europe |
| V115 | European Russia | European Russia | Europe |
| V116 | European Russia | European Russia | Europe |
| V117 | European Russia | European Russia | Europe |
| V119 | European Russia | European Russia | Europe |
| V120 | European Russia | European Russia | Europe |

| | | | |
|---|---|---|---|
| V126 | European Russia | European Russia | Europe |
| V132 | European Russia | European Russia | Europe |
| V134 | European Russia | European Russia | Europe |
| V136 | European Russia | European Russia | Europe |
| V141 | European Russia | European Russia | Europe |
| V143 | European Russia | European Russia | Europe |
| V3064 | hybrid Scandinavia | Scandinavia | Europe |
| V3065 | hybrid Scandinavia | Scandinavia | Europe |
| V3069 | hybrid Scandinavia | Scandinavia | Europe |
| VILLAF000001 | Village dog Afghanistan | dog | not applicable |
| VILLAF000002 | Village dog Afghanistan | dog | not applicable |
| VILLAF000003 | Village dog Afghanistan | dog | not applicable |
| VILLAZ000002 | Village dog Azerbaijan | dog | not applicable |
| VILLAZ000003 | Village dog Azerbaijan | dog | not applicable |
| VILLAZ000004 | Village dog Azerbaijan | dog | not applicable |
| VILLAZ000005 | Village dog Azerbaijan | dog | not applicable |
| VILLAZ000007 | Village dog Azerbaijan | dog | not applicable |
| VILLBG000001 | Village dog Bulgaria | dog | not applicable |
| VILLBG000002 | Village dog Bulgaria | dog | not applicable |
| VILLCG000001 | Village dog Congo | dog | not applicable |
| VILLCG000004 | Village dog Congo | dog | not applicable |
| VILLCG000006 | Village dog Congo | dog | not applicable |
| VILLCG000011 | Village dog Congo | dog | not applicable |
| VILLCG000013 | Village dog Congo | dog | not applicable |
| VILLCN000005 | Village dog China | dog | not applicable |
| VILLCN000013 | Village dog China | dog | not applicable |
| VILLCN000088 | Village dog China | dog | not applicable |
| VILLCN000145 | Village dog China | dog | not applicable |
| VILLCN000153 | Village dog China | dog | not applicable |
| VILLIR000021 | Village dog Iran | dog | not applicable |
| VILLIR000023 | Village dog Iran | dog | not applicable |
| VILLIR000024 | Village dog Iran | dog | not applicable |
| VILLIR000026 | Village dog Iran | dog | not applicable |
| VILLIR000027 | Village dog Iran | dog | not applicable |
| VILLKE000002 | Village dog Kenya | dog | not applicable |

| | | | |
|---|---|---|---|
| VILLKE000009 | Village dog Kenya | dog | not applicable |
| VILLKE000010 | Village dog Kenya | dog | not applicable |
| VILLKE000011 | Village dog Kenya | dog | not applicable |
| VILLKE000019 | Village dog Kenya | dog | not applicable |
| VILLLR000005 | Village dog Liberia | dog | not applicable |
| VILLLR000011 | Village dog Liberia | dog | not applicable |
| VILLLR000014 | Village dog Liberia | dog | not applicable |
| VILLLR000016 | Village dog Liberia | dog | not applicable |
| VILLLR000017 | Village dog Liberia | dog | not applicable |
| VILLTJ000001 | Village dog Tajikistan | dog | not applicable |
| VILLUZ000002 | Village dog Uzbekistan | dog | not applicable |
| VILLUZ000003 | Village dog Uzbekistan | dog | not applicable |
| VILLUZ000004 | Village dog Uzbekistan | dog | not applicable |
| VILLUZ000005 | Village dog Uzbekistan | dog | not applicable |
| VILLUZ000006 | Village dog Uzbekistan | dog | not applicable |
| VIZS000007 | Vizsla | dog | not applicable |
| W1023 | Italy | Italy | Europe |
| W11 | Finland | Finland | Europe |
| W14 | Finland | Finland | Europe |
| W1456 | Italy | Italy | Europe |
| W16 | Finland | Finland | Europe |
| W21 | Finland | Finland | Europe |
| W26 | Finland | Finland | Europe |
| W29 | Finland | Finland | Europe |
| W32 | Finland | Finland | Europe |
| W4 | Finland | Finland | Europe |
| W44 | Finland | Finland | Europe |
| W45 | Finland | Finland | Europe |
| W893 | Italy | Italy | Europe |
| WEIM000002 | Weimaraner | dog | not applicable |
| WEIM000005 | Weimaraner | dog | not applicable |
| WHPG000001 | Wirehaired Pointing Griffon | dog | not applicable |
| WHPG000002 | Wirehaired Pointing Griffon | dog | not applicable |
| WIT11 | Italy | Italy | Europe |
| WIT13 | Italy | Italy | Europe |

| | | | |
|---|---|---|---|
| WIT14 | Italy | Italy | Europe |
| WIT16 | Italy | Italy | Europe |
| WIT17 | Italy | Italy | Europe |
| WIT18 | Italy | Italy | Europe |
| WIT19 | Italy | Italy | Europe |
| WIT20 | Italy | Italy | Europe |
| WIT21 | Italy | Italy | Europe |
| WIT22 | Italy | Italy | Europe |
| WIT23 | Italy | Italy | Europe |
| wSierraMorena | Spain | Spain | Europe |
| Yellowstone1_wolf | US | US | North America |
| Yellowstone2_wolf | US | US | North America |
| Yellowstone3_wolf | US | US | North America |
| ysa | US | US | North America |
| ysb | US | US | North America |
| ZERD000002 | Turkish Zerdava | dog | not applicable |
| ZERD000003 | Turkish Zerdava | dog | not applicable |
| Kenya_GoldenJackal | Golden Jackal | outgroup - golden jackal | not applicable |
| cac | Coyote | outgroup - coyote | not applicable |

**Table S2.** Information about previously published genomes

| VCF_ID | Population | MeanDepth | Source | Passed quality filters |
|---|---|---|---|---|
| 140447_S11 | not applicable | 30,83 | Plassais et al. (2019) | yes |
| 149323_S6 | not applicable | 28,84 | Plassais et al. (2019) | yes |
| 165414_S20 | not applicable | 28,01 | Plassais et al. (2019) | yes |
| 171515_S19 | not applicable | 28,2 | Plassais et al. (2019) | yes |
| 173006_S10 | not applicable | 31,3 | Plassais et al. (2019) | yes |
| 173486_S3 | not applicable | 27,61 | Plassais et al. (2019) | yes |
| AKBH000001 | not applicable | 19,13 | Dog10K Consortium (Ostrander | yes |
| AKBH000003 | not applicable | 19,01 | Dog10K Consortium (Ostrander | yes |
| AMAL000001 | not applicable | 17,34 | Dog10K Consortium (Ostrander | yes |
| AMAL000002 | not applicable | 17,12 | Dog10K Consortium (Ostrander | yes |
| AMST000001 | not applicable | 17,15 | Dog10K Consortium (Ostrander | yes |
| AMST000006 | not applicable | 19,2 | Dog10K Consortium (Ostrander | yes |
| ANAT000003 | not applicable | 18 | Freedman et al. (2014) | yes |
| ANAT000004 | not applicable | 19,83 | Dog10K Consortium (Ostrander | yes |
| Arctic_BaffinIsl_CD130_RKW7639 | North America | 48,22 | Robinson et al. (2019) | yes |
| Arctic_EllesmereIsl_GF44_RKW7640 | North America | 47,38 | Robinson et al. (2019) | yes |
| Arctic_Nunavut_CB177_RKW7649 | North America | 34,85 | Robinson et al. (2019) | yes |
| Arctic_VictoriaIsl_CB215_RKW7619 | North America | 30,36 | Robinson et al. (2019) | yes |
| AUSS000001 | not applicable | 18,57 | Dog10K Consortium (Ostrander | yes |
| AUSS000003 | not applicable | 20,06 | Dog10K Consortium (Ostrander | yes |
| BEAU000004 | not applicable | 17,39 | Dog10K Consortium (Ostrander | yes |
| BEAU000007 | not applicable | 17,84 | Dog10K Consortium (Ostrander | yes |
| BELS000001 | not applicable | 18,29 | Dog10K Consortium (Ostrander | yes |
| BELS000003 | not applicable | 20,55 | Dog10K Consortium (Ostrander | yes |
| BGI-01505070001 | Tibetan | 25,41 | Plassais et al. (2019) | yes |
| BGI-01505070002 | Tibetan | 26,1 | Plassais et al. (2019) | yes |
| BGI-01505070003 | Tibetan | 26,25 | Plassais et al. (2019) | yes |
| BGI-01505070004 | Tibetan | 23,06 | Plassais et al. (2019) | yes |
| BGI-01505070005 | Tibetan | 25,52 | Plassais et al. (2019) | yes |
| BGI-01505070006 | East Asia | 26,21 | Plassais et al. (2019) | yes |
| BGI-01505070007 | East Asia | 23,68 | Plassais et al. (2019) | yes |
| BGI-01505070008 | Tibetan | 26,73 | Plassais et al. (2019) | yes |
| BLDH000002 | not applicable | 19,26 | Dog10K Consortium (Ostrander | yes |

| | | | | |
|---|---|---|---|---|
| BLDH000003 | not applicable | 16,98 | Dog10K Consortium (Ostrander | yes |
| BMAL000001 | not applicable | 19,63 | Dog10K Consortium (Ostrander | yes |
| BMAL000002 | not applicable | 17,28 | Dog10K Consortium (Ostrander | yes |
| BMSH000005 | not applicable | 15,12 | Dog10K Consortium (Ostrander | yes |
| BMSH000006 | not applicable | 17,98 | Dog10K Consortium (Ostrander | yes |
| BOER000002 | not applicable | 19,25 | Dog10K Consortium (Ostrander | yes |
| BOER000003 | not applicable | 18,73 | Dog10K Consortium (Ostrander | yes |
| BOXR000001 | not applicable | 21,04 | Dog10K Consortium (Ostrander | yes |
| BOXR000005 | not applicable | 20,35 | Dog10K Consortium (Ostrander | yes |
| BRAC000001 | not applicable | 21,13 | Dog10K Consortium (Ostrander | yes |
| BRAC000004 | not applicable | 19,46 | Dog10K Consortium (Ostrander | yes |
| BRIA000005 | not applicable | 19,72 | Dog10K Consortium (Ostrander | yes |
| BRIA000007 | not applicable | 18,53 | Dog10K Consortium (Ostrander | yes |
| BRMD000001 | not applicable | 19,52 | Dog10K Consortium (Ostrander | yes |
| BRMD000004 | not applicable | 0,82 | Dog10K Consortium (Ostrander | yes |
| BRTR000002 | not applicable | 18,99 | Dog10K Consortium (Ostrander | yes |
| BRTR000004 | not applicable | 17,48 | Dog10K Consortium (Ostrander | yes |
| CAUC000001 | not applicable | 18,62 | Dog10K Consortium (Ostrander | yes |
| CCRT000003 | not applicable | 18,35 | Dog10K Consortium (Ostrander | yes |
| CCRT000004 | not applicable | 18,65 | Dog10K Consortium (Ostrander | yes |
| CHBR000001 | not applicable | 18,16 | Dog10K Consortium (Ostrander | yes |
| CHBR000002 | not applicable | 17,42 | Dog10K Consortium (Ostrander | yes |
| CLUPAZ000001 | West Asia | 21,93 | Dog10K Consortium (Ostrander | yes |
| CLUPCN000001 | East Asia | 18,08 | Dog10K Consortium (Ostrander | yes |
| CLUPCN000002 | Tibetan | 17,2 | Dog10K Consortium (Ostrander | yes |
| CLUPCN000003 | Tibetan | 17,42 | Dog10K Consortium (Ostrander | yes |
| CLUPCN000004 | East Asia | 19,28 | Dog10K Consortium (Ostrander | yes |
| CLUPCN000005 | East Asia | 19,58 | Dog10K Consortium (Ostrander | yes |
| CLUPCN000006 | East Asia | 19,51 | Dog10K Consortium (Ostrander | yes |
| CLUPCN000007 | East Asia | 19,53 | Dog10K Consortium (Ostrander | yes |
| CLUPCN000008 | East Asia | 18,4 | Dog10K Consortium (Ostrander | yes |
| CLUPCN000009 | East Asia | 19,37 | Dog10K Consortium (Ostrander | yes |
| CLUPCN000010 | East Asia | 20,04 | Dog10K Consortium (Ostrander | yes |
| CLUPIR000001 | West Asia | 20,9 | Dog10K Consortium (Ostrander | yes |
| CLUPIR000002 | West Asia | 22,06 | Dog10K Consortium (Ostrander | yes |

| | | | | |
|---|---|---|---|---|
| CLUPIR000003 | West Asia | 22,74 | Dog10K Consortium (Ostrander | yes |
| CLUPIR000004 | West Asia | 23,88 | Dog10K Consortium (Ostrander | yes |
| CLUPIR000005 | West Asia | 24,07 | Dog10K Consortium (Ostrander | yes |
| CLUPIR000006 | West Asia | 22,68 | Dog10K Consortium (Ostrander | yes |
| CLUPKG000001 | Central Asia | 19,06 | Dog10K Consortium (Ostrander | yes |
| CLUPKZ000002 | Central Asia | 19,56 | Dog10K Consortium (Ostrander | yes |
| CLUPRU000001 | Central Asia | 18,8 | Dog10K Consortium (Ostrander | yes |
| CLUPRU000002 | Central Asia | 17,78 | Dog10K Consortium (Ostrander | yes |
| CLUPRU000003 | Central Asia | 18,3 | Dog10K Consortium (Ostrander | yes |
| CLUPRU000004 | East Asia | 19,46 | Dog10K Consortium (Ostrander | yes |
| CLUPRU000005 | Karelian | 16,84 | Dog10K Consortium (Ostrander | yes |
| CLUPRU000006 | Karelian | 17,67 | Dog10K Consortium (Ostrander | yes |
| CLUPRU000007 | Karelian | 20,52 | Dog10K Consortium (Ostrander | yes |
| CLUPRU000008 | Karelian | 19,28 | Dog10K Consortium (Ostrander | yes |
| CLUPRU000018 | West Asia | 17,11 | Dog10K Consortium (Ostrander | yes |
| CLUPRU000019 | Karelian | 17,94 | Dog10K Consortium (Ostrander | yes |
| CLUPRU000020 | Karelian | 20,77 | Dog10K Consortium (Ostrander | yes |
| CLUPTJ000001 | Central Asia | 15,44 | Dog10K Consortium (Ostrander | yes |
| CLUPTJ000002 | Central Asia | 20,25 | Dog10K Consortium (Ostrander | yes |
| CLUPTJ000003 | Central Asia | 20,97 | Dog10K Consortium (Ostrander | yes |
| CLUPTJ000004 | Central Asia | 20,34 | Dog10K Consortium (Ostrander | yes |
| CLUPTJ000005 | Central Asia | 20,09 | Dog10K Consortium (Ostrander | yes |
| CLUPTJ000006 | Central Asia | 22,07 | Dog10K Consortium (Ostrander | yes |
| CLUPTJ000007 | Central Asia | 17,44 | Dog10K Consortium (Ostrander | yes |
| CNCS000003 | not applicable | 16,97 | Dog10K Consortium (Ostrander | yes |
| CNCS000004 | not applicable | 20,13 | Dog10K Consortium (Ostrander | yes |
| COLL000007 | not applicable | 20,45 | Dog10K Consortium (Ostrander | yes |
| COLL000013 | not applicable | 27,44 | Dog10K Consortium (Ostrander | yes |
| COOK000002 | not applicable | 19,07 | Dog10K Consortium (Ostrander | yes |
| COOK000007 | not applicable | 11,39 | Dog10K Consortium (Ostrander | yes |
| CZEC000003 | not applicable | 19,21 | Dog10K Consortium (Ostrander | yes |
| CZWO000001 | not applicable | 19,21 | Dog10K Consortium (Ostrander | yes |
| D-00-12 | Scandinavian | 29,08 | Kardos et al. (2018) | yes |
| D-06-14 | Scandinavian | 41,28 | Kardos et al. (2018) | yes |
| D-07-16 | Scandinavian | 42,12 | Kardos et al. (2018) | yes |

| | | | | |
|---|---|---|---|---|
| D-08-10 | Scandinavian | 14,91 | Kardos et al. (2018) | yes |
| D-10-68 | Scandinavian | 13,9 | Kardos et al. (2018) | yes |
| D-11-58 | Scandinavian | 11,69 | Kardos et al. (2018) | yes |
| D-85-02 | Scandinavian | 48,96 | Kardos et al. (2018) | yes |
| DALM000002 | not applicable | 16,65 | Dog10K Consortium (Ostrander | yes |
| Dalmatian01 | not applicable | 23,52 | Plassais et al. (2019) | yes |
| DEER000003 | not applicable | 20,46 | Dog10K Consortium (Ostrander | yes |
| DEER000004 | not applicable | 21 | Dog10K Consortium (Ostrander | yes |
| Dhole_BerlinZoo | not applicable | 13,04 | Gopalakrishnan et al. (2018) | yes |
| EFXH000002 | not applicable | 17,49 | Dog10K Consortium (Ostrander | yes |
| EFXH000003 | not applicable | 15,5 | Dog10K Consortium (Ostrander | yes |
| ENTB000001 | not applicable | 19,16 | Dog10K Consortium (Ostrander | yes |
| ENTB000002 | not applicable | 17,5 | Dog10K Consortium (Ostrander | yes |
| ESMD000002 | not applicable | 18,71 | Dog10K Consortium (Ostrander | yes |
| ESMD000003 | not applicable | 21,72 | Dog10K Consortium (Ostrander | yes |
| FlatcoatedRetriever03 | not applicable | 37,63 | Plassais et al. (2019) | yes |
| FLCR000002 | not applicable | 20,01 | Dog10K Consortium (Ostrander | yes |
| G100-12 | Scandinavian | 13,72 | Kardos et al. (2018) | yes |
| G100-14 | Scandinavian | 42,86 | Kardos et al. (2018) | yes |
| G109-11 | Scandinavian | 12,85 | Kardos et al. (2018) | yes |
| G126-13 | Scandinavian | 11,92 | Kardos et al. (2018) | yes |
| G139-12 | Scandinavian | 11,43 | Kardos et al. (2018) | yes |
| G31-13 | Scandinavian | 12,91 | Kardos et al. (2018) | yes |
| G37-10 | Scandinavian | 13,23 | Kardos et al. (2018) | yes |
| G50-12 | Scandinavian | 12,13 | Kardos et al. (2018) | yes |
| GALG000002 | not applicable | 19,14 | Dog10K Consortium (Ostrander | yes |
| GALG000003 | not applicable | 18 | Dog10K Consortium (Ostrander | yes |
| glw | North America | 30,84 | Fan et al. (2016) | yes |
| GORD000002 | not applicable | 18,37 | Dog10K Consortium (Ostrander | yes |
| GPYR000003 | not applicable | 18,84 | Dog10K Consortium (Ostrander | yes |
| GPYR000004 | not applicable | 19,61 | Dog10K Consortium (Ostrander | yes |
| GreaterSwissMountainDog01 | not applicable | 38,14 | Plassais et al. (2019) | yes |
| GreyWolf_AtlanticCoast | North America | 8,74 | Sinding et al. (2018) | yes |
| GreyWolf_BaffinNorth | North America | 15,00 | Sinding et al. (2018) | yes |
| GreyWolf_BaffinSouth | North America | 13,70 | Sinding et al. (2018) | yes |

| | | | | |
|---|---|---|---|---|
| GreyWolf_BanksIsland | North America | 16,58 | Sinding et al. (2018) | yes |
| GreyWolf_EllesmereIsland | North America | 8,40 | Gopalakrishnan et al. (2018) | no |
| GreyWolf_Greenland | North America | 11,94 | Gopalakrishnan et al. (2018) | yes |
| GreyWolf_Montana | North America | 28,74 | Perri et al. (2021) | yes |
| GreyWolf_PacificCoast | North America | 13,96 | Sinding et al. (2018) | yes |
| GreyWolf_StLawrenceIsland | North America | 12,28 | Sinding et al. (2018) | yes |
| GreyWolf_Toronto | North America | 8,99 | Sinding et al. (2018) | yes |
| GreyWolf_VictoriaIsland | North America | 12,79 | Sinding et al. (2018) | yes |
| GRSD000002 | not applicable | 18,49 | Dog10K Consortium (Ostrander | yes |
| GRSD000003 | not applicable | 17,07 | Dog10K Consortium (Ostrander | yes |
| GSMD000004 | not applicable | 19,9 | Dog10K Consortium (Ostrander | yes |
| HOVA000003 | not applicable | 16,62 | Dog10K Consortium (Ostrander | yes |
| HOVA000004 | not applicable | 17,68 | Dog10K Consortium (Ostrander | yes |
| HWHP000001 | not applicable | 17,22 | Dog10K Consortium (Ostrander | yes |
| IBIZ000010 | not applicable | 20,45 | Dog10K Consortium (Ostrander | yes |
| IBIZ000011 | not applicable | 17,82 | Dog10K Consortium (Ostrander | yes |
| inw | Indian | 25,88 | Fan et al. (2016) | yes |
| Iran_wolf | West Asia | 24,05 | vonHoldt et al. (2016) | yes |
| IRSE000001 | not applicable | 22 | Dog10K Consortium (Ostrander | yes |
| IRSE000003 | not applicable | 20,57 | Dog10K Consortium (Ostrander | yes |
| irw | West Asia | 27,56 | Fan et al. (2016) | yes |
| IsleRoyaleNP_CL141_JRRW018 | North America | 24,18 | Robinson et al. (2019) | yes |
| IsleRoyaleNP_CL189_RWJR008 | North America | 24,34 | Robinson et al. (2019) | yes |
| IsleRoyaleNP_CL61_RWJR005 | North America | 23,37 | Robinson et al. (2019) | yes |
| IWSP000001 | not applicable | 19,75 | Dog10K Consortium (Ostrander | yes |
| IWSP000002 | not applicable | 19,44 | Dog10K Consortium (Ostrander | yes |
| KANG000001 | not applicable | 20,41 | Dog10K Consortium (Ostrander | yes |
| KANG000003 | not applicable | 20,1 | Dog10K Consortium (Ostrander | yes |
| KARS000005 | not applicable | 23,66 | Dog10K Consortium (Ostrander | yes |
| KARS000006 | not applicable | 22,67 | Dog10K Consortium (Ostrander | yes |
| KUVZ000002 | not applicable | 16,13 | Dog10K Consortium (Ostrander | yes |
| KUVZ000007 | not applicable | 19,48 | Dog10K Consortium (Ostrander | yes |
| L253 | NW Iberia | 13,6 | | yes |
| L409 | NW Iberia | 15,2 | | yes |
| L514 | NW Iberia | 13,99 | | yes |

| | | | | |
|---|---|---|---|---|
| L547 | NW Iberia | 15,27 | | yes |
| L552 | NW Iberia | 14,05 | | yes |
| L588 | NW Iberia | 16,18 | | yes |
| L844 | NW Iberia | 13,66 | | yes |
| LAGO000001 | not applicable | 16,69 | Dog10K Consortium (Ostrander | yes |
| LAGO000004 | not applicable | 18,36 | Dog10K Consortium (Ostrander | yes |
| Lcu2_Pastora | not applicable | 10,64 | Plassais et al. (2019) | yes |
| LEOP000003 | not applicable | 16,95 | Dog10K Consortium (Ostrander | yes |
| LEOP000008 | not applicable | 19,04 | Dog10K Consortium (Ostrander | yes |
| LUP006817 | Dinaric-Balkan | 17 | Ostrander et al. (2019) | yes |
| LUP006818 | Dinaric-Balkan | 17,16 | Ostrander et al. (2019) | yes |
| LUP006819 | Dinaric-Balkan | 15,96 | Ostrander et al. (2019) | yes |
| LUP006820 | Dinaric-Balkan | 16,09 | Ostrander et al. (2019) | yes |
| LUP006821 | Dinaric-Balkan | 16,47 | Ostrander et al. (2019) | yes |
| LUP006822 | Dinaric-Balkan | 19,02 | Ostrander et al. (2019) | yes |
| LUP006823 | Dinaric-Balkan | 16,11 | Ostrander et al. (2019) | yes |
| LUP006824 | Dinaric-Balkan | 15,65 | Ostrander et al. (2019) | yes |
| LUP006825 | Dinaric-Balkan | 18,47 | Ostrander et al. (2019) | yes |
| LUP006826 | Dinaric-Balkan | 13,9 | Ostrander et al. (2019) | yes |
| LUP006827 | Dinaric-Balkan | 19,28 | Ostrander et al. (2019) | yes |
| LUP006828 | Dinaric-Balkan | 15,74 | Ostrander et al. (2019) | yes |
| LUP006829 | NW Iberia | 16,71 | Ostrander et al. (2019) | yes |
| LUP006830 | NW Iberia | 18,22 | Ostrander et al. (2019) | yes |
| LUPWCHN00003 | East Asia | 13,86 | Plassais et al. (2019) | yes |
| LUPWCHN00010 | East Asia | 15,27 | Plassais et al. (2019) | yes |
| LUPZCHN00006 | East Asia | 12,22 | Plassais et al. (2019) | yes |
| LUPZCHN00009 | East Asia | 27,13 | Plassais et al. (2019) | yes |
| LUPZCHN00013 | East Asia | 12,31 | Plassais et al. (2019) | no |
| M-01-06 | Scandinavian | 25,42 | Kardos et al. (2018) | yes |
| M-06-03 | Scandinavian | 37,82 | Kardos et al. (2018) | yes |
| M-09-05 | Scandinavian | 13,26 | Kardos et al. (2018) | yes |
| M-10-04 | Scandinavian | 16,83 | Kardos et al. (2018) | yes |
| M-98-08 | Scandinavian | 40,69 | Kardos et al. (2018) | yes |
| Maremma01 | not applicable | 26,83 | Plassais et al. (2019) | yes |
| MARM000006 | not applicable | 20,05 | Dog10K Consortium (Ostrander | yes |

| | | | | |
|---|---|---|---|---|
| MastinoAbruzzese01 | not applicable | 26,31 | Plassais et al. (2019) | yes |
| Mexican_wolf | North America | 21,70 | vonHoldt et al. (2016) | yes |
| Minnesota_RKW119_RWJR007 | North America | 22,02 | Robinson et al. (2019) | yes |
| Minnesota_RKW2515_RWJR016 | North America | 24,45 | Robinson et al. (2019) | yes |
| Minnesota_RKW2518_RWJR012 | North America | 23,77 | Robinson et al. (2019) | yes |
| Minnesota_RKW2523_RWJR009 | North America | 24,31 | Robinson et al. (2019) | yes |
| Minnesota_RKW2524_RWJR003 | North America | 21,63 | Robinson et al. (2019) | yes |
| Minnesota_wolf | North America | 22,38 | vonHoldt et al. (2016) | yes |
| Mongolian_wolf | Central Asia | 19,71 | vonHoldt et al. (2016) | yes |
| MW122 | NW Iberia | 15,84 | Ciucani et al. (2023) | yes |
| MW127 | NW Iberia | 14,71 | Ciucani et al. (2023) | yes |
| mxa | North America | 24,44 | Fan et al. (2016) | yes |
| NEAP000004 | not applicable | 16,66 | Dog10K Consortium (Ostrander | yes |
| NEAP000005 | not applicable | 19,64 | Dog10K Consortium (Ostrander | yes |
| NELK000002 | not applicable | 18,02 | Dog10K Consortium (Ostrander | yes |
| NEWF000002 | not applicable | 17,48 | Dog10K Consortium (Ostrander | yes |
| NEWF000006 | not applicable | 17,67 | Dog10K Consortium (Ostrander | yes |
| NorwegianElkhound02 | not applicable | 37,04 | Plassais et al. (2019) | yes |
| Penelope | NW Iberia | 26,4 | Frantz et al. (2016) | yes |
| PITB000001 | not applicable | 18,57 | Dog10K Consortium (Ostrander | yes |
| PITB000002 | not applicable | 18,21 | Dog10K Consortium (Ostrander | yes |
| POPT000001 | not applicable | 18,1 | Dog10K Consortium (Ostrander | yes |
| PortugueseWaterDog09 | not applicable | 31 | Plassais et al. (2019) | yes |
| POSD000005 | not applicable | 16,65 | Dog10K Consortium (Ostrander | yes |
| PPOD000004 | not applicable | 18,16 | Dog10K Consortium (Ostrander | yes |
| PPOP000001 | not applicable | 16,56 | Dog10K Consortium (Ostrander | yes |
| PT61 | not applicable | 13,73 | Plassais et al. (2019) | yes |
| ptw | NW Iberia | 30,75 | Fan et al. (2016) | yes |
| PTWD000008 | not applicable | 18,91 | Dog10K Consortium (Ostrander | yes |
| PYMF000005 | not applicable | 18,19 | Dog10K Consortium (Ostrander | yes |
| PYMF000006 | not applicable | 17,38 | Dog10K Consortium (Ostrander | yes |
| Qinghai_wolf | NW Iberia | 22,77 | vonHoldt et al. (2016) | yes |
| Quebec_MontTremblantNP_voyou0833M_RWBH001 | North America | 24,48 | Robinson et al. (2019) | yes |
| RHOD000001 | not applicable | 18,01 | Dog10K Consortium (Ostrander | yes |
| RKW13451 | East Asia | 27,57 | Freedman et al. (2014) | yes |

| | | | | |
|---|---|---|---|---|
| RKW481_SF5 | not applicable | 26,4 | Campana et al. (2016) | yes |
| SAAR000002 | not applicable | 16,23 | Dog10K Consortium (Ostrander | yes |
| SAAR000005 | not applicable | 17,92 | Dog10K Consortium (Ostrander | yes |
| SALU000001 | not applicable | 19,33 | Dog10K Consortium (Ostrander | yes |
| SALU000002 | not applicable | 18,85 | Dog10K Consortium (Ostrander | no |
| SAMO000002 | not applicable | 17,57 | Dog10K Consortium (Ostrander | yes |
| Samoyed01 | not applicable | 44,78 | Plassais et al. (2019) | yes |
| SGGO000002 | not applicable | 18,36 | Dog10K Consortium (Ostrander | yes |
| SGGO000004 | not applicable | 19,45 | Dog10K Consortium (Ostrander | yes |
| SLOU000001 | not applicable | 17,18 | Dog10K Consortium (Ostrander | yes |
| SLOU000003 | not applicable | 17,6 | Dog10K Consortium (Ostrander | yes |
| SPIN000001 | not applicable | 18,46 | Dog10K Consortium (Ostrander | yes |
| SPIN000004 | not applicable | 21,29 | Dog10K Consortium (Ostrander | yes |
| SPMT000002 | not applicable | 16,42 | Dog10K Consortium (Ostrander | yes |
| SPMT000005 | not applicable | 19,74 | Dog10K Consortium (Ostrander | yes |
| spw | NW Iberia | 24,92 | Fan et al. (2016) | yes |
| SPWD000002 | not applicable | 19,16 | Dog10K Consortium (Ostrander | yes |
| SPWD000003 | not applicable | 20,63 | Dog10K Consortium (Ostrander | yes |
| SRAB000001 | not applicable | 17,92 | Dog10K Consortium (Ostrander | yes |
| SRAB000002 | not applicable | 17,97 | Dog10K Consortium (Ostrander | yes |
| TERV000005 | not applicable | 19,88 | Dog10K Consortium (Ostrander | yes |
| TOSA000002 | not applicable | 16,74 | Dog10K Consortium (Ostrander | yes |
| TOSA000003 | not applicable | 19,31 | Dog10K Consortium (Ostrander | yes |
| TURM000002 | not applicable | 18,8 | Dog10K Consortium (Ostrander | yes |
| TURM000004 | not applicable | 19,53 | Dog10K Consortium (Ostrander | yes |
| TURV000001 | not applicable | 16,27 | Dog10K Consortium (Ostrander | yes |
| TWCH000001 | not applicable | 15,27 | Dog10K Consortium (Ostrander | yes |
| TWCH000002 | not applicable | 16,47 | Dog10K Consortium (Ostrander | yes |
| V113 | Karelian | 39,69 | Smeds et al. (2021) | yes |
| V114 | Karelian | 39,9 | Smeds et al. (2021) | yes |
| V115 | Karelian | 35,29 | Smeds et al. (2021) | yes |
| V116 | Karelian | 40,28 | Smeds et al. (2021) | yes |
| V117 | Karelian | 40,78 | Smeds et al. (2021) | yes |
| V119 | Karelian | 40,08 | Smeds et al. (2021) | yes |
| V120 | Karelian | 40,73 | Smeds et al. (2021) | yes |

| | | | | |
|---|---|---|---|---|
| V126 | Karelian | 42,6 | Smeds et al. (2021) | yes |
| V132 | Karelian | 43,79 | Smeds et al. (2021) | yes |
| V134 | Karelian | 39,43 | Smeds et al. (2021) | yes |
| V136 | Karelian | 42,76 | Smeds et al. (2021) | yes |
| V141 | Karelian | 40,27 | Smeds et al. (2021) | yes |
| V143 | Karelian | 42,15 | Smeds et al. (2021) | yes |
| V3064 | Scandinavian | 42,27 | Smeds et al. (2021) | yes |
| V3065 | Scandinavian | 42,77 | Smeds et al. (2021) | yes |
| V3069 | Scandinavian | 41,34 | Smeds et al. (2021) | yes |
| VILLAF000001 | not applicable | 18,86 | Dog10K Consortium (Ostrander | yes |
| VILLAF000002 | not applicable | 21,75 | Dog10K Consortium (Ostrander | yes |
| VILLAF000003 | not applicable | 19,77 | Dog10K Consortium (Ostrander | yes |
| VILLAZ000002 | not applicable | 18,91 | Dog10K Consortium (Ostrander | yes |
| VILLAZ000003 | not applicable | 18,66 | Dog10K Consortium (Ostrander | yes |
| VILLAZ000004 | not applicable | 17,09 | Dog10K Consortium (Ostrander | yes |
| VILLAZ000005 | not applicable | 23,18 | Dog10K Consortium (Ostrander | yes |
| VILLAZ000007 | not applicable | 19,11 | Dog10K Consortium (Ostrander | yes |
| VILLBG000001 | not applicable | 15,37 | Ostrander et al. (2019) | yes |
| VILLBG000002 | not applicable | 13,97 | Ostrander et al. (2019) | yes |
| VILLCG000001 | not applicable | 19,94 | Dog10K Consortium (Ostrander | yes |
| VILLCG000004 | not applicable | 22,56 | Dog10K Consortium (Ostrander | yes |
| VILLCG000006 | not applicable | 18,38 | Dog10K Consortium (Ostrander | yes |
| VILLCG000011 | not applicable | 20,3 | Dog10K Consortium (Ostrander | yes |
| VILLCG000013 | not applicable | 20,1 | Dog10K Consortium (Ostrander | yes |
| VILLCN000005 | not applicable | 20,76 | Dog10K Consortium (Ostrander | yes |
| VILLCN000013 | not applicable | 23,82 | Dog10K Consortium (Ostrander | yes |
| VILLCN000088 | not applicable | 21,64 | Dog10K Consortium (Ostrander | yes |
| VILLCN000145 | not applicable | 19,47 | Dog10K Consortium (Ostrander | yes |
| VILLCN000153 | not applicable | 21,9 | Dog10K Consortium (Ostrander | yes |
| VILLIR000021 | not applicable | 23,26 | Dog10K Consortium (Ostrander | yes |
| VILLIR000023 | not applicable | 22,24 | Dog10K Consortium (Ostrander | yes |
| VILLIR000024 | not applicable | 22,23 | Dog10K Consortium (Ostrander | yes |
| VILLIR000026 | not applicable | 23,72 | Dog10K Consortium (Ostrander | yes |
| VILLIR000027 | not applicable | 22,26 | Dog10K Consortium (Ostrander | yes |
| VILLKE000002 | not applicable | 21,89 | Dog10K Consortium (Ostrander | yes |

| | | | | |
|---|---|---|---|---|
| VILLKE000009 | not applicable | 21,02 | Dog10K Consortium (Ostrander | yes |
| VILLKE000010 | not applicable | 21,76 | Dog10K Consortium (Ostrander | yes |
| VILLKE000011 | not applicable | 20,67 | Dog10K Consortium (Ostrander | yes |
| VILLKE000019 | not applicable | 21,09 | Dog10K Consortium (Ostrander | yes |
| VILLLR000005 | not applicable | 20,68 | Dog10K Consortium (Ostrander | yes |
| VILLLR000011 | not applicable | 23,76 | Dog10K Consortium (Ostrander | yes |
| VILLLR000014 | not applicable | 21,7 | Dog10K Consortium (Ostrander | yes |
| VILLLR000016 | not applicable | 19,23 | Dog10K Consortium (Ostrander | yes |
| VILLLR000017 | not applicable | 18,58 | Dog10K Consortium (Ostrander | yes |
| VILLTJ000001 | not applicable | 19,62 | Dog10K Consortium (Ostrander | yes |
| VILLUZ000002 | not applicable | 18,94 | Dog10K Consortium (Ostrander | yes |
| VILLUZ000003 | not applicable | 19,43 | Dog10K Consortium (Ostrander | yes |
| VILLUZ000004 | not applicable | 19,08 | Dog10K Consortium (Ostrander | yes |
| VILLUZ000005 | not applicable | 19,35 | Dog10K Consortium (Ostrander | yes |
| VILLUZ000006 | not applicable | 18,56 | Dog10K Consortium (Ostrander | yes |
| VIZS000007 | not applicable | 17,8 | Dog10K Consortium (Ostrander | yes |
| W1023 | Italian Peninsula | 25,19 | Battilani et al. (2025) | yes |
| W11 | Karelian | 21,02 | Smeds et al. (2019) | yes |
| W14 | Karelian | 31,73 | Smeds et al. (2019) | yes |
| W1456 | Italian Peninsula | 27,09 | Battilani et al. (2025) | yes |
| W16 | Karelian | 26 | Smeds et al. (2021) | yes |
| W21 | Karelian | 29,82 | Smeds et al. (2019) | yes |
| W26 | Karelian | 34,7 | Smeds et al. (2019) | yes |
| W29 | Karelian | 23,91 | Smeds et al. (2021) | yes |
| W32 | Karelian | 21,94 | Smeds et al. (2021) | yes |
| W4 | Karelian | 27,96 | Smeds et al. (2021) | yes |
| W44 | Karelian | 26,17 | Smeds et al. (2019) | yes |
| W45 | Karelian | 27,64 | Smeds et al. (2021) | yes |
| W893 | Italian Peninsula | 20,05 | Battilani et al. (2025) | yes |
| WEIM000002 | not applicable | 18,16 | Dog10K Consortium (Ostrander | yes |
| WEIM000005 | not applicable | 17,6 | Dog10K Consortium (Ostrander | yes |
| WHPG000001 | not applicable | 19,04 | Dog10K Consortium (Ostrander | yes |
| WHPG000002 | not applicable | 19,62 | Dog10K Consortium (Ostrander | yes |
| WIT11 | Italian Peninsula | 16,01 | Battilani et al. (2024) | yes |
| WIT13 | Italian Peninsula | 12,65 | Battilani et al. (2024) | yes |

| | | | | |
|---|---|---|---|---|
| WIT14 | Italian Peninsula | 16,92 | Battilani et al. (2024) | yes |
| WIT16 | Italian Peninsula | 18,88 | Battilani et al. (2024) | yes |
| WIT17 | Italian Peninsula | 14,16 | Battilani et al. (2024) | yes |
| WIT18 | Italian Peninsula | 19,97 | Battilani et al. (2024) | yes |
| WIT19 | Italian Peninsula | 14,24 | Battilani et al. (2024) | yes |
| WIT20 | Italian Peninsula | 18,86 | Battilani et al. (2024) | yes |
| WIT21 | Italian Peninsula | 14,02 | Battilani et al. (2024) | yes |
| WIT22 | Italian Peninsula | 12,24 | Battilani et al. (2024) | yes |
| WIT23 | Italian Peninsula | 16,76 | Battilani et al. (2024) | yes |
| wSierraMorena | NW Iberia | 14,53 | Gomez-Sanchez et al. (2018) | yes |
| Yellowstone1_wolf | North America | 6,07 | vonHoldt et al. (2016) | no |
| Yellowstone2_wolf | North America | 23,67 | vonHoldt et al. (2016) | yes |
| Yellowstone3_wolf | North America | 22,16 | vonHoldt et al. (2016) | yes |
| ysa | North America | 26,86 | Fan et al. (2016) | yes |
| ysb | North America | 25,08 | Fan et al. (2016) | yes |
| ZERD000002 | not applicable | 18,84 | Dog10K Consortium (Ostrander | yes |
| ZERD000003 | not applicable | 19,22 | Dog10K Consortium (Ostrander | yes |
| Kenya_GoldenJackal | not applicable | 22,46 | vonHoldt et al. (2016) | used only for genetic loa |
| cac | not applicable | 25,36 | Plassais et al. (2019) | used only for genetic loa |

**Table S2.** Information about previously published genomes

| VCF_ID | Passed dog introgression filters | Passed relatedness filters |
|---|---|---|
| 140447_S11 | no | NA |
| 149323_S6 | no | NA |
| 165414_S20 | no | NA |
| 171515_S19 | no | NA |
| 173006_S10 | no | NA |
| 173486_S3 | no | NA |
| AKBH000001 | no | NA |
| AKBH000003 | no | NA |
| AMAL000001 | no | NA |
| AMAL000002 | no | NA |
| AMST000001 | no | NA |
| AMST000006 | no | NA |
| ANAT000003 | no | NA |
| ANAT000004 | no | NA |
| Arctic_BaffinIsl_CD130_RKW7639 | yes | yes |
| Arctic_EllesmereIsl_GF44_RKW7640 | yes | yes |
| Arctic_Nunavut_CB177_RKW7649 | yes | yes |
| Arctic_VictoriaIsl_CB215_RKW7619 | yes | yes |
| AUSS000001 | no | NA |
| AUSS000003 | no | NA |
| BEAU000004 | no | NA |
| BEAU000007 | no | NA |
| BELS000001 | no | NA |
| BELS000003 | no | NA |
| BGI-01505070001 | yes | yes |
| BGI-01505070002 | yes | yes |
| BGI-01505070003 | yes | yes |
| BGI-01505070004 | yes | yes |
| BGI-01505070005 | yes | yes |
| BGI-01505070006 | yes | yes |
| BGI-01505070007 | yes | yes |
| BGI-01505070008 | yes | yes |
| BLDH000002 | no | NA |

| | | |
|---|---|---|
| BLDH000003 | no | NA |
| BMAL000001 | no | NA |
| BMAL000002 | no | NA |
| BMSH000005 | no | NA |
| BMSH000006 | no | NA |
| BOER000002 | no | NA |
| BOER000003 | no | NA |
| BOXR000001 | no | NA |
| BOXR000005 | no | NA |
| BRAC000001 | no | NA |
| BRAC000004 | no | NA |
| BRIA000005 | no | NA |
| BRIA000007 | no | NA |
| BRMD000001 | no | NA |
| BRMD000004 | no | NA |
| BRTR000002 | no | NA |
| BRTR000004 | no | NA |
| CAUC000001 | no | NA |
| CCRT000003 | no | NA |
| CCRT000004 | no | NA |
| CHBR000001 | no | NA |
| CHBR000002 | no | NA |
| CLUPAZ000001 | yes | yes |
| CLUPCN000001 | yes | yes |
| CLUPCN000002 | yes | no |
| CLUPCN000003 | yes | yes |
| CLUPCN000004 | yes | no |
| CLUPCN000005 | yes | yes |
| CLUPCN000006 | yes | yes |
| CLUPCN000007 | yes | yes |
| CLUPCN000008 | yes | yes |
| CLUPCN000009 | yes | yes |
| CLUPCN000010 | yes | yes |
| CLUPIR000001 | yes | yes |
| CLUPIR000002 | yes | no |

| | | |
|---|---|---|
| CLUPIR000003 | yes | yes |
| CLUPIR000004 | yes | yes |
| CLUPIR000005 | yes | yes |
| CLUPIR000006 | yes | no |
| CLUPKG000001 | yes | yes |
| CLUPKZ000002 | yes | yes |
| CLUPRU000001 | yes | yes |
| CLUPRU000002 | yes | yes |
| CLUPRU000003 | yes | yes |
| CLUPRU000004 | yes | yes |
| CLUPRU000005 | yes | yes |
| CLUPRU000006 | yes | no |
| CLUPRU000007 | yes | yes |
| CLUPRU000008 | yes | yes |
| CLUPRU000018 | no | NA |
| CLUPRU000019 | yes | yes |
| CLUPRU000020 | yes | yes |
| CLUPTJ000001 | yes | yes |
| CLUPTJ000002 | yes | yes |
| CLUPTJ000003 | yes | yes |
| CLUPTJ000004 | yes | yes |
| CLUPTJ000005 | yes | yes |
| CLUPTJ000006 | yes | yes |
| CLUPTJ000007 | yes | yes |
| CNCS000003 | no | NA |
| CNCS000004 | no | NA |
| COLL000007 | no | NA |
| COLL000013 | no | NA |
| COOK000002 | no | NA |
| COOK000007 | no | NA |
| CZEC000003 | no | NA |
| CZWO000001 | no | NA |
| D-00-12 | yes | yes |
| D-06-14 | yes | yes |
| D-07-16 | yes | yes |

| | | |
|---|---|---|
| D-08-10 | yes | yes |
| D-10-68 | yes | yes |
| D-11-58 | yes | yes |
| D-85-02 | yes | yes |
| DALM000002 | no | NA |
| Dalmatian01 | no | NA |
| DEER000003 | no | NA |
| DEER000004 | no | NA |
| Dhole_BerlinZoo | yes | yes |
| EFXH000002 | no | NA |
| EFXH000003 | no | NA |
| ENTB000001 | no | NA |
| ENTB000002 | no | NA |
| ESMD000002 | no | NA |
| ESMD000003 | no | NA |
| FlatcoatedRetriever03 | no | NA |
| FLCR000002 | no | NA |
| G100-12 | yes | yes |
| G100-14 | yes | yes |
| G109-11 | yes | no |
| G126-13 | yes | yes |
| G139-12 | yes | yes |
| G31-13 | yes | no |
| G37-10 | yes | no |
| G50-12 | yes | yes |
| GALG000002 | no | NA |
| GALG000003 | no | NA |
| glw | yes | yes |
| GORD000002 | no | NA |
| GPYR000003 | no | NA |
| GPYR000004 | no | NA |
| GreaterSwissMountainDog01 | no | NA |
| GreyWolf_AtlanticCoast | yes | yes |
| GreyWolf_BaffinNorth | yes | yes |
| GreyWolf_BaffinSouth | yes | yes |

| | | |
|---|---|---|
| GreyWolf_BanksIsland | yes | yes |
| GreyWolf_EllesmereIsland | NA | NA |
| GreyWolf_Greenland | yes | yes |
| GreyWolf_Montana | yes | yes |
| GreyWolf_PacificCoast | yes | yes |
| GreyWolf_StLawrenceIsland | yes | yes |
| GreyWolf_Toronto | yes | yes |
| GreyWolf_VictoriaIsland | yes | yes |
| GRSD000002 | no | NA |
| GRSD000003 | no | NA |
| GSMD000004 | no | NA |
| HOVA000003 | no | NA |
| HOVA000004 | no | NA |
| HWHP000001 | no | NA |
| IBIZ000010 | no | NA |
| IBIZ000011 | no | NA |
| inw | yes | yes |
| Iran_wolf | yes | no |
| IRSE000001 | no | NA |
| IRSE000003 | no | NA |
| irw | yes | yes |
| IsleRoyaleNP_CL141_JRRW018 | yes | yes |
| IsleRoyaleNP_CL189_RWJR008 | yes | yes |
| IsleRoyaleNP_CL61_RWJR005 | yes | yes |
| IWSP000001 | no | NA |
| IWSP000002 | no | NA |
| KANG000001 | no | NA |
| KANG000003 | no | NA |
| KARS000005 | no | NA |
| KARS000006 | no | NA |
| KUVZ000002 | no | NA |
| KUVZ000007 | no | NA |
| L253 | yes | yes |
| L409 | yes | yes |
| L514 | yes | yes |

| | | |
|---|---|---|
| L547 | yes | yes |
| L552 | yes | yes |
| L588 | yes | yes |
| L844 | yes | no |
| LAGO000001 | no | NA |
| LAGO000004 | no | NA |
| Lcu2_Pastora | yes | yes |
| LEOP000003 | no | NA |
| LEOP000008 | no | NA |
| LUP006817 | yes | yes |
| LUP006818 | yes | yes |
| LUP006819 | yes | yes |
| LUP006820 | yes | yes |
| LUP006821 | yes | yes |
| LUP006822 | yes | yes |
| LUP006823 | yes | yes |
| LUP006824 | yes | yes |
| LUP006825 | yes | yes |
| LUP006826 | yes | yes |
| LUP006827 | yes | yes |
| LUP006828 | yes | yes |
| LUP006829 | yes | yes |
| LUP006830 | yes | yes |
| LUPWCHN00003 | no | NA |
| LUPWCHN00010 | yes | yes |
| LUPZCHN00006 | no | NA |
| LUPZCHN00009 | yes | yes |
| LUPZCHN00013 | NA | NA |
| M-01-06 | yes | yes |
| M-06-03 | yes | yes |
| M-09-05 | yes | yes |
| M-10-04 | yes | yes |
| M-98-08 | yes | yes |
| Maremma01 | no | NA |
| MARM000006 | no | NA |

| | | |
|---|---|---|
| MastinoAbruzzese01 | no | NA |
| Mexican_wolf | yes | no |
| Minnesota_RKW119_RWJR007 | yes | yes |
| Minnesota_RKW2515_RWJR016 | yes | yes |
| Minnesota_RKW2518_RWJR012 | yes | yes |
| Minnesota_RKW2523_RWJR009 | yes | yes |
| Minnesota_RKW2524_RWJR003 | yes | yes |
| Minnesota_wolf | yes | no |
| Mongolian_wolf | yes | no |
| MW122 | yes | yes |
| MW127 | yes | yes |
| mxa | yes | yes |
| NEAP000004 | no | NA |
| NEAP000005 | no | NA |
| NELK000002 | no | NA |
| NEWF000002 | no | NA |
| NEWF000006 | no | NA |
| NorwegianElkhound02 | no | NA |
| Penelope | yes | yes |
| PITB000001 | no | NA |
| PITB000002 | no | NA |
| POPT000001 | no | NA |
| PortugueseWaterDog09 | no | NA |
| POSD000005 | no | NA |
| PPOD000004 | no | NA |
| PPOP000001 | no | NA |
| PT61 | no | NA |
| ptw | yes | yes |
| PTWD000008 | no | NA |
| PYMF000005 | no | NA |
| PYMF000006 | no | NA |
| Qinghai_wolf | yes | no |
| Quebec_MontTremblantNP_voyou0833M_RWBH001 | yes | yes |
| RHOD000001 | no | NA |
| RKW13451 | yes | yes |

| | | |
|---|---|---|
| RKW481_SF5 | yes | yes |
| SAAR000002 | no | NA |
| SAAR000005 | no | NA |
| SALU000001 | no | NA |
| SALU000002 | NA | NA |
| SAMO000002 | no | NA |
| Samoyed01 | no | NA |
| SGGO000002 | no | NA |
| SGGO000004 | no | NA |
| SLOU000001 | no | NA |
| SLOU000003 | no | NA |
| SPIN000001 | no | NA |
| SPIN000004 | no | NA |
| SPMT000002 | no | NA |
| SPMT000005 | no | NA |
| spw | no | NA |
| SPWD000002 | no | NA |
| SPWD000003 | no | NA |
| SRAB000001 | no | NA |
| SRAB000002 | no | NA |
| TERV000005 | no | NA |
| TOSA000002 | no | NA |
| TOSA000003 | no | NA |
| TURM000002 | no | NA |
| TURM000004 | no | NA |
| TURV000001 | no | NA |
| TWCH000001 | no | NA |
| TWCH000002 | no | NA |
| V113 | yes | yes |
| V114 | yes | yes |
| V115 | yes | yes |
| V116 | yes | yes |
| V117 | yes | yes |
| V119 | yes | yes |
| V120 | yes | yes |

| | | |
|---|---|---|
| V126 | yes | yes |
| V132 | yes | yes |
| V134 | yes | yes |
| V136 | yes | yes |
| V141 | yes | yes |
| V143 | yes | yes |
| V3064 | no | NA |
| V3065 | no | NA |
| V3069 | no | NA |
| VILLAF000001 | no | NA |
| VILLAF000002 | no | NA |
| VILLAF000003 | no | NA |
| VILLAZ000002 | no | NA |
| VILLAZ000003 | no | NA |
| VILLAZ000004 | no | NA |
| VILLAZ000005 | no | NA |
| VILLAZ000007 | no | NA |
| VILLBG000001 | no | NA |
| VILLBG000002 | no | NA |
| VILLCG000001 | no | NA |
| VILLCG000004 | no | NA |
| VILLCG000006 | no | NA |
| VILLCG000011 | no | NA |
| VILLCG000013 | no | NA |
| VILLCN000005 | no | NA |
| VILLCN000013 | no | NA |
| VILLCN000088 | no | NA |
| VILLCN000145 | no | NA |
| VILLCN000153 | no | NA |
| VILLIR000021 | no | NA |
| VILLIR000023 | no | NA |
| VILLIR000024 | no | NA |
| VILLIR000026 | no | NA |
| VILLIR000027 | no | NA |
| VILLKE000002 | no | NA |

| | | |
|---|---|---|
| VILLKE000009 | no | NA |
| VILLKE000010 | no | NA |
| VILLKE000011 | no | NA |
| VILLKE000019 | no | NA |
| VILLLR000005 | no | NA |
| VILLLR000011 | no | NA |
| VILLLR000014 | no | NA |
| VILLLR000016 | no | NA |
| VILLLR000017 | no | NA |
| VILLTJ000001 | no | NA |
| VILLUZ000002 | no | NA |
| VILLUZ000003 | no | NA |
| VILLUZ000004 | no | NA |
| VILLUZ000005 | no | NA |
| VILLUZ000006 | no | NA |
| VIZS000007 | no | NA |
| W1023 | yes | yes |
| W11 | yes | yes |
| W14 | yes | yes |
| W1456 | yes | yes |
| W16 | yes | yes |
| W21 | yes | yes |
| W26 | yes | yes |
| W29 | yes | no |
| W32 | yes | yes |
| W4 | yes | yes |
| W44 | yes | yes |
| W45 | yes | yes |
| W893 | yes | yes |
| WEIM000002 | no | NA |
| WEIM000005 | no | NA |
| WHPG000001 | no | NA |
| WHPG000002 | no | NA |
| WIT11 | yes | yes |
| WIT13 | yes | yes |

| | | |
|---|---|---|
| WIT14 | yes | yes |
| WIT16 | yes | yes |
| WIT17 | yes | yes |
| WIT18 | no | NA |
| WIT19 | yes | yes |
| WIT20 | yes | yes |
| WIT21 | yes | yes |
| WIT22 | yes | yes |
| WIT23 | yes | yes |
| wSierraMorena | no | yes |
| Yellowstone1_wolf | NA | NA |
| Yellowstone2_wolf | yes | no |
| Yellowstone3_wolf | yes | no |
| ysa | yes | yes |
| ysb | yes | yes |
| ZERD000002 | no | NA |
| ZERD000003 | no | NA |
| Kenya_GoldenJackal | d pseudohaploidization | |
| cac | d pseudohaploidization | |

**Table S3.** Ancestry proportion assigned to the gray wolf cluster (Q_wolf_) estimated using ADMIXTURE at K = 2 for European wolves and dogs. Values range

| Number | Species | Population/Breed | Sample ID | Qwolf |
|---|---|---|---|---|
| 1 | *Canis lupus familiaris* | Golden Retriever | 140447_S11 | 0,000 |
| 2 | *Canis lupus familiaris* | Labrador Retriever | 149323_S6 | 0,000 |
| 3 | *Canis lupus familiaris* | Rottweiler | 165414_S20 | 0,000 |
| 4 | *Canis lupus familiaris* | Rottweiler | 171515_S19 | 0,000 |
| 5 | *Canis lupus familiaris* | Golden Retriever | 173006_S10 | 0,000 |
| 6 | *Canis lupus familiaris* | Labrador Retriever | 173486_S3 | 0,000 |
| 7 | *Canis lupus familiaris* | Akbash | AKBH000001 | 0,019 |
| 8 | *Canis lupus familiaris* | Akbash | AKBH000003 | 0,007 |
| 9 | *Canis lupus familiaris* | Alaskan Malamute | AMAL000001 | 0,106 |
| 10 | *Canis lupus familiaris* | Alaskan Malamute | AMAL000002 | 0,101 |
| 11 | *Canis lupus familiaris* | American Staffordshire Terrier | AMST000001 | 0,000 |
| 12 | *Canis lupus familiaris* | American Staffordshire Terrier | AMST000006 | 0,000 |
| 13 | *Canis lupus familiaris* | Anatolian Shepherd Dog | ANAT000003 | 0,021 |
| 14 | *Canis lupus familiaris* | Anatolian Shepherd Dog | ANAT000004 | 0,020 |
| 15 | *Canis lupus* | Dinaric-Balkan | AP_07M5 | 0,903 |
| 16 | *Canis lupus* | Dinaric-Balkan | AP_082A | 0,844 |
| 17 | *Canis lupus* | Dinaric-Balkan | AP_0844 | 0,868 |
| 18 | *Canis lupus* | Dinaric-Balkan | AP_085T | 1,000 |
| 19 | *Canis lupus* | Dinaric-Balkan | AP_085Y | 0,895 |
| 20 | *Canis lupus* | Dinaric-Balkan | AP_086E | 1,000 |
| 21 | *Canis lupus* | Dinaric-Balkan | AP_086F | 1,000 |
| 22 | *Canis lupus* | Dinaric-Balkan | AP_086Y | 0,897 |
| 23 | *Canis lupus* | Dinaric-Balkan | AP_0870 | 0,890 |
| 24 | *Canis lupus* | Dinaric-Balkan | AP_0872 | 0,929 |
| 25 | *Canis lupus* | Dinaric-Balkan | AP_0878 | 0,905 |
| 26 | *Canis lupus* | Dinaric-Balkan | AP_087A | 1,000 |
| 27 | *Canis lupus* | Dinaric-Balkan | AP_087E | 1,000 |
| 28 | *Canis lupus* | Dinaric-Balkan | AP_087F | 0,867 |
| 29 | *Canis lupus* | Dinaric-Balkan | AP_087L | 1,000 |
| 30 | *Canis lupus* | Dinaric-Balkan | AP_087U | 0,998 |
| 31 | *Canis lupus* | Dinaric-Balkan | AP_087Y | 0,863 |
| 32 | *Canis lupus* | Dinaric-Balkan | AP_0882 | 1,000 |
| 33 | *Canis lupus* | Dinaric-Balkan | AP_0886 | 0,793 |

**Table S3.** Ancestry proportion assigned to the gray wolf cluster (Q_wolf_) estimated using ADMIXTURE at K = 2 for European wolves and dogs. Values range

| Number | Species | Population/Breed | Sample ID | Qwolf |
|---|---|---|---|---|
| 34 | *Canis lupus* | Dinaric-Balkan | AP_088F | 0,909 |
| 35 | *Canis lupus* | Dinaric-Balkan | AP_088L | 1,000 |
| 36 | *Canis lupus* | Dinaric-Balkan | AP_088P | 0,912 |
| 37 | *Canis lupus* | Dinaric-Balkan | AP_08A1 | 1,000 |
| 38 | *Canis lupus* | Dinaric-Balkan | AP_08A3 | 1,000 |
| 39 | *Canis lupus* | Dinaric-Balkan | AP_08A6 | 0,972 |
| 40 | *Canis lupus* | Dinaric-Balkan | AP_08AK | 0,879 |
| 41 | *Canis lupus* | Dinaric-Balkan | AP_08C3 | 1,000 |
| 42 | *Canis lupus* | Dinaric-Balkan | AP_08C5 | 1,000 |
| 43 | *Canis lupus familiaris* | Australian Shepherd | AUSS000001 | 0,000 |
| 44 | *Canis lupus familiaris* | Australian Shepherd | AUSS000003 | 0,000 |
| 45 | *Canis lupus familiaris* | Beauceron | BEAU000004 | 0,000 |
| 46 | *Canis lupus familiaris* | Beauceron | BEAU000007 | 0,000 |
| 47 | *Canis lupus familiaris* | Belgian Sheepdog | BELS000001 | 0,000 |
| 48 | *Canis lupus familiaris* | Belgian Sheepdog | BELS000003 | 0,000 |
| 49 | *Canis lupus familiaris* | Bloodhound | BLDH000002 | 0,000 |
| 50 | *Canis lupus familiaris* | Bloodhound | BLDH000003 | 0,000 |
| 51 | *Canis lupus familiaris* | Belgian Malinois | BMAL000001 | 0,000 |
| 52 | *Canis lupus familiaris* | Belgian Malinois | BMAL000002 | 0,000 |
| 53 | *Canis lupus familiaris* | Bavarian Mountain Scent Hound | BMSH000005 | 0,000 |
| 54 | *Canis lupus familiaris* | Bavarian Mountain Scent Hound | BMSH000006 | 0,000 |
| 55 | *Canis lupus familiaris* | Boerboel | BOER000002 | 0,000 |
| 56 | *Canis lupus familiaris* | Boerboel | BOER000003 | 0,000 |
| 57 | *Canis lupus familiaris* | Boxer | BOXR000001 | 0,000 |
| 58 | *Canis lupus familiaris* | Boxer | BOXR000005 | 0,000 |
| 59 | *Canis lupus familiaris* | Bracco Italiano | BRAC000001 | 0,000 |
| 60 | *Canis lupus familiaris* | Bracco Italiano | BRAC000004 | 0,000 |
| 61 | *Canis lupus familiaris* | Briard | BRIA000005 | 0,000 |
| 62 | *Canis lupus familiaris* | Briard | BRIA000007 | 0,000 |
| 63 | *Canis lupus familiaris* | Bernese Mountain Dog | BRMD000001 | 0,000 |
| 64 | *Canis lupus familiaris* | Bernese Mountain Dog | BRMD000004 | 0,000 |
| 65 | *Canis lupus familiaris* | Black Russian Terrier | BRTR000002 | 0,000 |
| 66 | *Canis lupus familiaris* | Black Russian Terrier | BRTR000004 | 0,000 |

**Table S3.** Ancestry proportion assigned to the gray wolf cluster (Q_wolf_) estimated using ADMIXTURE at K = 2 for European wolves and dogs. Values range

| Number | Species | Population/Breed | Sample ID | Qwolf |
|---|---|---|---|---|
| 67 | *Canis lupus* | Dinaric-Balkan | BiH_244 | 1,000 |
| 68 | *Canis lupus* | Dinaric-Balkan | BiH_245 | 1,000 |
| 69 | *Canis lupus familiaris* | Caucasian Shepherd Dog | CAUC000001 | 0,013 |
| 70 | *Canis lupus familiaris* | Curly-coated Retriever | CCRT000003 | 0,000 |
| 71 | *Canis lupus familiaris* | Curly-coated Retriever | CCRT000004 | 0,000 |
| 72 | *Canis lupus familiaris* | Chesapeake Bay Retriever | CHBR000001 | 0,000 |
| 73 | *Canis lupus familiaris* | Chesapeake Bay Retriever | CHBR000002 | 0,000 |
| 74 | *Canis lupus* | Dinaric-Balkan | CH_0LLU | 0,854 |
| 75 | *Canis lupus* | Dinaric-Balkan | CH_0LLY | 0,860 |
| 76 | *Canis lupus* | Dinaric-Balkan | CH_0LM3 | 0,868 |
| 77 | *Canis lupus* | Dinaric-Balkan | CK_0PET | 0,721 |
| 78 | *Canis lupus* | Karelian | CLUPRU000005 | 0,974 |
| 79 | *Canis lupus* | Karelian | CLUPRU000006 | 0,976 |
| 80 | *Canis lupus* | Karelian | CLUPRU000007 | 0,973 |
| 81 | *Canis lupus* | Karelian | CLUPRU000008 | 0,970 |
| 82 | *Canis lupus* | Karelian | CLUPRU000019 | 0,969 |
| 83 | *Canis lupus* | Karelian | CLUPRU000020 | 0,966 |
| 84 | *Canis lupus familiaris* | Cane Corso | CNCS000003 | 0,000 |
| 85 | *Canis lupus familiaris* | Cane Corso | CNCS000004 | 0,000 |
| 86 | *Canis lupus familiaris* | Collie | COLL000007 | 0,000 |
| 87 | *Canis lupus familiaris* | Collie | COLL000013 | 0,000 |
| 88 | *Canis lupus familiaris* | Chinook | COOK000002 | 0,000 |
| 89 | *Canis lupus familiaris* | Chinook | COOK000007 | 0,000 |
| 90 | *Canis lupus familiaris* | Czechoslovakian Wolfdog | CZEC000003 | 0,280 |
| 91 | *Canis lupus familiaris* | Czechoslovakian Wolfdog | CZWO000001 | 0,280 |
| 92 | *Canis lupus* | Scandinavian | D-00-12 | 1,000 |
| 93 | *Canis lupus* | Scandinavian | D-06-14 | 1,000 |
| 94 | *Canis lupus* | Scandinavian | D-07-16 | 1,000 |
| 95 | *Canis lupus* | Scandinavian | D-08-10 | 1,000 |
| 96 | *Canis lupus* | Scandinavian | D-10-68 | 1,000 |
| 97 | *Canis lupus* | Scandinavian | D-11-58 | 1,000 |
| 98 | *Canis lupus* | Scandinavian | D-85-02 | 1,000 |
| 99 | *Canis lupus familiaris* | Dalmatian | DALM000002 | 0,000 |

**Table S3.** Ancestry proportion assigned to the gray wolf cluster (Q_wolf_) estimated using ADMIXTURE at K = 2 for European wolves and dogs. Values range

| Number | Species | Population/Breed | Sample ID | Qwolf |
|---|---|---|---|---|
| 100 | *Canis lupus familiaris* | Scottish Deerhound | DEER000003 | 0,000 |
| 101 | *Canis lupus familiaris* | Scottish Deerhound | DEER000004 | 0,000 |
| 102 | *Canis lupus familiaris* | Village dog Turkey | DTR001 | 0,021 |
| 103 | *Canis lupus familiaris* | Village dog Turkey | DTR002 | 0,013 |
| 104 | *Canis lupus familiaris* | Village dog Turkey | DTR003 | 0,017 |
| 105 | *Canis lupus familiaris* | Dalmatian | Dalmatian01 | 0,000 |
| 106 | *Canis lupus familiaris* | English Foxhound | EFXH000002 | 0,000 |
| 107 | *Canis lupus familiaris* | English Foxhound | EFXH000003 | 0,000 |
| 108 | *Canis lupus familiaris* | Entlebucher Mountain Dog | ENTB000001 | 0,000 |
| 109 | *Canis lupus familiaris* | Entlebucher Mountain Dog | ENTB000002 | 0,000 |
| 110 | *Canis lupus familiaris* | Estrela Mountain Dog | ESMD000002 | 0,000 |
| 111 | *Canis lupus familiaris* | Estrela Mountain Dog | ESMD000003 | 0,000 |
| 112 | *Canis lupus familiaris* | Flat-Coated Retriever | FLCR000002 | 0,000 |
| 113 | *Canis lupus familiaris* | Flat-Coated Retriever | FlatcoatedRetriever03 | 0,000 |
| 114 | *Canis lupus* | Scandinavian | G100-12 | 1,000 |
| 115 | *Canis lupus* | Scandinavian | G100-14 | 1,000 |
| 116 | *Canis lupus* | Scandinavian | G109-11 | 1,000 |
| 117 | *Canis lupus* | Scandinavian | G126-13 | 1,000 |
| 118 | *Canis lupus* | Scandinavian | G139-12 | 1,000 |
| 119 | *Canis lupus* | Scandinavian | G31-13 | 1,000 |
| 120 | *Canis lupus* | Scandinavian | G37-10 | 1,000 |
| 121 | *Canis lupus* | Scandinavian | G50-12 | 1,000 |
| 122 | *Canis lupus familiaris* | Galgo Espanol | GALG000002 | 0,000 |
| 123 | *Canis lupus familiaris* | Galgo Espanol | GALG000003 | 0,000 |
| 124 | *Canis lupus familiaris* | Gordon Setter | GORD000002 | 0,000 |
| 125 | *Canis lupus familiaris* | Great Pyrenees | GPYR000003 | 0,000 |
| 126 | *Canis lupus familiaris* | Great Pyrenees | GPYR000004 | 0,000 |
| 127 | *Canis lupus familiaris* | German Shepherd | GRSD000002 | 0,000 |
| 128 | *Canis lupus familiaris* | German Shepherd | GRSD000003 | 0,000 |
| 129 | *Canis lupus familiaris* | Greater Swiss Mountain Dog | GSMD000004 | 0,000 |
| 130 | *Canis lupus familiaris* | Greater Swiss Mountain Dog | GreaterSwissMountainDog01 | 0,000 |
| 131 | *Canis lupus familiaris* | Hovawart | HOVA000003 | 0,000 |
| 132 | *Canis lupus familiaris* | Hovawart | HOVA000004 | 0,000 |

**Table S3.** Ancestry proportion assigned to the gray wolf cluster (Q_wolf_) estimated using ADMIXTURE at K = 2 for European wolves and dogs. Values range

| Number | Species | Population/Breed | Sample ID | Qwolf |
|---|---|---|---|---|
| 133 | *Canis lupus familiaris* | Vizsla | HWHP000001 | 0,000 |
| 134 | *Canis lupus familiaris* | Ibizan Hound | IBIZ000010 | 0,000 |
| 135 | *Canis lupus familiaris* | Ibizan Hound | IBIZ000011 | 0,000 |
| 136 | *Canis lupus familiaris* | Irish Setter | IRSE000001 | 0,000 |
| 137 | *Canis lupus familiaris* | Irish Setter | IRSE000003 | 0,000 |
| 138 | *Canis lupus familiaris* | Irish Water Spaniel | IWSP000001 | 0,000 |
| 139 | *Canis lupus familiaris* | Irish Water Spaniel | IWSP000002 | 0,000 |
| 140 | *Canis lupus* | NW Iberia | JAL7480 | 1,000 |
| 141 | *Canis lupus* | NW Iberia | JAL7485 | 0,893 |
| 142 | *Canis lupus* | NW Iberia | JAL7486 | 1,000 |
| 143 | *Canis lupus* | NW Iberia | JAL7491 | 0,973 |
| 144 | *Canis lupus* | NW Iberia | JAL7492 | 0,978 |
| 145 | *Canis lupus familiaris* | Kangal | KANG000001 | 0,021 |
| 146 | *Canis lupus familiaris* | Kangal | KANG000003 | 0,019 |
| 147 | *Canis lupus familiaris* | Kars | KARS000005 | 0,016 |
| 148 | *Canis lupus familiaris* | Kars | KARS000006 | 0,016 |
| 149 | *Canis lupus familiaris* | Kuvasz | KUVZ000002 | 0,000 |
| 150 | *Canis lupus familiaris* | Kuvasz | KUVZ000007 | 0,000 |
| 151 | *Canis lupus* | NW Iberia | L058 | 0,866 |
| 152 | *Canis lupus* | NW Iberia | L166 | 0,999 |
| 153 | *Canis lupus* | NW Iberia | L186 | 0,925 |
| 154 | *Canis lupus* | NW Iberia | L253 | 1,000 |
| 155 | *Canis lupus* | NW Iberia | L352 | 0,844 |
| 156 | *Canis lupus* | NW Iberia | L409 | 1,000 |
| 157 | *Canis lupus* | NW Iberia | L514 | 1,000 |
| 158 | *Canis lupus* | NW Iberia | L547 | 1,000 |
| 159 | *Canis lupus* | NW Iberia | L552 | 1,000 |
| 160 | *Canis lupus* | NW Iberia | L588 | 1,000 |
| 161 | *Canis lupus* | NW Iberia | L781 | 0,918 |
| 162 | *Canis lupus* | NW Iberia | L800 | 0,908 |
| 163 | *Canis lupus* | NW Iberia | L844 | 1,000 |
| 164 | *Canis lupus familiaris* | Lagotto Romagnolo | LAGO000001 | 0,000 |
| 165 | *Canis lupus familiaris* | Lagotto Romagnolo | LAGO000004 | 0,000 |

**Table S3.** Ancestry proportion assigned to the gray wolf cluster (Q_wolf_) estimated using ADMIXTURE at K = 2 for European wolves and dogs. Values range

| Number | Species | Population/Breed | Sample ID | Qwolf |
|---|---|---|---|---|
| 166 | *Canis lupus familiaris* | Catahoula Leopard Dog | LEOP000003 | 0,000 |
| 167 | *Canis lupus familiaris* | Catahoula Leopard Dog | LEOP000008 | 0,000 |
| 168 | *Canis lupus* | Dinaric-Balkan | LUP006817 | 0,980 |
| 169 | *Canis lupus* | Dinaric-Balkan | LUP006818 | 0,952 |
| 170 | *Canis lupus* | Dinaric-Balkan | LUP006819 | 0,970 |
| 171 | *Canis lupus* | Dinaric-Balkan | LUP006820 | 0,946 |
| 172 | *Canis lupus* | Dinaric-Balkan | LUP006821 | 0,975 |
| 173 | *Canis lupus* | Dinaric-Balkan | LUP006822 | 0,972 |
| 174 | *Canis lupus* | Dinaric-Balkan | LUP006823 | 0,977 |
| 175 | *Canis lupus* | Dinaric-Balkan | LUP006824 | 0,964 |
| 176 | *Canis lupus* | Dinaric-Balkan | LUP006825 | 0,978 |
| 177 | *Canis lupus* | Dinaric-Balkan | LUP006826 | 0,973 |
| 178 | *Canis lupus* | Dinaric-Balkan | LUP006827 | 0,969 |
| 179 | *Canis lupus* | Dinaric-Balkan | LUP006828 | 0,976 |
| 180 | *Canis lupus* | NW Iberia | LUP006829 | 0,994 |
| 181 | *Canis lupus* | NW Iberia | LUP006830 | 1,000 |
| 182 | *Canis lupus* | Scandinavian | M-01-06 | 1,000 |
| 183 | *Canis lupus* | Scandinavian | M-06-03 | 1,000 |
| 184 | *Canis lupus* | Scandinavian | M-09-05 | 1,000 |
| 185 | *Canis lupus* | Scandinavian | M-10-04 | 1,000 |
| 186 | *Canis lupus* | Scandinavian | M-98-08 | 1,000 |
| 187 | *Canis lupus* | Dinaric-Balkan | M2C3H | 0,908 |
| 188 | *Canis lupus* | Dinaric-Balkan | M2CHL | 0,863 |
| 189 | *Canis lupus* | Dinaric-Balkan | M2E71 | 0,768 |
| 190 | *Canis lupus familiaris* | Maremma Sheepdog | MARM000006 | 0,000 |
| 191 | *Canis lupus* | Dinaric-Balkan | MSV2AC | 1,000 |
| 192 | *Canis lupus* | NW Iberia | MW122 | 1,000 |
| 193 | *Canis lupus* | NW Iberia | MW127 | 1,000 |
| 194 | *Canis lupus familiaris* | Maremma Sheepdog | Maremma01 | 0,000 |
| 195 | *Canis lupus familiaris* | Mastino Abruzzese | MastinoAbruzzese01 | 0,000 |
| 196 | *Canis lupus familiaris* | Neapolitan Mastiff | NEAP000004 | 0,000 |
| 197 | *Canis lupus familiaris* | Neapolitan Mastiff | NEAP000005 | 0,000 |
| 198 | *Canis lupus familiaris* | Norwegian Elkhound | NELK000002 | 0,000 |

**Table S3.** Ancestry proportion assigned to the gray wolf cluster (Q_wolf_) estimated using ADMIXTURE at K = 2 for European wolves and dogs. Values range

| Number | Species | Population/Breed | Sample ID | Qwolf |
|---|---|---|---|---|
| 199 | *Canis lupus familiaris* | Newfoundland | NEWF000002 | 0,000 |
| 200 | *Canis lupus familiaris* | Newfoundland | NEWF000006 | 0,000 |
| 201 | *Canis lupus familiaris* | Norwegian Elkhound | NorwegianElkhound02 | 0,000 |
| 202 | *Canis lupus familiaris* | Pit Bull Terrier | PITB000001 | 0,000 |
| 203 | *Canis lupus familiaris* | Pit Bull Terrier | PITB000002 | 0,000 |
| 204 | *Canis lupus familiaris* | Portuguese Pointer | POPT000001 | 0,000 |
| 205 | *Canis lupus familiaris* | Portuguese Sheepdog | POSD000005 | 0,000 |
| 206 | *Canis lupus familiaris* | Portuguese Podengo | PPOD000004 | 0,000 |
| 207 | *Canis lupus familiaris* | Portuguese Podengo | PPOP000001 | 0,000 |
| 208 | *Canis lupus familiaris* | Village dog Portugal | PT61 | 0,000 |
| 209 | *Canis lupus familiaris* | Portuguese Water Dog | PTWD000008 | 0,000 |
| 210 | *Canis lupus familiaris* | Pyrenean Mastiff | PYMF000005 | 0,000 |
| 211 | *Canis lupus familiaris* | Pyrenean Mastiff | PYMF000006 | 0,000 |
| 212 | *Canis lupus* | NW Iberia | Penelope | 1,000 |
| 213 | *Canis lupus familiaris* | Portuguese Water Dog | PortugueseWaterDog09 | 0,000 |
| 214 | *Canis lupus familiaris* | Rhodesian Ridgeback | RHOD000001 | 0,000 |
| 215 | *Canis lupus familiaris* | Saarloos Wolfdog | SAAR000002 | 0,290 |
| 216 | *Canis lupus familiaris* | Saarloos Wolfdog | SAAR000005 | 0,277 |
| 217 | *Canis lupus familiaris* | Saluki | SALU000001 | 0,030 |
| 218 | *Canis lupus familiaris* | Saluki | SALU000002 | 0,030 |
| 219 | *Canis lupus familiaris* | Samoyed | SAMO000002 | 0,032 |
| 220 | *Canis lupus familiaris* | Segugio Italiano | SGGO000002 | 0,000 |
| 221 | *Canis lupus familiaris* | Segugio Italiano | SGGO000004 | 0,000 |
| 222 | *Canis lupus familiaris* | Sloughi | SLOU000001 | 0,000 |
| 223 | *Canis lupus familiaris* | Sloughi | SLOU000003 | 0,000 |
| 224 | *Canis lupus familiaris* | Spinone Italiano | SPIN000001 | 0,000 |
| 225 | *Canis lupus familiaris* | Spinone Italiano | SPIN000004 | 0,000 |
| 226 | *Canis lupus familiaris* | Spanish Mastiff | SPMT000002 | 0,000 |
| 227 | *Canis lupus familiaris* | Spanish Mastiff | SPMT000005 | 0,000 |
| 228 | *Canis lupus familiaris* | Spanish Water Dog | SPWD000002 | 0,000 |
| 229 | *Canis lupus familiaris* | Spanish Water Dog | SPWD000003 | 0,000 |
| 230 | *Canis lupus familiaris* | Sarabi | SRAB000001 | 0,020 |
| 231 | *Canis lupus familiaris* | Sarabi | SRAB000002 | 0,024 |

**Table S3.** Ancestry proportion assigned to the gray wolf cluster (Q_wolf_) estimated using ADMIXTURE at K = 2 for European wolves and dogs. Values range

| Number | Species | Population/Breed | Sample ID | Qwolf |
|---|---|---|---|---|
| 232 | *Canis lupus familiaris* | Samoyed | Samoyed01 | 0,026 |
| 233 | *Canis lupus familiaris* | Belgian Tervuren | TERV000005 | 0,000 |
| 234 | *Canis lupus familiaris* | Tosa Inu | TOSA000002 | 0,019 |
| 235 | *Canis lupus familiaris* | Tosa Inu | TOSA000003 | 0,013 |
| 236 | *Canis lupus familiaris* | Turkish Mastiff | TURM000002 | 0,018 |
| 237 | *Canis lupus familiaris* | Turkish Mastiff | TURM000004 | 0,019 |
| 238 | *Canis lupus familiaris* | Belgian Tervuren | TURV000001 | 0,000 |
| 239 | *Canis lupus familiaris* | Treeing Walker Coonhound | TWCH000001 | 0,000 |
| 240 | *Canis lupus familiaris* | Treeing Walker Coonhound | TWCH000002 | 0,000 |
| 241 | *Canis lupus* | Karelian | V113 | 0,983 |
| 242 | *Canis lupus* | Karelian | V114 | 0,969 |
| 243 | *Canis lupus* | Karelian | V115 | 0,976 |
| 244 | *Canis lupus* | Karelian | V116 | 0,968 |
| 245 | *Canis lupus* | Karelian | V117 | 0,976 |
| 246 | *Canis lupus* | Karelian | V119 | 1,000 |
| 247 | *Canis lupus* | Karelian | V120 | 0,976 |
| 248 | *Canis lupus* | Karelian | V126 | 0,991 |
| 249 | *Canis lupus* | Karelian | V132 | 0,970 |
| 250 | *Canis lupus* | Karelian | V134 | 0,996 |
| 251 | *Canis lupus* | Karelian | V136 | 0,981 |
| 252 | *Canis lupus* | Karelian | V141 | 0,997 |
| 253 | *Canis lupus* | Karelian | V143 | 0,995 |
| 254 | *Canis lupus* | Scandinavian | V3064 | 0,533 |
| 255 | *Canis lupus* | Scandinavian | V3065 | 0,536 |
| 256 | *Canis lupus* | Scandinavian | V3069 | 0,539 |
| 257 | *Canis lupus familiaris* | Village dog Afghanistan | VILLAF000001 | 0,033 |
| 258 | *Canis lupus familiaris* | Village dog Afghanistan | VILLAF000002 | 0,051 |
| 259 | *Canis lupus familiaris* | Village dog Afghanistan | VILLAF000003 | 0,038 |
| 260 | *Canis lupus familiaris* | Village dog Azerbaijan | VILLAZ000002 | 0,028 |
| 261 | *Canis lupus familiaris* | Village dog Azerbaijan | VILLAZ000003 | 0,021 |
| 262 | *Canis lupus familiaris* | Village dog Azerbaijan | VILLAZ000004 | 0,022 |
| 263 | *Canis lupus familiaris* | Village dog Azerbaijan | VILLAZ000005 | 0,020 |
| 264 | *Canis lupus familiaris* | Village dog Azerbaijan | VILLAZ000007 | 0,017 |

**Table S3.** Ancestry proportion assigned to the gray wolf cluster (Q_wolf_) estimated using ADMIXTURE at K = 2 for European wolves and dogs. Values range

| Number | Species | Population/Breed | Sample ID | Qwolf |
|---|---|---|---|---|
| 265 | *Canis lupus familiaris* | Village dog Bulgaria | VILLBG000001 | 0,000 |
| 266 | *Canis lupus familiaris* | Village dog Bulgaria | VILLBG000002 | 0,000 |
| 267 | *Canis lupus familiaris* | Village dog Congo | VILLCG000001 | 0,000 |
| 268 | *Canis lupus familiaris* | Village dog Congo | VILLCG000004 | 0,000 |
| 269 | *Canis lupus familiaris* | Village dog Congo | VILLCG000006 | 0,000 |
| 270 | *Canis lupus familiaris* | Village dog Congo | VILLCG000011 | 0,000 |
| 271 | *Canis lupus familiaris* | Village dog Congo | VILLCG000013 | 0,000 |
| 272 | *Canis lupus familiaris* | Village dog China | VILLCN000005 | 0,171 |
| 273 | *Canis lupus familiaris* | Village dog China | VILLCN000013 | 0,164 |
| 274 | *Canis lupus familiaris* | Village dog China | VILLCN000088 | 0,135 |
| 275 | *Canis lupus familiaris* | Village dog China | VILLCN000145 | 0,117 |
| 276 | *Canis lupus familiaris* | Village dog China | VILLCN000153 | 0,073 |
| 277 | *Canis lupus familiaris* | Village dog Iran | VILLIR000021 | 0,023 |
| 278 | *Canis lupus familiaris* | Village dog Iran | VILLIR000023 | 0,000 |
| 279 | *Canis lupus familiaris* | Village dog Iran | VILLIR000024 | 0,034 |
| 280 | *Canis lupus familiaris* | Village dog Iran | VILLIR000026 | 0,027 |
| 281 | *Canis lupus familiaris* | Village dog Iran | VILLIR000027 | 0,054 |
| 282 | *Canis lupus familiaris* | Village dog Kenya | VILLKE000002 | 0,014 |
| 283 | *Canis lupus familiaris* | Village dog Kenya | VILLKE000009 | 0,011 |
| 284 | *Canis lupus familiaris* | Village dog Kenya | VILLKE000010 | 0,023 |
| 285 | *Canis lupus familiaris* | Village dog Kenya | VILLKE000011 | 0,022 |
| 286 | *Canis lupus familiaris* | Village dog Kenya | VILLKE000019 | 0,023 |
| 287 | *Canis lupus familiaris* | Village dog Liberia | VILLLR000005 | 0,009 |
| 288 | *Canis lupus familiaris* | Village dog Liberia | VILLLR000011 | 0,014 |
| 289 | *Canis lupus familiaris* | Village dog Liberia | VILLLR000014 | 0,024 |
| 290 | *Canis lupus familiaris* | Village dog Liberia | VILLLR000016 | 0,000 |
| 291 | *Canis lupus familiaris* | Village dog Liberia | VILLLR000017 | 0,004 |
| 292 | *Canis lupus familiaris* | Village dog Tajikistan | VILLTJ000001 | 0,040 |
| 293 | *Canis lupus familiaris* | Village dog Uzbekistan | VILLUZ000002 | 0,010 |
| 294 | *Canis lupus familiaris* | Village dog Uzbekistan | VILLUZ000003 | 0,019 |
| 295 | *Canis lupus familiaris* | Village dog Uzbekistan | VILLUZ000004 | 0,020 |
| 296 | *Canis lupus familiaris* | Village dog Uzbekistan | VILLUZ000005 | 0,022 |
| 297 | *Canis lupus familiaris* | Village dog Uzbekistan | VILLUZ000006 | 0,016 |

**Table S3.** Ancestry proportion assigned to the gray wolf cluster (Q_wolf_) estimated using ADMIXTURE at K = 2 for European wolves and dogs. Values range

| Number | Species | Population/Breed | Sample ID | Qwolf |
|---|---|---|---|---|
| 298 | *Canis lupus familiaris* | Vizsla | VIZS000007 | 0,000 |
| 299 | *Canis lupus* | Italian Peninsula | W1023 | 1,000 |
| 300 | *Canis lupus* | Karelian | W11 | 0,995 |
| 301 | *Canis lupus familiaris* | dogsItaly | W1129 | 0,000 |
| 302 | *Canis lupus familiaris* | dogsItaly | W1208 | 0,000 |
| 303 | *Canis lupus familiaris* | dogsItaly | W1297 | 0,005 |
| 304 | *Canis lupus* | Karelian | W14 | 0,995 |
| 305 | *Canis lupus familiaris* | dogsItaly | W1431 | 0,000 |
| 306 | *Canis lupus familiaris* | dogsItaly | W1432 | 0,000 |
| 307 | *Canis lupus* | Italian Peninsula | W1433 | 0,831 |
| 308 | *Canis lupus familiaris* | dogsItaly | W1449 | 0,000 |
| 309 | *Canis lupus* | Italian Peninsula | W1456 | 0,901 |
| 310 | *Canis lupus* | Italian Peninsula | W1551 | 0,993 |
| 311 | *Canis lupus familiaris* | dogsItaly | W1564 | 0,000 |
| 312 | *Canis lupus familiaris* | dogsItaly | W1578 | 0,000 |
| 313 | *Canis lupus* | Central European | W1593 | 0,957 |
| 314 | *Canis lupus* | Karelian | W16 | 1,000 |
| 315 | *Canis lupus* | Italian Peninsula | W1869 | 0,910 |
| 316 | *Canis lupus* | Central European | W191598 | 0,470 |
| 317 | *Canis lupus* | Central European | W200546 | 0,719 |
| 318 | *Canis lupus* | Italian Peninsula | W2030 | 1,000 |
| 319 | *Canis lupus* | Karelian | W21 | 1,000 |
| 320 | *Canis lupus* | Italian Peninsula | W2124 | 0,992 |
| 321 | *Canis lupus* | Italian Peninsula | W2185 | 0,983 |
| 322 | *Canis lupus* | Italian Peninsula | W2192 | 0,887 |
| 323 | *Canis lupus* | Italian Peninsula | W2193 | 0,983 |
| 324 | *Canis lupus* | Central European | W224957 | 0,471 |
| 325 | *Canis lupus* | Italian Peninsula | W2265 | 0,997 |
| 326 | *Canis lupus* | Central European | W226855 | 0,801 |
| 327 | *Canis lupus* | Italian Peninsula | W2280 | 1,000 |
| 328 | *Canis lupus* | Italian Peninsula | W2291 | 0,992 |
| 329 | *Canis lupus* | Italian Peninsula | W2371F | 1,000 |
| 330 | *Canis lupus* | Italian Peninsula | W2419 | 0,877 |

**Table S3.** Ancestry proportion assigned to the gray wolf cluster (Q_wolf_) estimated using ADMIXTURE at K = 2 for European wolves and dogs. Values range

| Number | Species | Population/Breed | Sample ID | Qwolf |
|---|---|---|---|---|
| 331 | *Canis lupus* | Italian Peninsula | W2430F | 0,982 |
| 332 | *Canis lupus* | Italian Peninsula | W2470 | 0,989 |
| 333 | *Canis lupus* | Italian Peninsula | W2487F | 1,000 |
| 334 | *Canis lupus* | Italian Peninsula | W2500M | 1,000 |
| 335 | *Canis lupus* | Italian Peninsula | W2594M | 1,000 |
| 336 | *Canis lupus* | Karelian | W26 | 1,000 |
| 337 | *Canis lupus* | Italian Peninsula | W2623M | 0,988 |
| 338 | *Canis lupus* | Italian Peninsula | W2685F | 0,990 |
| 339 | *Canis lupus* | Italian Peninsula | W2707M | 1,000 |
| 340 | *Canis lupus* | Italian Peninsula | W2725 | 0,820 |
| 341 | *Canis lupus* | Italian Peninsula | W2878M | 1,000 |
| 342 | *Canis lupus* | Italian Peninsula | W2883F | 1,000 |
| 343 | *Canis lupus* | Italian Peninsula | W2886 | 0,977 |
| 344 | *Canis lupus* | Karelian | W29 | 1,000 |
| 345 | *Canis lupus* | Italian Peninsula | W2910 | 1,000 |
| 346 | *Canis lupus* | Italian Peninsula | W2951 | 1,000 |
| 347 | *Canis lupus* | Karelian | W32 | 1,000 |
| 348 | *Canis lupus* | Karelian | W4 | 1,000 |
| 349 | *Canis lupus* | Karelian | W44 | 0,995 |
| 350 | *Canis lupus* | Karelian | W45 | 0,990 |
| 351 | *Canis lupus* | Italian Peninsula | W893 | 1,000 |
| 352 | *Canis lupus familiaris* | Weimaraner | WEIM000002 | 0,000 |
| 353 | *Canis lupus familiaris* | Weimaraner | WEIM000005 | 0,000 |
| 354 | *Canis lupus familiaris* | Wirehaired Pointing Griffon | WHPG000001 | 0,000 |
| 355 | *Canis lupus familiaris* | Wirehaired Pointing Griffon | WHPG000002 | 0,000 |
| 356 | *Canis lupus* | Italian Peninsula | WIT11 | 1,000 |
| 357 | *Canis lupus* | Italian Peninsula | WIT13 | 1,000 |
| 358 | *Canis lupus* | Italian Peninsula | WIT14 | 1,000 |
| 359 | *Canis lupus* | Italian Peninsula | WIT16 | 1,000 |
| 360 | *Canis lupus* | Italian Peninsula | WIT17 | 0,993 |
| 361 | *Canis lupus* | Italian Peninsula | WIT18 | 0,916 |
| 362 | *Canis lupus* | Italian Peninsula | WIT19 | 0,986 |
| 363 | *Canis lupus* | Italian Peninsula | WIT20 | 1,000 |

**Table S3.** Ancestry proportion assigned to the gray wolf cluster (Q_wolf_) estimated using ADMIXTURE at K = 2 for European wolves and dogs. Values range

| Number | Species | Population/Breed | Sample ID | Qwolf |
|---|---|---|---|---|
| 364 | *Canis lupus* | Italian Peninsula | WIT21 | 0,986 |
| 365 | *Canis lupus* | Italian Peninsula | WIT22 | 1,000 |
| 366 | *Canis lupus* | Italian Peninsula | WIT23 | 1,000 |
| 367 | *Canis lupus familiaris* | Turkish Zerdava | ZERD000002 | 0,007 |
| 368 | *Canis lupus familiaris* | Turkish Zerdava | ZERD000003 | 0,007 |
| 369 | *Canis lupus* | NW Iberia | ptw | 1,000 |
| 370 | *Canis lupus* | NW Iberia | spw | 0,847 |
| 371 | *Canis lupus* | Italian Peninsula | w361703 | 1,000 |
| 372 | *Canis lupus* | Italian Peninsula | w361756 | 0,959 |
| 373 | *Canis lupus* | Italian Peninsula | w484049 | 0,895 |
| 374 | *Canis lupus* | Italian Peninsula | w4902419 | 0,941 |
| 375 | *Canis lupus* | Italian Peninsula | w4926402 | 0,946 |
| 376 | *Canis lupus* | NW Iberia | wSierraMorena | 0,633 |

[illegible] from 0 (entirely assigned to the domestic dog cluster) to 1 (entirely assigned to the gray wolf cluster). Samples with Q_wolf_ < 0.95 were considered potent

from 0 (entirely assigned to the domestic dog cluster) to 1 (entirely assigned to the gray wolf cluster). Samples with Q_wolf_ < 0.95 were considered potent

from 0 (entirely assigned to the domestic dog cluster) to 1 (entirely assigned to the gray wolf cluster). Samples with Q_wolf_ < 0.95 were considered potent

from 0 (entirely assigned to the domestic dog cluster) to 1 (entirely assigned to the gray wolf cluster). Samples with Q_wolf_ < 0.95 were considered potent

from 0 (entirely assigned to the domestic dog cluster) to 1 (entirely assigned to the gray wolf cluster). Samples with Q_wolf_ < 0.95 were considered potent

: from 0 (entirely assigned to the domestic dog cluster) to 1 (entirely assigned to the gray wolf cluster). Samples with Q_wolf_ < 0.95 were considered potent

from 0 (entirely assigned to the domestic dog cluster) to 1 (entirely assigned to the gray wolf cluster). Samples with Q_wolf_ < 0.95 were considered potent

from 0 (entirely assigned to the domestic dog cluster) to 1 (entirely assigned to the gray wolf cluster). Samples with Q_wolf_ < 0.95 were considered potent

: from 0 (entirely assigned to the domestic dog cluster) to 1 (entirely assigned to the gray wolf cluster). Samples with Q_wolf_ < 0.95 were considered potent [illegible] from 0 (entirely assigned to the domestic dog cluster) to 1 (entirely assigned to the gray wolf cluster). Samples with Q_wolf_ < 0.95 were considered potent

from 0 (entirely assigned to the domestic dog cluster) to 1 (entirely assigned to the gray wolf cluster). Samples with Q_wolf_ < 0.95 were considered potent

from 0 (entirely assigned to the domestic dog cluster) to 1 (entirely assigned to the gray wolf cluster). Samples with Q_wolf_ < 0.95 were considered potent

ially admixed and were excluded from subsequent analyses.

ially admixed and were excluded from subsequent analyses.

ially admixed and were excluded from subsequent analyses.

ially admixed and were excluded from subsequent analyses.

ially admixed and were excluded from subsequent analyses.

ially admixed and were excluded from subsequent analyses.

ially admixed and were excluded from subsequent analyses.

ially admixed and were excluded from subsequent analyses.

ially admixed and were excluded from subsequent analyses.

ially admixed and were excluded from subsequent analyses.

ially admixed and were excluded from subsequent analyses.

ially admixed and were excluded from subsequent analyses.

**Table S4.** Ancestry proportion assigned to the gray wolf cluster (Q_wolf_) estimated using ADMIXTURE at K = 2 for Asian wolves and

| **Number** | **Species** | **Population/Breed** |
|---|---|---|
| 1 | *Canis lupus familiaris* | Golden Retriever |
| 2 | *Canis lupus familiaris* | Labrador Retriever |
| 3 | *Canis lupus familiaris* | Rottweiler |
| 4 | *Canis lupus familiaris* | Rottweiler |
| 5 | *Canis lupus familiaris* | Golden Retriever |
| 6 | *Canis lupus familiaris* | Labrador Retriever |
| 7 | *Canis lupus familiaris* | Akbash |
| 8 | *Canis lupus familiaris* | Akbash |
| 9 | *Canis lupus familiaris* | Alaskan Malamute |
| 10 | *Canis lupus familiaris* | Alaskan Malamute |
| 11 | *Canis lupus familiaris* | American Staffordshire Terrier |
| 12 | *Canis lupus familiaris* | American Staffordshire Terrier |
| 13 | *Canis lupus familiaris* | Anatolian Shepherd Dog |
| 14 | *Canis lupus familiaris* | Anatolian Shepherd Dog |
| 15 | *Canis lupus familiaris* | Australian Shepherd |
| 16 | *Canis lupus familiaris* | Australian Shepherd |
| 17 | *Canis lupus familiaris* | Beauceron |
| 18 | *Canis lupus familiaris* | Beauceron |
| 19 | *Canis lupus familiaris* | Belgian Sheepdog |
| 20 | *Canis lupus familiaris* | Belgian Sheepdog |
| 21 | *Canis lupus* | East Asia |
| 22 | *Canis lupus* | East Asia |
| 23 | *Canis lupus familiaris* | Bloodhound |
| 24 | *Canis lupus familiaris* | Bloodhound |
| 25 | *Canis lupus familiaris* | Belgian Malinois |
| 26 | *Canis lupus familiaris* | Belgian Malinois |
| 27 | *Canis lupus familiaris* | Bavarian Mountain Scent Hound |
| 28 | *Canis lupus familiaris* | Bavarian Mountain Scent Hound |
| 29 | *Canis lupus familiaris* | Boerboel |
| 30 | *Canis lupus familiaris* | Boerboel |
| 31 | *Canis lupus familiaris* | Boxer |
| 32 | *Canis lupus familiaris* | Boxer |
| 33 | *Canis lupus familiaris* | Bracco Italiano |

**Table S4.** Ancestry proportion assigned to the gray wolf cluster (Q_wolf_) estimated using ADMIXTURE at K = 2 for Asian wolves and

| Number | Species | Population/Breed |
|---|---|---|
| 34 | *Canis lupus familiaris* | Bracco Italiano |
| 35 | *Canis lupus familiaris* | Briard |
| 36 | *Canis lupus familiaris* | Briard |
| 37 | *Canis lupus familiaris* | Bernese Mountain Dog |
| 38 | *Canis lupus familiaris* | Bernese Mountain Dog |
| 39 | *Canis lupus familiaris* | Black Russian Terrier |
| 40 | *Canis lupus familiaris* | Black Russian Terrier |
| 41 | *Canis lupus familiaris* | Caucasian Shepherd Dog |
| 42 | *Canis lupus familiaris* | Curly-coated Retriever |
| 43 | *Canis lupus familiaris* | Curly-coated Retriever |
| 44 | *Canis lupus familiaris* | Chesapeake Bay Retriever |
| 45 | *Canis lupus familiaris* | Chesapeake Bay Retriever |
| 46 | *Canis lupus* | West Asia |
| 47 | *Canis lupus* | East Asia |
| 48 | *Canis lupus* | East Asia |
| 49 | *Canis lupus* | East Asia |
| 50 | *Canis lupus* | East Asia |
| 51 | *Canis lupus* | East Asia |
| 52 | *Canis lupus* | East Asia |
| 53 | *Canis lupus* | East Asia |
| 54 | *Canis lupus* | East Asia |
| 55 | *Canis lupus* | West Asia |
| 56 | *Canis lupus* | West Asia |
| 57 | *Canis lupus* | West Asia |
| 58 | *Canis lupus* | West Asia |
| 59 | *Canis lupus* | West Asia |
| 60 | *Canis lupus* | West Asia |
| 61 | *Canis lupus* | Central Asia |
| 62 | *Canis lupus* | Central Asia |
| 63 | *Canis lupus* | Central Asia |
| 64 | *Canis lupus* | Central Asia |
| 65 | *Canis lupus* | Central Asia |
| 66 | *Canis lupus* | East Asia |

**Table S4.** Ancestry proportion assigned to the gray wolf cluster (Q_wolf_) estimated using ADMIXTURE at K = 2 for Asian wolves and

| Number | Species | Population/Breed |
| --- | --- | --- |
| 67 | *Canis lupus* | West Asia |
| 68 | *Canis lupus* | Central Asia |
| 69 | *Canis lupus* | Central Asia |
| 70 | *Canis lupus* | Central Asia |
| 71 | *Canis lupus* | Central Asia |
| 72 | *Canis lupus* | Central Asia |
| 73 | *Canis lupus* | Central Asia |
| 74 | *Canis lupus* | Central Asia |
| 75 | *Canis lupus familiaris* | Cane Corso |
| 76 | *Canis lupus familiaris* | Cane Corso |
| 77 | *Canis lupus familiaris* | Collie |
| 78 | *Canis lupus familiaris* | Collie |
| 79 | *Canis lupus familiaris* | Chinook |
| 80 | *Canis lupus familiaris* | Chinook |
| 81 | *Canis lupus familiaris* | Czechoslovakian Wolfdog |
| 82 | *Canis lupus familiaris* | Czechoslovakian Wolfdog |
| 83 | *Canis lupus familiaris* | Dalmatian |
| 84 | *Canis lupus familiaris* | Scottish Deerhound |
| 85 | *Canis lupus familiaris* | Scottish Deerhound |
| 86 | *Canis lupus familiaris* | Village dog Turkey |
| 87 | *Canis lupus familiaris* | Village dog Turkey |
| 88 | *Canis lupus familiaris* | Village dog Turkey |
| 89 | *Canis lupus familiaris* | Dalmatian |
| 90 | *Canis lupus familiaris* | English Foxhound |
| 91 | *Canis lupus familiaris* | English Foxhound |
| 92 | *Canis lupus familiaris* | Entlebucher Mountain Dog |
| 93 | *Canis lupus familiaris* | Entlebucher Mountain Dog |
| 94 | *Canis lupus familiaris* | Estrela Mountain Dog |
| 95 | *Canis lupus familiaris* | Estrela Mountain Dog |
| 96 | *Canis lupus familiaris* | Flat-Coated Retriever |
| 97 | *Canis lupus familiaris* | Flat-Coated Retriever |
| 98 | *Canis lupus familiaris* | Galgo Espanol |
| 99 | *Canis lupus familiaris* | Galgo Espanol |

**Table S4.** Ancestry proportion assigned to the gray wolf cluster (Q_wolf_) estimated using ADMIXTURE at K = 2 for Asian wolves and

| Number | Species | Population/Breed |
|---|---|---|
| 100 | *Canis lupus familiaris* | Gordon Setter |
| 101 | *Canis lupus familiaris* | Great Pyrenees |
| 102 | *Canis lupus familiaris* | Great Pyrenees |
| 103 | *Canis lupus familiaris* | German Shepherd |
| 104 | *Canis lupus familiaris* | German Shepherd |
| 105 | *Canis lupus familiaris* | Greater Swiss Mountain Dog |
| 106 | *Canis lupus familiaris* | Greater Swiss Mountain Dog |
| 107 | *Canis lupus familiaris* | Hovawart |
| 108 | *Canis lupus familiaris* | Hovawart |
| 109 | *Canis lupus familiaris* | Vizsla |
| 110 | *Canis lupus familiaris* | Ibizan Hound |
| 111 | *Canis lupus familiaris* | Ibizan Hound |
| 112 | *Canis lupus familiaris* | Irish Setter |
| 113 | *Canis lupus familiaris* | Irish Setter |
| 114 | *Canis lupus familiaris* | Irish Water Spaniel |
| 115 | *Canis lupus familiaris* | Irish Water Spaniel |
| 116 | *Canis lupus* | West Asia |
| 117 | *Canis lupus familiaris* | Kangal |
| 118 | *Canis lupus familiaris* | Kangal |
| 119 | *Canis lupus familiaris* | Kars |
| 120 | *Canis lupus familiaris* | Kars |
| 121 | *Canis lupus familiaris* | Kuvasz |
| 122 | *Canis lupus familiaris* | Kuvasz |
| 123 | *Canis lupus familiaris* | Lagotto Romagnolo |
| 124 | *Canis lupus familiaris* | Lagotto Romagnolo |
| 125 | *Canis lupus familiaris* | Catahoula Leopard Dog |
| 126 | *Canis lupus familiaris* | Catahoula Leopard Dog |
| 127 | *Canis lupus* | East Asia |
| 128 | *Canis lupus* | East Asia |
| 129 | *Canis lupus* | East Asia |
| 130 | *Canis lupus* | East Asia |
| 131 | *Canis lupus* | East Asia |
| 132 | *Canis lupus familiaris* | Maremma Sheepdog |

**Table S4.** Ancestry proportion assigned to the gray wolf cluster (Q_wolf_) estimated using ADMIXTURE at K = 2 for Asian wolves and

| **Number** | **Species** | **Population/Breed** |
|---|---|---|
| 133 | *Canis lupus familiaris* | Maremma Sheepdog |
| 134 | *Canis lupus familiaris* | Mastino Abruzzese |
| 135 | *Canis lupus* | Central Asia |
| 136 | *Canis lupus familiaris* | Neapolitan Mastiff |
| 137 | *Canis lupus familiaris* | Neapolitan Mastiff |
| 138 | *Canis lupus familiaris* | Norwegian Elkhound |
| 139 | *Canis lupus familiaris* | Newfoundland |
| 140 | *Canis lupus familiaris* | Newfoundland |
| 141 | *Canis lupus familiaris* | Norwegian Elkhound |
| 142 | *Canis lupus familiaris* | Pit Bull Terrier |
| 143 | *Canis lupus familiaris* | Pit Bull Terrier |
| 144 | *Canis lupus familiaris* | Portuguese Pointer |
| 145 | *Canis lupus familiaris* | Portuguese Sheepdog |
| 146 | *Canis lupus familiaris* | Portuguese Podengo |
| 147 | *Canis lupus familiaris* | Portuguese Podengo |
| 148 | *Canis lupus familiaris* | Village dog Portugal |
| 149 | *Canis lupus familiaris* | Portuguese Water Dog |
| 150 | *Canis lupus familiaris* | Pyrenean Mastiff |
| 151 | *Canis lupus familiaris* | Pyrenean Mastiff |
| 152 | *Canis lupus familiaris* | Portuguese Water Dog |
| 153 | *Canis lupus familiaris* | Rhodesian Ridgeback |
| 154 | *Canis lupus* | East Asia |
| 155 | *Canis lupus familiaris* | Saarloos Wolfdog |
| 156 | *Canis lupus familiaris* | Saarloos Wolfdog |
| 157 | *Canis lupus familiaris* | Saluki |
| 158 | *Canis lupus familiaris* | Saluki |
| 159 | *Canis lupus familiaris* | Samoyed |
| 160 | *Canis lupus familiaris* | Segugio Italiano |
| 161 | *Canis lupus familiaris* | Segugio Italiano |
| 162 | *Canis lupus familiaris* | Sloughi |
| 163 | *Canis lupus familiaris* | Sloughi |
| 164 | *Canis lupus familiaris* | Spinone Italiano |
| 165 | *Canis lupus familiaris* | Spinone Italiano |

**Table S4.** Ancestry proportion assigned to the gray wolf cluster (Q_wolf_) estimated using ADMIXTURE at K = 2 for Asian wolves and

| Number | Species | Population/Breed |
|---|---|---|
| 166 | *Canis lupus familiaris* | Spanish Mastiff |
| 167 | *Canis lupus familiaris* | Spanish Mastiff |
| 168 | *Canis lupus familiaris* | Spanish Water Dog |
| 169 | *Canis lupus familiaris* | Spanish Water Dog |
| 170 | *Canis lupus familiaris* | Sarabi |
| 171 | *Canis lupus familiaris* | Sarabi |
| 172 | *Canis lupus familiaris* | Samoyed |
| 173 | *Canis lupus familiaris* | Belgian Tervuren |
| 174 | *Canis lupus familiaris* | Tosa Inu |
| 175 | *Canis lupus familiaris* | Tosa Inu |
| 176 | *Canis lupus familiaris* | Turkish Mastiff |
| 177 | *Canis lupus familiaris* | Turkish Mastiff |
| 178 | *Canis lupus familiaris* | Belgian Tervuren |
| 179 | *Canis lupus familiaris* | Treeing Walker Coonhound |
| 180 | *Canis lupus familiaris* | Treeing Walker Coonhound |
| 181 | *Canis lupus familiaris* | Village dog Afghanistan |
| 182 | *Canis lupus familiaris* | Village dog Afghanistan |
| 183 | *Canis lupus familiaris* | Village dog Afghanistan |
| 184 | *Canis lupus familiaris* | Village dog Azerbaijan |
| 185 | *Canis lupus familiaris* | Village dog Azerbaijan |
| 186 | *Canis lupus familiaris* | Village dog Azerbaijan |
| 187 | *Canis lupus familiaris* | Village dog Azerbaijan |
| 188 | *Canis lupus familiaris* | Village dog Azerbaijan |
| 189 | *Canis lupus familiaris* | Village dog Bulgaria |
| 190 | *Canis lupus familiaris* | Village dog Bulgaria |
| 191 | *Canis lupus familiaris* | Village dog Congo |
| 192 | *Canis lupus familiaris* | Village dog Congo |
| 193 | *Canis lupus familiaris* | Village dog Congo |
| 194 | *Canis lupus familiaris* | Village dog Congo |
| 195 | *Canis lupus familiaris* | Village dog Congo |
| 196 | *Canis lupus familiaris* | Village dog China |
| 197 | *Canis lupus familiaris* | Village dog China |
| 198 | *Canis lupus familiaris* | Village dog China |

**Table S4.** Ancestry proportion assigned to the gray wolf cluster (Q_wolf_) estimated using ADMIXTURE at K = 2 for Asian wolves and

| Number | Species | Population/Breed |
|---|---|---|
| 199 | *Canis lupus familiaris* | Village dog China |
| 200 | *Canis lupus familiaris* | Village dog China |
| 201 | *Canis lupus familiaris* | Village dog Iran |
| 202 | *Canis lupus familiaris* | Village dog Iran |
| 203 | *Canis lupus familiaris* | Village dog Iran |
| 204 | *Canis lupus familiaris* | Village dog Iran |
| 205 | *Canis lupus familiaris* | Village dog Iran |
| 206 | *Canis lupus familiaris* | Village dog Kenya |
| 207 | *Canis lupus familiaris* | Village dog Kenya |
| 208 | *Canis lupus familiaris* | Village dog Kenya |
| 209 | *Canis lupus familiaris* | Village dog Kenya |
| 210 | *Canis lupus familiaris* | Village dog Kenya |
| 211 | *Canis lupus familiaris* | Village dog Liberia |
| 212 | *Canis lupus familiaris* | Village dog Liberia |
| 213 | *Canis lupus familiaris* | Village dog Liberia |
| 214 | *Canis lupus familiaris* | Village dog Liberia |
| 215 | *Canis lupus familiaris* | Village dog Liberia |
| 216 | *Canis lupus familiaris* | Village dog Tajikistan |
| 217 | *Canis lupus familiaris* | Village dog Uzbekistan |
| 218 | *Canis lupus familiaris* | Village dog Uzbekistan |
| 219 | *Canis lupus familiaris* | Village dog Uzbekistan |
| 220 | *Canis lupus familiaris* | Village dog Uzbekistan |
| 221 | *Canis lupus familiaris* | Village dog Uzbekistan |
| 222 | *Canis lupus familiaris* | Vizsla |
| 223 | *Canis lupus familiaris* | Weimaraner |
| 224 | *Canis lupus familiaris* | Weimaraner |
| 225 | *Canis lupus familiaris* | Wirehaired Pointing Griffon |
| 226 | *Canis lupus familiaris* | Wirehaired Pointing Griffon |
| 227 | *Canis lupus* | Turkey |
| 228 | *Canis lupus* | Turkey |
| 229 | *Canis lupus* | Turkey |
| 230 | *Canis lupus* | Turkey |
| 231 | *Canis lupus* | Turkey |

**Table S4.** Ancestry proportion assigned to the gray wolf cluster (Q_wolf_) estimated using ADMIXTURE at K = 2 for Asian wolves and

| Number | Species | Population/Breed |
|---|---|---|
| 232 | *Canis lupus* | Turkey |
| 233 | *Canis lupus* | Turkey |
| 234 | *Canis lupus* | Turkey |
| 235 | *Canis lupus* | Turkey |
| 236 | *Canis lupus* | Turkey |
| 237 | *Canis lupus* | Turkey |
| 238 | *Canis lupus* | Turkey |
| 239 | *Canis lupus* | Turkey |
| 240 | *Canis lupus* | Turkey |
| 241 | *Canis lupus* | Turkey |
| 242 | *Canis lupus* | Turkey |
| 243 | *Canis lupus familiaris* | Turkish Zerdava |
| 244 | *Canis lupus familiaris* | Turkish Zerdava |
| 245 | *Canis lupus* | Indian |
| 246 | *Canis lupus* | West Asia |

dogs. Values range from 0 (entirely assigned to the domestic dog cluster) to 1 (entirely assigned to the gray wolf cluster). Samples with Q_wolf_ < 0.95 were co

| Sample | Qwolf |
|---|---|
| 140447_S11 | 0,000 |
| 149323_S6 | 0,000 |
| 165414_S20 | 0,000 |
| 171515_S19 | 0,000 |
| 173006_S10 | 0,000 |
| 173486_S3 | 0,000 |
| AKBH000001 | 0,036 |
| AKBH000003 | 0,021 |
| AMAL000001 | 0,150 |
| AMAL000002 | 0,144 |
| AMST000001 | 0,000 |
| AMST000006 | 0,000 |
| ANAT000003 | 0,038 |
| ANAT000004 | 0,038 |
| AUSS000001 | 0,000 |
| AUSS000003 | 0,000 |
| BEAU000004 | 0,000 |
| BEAU000007 | 0,000 |
| BELS000001 | 0,000 |
| BELS000003 | 0,000 |
| BGI-01505070006 | 0,998 |
| BGI-01505070007 | 1,000 |
| BLDH000002 | 0,000 |
| BLDH000003 | 0,000 |
| BMAL000001 | 0,000 |
| BMAL000002 | 0,000 |
| BMSH000005 | 0,000 |
| BMSH000006 | 0,000 |
| BOER000002 | 0,000 |
| BOER000003 | 0,000 |
| BOXR000001 | 0,000 |
| BOXR000005 | 0,000 |
| BRAC000001 | 0,000 |

dogs. Values range from 0 (entirely assigned to the domestic dog cluster) to 1 (entirely assigned to the gray wolf cluster). Samples with Q_wolf_ < 0.95 were co

| **Sample** | **Qwolf** |
|---|---|
| BRAC000004 | 0,000 |
| BRIA000005 | 0,000 |
| BRIA000007 | 0,000 |
| BRMD000001 | 0,000 |
| BRMD000004 | 0,000 |
| BRTR000002 | 0,000 |
| BRTR000004 | 0,000 |
| CAUC000001 | 0,027 |
| CCRT000003 | 0,000 |
| CCRT000004 | 0,000 |
| CHBR000001 | 0,000 |
| CHBR000002 | 0,000 |
| CLUPAZ000001 | 1,000 |
| CLUPCN000001 | 0,965 |
| CLUPCN000004 | 1,000 |
| CLUPCN000005 | 1,000 |
| CLUPCN000006 | 1,000 |
| CLUPCN000007 | 1,000 |
| CLUPCN000008 | 0,958 |
| CLUPCN000009 | 1,000 |
| CLUPCN000010 | 1,000 |
| CLUPIR000001 | 1,000 |
| CLUPIR000002 | 0,956 |
| CLUPIR000003 | 1,000 |
| CLUPIR000004 | 0,955 |
| CLUPIR000005 | 1,000 |
| CLUPIR000006 | 1,000 |
| CLUPKG000001 | 0,946 |
| CLUPKZ000002 | 1,000 |
| CLUPRU000001 | 0,992 |
| CLUPRU000002 | 0,983 |
| CLUPRU000003 | 0,972 |
| CLUPRU000004 | 0,966 |

dogs. Values range from 0 (entirely assigned to the domestic dog cluster) to 1 (entirely assigned to the gray wolf cluster). Samples with Q_wolf_ < 0.95 were co

| Sample | Qwolf |
|---|---|
| CLUPRU000018 | 0,944 |
| CLUPTJ000001 | 1,000 |
| CLUPTJ000002 | 1,000 |
| CLUPTJ000003 | 1,000 |
| CLUPTJ000004 | 1,000 |
| CLUPTJ000005 | 1,000 |
| CLUPTJ000006 | 1,000 |
| CLUPTJ000007 | 1,000 |
| CNCS000003 | 0,000 |
| CNCS000004 | 0,000 |
| COLL000007 | 0,000 |
| COLL000013 | 0,000 |
| COOK000002 | 0,000 |
| COOK000007 | 0,000 |
| CZEC000003 | 0,306 |
| CZWO000001 | 0,306 |
| DALM000002 | 0,000 |
| DEER000003 | 0,000 |
| DEER000004 | 0,000 |
| DTR001 | 0,039 |
| DTR002 | 0,031 |
| DTR003 | 0,035 |
| Dalmatian01 | 0,000 |
| EFXH000002 | 0,000 |
| EFXH000003 | 0,000 |
| ENTB000001 | 0,000 |
| ENTB000002 | 0,000 |
| ESMD000002 | 0,000 |
| ESMD000003 | 0,000 |
| FLCR000002 | 0,000 |
| FlatcoatedRetriever03 | 0,000 |
| GALG000002 | 0,000 |
| GALG000003 | 0,000 |

dogs. Values range from 0 (entirely assigned to the domestic dog cluster) to 1 (entirely assigned to the gray wolf cluster). Samples with Q_wolf_ < 0.95 were co

| Sample | Qwolf |
|---|---|
| GORD000002 | 0,000 |
| GPYR000003 | 0,000 |
| GPYR000004 | 0,000 |
| GRSD000002 | 0,000 |
| GRSD000003 | 0,000 |
| GSMD000004 | 0,000 |
| GreaterSwissMountainDog01 | 0,000 |
| HOVA000003 | 0,000 |
| HOVA000004 | 0,000 |
| HWHP000001 | 0,000 |
| IBIZ000010 | 0,000 |
| IBIZ000011 | 0,000 |
| IRSE000001 | 0,000 |
| IRSE000003 | 0,000 |
| IWSP000001 | 0,000 |
| IWSP000002 | 0,000 |
| Iran_wolf | 1,000 |
| KANG000001 | 0,040 |
| KANG000003 | 0,038 |
| KARS000005 | 0,034 |
| KARS000006 | 0,032 |
| KUVZ000002 | 0,000 |
| KUVZ000007 | 0,000 |
| LAGO000001 | 0,000 |
| LAGO000004 | 0,000 |
| LEOP000003 | 0,000 |
| LEOP000008 | 0,000 |
| LUPWCHN00003 | 0,926 |
| LUPWCHN00010 | 1,000 |
| LUPZCHN00006 | 0,941 |
| LUPZCHN00009 | 1,000 |
| LUPZCHN00013 | 0,972 |
| MARM000006 | 0,000 |

dogs. Values range from 0 (entirely assigned to the domestic dog cluster) to 1 (entirely assigned to the gray wolf cluster). Samples with Q_wolf_ < 0.95 were co

| Sample | Qwolf |
|---|---|
| Maremma01 | 0,000 |
| MastinoAbruzzese01 | 0,000 |
| Mongolian_wolf | 0,963 |
| NEAP000004 | 0,000 |
| NEAP000005 | 0,000 |
| NELK000002 | 0,000 |
| NEWF000002 | 0,000 |
| NEWF000006 | 0,000 |
| NorwegianElkhound02 | 0,000 |
| PITB000001 | 0,000 |
| PITB000002 | 0,000 |
| POPT000001 | 0,000 |
| POSD000005 | 0,000 |
| PPOD000004 | 0,000 |
| PPOP000001 | 0,000 |
| PT61 | 0,000 |
| PTWD000008 | 0,000 |
| PYMF000005 | 0,000 |
| PYMF000006 | 0,000 |
| PortugueseWaterDog09 | 0,000 |
| RHOD000001 | 0,000 |
| RKW13451 | 0,955 |
| SAAR000002 | 0,335 |
| SAAR000005 | 0,307 |
| SALU000001 | 0,051 |
| SALU000002 | 0,051 |
| SAMO000002 | 0,052 |
| SGGO000002 | 0,000 |
| SGGO000004 | 0,000 |
| SLOU000001 | 0,000 |
| SLOU000003 | 0,000 |
| SPIN000001 | 0,000 |
| SPIN000004 | 0,000 |

dogs. Values range from 0 (entirely assigned to the domestic dog cluster) to 1 (entirely assigned to the gray wolf cluster). Samples with Q_wolf_ < 0.95 were co

| **Sample** | **Qwolf** |
| --- | --- |
| SPMT000002 | 0,000 |
| SPMT000005 | 0,000 |
| SPWD000002 | 0,000 |
| SPWD000003 | 0,000 |
| SRAB000001 | 0,038 |
| SRAB000002 | 0,045 |
| Samoyed01 | 0,044 |
| TERV000005 | 0,000 |
| TOSA000002 | 0,033 |
| TOSA000003 | 0,027 |
| TURM000002 | 0,035 |
| TURM000004 | 0,037 |
| TURV000001 | 0,000 |
| TWCH000001 | 0,000 |
| TWCH000002 | 0,000 |
| VILLAF000001 | 0,053 |
| VILLAF000002 | 0,077 |
| VILLAF000003 | 0,059 |
| VILLAZ000002 | 0,047 |
| VILLAZ000003 | 0,038 |
| VILLAZ000004 | 0,039 |
| VILLAZ000005 | 0,039 |
| VILLAZ000007 | 0,033 |
| VILLBG000001 | 0,000 |
| VILLBG000002 | 0,000 |
| VILLCG000001 | 0,000 |
| VILLCG000004 | 0,000 |
| VILLCG000006 | 0,000 |
| VILLCG000011 | 0,000 |
| VILLCG000013 | 0,000 |
| VILLCN000005 | 0,230 |
| VILLCN000013 | 0,222 |
| VILLCN000088 | 0,184 |

dogs. Values range from 0 (entirely assigned to the domestic dog cluster) to 1 (entirely assigned to the gray wolf cluster). Samples with Q_wolf_ < 0.95 were co

| Sample | Qwolf |
|---|---|
| VILLCN000145 | 0,162 |
| VILLCN000153 | 0,105 |
| VILLIR000021 | 0,042 |
| VILLIR000023 | 0,000 |
| VILLIR000024 | 0,056 |
| VILLIR000026 | 0,046 |
| VILLIR000027 | 0,081 |
| VILLKE000002 | 0,034 |
| VILLKE000009 | 0,028 |
| VILLKE000010 | 0,046 |
| VILLKE000011 | 0,045 |
| VILLKE000019 | 0,046 |
| VILLLR000005 | 0,026 |
| VILLLR000011 | 0,035 |
| VILLLR000014 | 0,046 |
| VILLLR000016 | 0,009 |
| VILLLR000017 | 0,017 |
| VILLTJ000001 | 0,063 |
| VILLUZ000002 | 0,025 |
| VILLUZ000003 | 0,038 |
| VILLUZ000004 | 0,039 |
| VILLUZ000005 | 0,039 |
| VILLUZ000006 | 0,034 |
| VIZS000007 | 0,000 |
| WEIM000002 | 0,000 |
| WEIM000005 | 0,000 |
| WHPG000001 | 0,000 |
| WHPG000002 | 0,000 |
| WTR025 | 1,000 |
| WTR026 | 1,000 |
| WTR027 | 1,000 |
| WTR028 | 1,000 |
| WTR029 | 1,000 |

dogs. Values range from 0 (entirely assigned to the domestic dog cluster) to 1 (entirely assigned to the gray wolf cluster). Samples with Q_wolf_ < 0.95 were co

| **Sample** | **Qwolf** |
|---|---|
| WTR030 | 1,000 |
| WTR031 | 1,000 |
| WTR032 | 1,000 |
| WTR033 | 1,000 |
| WTR034 | 1,000 |
| WTR035 | 1,000 |
| WTR037 | 0,990 |
| WTR038 | 1,000 |
| WTR039 | 1,000 |
| WTR040 | 1,000 |
| WTR041 | 1,000 |
| ZERD000002 | 0,020 |
| ZERD000003 | 0,019 |
| inw | 1,000 |
| irw | 1,000 |

onsidered potentially admixed and were excluded from subsequent analyses.

onsidered potentially admixed and were excluded from subsequent analyses.

onsidered potentially admixed and were excluded from subsequent analyses.

onsidered potentially admixed and were excluded from subsequent analyses.

onsidered potentially admixed and were excluded from subsequent analyses.

onsidered potentially admixed and were excluded from subsequent analyses.

onsidered potentially admixed and were excluded from subsequent analyses.

onsidered potentially admixed and were excluded from subsequent analyses.

**Table S5.** Ancestry proportion assigned to the gray wolf cluster (Q_wolf_) estimated u Table S5. ADMIXTURE results for K=2 for North American w

| **Number** | **Species** | **Population/Breed** |
|---|---|---|
| 1 | *Canis lupus familiaris* | Golden Retriever |
| 2 | *Canis lupus familiaris* | Labrador Retriever |
| 3 | *Canis lupus familiaris* | Rottweiler |
| 4 | *Canis lupus familiaris* | Akbash |
| 5 | *Canis lupus familiaris* | Alaskan Malamute |
| 6 | *Canis lupus familiaris* | American Staffordshire Terrier |
| 7 | *Canis lupus familiaris* | Anatolian Shepherd Dog |
| 8 | *Canis lupus familiaris* | Australian Shepherd |
| 9 | *Canis lupus* | North America |
| 10 | *Canis lupus* | North America |
| 11 | *Canis lupus* | North America |
| 12 | *Canis lupus* | North America |
| 13 | *Canis lupus familiaris* | Beauceron |
| 14 | *Canis lupus familiaris* | Belgian Sheepdog |
| 15 | *Canis lupus familiaris* | Bloodhound |
| 16 | *Canis lupus familiaris* | Belgian Malinois |
| 17 | *Canis lupus familiaris* | Bavarian Mountain Scent Hound |
| 18 | *Canis lupus familiaris* | Boerboel |
| 19 | *Canis lupus familiaris* | Boxer |
| 20 | *Canis lupus familiaris* | Bracco Italiano |
| 21 | *Canis lupus familiaris* | Briard |
| 22 | *Canis lupus familiaris* | Bernese Mountain Dog |
| 23 | *Canis lupus familiaris* | Black Russian Terrier |
| 24 | *Canis lupus familiaris* | Caucasian Shepherd Dog |
| 25 | *Canis lupus familiaris* | Curly-coated Retriever |
| 26 | *Canis lupus familiaris* | Chesapeake Bay Retriever |
| 27 | *Canis lupus familiaris* | Cane Corso |
| 28 | *Canis lupus familiaris* | Collie |
| 29 | *Canis lupus familiaris* | Chinook |
| 30 | *Canis lupus familiaris* | Dalmatian |
| 31 | *Canis lupus familiaris* | Scottish Deerhound |
| 32 | *Canis lupus familiaris* | English Foxhound |
| 33 | *Canis lupus familiaris* | Entlebucher Mountain Dog |

**Table S5.** Ancestry proportion assigned to the gray wolf cluster (Q_wolf_) estimated u Table S5. ADMIXTURE results for K=2 for North American w

| **Number** | **Species** | **Population/Breed** |
|---|---|---|
| 34 | *Canis lupus familiaris* | Estrela Mountain Dog |
| 35 | *Canis lupus familiaris* | Flat-Coated Retriever |
| 36 | *Canis lupus familiaris* | Galgo Espanol |
| 37 | *Canis lupus familiaris* | Gordon Setter |
| 38 | *Canis lupus familiaris* | Great Pyrenees |
| 39 | *Canis lupus familiaris* | German Shepherd |
| 40 | *Canis lupus familiaris* | German Shepherd |
| 41 | *Canis lupus* | North America |
| 42 | *Canis lupus* | North America |
| 43 | *Canis lupus* | North America |
| 44 | *Canis lupus* | North America |
| 45 | *Canis lupus* | North America |
| 46 | *Canis lupus* | North America |
| 47 | *Canis lupus* | North America |
| 48 | *Canis lupus* | North America |
| 49 | *Canis lupus* | North America |
| 50 | *Canis lupus* | North America |
| 51 | *Canis lupus* | North America |
| 52 | *Canis lupus familiaris* | Hovawart |
| 53 | *Canis lupus familiaris* | Vizsla |
| 54 | *Canis lupus familiaris* | Ibizan Hound |
| 55 | *Canis lupus familiaris* | Irish Setter |
| 56 | *Canis lupus familiaris* | Irish Water Spaniel |
| 57 | *Canis lupus* | North America |
| 58 | *Canis lupus* | North America |
| 59 | *Canis lupus* | North America |
| 60 | *Canis lupus familiaris* | Kuvasz |
| 61 | *Canis lupus familiaris* | Lagotto Romagnolo |
| 62 | *Canis lupus familiaris* | Catahoula Leopard Dog |
| 63 | *Canis lupus familiaris* | Maremma Sheepdog |
| 64 | *Canis lupus familiaris* | Mastino Abruzzese |
| 65 | *Canis lupus* | North America |
| 66 | *Canis lupus* | North America |

**Table S5.** Ancestry proportion assigned to the gray wolf cluster (Q_wolf_) estimated u Table S5. ADMIXTURE results for K=2 for North American w

| **Number** | **Species** | **Population/Breed** |
|---|---|---|
| 67 | *Canis lupus* | North America |
| 68 | *Canis lupus* | North America |
| 69 | *Canis lupus* | North America |
| 70 | *Canis lupus* | North America |
| 71 | *Canis lupus* | North America |
| 72 | *Canis lupus familiaris* | Neapolitan Mastiff |
| 73 | *Canis lupus familiaris* | Norwegian Elkhound |
| 74 | *Canis lupus familiaris* | Newfoundland |
| 75 | *Canis lupus familiaris* | Pit Bull Terrier |
| 76 | *Canis lupus familiaris* | Portuguese Pointer |
| 77 | *Canis lupus familiaris* | Portuguese Sheepdog |
| 78 | *Canis lupus familiaris* | Portuguese Podengo |
| 79 | *Canis lupus familiaris* | Pyrenean Mastiff |
| 80 | *Canis lupus familiaris* | Portuguese Water Dog |
| 81 | *Canis lupus* | North America |
| 82 | *Canis lupus familiaris* | Rhodesian Ridgeback |
| 83 | *Canis lupus familiaris* | Saluki |
| 84 | *Canis lupus familiaris* | Samoyed |
| 85 | *Canis lupus familiaris* | Segugio Italiano |
| 86 | *Canis lupus familiaris* | Sloughi |
| 87 | *Canis lupus familiaris* | Spinone Italiano |
| 88 | *Canis lupus familiaris* | Spanish Mastiff |
| 89 | *Canis lupus familiaris* | Spanish Water Dog |
| 90 | *Canis lupus familiaris* | Belgian Tervuren |
| 91 | *Canis lupus familiaris* | Tosa Inu |
| 92 | *Canis lupus familiaris* | Treeing Walker Coonhound |
| 93 | *Canis lupus familiaris* | Weimaraner |
| 94 | *Canis lupus familiaris* | Wirehaired Pointing Griffon |
| 95 | *Canis lupus* | North America |
| 96 | *Canis lupus* | North America |
| 97 | *Canis lupus* | North America |
| 98 | *Canis lupus* | North America |
| 99 | *Canis lupus* | North America |

**Table S5.** Ancestry proportion assigned to the gray wolf cluster (Q_wolf_) estimated u Table S5. ADMIXTURE results for K=2 for North American w

| **Number** | **Species** | **Population/Breed** |
|---|---|---|
| 100 | *Canis lupus* | North America |
| 101 | *Canis lupus* | North America |

olves and dogs

| Sample | Qwolf |
|---|---|
| 140447_S11 | 0,000 |
| 149323_S6 | 0,000 |
| 165414_S20 | 0,000 |
| AKBH000001 | 0,030 |
| AMAL000002 | 0,130 |
| AMST000006 | 0,000 |
| ANAT000003 | 0,040 |
| AUSS000001 | 0,000 |
| Arctic_BaffinIsl_CD130_RKW7639 | 1,000 |
| Arctic_EllesmereIsl_GF44_RKW7640 | 1,000 |
| Arctic_Nunavut_CB177_RKW7649 | 1,000 |
| Arctic_VictoriaIsl_CB215_RKW7619 | 1,000 |
| BEAU000007 | 0,000 |
| BELS000003 | 0,000 |
| BLDH000002 | 0,000 |
| BMAL000001 | 0,000 |
| BMSH000005 | 0,000 |
| BOER000002 | 0,000 |
| BOXR000001 | 0,000 |
| BRAC000001 | 0,000 |
| BRIA000005 | 0,000 |
| BRMD000001 | 0,000 |
| BRTR000002 | 0,000 |
| CAUC000001 | 0,020 |
| CCRT000004 | 0,000 |
| CHBR000001 | 0,000 |
| CNCS000003 | 0,000 |
| COLL000007 | 0,000 |
| COOK000002 | 0,000 |
| DALM000002 | 0,000 |
| DEER000003 | 0,000 |
| EFXH000003 | 0,000 |
| ENTB000002 | 0,000 |

olves and dogs

| Sample | Qwolf |
| --- | --- |
| ESMD000002 | 0,000 |
| FLCR000002 | 0,000 |
| GALG000002 | 0,000 |
| GORD000002 | 0,000 |
| GPYR000003 | 0,000 |
| GRSD000002 | 0,000 |
| GRSD000003 | 0,000 |
| GreyWolf_AtlanticCoast | 1,000 |
| GreyWolf_BaffinNorth | 1,000 |
| GreyWolf_BaffinSouth | 1,000 |
| GreyWolf_BanksIsland | 1,000 |
| GreyWolf_EllesmereIsland | 1,000 |
| GreyWolf_Greenland | 1,000 |
| GreyWolf_Montana | 1,000 |
| GreyWolf_PacificCoast | 1,000 |
| GreyWolf_StLawrenceIsland | 1,000 |
| GreyWolf_Toronto | 1,000 |
| GreyWolf_VictoriaIsland | 1,000 |
| HOVA000003 | 0,000 |
| HWHP000001 | 0,000 |
| IBIZ000010 | 0,000 |
| IRSE000001 | 0,000 |
| IWSP000001 | 0,000 |
| IsleRoyaleNP_CL141_JRRW018 | 1,000 |
| IsleRoyaleNP_CL189_RWJR008 | 1,000 |
| IsleRoyaleNP_CL61_RWJR005 | 1,000 |
| KUVZ000002 | 0,000 |
| LAGO000001 | 0,000 |
| LEOP000003 | 0,000 |
| MARM000006 | 0,000 |
| MastinoAbruzzese01 | 0,000 |
| Mexican_wolf | 1,000 |
| Minnesota_RKW119_RWJR007 | 1,000 |

olves and dogs

| Sample | Qwolf |
|---|---|
| Minnesota_RKW2515_RWJR016 | 1,000 |
| Minnesota_RKW2518_RWJR012 | 1,000 |
| Minnesota_RKW2523_RWJR009 | 1,000 |
| Minnesota_RKW2524_RWJR003 | 1,000 |
| Minnesota_wolf | 1,000 |
| NEAP000004 | 0,000 |
| NELK000002 | 0,000 |
| NEWF000002 | 0,000 |
| PITB000001 | 0,000 |
| POPT000001 | 0,000 |
| POSD000005 | 0,000 |
| PPOD000004 | 0,000 |
| PYMF000005 | 0,000 |
| PortugueseWaterDog09 | 0,000 |
| Quebec_MontTremblantNP_voyou0833M_RWBH001 | 1,000 |
| RHOD000001 | 0,000 |
| SALU000001 | 0,050 |
| SAMO000002 | 0,050 |
| SGGO000002 | 0,000 |
| SLOU000001 | 0,000 |
| SPIN000001 | 0,000 |
| SPMT000005 | 0,000 |
| SPWD000003 | 0,000 |
| TERV000005 | 0,000 |
| TOSA000002 | 0,030 |
| TWCH000001 | 0,000 |
| WEIM000002 | 0,000 |
| WHPG000001 | 0,000 |
| Yellowstone1_wolf | 1,000 |
| Yellowstone2_wolf | 1,000 |
| Yellowstone3_wolf | 1,000 |
| glw | 1,000 |
| mxa | 1,000 |

olves and dogs

| Sample | Qwolf |
| --- | --- |
| ysa | 1,000 |
| ysb | 1,000 |

**Table S6.** Results of kinship analyses for those pairs displaying indexes values beyond the thresholds for first-grade relatives.

| Continent | Population | Individual A | Individual B | R0 | R1 | KING |
|---|---|---|---|---|---|---|
| Europe | Dinaric-Balkan | AP_085T | AP_08C5 | 0,062576 | 0,753192 | 0,267938 |
| Europe | Dinaric-Balkan | AP_086E | AP_086F | 0,073342 | 0,744412 | 0,260953 |
| Europe | Karelian | CLUPRU000006 | CLUPRU000007 | 0,075551 | 0,634166 | 0,242451 |
| Europe | Karelian | W26 | W29 | 0,105713 | 0,613502 | 0,223755 |
| Europe | NW Iberia | JAL7480 | L844 | 0,000026 | 71,050371 | 0,496487 |
| Europe | Scandinavian | G100-12 | G109-11 | 0,032372 | 0,805319 | 0,291414 |
| Europe | Scandinavian | G100-14 | G31-13 | 0,000303 | 0,69556 | 0,290742 |
| Europe | Scandinavian | G37-10 | M-10-04 | 0,062951 | 0,833703 | 0,278684 |
| Asia | Central Asia | BGI-01505070004 | Mongolian_wolf | 0,000008 | 125,992551 | 0,498018 |
| Asia | East Asia | BGI-01505070003 | Qinghai_wolf | 0,000014 | 91,2259 | 0,497264 |
| Asia | East Asia | CLUPCN000004 | CLUPCN000005 | 0,073407 | 0,781758 | 0,266141 |
| Asia | Tibetan | CLUPCN000002 | CLUPCN000003 | 0,059626 | 0,880555 | 0,286327 |
| Asia | Turkey | WTR027 | WTR028 | 0,000005 | 1,2812 | 0,359642 |
| Asia | Turkey | WTR025 | WTR035 | 0,05486 | 0,703514 | 0,264447 |
| Asia | Turkey | WTR039 | WTR041 | 0,095105 | 0,698897 | 0,242763 |
| Asia | Turkey | WTR027 | WTR041 | 0,055634 | 0,482126 | 0,221159 |
| Asia | Turkey | WTR027 | WTR031 | 0,053615 | 0,473462 | 0,219976 |
| Asia | Turkey | WTR027 | WTR039 | 0,061397 | 0,4591 | 0,213081 |
| Asia | Turkey | WTR031 | WTR039 | 0,109216 | 0,517055 | 0,204342 |
| Asia | West Asia | CLUPIR000002 | CLUPIR000004 | 0,071881 | 0,687275 | 0,25309 |
| Asia | West Asia | CLUPIR000005 | CLUPIR000006 | 0,088863 | 0,666141 | 0,240972 |
| Asia | West Asia | Iran_wolf | irw | 0,000009 | 154,361535 | 0,498379 |
| North America | North America | Yellowstone2_wolf | ysa | 0,000014 | 126,766712 | 0,498025 |
| North America | North America | Minnesota_wolf | glw | 0,000014 | 87,805783 | 0,497158 |
| North America | North America | Yellowstone3_wolf | ysb | 0,000009 | 83,628997 | 0,497022 |
| North America | North America | Mexican_wolf | mxa | 0,000044 | 42,894663 | 0,494207 |
| North America | North America | Yellowstone2_wolf | Yellowstone3_wolf | 0,000275 | 0,512881 | 0,253058 |
| North America | North America | Yellowstone3_wolf | ysa | 0,000474 | 0,511367 | 0,2526 |
| North America | North America | Yellowstone2_wolf | ysb | 0,000478 | 0,50893 | 0,252002 |

**Table S7.** Final list of wolf samples used for the analyses and their population.

| Continent | Population | VCF_ID |
|---|---|---|
| Asia | Central Asia | CLUPKG000001 |
| Asia | Central Asia | CLUPKZ000002 |
| Asia | Central Asia | CLUPRU000001 |
| Asia | Central Asia | CLUPRU000002 |
| Asia | Central Asia | CLUPRU000003 |
| Asia | Central Asia | CLUPTJ000001 |
| Asia | Central Asia | CLUPTJ000002 |
| Asia | Central Asia | CLUPTJ000003 |
| Asia | Central Asia | CLUPTJ000004 |
| Asia | Central Asia | CLUPTJ000005 |
| Asia | Central Asia | CLUPTJ000006 |
| Asia | Central Asia | CLUPTJ000007 |
| Asia | East Asia | BGI-01505070006 |
| Asia | East Asia | BGI-01505070007 |
| Asia | East Asia | CLUPCN000001 |
| Asia | East Asia | CLUPCN000005 |
| Asia | East Asia | CLUPCN000006 |
| Asia | East Asia | CLUPCN000007 |
| Asia | East Asia | CLUPCN000008 |
| Asia | East Asia | CLUPCN000009 |
| Asia | East Asia | CLUPCN000010 |
| Asia | East Asia | CLUPRU000004 |
| Asia | East Asia | LUPWCHN00010 |
| Asia | East Asia | LUPZCHN00009 |
| Asia | East Asia | RKW13451 |
| Asia | Indian | inw |
| Asia | Tibetan | BGI-01505070001 |
| Asia | Tibetan | BGI-01505070002 |
| Asia | Tibetan | BGI-01505070003 |
| Asia | Tibetan | BGI-01505070004 |
| Asia | Tibetan | BGI-01505070005 |
| Asia | Tibetan | BGI-01505070008 |
| Asia | Tibetan | CLUPCN000003 |

**Table S7.** Final list of wolf samples used for the analyses and their population.

| | | |
|---|---|---|
| Asia | Turkey | WTR026 |
| Asia | Turkey | WTR028 |
| Asia | Turkey | WTR029 |
| Asia | Turkey | WTR030 |
| Asia | Turkey | WTR031 |
| Asia | Turkey | WTR032 |
| Asia | Turkey | WTR033 |
| Asia | Turkey | WTR034 |
| Asia | Turkey | WTR035 |
| Asia | Turkey | WTR037 |
| Asia | Turkey | WTR038 |
| Asia | Turkey | WTR040 |
| Asia | Turkey | WTR041 |
| Asia | West Asia | CLUPAZ000001 |
| Asia | West Asia | CLUPIR000001 |
| Asia | West Asia | CLUPIR000003 |
| Asia | West Asia | CLUPIR000004 |
| Asia | West Asia | CLUPIR000005 |
| Asia | West Asia | irw |
| Europe | Dinaric-Balkan | AP_085T |
| Europe | Dinaric-Balkan | AP_086E |
| Europe | Dinaric-Balkan | AP_087A |
| Europe | Dinaric-Balkan | AP_087E |
| Europe | Dinaric-Balkan | AP_087L |
| Europe | Dinaric-Balkan | AP_087U |
| Europe | Dinaric-Balkan | AP_0882 |
| Europe | Dinaric-Balkan | AP_088L |
| Europe | Dinaric-Balkan | AP_08A1 |
| Europe | Dinaric-Balkan | AP_08A3 |
| Europe | Dinaric-Balkan | AP_08A6 |
| Europe | Dinaric-Balkan | AP_08C3 |
| Europe | Dinaric-Balkan | BiH_244 |
| Europe | Dinaric-Balkan | BiH_245 |
| Europe | Dinaric-Balkan | LUP006817 |

**Table S7.** Final list of wolf samples used for the analyses and their population.

| | | |
|---|---|---|
| Europe | Dinaric-Balkan | LUP006818 |
| Europe | Dinaric-Balkan | LUP006819 |
| Europe | Dinaric-Balkan | LUP006820 |
| Europe | Dinaric-Balkan | LUP006821 |
| Europe | Dinaric-Balkan | LUP006822 |
| Europe | Dinaric-Balkan | LUP006823 |
| Europe | Dinaric-Balkan | LUP006824 |
| Europe | Dinaric-Balkan | LUP006825 |
| Europe | Dinaric-Balkan | LUP006826 |
| Europe | Dinaric-Balkan | LUP006827 |
| Europe | Dinaric-Balkan | LUP006828 |
| Europe | Dinaric-Balkan | MSV2AC |
| Europe | Italian Peninsula | W1023 |
| Europe | Italian Peninsula | W1551 |
| Europe | Italian Peninsula | W1593 |
| Europe | Italian Peninsula | W2030 |
| Europe | Italian Peninsula | W2124 |
| Europe | Italian Peninsula | W2185 |
| Europe | Italian Peninsula | W2193 |
| Europe | Italian Peninsula | W2265 |
| Europe | Italian Peninsula | W2280 |
| Europe | Italian Peninsula | W2291 |
| Europe | Italian Peninsula | W2371F |
| Europe | Italian Peninsula | W2430F |
| Europe | Italian Peninsula | W2487F |
| Europe | Italian Peninsula | W2594M |
| Europe | Italian Peninsula | W2623M |
| Europe | Italian Peninsula | W2707M |
| Europe | Italian Peninsula | W2878M |
| Europe | Italian Peninsula | W2883F |
| Europe | Italian Peninsula | W2886 |
| Europe | Italian Peninsula | W2910 |
| Europe | Italian Peninsula | W2951 |
| Europe | Italian Peninsula | w361703 |

**Table S7.** Final list of wolf samples used for the analyses and their population.

| | | |
|---|---|---|
| Europe | Italian Peninsula | w361756 |
| Europe | Italian Peninsula | w4926402 |
| Europe | Italian Peninsula | W893 |
| Europe | Italian Peninsula | WIT11 |
| Europe | Italian Peninsula | WIT13 |
| Europe | Italian Peninsula | WIT14 |
| Europe | Italian Peninsula | WIT16 |
| Europe | Italian Peninsula | WIT17 |
| Europe | Italian Peninsula | WIT19 |
| Europe | Italian Peninsula | WIT20 |
| Europe | Italian Peninsula | WIT21 |
| Europe | Italian Peninsula | WIT22 |
| Europe | Italian Peninsula | WIT23 |
| Europe | Karelian | CLUPRU000005 |
| Europe | Karelian | CLUPRU000007 |
| Europe | Karelian | CLUPRU000008 |
| Europe | Karelian | CLUPRU000019 |
| Europe | Karelian | CLUPRU000020 |
| Europe | Karelian | V113 |
| Europe | Karelian | V114 |
| Europe | Karelian | V115 |
| Europe | Karelian | V116 |
| Europe | Karelian | V117 |
| Europe | Karelian | V119 |
| Europe | Karelian | V120 |
| Europe | Karelian | V126 |
| Europe | Karelian | V132 |
| Europe | Karelian | V134 |
| Europe | Karelian | V136 |
| Europe | Karelian | V141 |
| Europe | Karelian | V143 |
| Europe | Karelian | W11 |
| Europe | Karelian | W14 |
| Europe | Karelian | W16 |

**Table S7.** Final list of wolf samples used for the analyses and their population.

| | | |
|---|---|---|
| Europe | Karelian | W21 |
| Europe | Karelian | W26 |
| Europe | Karelian | W32 |
| Europe | Karelian | W4 |
| Europe | Karelian | W44 |
| Europe | Karelian | W45 |
| Europe | NW Iberia | JAL7480 |
| Europe | NW Iberia | JAL7486 |
| Europe | NW Iberia | JAL7491 |
| Europe | NW Iberia | JAL7492 |
| Europe | NW Iberia | L166 |
| Europe | NW Iberia | L253 |
| Europe | NW Iberia | L409 |
| Europe | NW Iberia | L514 |
| Europe | NW Iberia | L547 |
| Europe | NW Iberia | L552 |
| Europe | NW Iberia | L588 |
| Europe | NW Iberia | LUP006829 |
| Europe | NW Iberia | LUP006830 |
| Europe | NW Iberia | MW122 |
| Europe | NW Iberia | MW127 |
| Europe | NW Iberia | Penelope |
| Europe | NW Iberia | ptw |
| Europe | Scandinavian | D-00-12 |
| Europe | Scandinavian | D-06-14 |
| Europe | Scandinavian | D-07-16 |
| Europe | Scandinavian | D-08-10 |
| Europe | Scandinavian | D-10-68 |
| Europe | Scandinavian | D-11-58 |
| Europe | Scandinavian | D-85-02 |
| Europe | Scandinavian | G100-12 |
| Europe | Scandinavian | G100-14 |
| Europe | Scandinavian | G126-13 |
| Europe | Scandinavian | G139-12 |

**Table S7.** Final list of wolf samples used for the analyses and their population.

| | | |
|---|---|---|
| Europe | Scandinavian | G50-12 |
| Europe | Scandinavian | M-01-06 |
| Europe | Scandinavian | M-06-03 |
| Europe | Scandinavian | M-09-05 |
| Europe | Scandinavian | M-10-04 |
| Europe | Scandinavian | M-98-08 |
| North America | North America | Arctic_BaffinIsl_CD130_RKW7639 |
| North America | North America | Arctic_EllesmereIsl_GF44_RKW7640 |
| North America | North America | Arctic_Nunavut_CB177_RKW7649 |
| North America | North America | Arctic_VictoriaIsl_CB215_RKW7619 |
| North America | North America | glw |
| North America | North America | GreyWolf_AtlanticCoast |
| North America | North America | GreyWolf_BaffinNorth |
| North America | North America | GreyWolf_BaffinSouth |
| North America | North America | GreyWolf_BanksIsland |
| North America | North America | GreyWolf_Greenland |
| North America | North America | GreyWolf_Montana |
| North America | North America | GreyWolf_PacificCoast |
| North America | North America | GreyWolf_StLawrenceIsland |
| North America | North America | GreyWolf_Toronto |
| North America | North America | GreyWolf_VictoriaIsland |
| North America | North America | IsleRoyaleNP_CL141_JRRW018 |
| North America | North America | IsleRoyaleNP_CL189_RWJR008 |
| North America | North America | IsleRoyaleNP_CL61_RWJR005 |
| North America | North America | Minnesota_RKW119_RWJR007 |
| North America | North America | Minnesota_RKW2515_RWJR016 |
| North America | North America | Minnesota_RKW2518_RWJR012 |
| North America | North America | Minnesota_RKW2523_RWJR009 |
| North America | North America | Minnesota_RKW2524_RWJR003 |
| North America | North America | mxa |
| North America | North America | Quebec_MontTremblantNP_voyou0833M_RWBH001 |
| North America | North America | ysa |
| North America | North America | ysb |

**Table S8.** ADMIXTURE results for K = 6 for European, Asian and North American wolves.

| Number | Continent | Population | Sample | | | Admixture_k6 | |
|---|---|---|---|---|---|---|---|
| 1 | Europe | Dinaric-Balkan | AP_085T | 1,000 | 0,000 | 0,000 | 0,000 |
| 2 | Europe | Dinaric-Balkan | AP_086E | 1,000 | 0,000 | 0,000 | 0,000 |
| 3 | Europe | Dinaric-Balkan | AP_087A | 1,000 | 0,000 | 0,000 | 0,000 |
| 4 | Europe | Dinaric-Balkan | AP_087E | 1,000 | 0,000 | 0,000 | 0,000 |
| 5 | Europe | Dinaric-Balkan | AP_087L | 1,000 | 0,000 | 0,000 | 0,000 |
| 6 | Europe | Dinaric-Balkan | AP_087U | 1,000 | 0,000 | 0,000 | 0,000 |
| 7 | Europe | Dinaric-Balkan | AP_0882 | 1,000 | 0,000 | 0,000 | 0,000 |
| 8 | Europe | Dinaric-Balkan | AP_088L | 1,000 | 0,000 | 0,000 | 0,000 |
| 9 | Europe | Dinaric-Balkan | AP_08A1 | 1,000 | 0,000 | 0,000 | 0,000 |
| 10 | Europe | Dinaric-Balkan | AP_08A3 | 1,000 | 0,000 | 0,000 | 0,000 |
| 11 | Europe | Dinaric-Balkan | AP_08A6 | 1,000 | 0,000 | 0,000 | 0,000 |
| 12 | Europe | Dinaric-Balkan | AP_08C3 | 1,000 | 0,000 | 0,000 | 0,000 |
| 13 | North America | North America | Arctic_BaffinIsl_CD130_RKW7639 | 0,000 | 0,000 | 1,000 | 0,000 |
| 14 | North America | North America | Arctic_EllesmereIsl_GF44_RKW7640 | 0,000 | 0,000 | 1,000 | 0,000 |
| 15 | North America | North America | Arctic_Nunavut_CB177_RKW7649 | 0,000 | 0,000 | 1,000 | 0,000 |
| 16 | North America | North America | Arctic_VictoriaIsl_CB215_RKW7619 | 0,000 | 0,000 | 1,000 | 0,000 |
| 17 | Asia | Tibetan | BGI-01505070001 | 0,000 | 0,000 | 0,000 | 0,000 |
| 18 | Asia | Tibetan | BGI-01505070002 | 0,000 | 0,000 | 0,000 | 0,000 |
| 19 | Asia | Tibetan | BGI-01505070003 | 0,000 | 0,000 | 0,000 | 0,000 |
| 20 | Asia | Tibetan | BGI-01505070004 | 0,000 | 0,000 | 0,167 | 0,009 |
| 21 | Asia | Tibetan | BGI-01505070005 | 0,000 | 0,000 | 0,154 | 0,007 |
| 22 | Asia | East Asia | BGI-01505070006 | 0,085 | 0,000 | 0,086 | 0,036 |
| 23 | Asia | East Asia | BGI-01505070007 | 0,056 | 0,000 | 0,082 | 0,031 |
| 24 | Asia | Tibetan | BGI-01505070008 | 0,000 | 0,000 | 0,000 | 0,000 |
| 25 | Europe | Dinaric-Balkan | BiH_244 | 1,000 | 0,000 | 0,000 | 0,000 |
| 26 | Europe | Dinaric-Balkan | BiH_245 | 1,000 | 0,000 | 0,000 | 0,000 |
| 27 | Asia | West Asia | CLUPAZ000001 | 0,095 | 0,000 | 0,000 | 0,000 |
| 28 | Asia | East Asia | CLUPCN000001 | 0,010 | 0,000 | 0,160 | 0,017 |
| 29 | Asia | Tibetan | CLUPCN000003 | 0,000 | 0,000 | 0,000 | 0,000 |
| 30 | Asia | East Asia | CLUPCN000005 | 0,081 | 0,000 | 0,060 | 0,032 |
| 31 | Asia | East Asia | CLUPCN000006 | 0,091 | 0,000 | 0,064 | 0,039 |
| 32 | Asia | East Asia | CLUPCN000007 | 0,087 | 0,000 | 0,068 | 0,034 |
| 33 | Asia | East Asia | CLUPCN000008 | 0,100 | 0,000 | 0,063 | 0,042 |

**Table S8.** ADMIXTURE results for K = 6 for European, Asian and North American wolves.

| Number | Continent | Population | Sample | | | Admixture_k6 | |
|---|---|---|---|---|---|---|---|
| 34 | Asia | East Asia | CLUPCN000009 | 0,096 | 0,000 | 0,060 | 0,039 |
| 35 | Asia | East Asia | CLUPCN000010 | 0,084 | 0,000 | 0,056 | 0,036 |
| 36 | Asia | West Asia | CLUPIR000001 | 0,068 | 0,000 | 0,000 | 0,000 |
| 37 | Asia | West Asia | CLUPIR000003 | 0,077 | 0,000 | 0,000 | 0,000 |
| 38 | Asia | West Asia | CLUPIR000004 | 0,069 | 0,003 | 0,000 | 0,000 |
| 39 | Asia | West Asia | CLUPIR000005 | 0,072 | 0,000 | 0,000 | 0,000 |
| 40 | Asia | Central Asia | CLUPKG000001 | 0,092 | 0,005 | 0,064 | 0,036 |
| 41 | Asia | Central Asia | CLUPKZ000002 | 0,086 | 0,000 | 0,071 | 0,035 |
| 42 | Asia | Central Asia | CLUPRU000001 | 0,090 | 0,000 | 0,103 | 0,046 |
| 43 | Asia | Central Asia | CLUPRU000002 | 0,096 | 0,000 | 0,110 | 0,049 |
| 44 | Asia | Central Asia | CLUPRU000003 | 0,086 | 0,000 | 0,253 | 0,053 |
| 45 | Asia | East Asia | CLUPRU000004 | 0,053 | 0,000 | 0,216 | 0,044 |
| 46 | Europe | Karelian | CLUPRU000005 | 0,375 | 0,009 | 0,000 | 0,336 |
| 47 | Europe | Karelian | CLUPRU000007 | 0,416 | 0,020 | 0,000 | 0,259 |
| 48 | Europe | Karelian | CLUPRU000008 | 0,408 | 0,025 | 0,000 | 0,266 |
| 49 | Europe | Karelian | CLUPRU000019 | 0,361 | 0,011 | 0,000 | 0,350 |
| 50 | Europe | Karelian | CLUPRU000020 | 0,360 | 0,004 | 0,000 | 0,331 |
| 51 | Asia | Central Asia | CLUPTJ000001 | 0,000 | 0,000 | 0,000 | 0,000 |
| 52 | Asia | Central Asia | CLUPTJ000002 | 0,000 | 0,000 | 0,000 | 0,000 |
| 53 | Asia | Central Asia | CLUPTJ000003 | 0,000 | 0,000 | 0,000 | 0,000 |
| 54 | Asia | Central Asia | CLUPTJ000004 | 0,000 | 0,000 | 0,000 | 0,000 |
| 55 | Asia | Central Asia | CLUPTJ000005 | 0,000 | 0,000 | 0,000 | 0,000 |
| 56 | Asia | Central Asia | CLUPTJ000006 | 0,000 | 0,000 | 0,000 | 0,000 |
| 57 | Asia | Central Asia | CLUPTJ000007 | 0,000 | 0,000 | 0,000 | 0,000 |
| 58 | Europe | Scandinavian | D-00-12 | 0,000 | 0,000 | 0,000 | 1,000 |
| 59 | Europe | Scandinavian | D-06-14 | 0,000 | 0,000 | 0,000 | 1,000 |
| 60 | Europe | Scandinavian | D-07-16 | 0,000 | 0,000 | 0,000 | 1,000 |
| 61 | Europe | Scandinavian | D-08-10 | 0,000 | 0,000 | 0,000 | 1,000 |
| 62 | Europe | Scandinavian | D-10-68 | 0,000 | 0,000 | 0,000 | 1,000 |
| 63 | Europe | Scandinavian | D-11-58 | 0,000 | 0,000 | 0,000 | 1,000 |
| 64 | Europe | Scandinavian | D-85-02 | 0,046 | 0,000 | 0,000 | 0,954 |
| 65 | Europe | Scandinavian | G100-12 | 0,004 | 0,000 | 0,000 | 0,996 |
| 66 | Europe | Scandinavian | G100-14 | 0,371 | 0,000 | 0,000 | 0,502 |

**Table S8.** ADMIXTURE results for K = 6 for European, Asian and North American wolves.

| **Number** | **Continent** | **Population** | **Sample** | | | **Admixture_k6** | |
|---|---|---|---|---|---|---|---|
| 67 | Europe | Scandinavian | G126-13 | 0,000 | 0,000 | 0,000 | 1,000 |
| 68 | Europe | Scandinavian | G139-12 | 0,038 | 0,000 | 0,000 | 0,962 |
| 69 | Europe | Scandinavian | G50-12 | 0,000 | 0,000 | 0,000 | 1,000 |
| 70 | North America | North America | GreyWolf_AtlanticCoast | 0,000 | 0,000 | 1,000 | 0,000 |
| 71 | North America | North America | GreyWolf_BaffinNorth | 0,000 | 0,000 | 1,000 | 0,000 |
| 72 | North America | North America | GreyWolf_BaffinSouth | 0,000 | 0,000 | 1,000 | 0,000 |
| 73 | North America | North America | GreyWolf_BanksIsland | 0,000 | 0,000 | 1,000 | 0,000 |
| 74 | North America | North America | GreyWolf_Greenland | 0,000 | 0,000 | 1,000 | 0,000 |
| 75 | North America | North America | GreyWolf_Montana | 0,000 | 0,000 | 1,000 | 0,000 |
| 76 | North America | North America | GreyWolf_PacificCoast | 0,000 | 0,000 | 1,000 | 0,000 |
| 77 | North America | North America | GreyWolf_StLawrenceIsland | 0,000 | 0,000 | 1,000 | 0,000 |
| 78 | North America | North America | GreyWolf_Toronto | 0,000 | 0,000 | 1,000 | 0,000 |
| 79 | North America | North America | GreyWolf_VictoriaIsland | 0,000 | 0,000 | 1,000 | 0,000 |
| 80 | North America | North America | IsleRoyaleNP_CL141_JRRW018 | 0,000 | 0,000 | 1,000 | 0,000 |
| 81 | North America | North America | IsleRoyaleNP_CL189_RWJR008 | 0,000 | 0,000 | 1,000 | 0,000 |
| 82 | North America | North America | IsleRoyaleNP_CL61_RWJR005 | 0,000 | 0,000 | 1,000 | 0,000 |
| 83 | Europe | NW Iberia | JAL7480 | 0,000 | 0,000 | 0,000 | 0,000 |
| 84 | Europe | NW Iberia | JAL7486 | 0,000 | 0,000 | 0,000 | 0,000 |
| 85 | Europe | NW Iberia | JAL7491 | 0,000 | 0,000 | 0,000 | 0,000 |
| 86 | Europe | NW Iberia | JAL7492 | 0,000 | 0,000 | 0,000 | 0,000 |
| 87 | Europe | NW Iberia | L166 | 0,000 | 0,000 | 0,000 | 0,000 |
| 88 | Europe | NW Iberia | L253 | 0,000 | 0,000 | 0,000 | 0,000 |
| 89 | Europe | NW Iberia | L409 | 0,000 | 0,000 | 0,000 | 0,000 |
| 90 | Europe | NW Iberia | L514 | 0,000 | 0,000 | 0,000 | 0,000 |
| 91 | Europe | NW Iberia | L547 | 0,000 | 0,000 | 0,000 | 0,000 |
| 92 | Europe | NW Iberia | L552 | 0,000 | 0,000 | 0,000 | 0,000 |
| 93 | Europe | NW Iberia | L588 | 0,000 | 0,000 | 0,000 | 0,000 |
| 94 | Europe | Dinaric-Balkan | LUP006817 | 1,000 | 0,000 | 0,000 | 0,000 |
| 95 | Europe | Dinaric-Balkan | LUP006818 | 0,842 | 0,012 | 0,000 | 0,029 |
| 96 | Europe | Dinaric-Balkan | LUP006819 | 0,976 | 0,011 | 0,000 | 0,013 |
| 97 | Europe | Dinaric-Balkan | LUP006820 | 0,956 | 0,016 | 0,000 | 0,010 |
| 98 | Europe | Dinaric-Balkan | LUP006821 | 1,000 | 0,000 | 0,000 | 0,000 |
| 99 | Europe | Dinaric-Balkan | LUP006822 | 0,983 | 0,004 | 0,000 | 0,000 |

**Table S8.** ADMIXTURE results for K = 6 for European, Asian and North American wolves.

| Number | Continent | Population | Sample | | | Admixture_k6 | |
|---|---|---|---|---|---|---|---|
| 100 | Europe | Dinaric-Balkan | LUP006823 | 1,000 | 0,000 | 0,000 | 0,000 |
| 101 | Europe | Dinaric-Balkan | LUP006824 | 0,894 | 0,015 | 0,000 | 0,025 |
| 102 | Europe | Dinaric-Balkan | LUP006825 | 1,000 | 0,000 | 0,000 | 0,000 |
| 103 | Europe | Dinaric-Balkan | LUP006826 | 1,000 | 0,000 | 0,000 | 0,000 |
| 104 | Europe | Dinaric-Balkan | LUP006827 | 1,000 | 0,000 | 0,000 | 0,000 |
| 105 | Europe | Dinaric-Balkan | LUP006828 | 1,000 | 0,000 | 0,000 | 0,000 |
| 106 | Europe | NW Iberia | LUP006829 | 0,000 | 0,013 | 0,000 | 0,000 |
| 107 | Europe | NW Iberia | LUP006830 | 0,000 | 0,000 | 0,000 | 0,000 |
| 108 | Asia | East Asia | LUPWCHN00010 | 0,058 | 0,000 | 0,088 | 0,033 |
| 109 | Asia | East Asia | LUPZCHN00009 | 0,063 | 0,000 | 0,087 | 0,037 |
| 110 | Europe | Scandinavian | M-01-06 | 0,000 | 0,000 | 0,000 | 1,000 |
| 111 | Europe | Scandinavian | M-06-03 | 0,000 | 0,000 | 0,000 | 1,000 |
| 112 | Europe | Scandinavian | M-09-05 | 0,000 | 0,000 | 0,000 | 1,000 |
| 113 | Europe | Scandinavian | M-10-04 | 0,106 | 0,000 | 0,000 | 0,894 |
| 114 | Europe | Scandinavian | M-98-08 | 0,000 | 0,000 | 0,000 | 1,000 |
| 115 | Europe | Dinaric-Balkan | MSV2AC | 1,000 | 0,000 | 0,000 | 0,000 |
| 116 | Europe | NW Iberia | MW122 | 0,000 | 0,000 | 0,000 | 0,000 |
| 117 | Europe | NW Iberia | MW127 | 0,000 | 0,000 | 0,000 | 0,000 |
| 118 | North America | North America | Minnesota_RKW119_RWJR007 | 0,000 | 0,000 | 1,000 | 0,000 |
| 119 | North America | North America | Minnesota_RKW2515_RWJR016 | 0,000 | 0,000 | 1,000 | 0,000 |
| 120 | North America | North America | Minnesota_RKW2518_RWJR012 | 0,000 | 0,000 | 1,000 | 0,000 |
| 121 | North America | North America | Minnesota_RKW2523_RWJR009 | 0,000 | 0,000 | 1,000 | 0,000 |
| 122 | North America | North America | Minnesota_RKW2524_RWJR003 | 0,000 | 0,000 | 1,000 | 0,000 |
| 123 | Europe | NW Iberia | Penelope | 0,000 | 0,000 | 0,000 | 0,000 |
| 124 | North America | North America | Quebec_MontTremblantNP_voyou0833M_RWBH001 | 0,000 | 0,000 | 0,968 | 0,000 |
| 125 | Asia | East Asia | RKW13451 | 0,000 | 0,000 | 0,155 | 0,000 |
| 126 | Europe | Karelian | V113 | 0,403 | 0,001 | 0,000 | 0,411 |
| 127 | Europe | Karelian | V114 | 0,381 | 0,004 | 0,000 | 0,358 |
| 128 | Europe | Karelian | V115 | 0,369 | 0,000 | 0,000 | 0,371 |
| 129 | Europe | Karelian | V116 | 0,359 | 0,000 | 0,000 | 0,348 |
| 130 | Europe | Karelian | V117 | 0,378 | 0,008 | 0,000 | 0,346 |
| 131 | Europe | Karelian | V119 | 0,390 | 0,000 | 0,000 | 0,471 |
| 132 | Europe | Karelian | V120 | 0,379 | 0,000 | 0,000 | 0,424 |

**Table S8.** ADMIXTURE results for K = 6 for European, Asian and North American wolves.

| Number | Continent | Population | Sample | | | Admixture_k6 | |
|---|---|---|---|---|---|---|---|
| 133 | Europe | Karelian | V126 | 0,350 | 0,000 | 0,000 | 0,473 |
| 134 | Europe | Karelian | V132 | 0,382 | 0,016 | 0,000 | 0,332 |
| 135 | Europe | Karelian | V134 | 0,392 | 0,000 | 0,000 | 0,433 |
| 136 | Europe | Karelian | V136 | 0,356 | 0,000 | 0,000 | 0,394 |
| 137 | Europe | Karelian | V141 | 0,409 | 0,000 | 0,000 | 0,416 |
| 138 | Europe | Karelian | V143 | 0,402 | 0,000 | 0,000 | 0,427 |
| 139 | Europe | Italian Peninsula | W1023 | 0,000 | 1,000 | 0,000 | 0,000 |
| 140 | Europe | Karelian | W11 | 0,343 | 0,000 | 0,000 | 0,488 |
| 141 | Europe | Karelian | W14 | 0,399 | 0,007 | 0,000 | 0,416 |
| 142 | Europe | Italian Peninsula | W1551 | 0,000 | 1,000 | 0,000 | 0,000 |
| 143 | Europe | Italian Peninsula | W1593 | 0,082 | 0,882 | 0,000 | 0,027 |
| 144 | Europe | Karelian | W16 | 0,404 | 0,000 | 0,000 | 0,474 |
| 145 | Europe | Italian Peninsula | W2030 | 0,000 | 1,000 | 0,000 | 0,000 |
| 146 | Europe | Karelian | W21 | 0,388 | 0,000 | 0,000 | 0,500 |
| 147 | Europe | Italian Peninsula | W2124 | 0,000 | 1,000 | 0,000 | 0,000 |
| 148 | Europe | Italian Peninsula | W2185 | 0,000 | 1,000 | 0,000 | 0,000 |
| 149 | Europe | Italian Peninsula | W2193 | 0,000 | 1,000 | 0,000 | 0,000 |
| 150 | Europe | Italian Peninsula | W2265 | 0,000 | 1,000 | 0,000 | 0,000 |
| 151 | Europe | Italian Peninsula | W2280 | 0,000 | 1,000 | 0,000 | 0,000 |
| 152 | Europe | Italian Peninsula | W2291 | 0,000 | 1,000 | 0,000 | 0,000 |
| 153 | Europe | Italian Peninsula | W2371F | 0,000 | 1,000 | 0,000 | 0,000 |
| 154 | Europe | Italian Peninsula | W2430F | 0,000 | 1,000 | 0,000 | 0,000 |
| 155 | Europe | Italian Peninsula | W2487F | 0,000 | 1,000 | 0,000 | 0,000 |
| 156 | Europe | Italian Peninsula | W2594M | 0,000 | 1,000 | 0,000 | 0,000 |
| 157 | Europe | Karelian | W26 | 0,390 | 0,000 | 0,000 | 0,487 |
| 158 | Europe | Italian Peninsula | W2623M | 0,000 | 1,000 | 0,000 | 0,000 |
| 159 | Europe | Italian Peninsula | W2707M | 0,000 | 1,000 | 0,000 | 0,000 |
| 160 | Europe | Italian Peninsula | W2878M | 0,000 | 1,000 | 0,000 | 0,000 |
| 161 | Europe | Italian Peninsula | W2883F | 0,000 | 1,000 | 0,000 | 0,000 |
| 162 | Europe | Italian Peninsula | W2886 | 0,048 | 0,924 | 0,000 | 0,029 |
| 163 | Europe | Italian Peninsula | W2910 | 0,000 | 1,000 | 0,000 | 0,000 |
| 164 | Europe | Italian Peninsula | W2951 | 0,000 | 1,000 | 0,000 | 0,000 |
| 165 | Europe | Karelian | W32 | 0,384 | 0,000 | 0,000 | 0,463 |

**Table S8.** ADMIXTURE results for K = 6 for European, Asian and North American wolves.

| Number | Continent | Population | Sample | | | Admixture_k6 | |
|---|---|---|---|---|---|---|---|
| 166 | Europe | Karelian | W4 | 0,397 | 0,000 | 0,000 | 0,489 |
| 167 | Europe | Karelian | W44 | 0,395 | 0,000 | 0,000 | 0,419 |
| 168 | Europe | Karelian | W45 | 0,377 | 0,000 | 0,000 | 0,439 |
| 169 | Europe | Italian Peninsula | W893 | 0,000 | 1,000 | 0,000 | 0,000 |
| 170 | Europe | Italian Peninsula | WIT11 | 0,000 | 1,000 | 0,000 | 0,000 |
| 171 | Europe | Italian Peninsula | WIT13 | 0,000 | 1,000 | 0,000 | 0,000 |
| 172 | Europe | Italian Peninsula | WIT14 | 0,000 | 1,000 | 0,000 | 0,000 |
| 173 | Europe | Italian Peninsula | WIT16 | 0,000 | 1,000 | 0,000 | 0,000 |
| 174 | Europe | Italian Peninsula | WIT17 | 0,000 | 1,000 | 0,000 | 0,000 |
| 175 | Europe | Italian Peninsula | WIT19 | 0,000 | 1,000 | 0,000 | 0,000 |
| 176 | Europe | Italian Peninsula | WIT20 | 0,000 | 1,000 | 0,000 | 0,000 |
| 177 | Europe | Italian Peninsula | WIT21 | 0,000 | 1,000 | 0,000 | 0,000 |
| 178 | Europe | Italian Peninsula | WIT22 | 0,000 | 1,000 | 0,000 | 0,000 |
| 179 | Europe | Italian Peninsula | WIT23 | 0,000 | 1,000 | 0,000 | 0,000 |
| 180 | Asia | Turkey | WTR026 | 0,108 | 0,000 | 0,000 | 0,000 |
| 181 | Asia | Turkey | WTR028 | 0,100 | 0,000 | 0,000 | 0,000 |
| 182 | Asia | Turkey | WTR029 | 0,094 | 0,000 | 0,000 | 0,000 |
| 183 | Asia | Turkey | WTR030 | 0,081 | 0,000 | 0,000 | 0,000 |
| 184 | Asia | Turkey | WTR031 | 0,044 | 0,000 | 0,000 | 0,000 |
| 185 | Asia | Turkey | WTR032 | 0,113 | 0,000 | 0,000 | 0,000 |
| 186 | Asia | Turkey | WTR033 | 0,069 | 0,000 | 0,000 | 0,000 |
| 187 | Asia | Turkey | WTR034 | 0,098 | 0,000 | 0,000 | 0,000 |
| 188 | Asia | Turkey | WTR035 | 0,111 | 0,000 | 0,000 | 0,000 |
| 189 | Asia | Turkey | WTR037 | 0,107 | 0,000 | 0,000 | 0,000 |
| 190 | Asia | Turkey | WTR038 | 0,101 | 0,000 | 0,000 | 0,000 |
| 191 | Asia | Turkey | WTR040 | 0,098 | 0,000 | 0,000 | 0,000 |
| 192 | Asia | Turkey | WTR041 | 0,034 | 0,000 | 0,000 | 0,000 |
| 193 | North America | North America | glw | 0,000 | 0,000 | 1,000 | 0,000 |
| 194 | Asia | Indian | inw | 0,000 | 0,000 | 0,000 | 0,000 |
| 195 | Asia | West Asia | irw | 0,007 | 0,000 | 0,000 | 0,000 |
| 196 | North America | North America | mxa | 0,000 | 0,016 | 0,925 | 0,000 |
| 197 | Europe | NW Iberia | ptw | 0,000 | 0,000 | 0,000 | 0,000 |
| 198 | Europe | Italian Peninsula | w361703 | 0,000 | 1,000 | 0,000 | 0,000 |

**Table S8.** ADMIXTURE results for K = 6 for European, Asian and North American wolves.

| Number | Continent | Population | Sample | | | Admixture_k6 | |
|---|---|---|---|---|---|---|---|
| 199 | Europe | Italian Peninsula | w361756 | 0,044 | 0,940 | 0,000 | 0,015 |
| 200 | Europe | Italian Peninsula | w4926402 | 0,000 | 1,000 | 0,000 | 0,000 |
| 201 | North America | North America | ysa | 0,000 | 0,000 | 1,000 | 0,000 |
| 202 | North America | North America | ysb | 0,000 | 0,000 | 1,000 | 0,000 |

| | |
|---|---|
| 0,000 | 0,000 |
| 0,000 | 0,000 |
| 0,000 | 0,000 |
| 0,000 | 0,000 |
| 0,000 | 0,000 |
| 0,000 | 0,000 |
| 0,000 | 0,000 |
| 0,000 | 0,000 |
| 0,000 | 0,000 |
| 0,000 | 0,000 |
| 0,000 | 0,000 |
| 0,000 | 0,000 |
| 0,000 | 0,000 |
| 0,000 | 0,000 |
| 0,000 | 0,000 |
| 0,000 | 0,000 |
| 1,000 | 0,000 |
| 1,000 | 0,000 |
| 1,000 | 0,000 |
| 0,824 | 0,000 |
| 0,839 | 0,000 |
| 0,791 | 0,001 |
| 0,831 | 0,000 |
| 1,000 | 0,000 |
| 0,000 | 0,000 |
| 0,000 | 0,000 |
| 0,905 | 0,000 |
| 0,813 | 0,000 |
| 1,000 | 0,000 |
| 0,827 | 0,000 |
| 0,806 | 0,000 |
| 0,810 | 0,001 |
| 0,796 | 0,000 |

| | |
|---|---|
| 0,805 | 0,000 |
| 0,824 | 0,000 |
| 0,932 | 0,000 |
| 0,923 | 0,000 |
| 0,928 | 0,000 |
| 0,928 | 0,000 |
| 0,792 | 0,011 |
| 0,806 | 0,003 |
| 0,753 | 0,009 |
| 0,741 | 0,004 |
| 0,605 | 0,003 |
| 0,687 | 0,000 |
| 0,231 | 0,050 |
| 0,237 | 0,068 |
| 0,233 | 0,069 |
| 0,243 | 0,036 |
| 0,267 | 0,038 |
| 1,000 | 0,000 |
| 1,000 | 0,000 |
| 1,000 | 0,000 |
| 1,000 | 0,000 |
| 1,000 | 0,000 |
| 1,000 | 0,000 |
| 1,000 | 0,000 |
| 0,000 | 0,000 |
| 0,000 | 0,000 |
| 0,000 | 0,000 |
| 0,000 | 0,000 |
| 0,000 | 0,000 |
| 0,000 | 0,000 |
| 0,000 | 0,000 |
| 0,000 | 0,000 |
| 0,111 | 0,017 |

| | |
|---|---|
| 0,000 | 0,000 |
| 0,000 | 0,000 |
| 0,000 | 0,000 |
| 0,000 | 0,000 |
| 0,000 | 0,000 |
| 0,000 | 0,000 |
| 0,000 | 0,000 |
| 0,000 | 0,000 |
| 0,000 | 0,000 |
| 0,000 | 0,000 |
| 0,000 | 0,000 |
| 0,000 | 0,000 |
| 0,000 | 0,000 |
| 0,000 | 0,000 |
| 0,000 | 0,000 |
| 0,000 | 0,000 |
| 0,000 | 1,000 |
| 0,000 | 1,000 |
| 0,000 | 1,000 |
| 0,000 | 1,000 |
| 0,000 | 1,000 |
| 0,000 | 1,000 |
| 0,000 | 1,000 |
| 0,000 | 1,000 |
| 0,000 | 1,000 |
| 0,000 | 1,000 |
| 0,000 | 1,000 |
| 0,000 | 0,000 |
| 0,079 | 0,038 |
| 0,000 | 0,000 |
| 0,000 | 0,018 |
| 0,000 | 0,000 |
| 0,000 | 0,012 |

| | |
|---|---|
| 0,000 | 0,000 |
| 0,030 | 0,037 |
| 0,000 | 0,000 |
| 0,000 | 0,000 |
| 0,000 | 0,000 |
| 0,000 | 0,000 |
| 0,000 | 0,987 |
| 0,000 | 1,000 |
| 0,821 | 0,000 |
| 0,812 | 0,001 |
| 0,000 | 0,000 |
| 0,000 | 0,000 |
| 0,000 | 0,000 |
| 0,000 | 0,000 |
| 0,000 | 0,000 |
| 0,000 | 0,000 |
| 0,000 | 1,000 |
| 0,000 | 1,000 |
| 0,000 | 0,000 |
| 0,000 | 0,000 |
| 0,000 | 0,000 |
| 0,000 | 0,000 |
| 0,000 | 0,000 |
| 0,000 | 1,000 |
| 0,032 | 0,000 |
| 0,845 | 0,000 |
| 0,157 | 0,028 |
| 0,223 | 0,035 |
| 0,235 | 0,025 |
| 0,270 | 0,023 |
| 0,229 | 0,039 |
| 0,122 | 0,017 |
| 0,175 | 0,022 |

| | |
|---|---|
| 0,160 | 0,018 |
| 0,216 | 0,055 |
| 0,159 | 0,016 |
| 0,222 | 0,028 |
| 0,154 | 0,021 |
| 0,150 | 0,021 |
| 0,000 | 0,000 |
| 0,148 | 0,022 |
| 0,145 | 0,033 |
| 0,000 | 0,000 |
| 0,009 | 0,000 |
| 0,121 | 0,002 |
| 0,000 | 0,000 |
| 0,097 | 0,015 |
| 0,000 | 0,000 |
| 0,000 | 0,000 |
| 0,000 | 0,000 |
| 0,000 | 0,000 |
| 0,000 | 0,000 |
| 0,000 | 0,000 |
| 0,000 | 0,000 |
| 0,000 | 0,000 |
| 0,000 | 0,000 |
| 0,000 | 0,000 |
| 0,112 | 0,011 |
| 0,000 | 0,000 |
| 0,000 | 0,000 |
| 0,000 | 0,000 |
| 0,000 | 0,000 |
| 0,000 | 0,000 |
| 0,000 | 0,000 |
| 0,000 | 0,000 |
| 0,135 | 0,018 |

| | |
|---|---|
| 0,099 | 0,015 |
| 0,161 | 0,025 |
| 0,167 | 0,017 |
| 0,000 | 0,000 |
| 0,000 | 0,000 |
| 0,000 | 0,000 |
| 0,000 | 0,000 |
| 0,000 | 0,000 |
| 0,000 | 0,000 |
| 0,000 | 0,000 |
| 0,000 | 0,000 |
| 0,000 | 0,000 |
| 0,000 | 0,000 |
| 0,000 | 0,000 |
| 0,892 | 0,000 |
| 0,900 | 0,000 |
| 0,906 | 0,000 |
| 0,919 | 0,000 |
| 0,956 | 0,000 |
| 0,887 | 0,000 |
| 0,931 | 0,000 |
| 0,902 | 0,000 |
| 0,889 | 0,000 |
| 0,893 | 0,000 |
| 0,899 | 0,000 |
| 0,902 | 0,000 |
| 0,966 | 0,000 |
| 0,000 | 0,000 |
| 1,000 | 0,000 |
| 0,993 | 0,000 |
| 0,059 | 0,000 |
| 0,000 | 1,000 |
| 0,000 | 0,000 |

| | |
|---|---|
| 0,000 | 0,000 |
| 0,000 | 0,000 |
| 0,000 | 0,000 |
| 0,000 | 0,000 |

**Table S9.** ADMIXTURE results for K = 5 for European wolves.

| Number | Continent | Population | Sample | Admixture_k5 | | | |
|---|---|---|---|---|---|---|---|
| 1 | Europe | Dinaric-Balkan | AP_085T | 1,000 | 0,000 | 0,000 | 0,000 |
| 2 | Europe | Dinaric-Balkan | AP_086E | 1,000 | 0,000 | 0,000 | 0,000 |
| 3 | Europe | Dinaric-Balkan | AP_087A | 1,000 | 0,000 | 0,000 | 0,000 |
| 4 | Europe | Dinaric-Balkan | AP_087E | 1,000 | 0,000 | 0,000 | 0,000 |
| 5 | Europe | Dinaric-Balkan | AP_087L | 1,000 | 0,000 | 0,000 | 0,000 |
| 6 | Europe | Dinaric-Balkan | AP_087U | 1,000 | 0,000 | 0,000 | 0,000 |
| 7 | Europe | Dinaric-Balkan | AP_0882 | 1,000 | 0,000 | 0,000 | 0,000 |
| 8 | Europe | Dinaric-Balkan | AP_088L | 1,000 | 0,000 | 0,000 | 0,000 |
| 9 | Europe | Dinaric-Balkan | AP_08A1 | 1,000 | 0,000 | 0,000 | 0,000 |
| 10 | Europe | Dinaric-Balkan | AP_08A3 | 1,000 | 0,000 | 0,000 | 0,000 |
| 11 | Europe | Dinaric-Balkan | AP_08A6 | 0,992 | 0,000 | 0,000 | 0,008 |
| 12 | Europe | Dinaric-Balkan | AP_08C3 | 1,000 | 0,000 | 0,000 | 0,000 |
| 13 | Europe | Dinaric-Balkan | BiH_244 | 1,000 | 0,000 | 0,000 | 0,000 |
| 14 | Europe | Dinaric-Balkan | BiH_245 | 1,000 | 0,000 | 0,000 | 0,000 |
| 15 | Europe | Karelian | CLUPRU000005 | 0,041 | 0,000 | 0,008 | 0,017 |
| 16 | Europe | Karelian | CLUPRU000007 | 0,160 | 0,011 | 0,000 | 0,050 |
| 17 | Europe | Karelian | CLUPRU000008 | 0,162 | 0,019 | 0,000 | 0,053 |
| 18 | Europe | Karelian | CLUPRU000019 | 0,008 | 0,000 | 0,000 | 0,000 |
| 19 | Europe | Karelian | CLUPRU000020 | 0,010 | 0,000 | 0,000 | 0,000 |
| 20 | Europe | Scandinavian | D-00-12 | 0,000 | 0,000 | 1,000 | 0,000 |
| 21 | Europe | Scandinavian | D-06-14 | 0,000 | 0,000 | 1,000 | 0,000 |
| 22 | Europe | Scandinavian | D-07-16 | 0,000 | 0,000 | 1,000 | 0,000 |
| 23 | Europe | Scandinavian | D-08-10 | 0,000 | 0,000 | 1,000 | 0,000 |
| 24 | Europe | Scandinavian | D-10-68 | 0,000 | 0,000 | 1,000 | 0,000 |
| 25 | Europe | Scandinavian | D-11-58 | 0,000 | 0,000 | 1,000 | 0,000 |
| 26 | Europe | Scandinavian | D-85-02 | 0,000 | 0,000 | 0,763 | 0,000 |
| 27 | Europe | Scandinavian | G100-12 | 0,000 | 0,000 | 0,785 | 0,000 |
| 28 | Europe | Scandinavian | G100-14 | 0,000 | 0,000 | 0,000 | 0,000 |
| 29 | Europe | Scandinavian | G126-13 | 0,000 | 0,000 | 1,000 | 0,000 |
| 30 | Europe | Scandinavian | G139-12 | 0,000 | 0,000 | 0,764 | 0,000 |
| 31 | Europe | Scandinavian | G50-12 | 0,000 | 0,000 | 1,000 | 0,000 |
| 32 | Europe | NW Iberia | JAL7480 | 0,000 | 0,000 | 0,000 | 1,000 |
| 33 | Europe | NW Iberia | JAL7486 | 0,000 | 0,000 | 0,000 | 1,000 |

**Table S9.** ADMIXTURE results for K = 5 for European wolves.

| Number | Continent | Population | Sample | Admixture_k5 | | | |
|---|---|---|---|---|---|---|---|
| 34 | Europe | NW Iberia | JAL7491 | 0,000 | 0,000 | 0,000 | 1,000 |
| 35 | Europe | NW Iberia | JAL7492 | 0,000 | 0,000 | 0,000 | 1,000 |
| 36 | Europe | NW Iberia | L166 | 0,000 | 0,000 | 0,000 | 1,000 |
| 37 | Europe | NW Iberia | L253 | 0,000 | 0,000 | 0,000 | 1,000 |
| 38 | Europe | NW Iberia | L409 | 0,000 | 0,000 | 0,000 | 1,000 |
| 39 | Europe | NW Iberia | L514 | 0,000 | 0,000 | 0,000 | 1,000 |
| 40 | Europe | NW Iberia | L547 | 0,000 | 0,000 | 0,000 | 1,000 |
| 41 | Europe | NW Iberia | L552 | 0,000 | 0,000 | 0,000 | 1,000 |
| 42 | Europe | NW Iberia | L588 | 0,000 | 0,000 | 0,000 | 1,000 |
| 43 | Europe | Dinaric-Balkan | LUP006817 | 0,985 | 0,000 | 0,000 | 0,015 |
| 44 | Europe | Dinaric-Balkan | LUP006818 | 0,761 | 0,018 | 0,000 | 0,054 |
| 45 | Europe | Dinaric-Balkan | LUP006819 | 0,864 | 0,016 | 0,000 | 0,023 |
| 46 | Europe | Dinaric-Balkan | LUP006820 | 0,846 | 0,020 | 0,000 | 0,039 |
| 47 | Europe | Dinaric-Balkan | LUP006821 | 0,933 | 0,000 | 0,000 | 0,013 |
| 48 | Europe | Dinaric-Balkan | LUP006822 | 0,890 | 0,011 | 0,000 | 0,041 |
| 49 | Europe | Dinaric-Balkan | LUP006823 | 1,000 | 0,000 | 0,000 | 0,000 |
| 50 | Europe | Dinaric-Balkan | LUP006824 | 0,786 | 0,018 | 0,000 | 0,053 |
| 51 | Europe | Dinaric-Balkan | LUP006825 | 0,906 | 0,000 | 0,000 | 0,025 |
| 52 | Europe | Dinaric-Balkan | LUP006826 | 0,913 | 0,006 | 0,000 | 0,028 |
| 53 | Europe | Dinaric-Balkan | LUP006827 | 1,000 | 0,000 | 0,000 | 0,000 |
| 54 | Europe | Dinaric-Balkan | LUP006828 | 1,000 | 0,000 | 0,000 | 0,000 |
| 55 | Europe | NW Iberia | LUP006829 | 0,000 | 0,000 | 0,000 | 1,000 |
| 56 | Europe | NW Iberia | LUP006830 | 0,000 | 0,000 | 0,000 | 1,000 |
| 57 | Europe | Scandinavian | M-01-06 | 0,000 | 0,000 | 1,000 | 0,000 |
| 58 | Europe | Scandinavian | M-06-03 | 0,000 | 0,000 | 1,000 | 0,000 |
| 59 | Europe | Scandinavian | M-09-05 | 0,000 | 0,000 | 1,000 | 0,000 |
| 60 | Europe | Scandinavian | M-10-04 | 0,000 | 0,000 | 0,674 | 0,000 |
| 61 | Europe | Scandinavian | M-98-08 | 0,000 | 0,000 | 1,000 | 0,000 |
| 62 | Europe | Dinaric-Balkan | MSV2AC | 1,000 | 0,000 | 0,000 | 0,000 |
| 63 | Europe | NW Iberia | MW122 | 0,000 | 0,000 | 0,000 | 1,000 |
| 64 | Europe | NW Iberia | MW127 | 0,000 | 0,000 | 0,000 | 1,000 |
| 65 | Europe | NW Iberia | Penelope | 0,000 | 0,000 | 0,000 | 1,000 |
| 66 | Europe | Karelian | V113 | 0,000 | 0,000 | 0,000 | 0,000 |

**Table S9.** ADMIXTURE results for K = 5 for European wolves.

| Number | Continent | Population | Sample | Admixture_k5 | | | |
|---|---|---|---|---|---|---|---|
| 67 | Europe | Karelian | V114 | 0,000 | 0,000 | 0,000 | 0,000 |
| 68 | Europe | Karelian | V115 | 0,000 | 0,000 | 0,000 | 0,000 |
| 69 | Europe | Karelian | V116 | 0,000 | 0,000 | 0,000 | 0,000 |
| 70 | Europe | Karelian | V117 | 0,039 | 0,000 | 0,000 | 0,006 |
| 71 | Europe | Karelian | V119 | 0,000 | 0,000 | 0,000 | 0,000 |
| 72 | Europe | Karelian | V120 | 0,000 | 0,000 | 0,000 | 0,000 |
| 73 | Europe | Karelian | V126 | 0,000 | 0,000 | 0,091 | 0,000 |
| 74 | Europe | Karelian | V132 | 0,081 | 0,005 | 0,011 | 0,029 |
| 75 | Europe | Karelian | V134 | 0,000 | 0,000 | 0,000 | 0,000 |
| 76 | Europe | Karelian | V136 | 0,000 | 0,000 | 0,001 | 0,000 |
| 77 | Europe | Karelian | V141 | 0,000 | 0,000 | 0,000 | 0,000 |
| 78 | Europe | Karelian | V143 | 0,000 | 0,000 | 0,000 | 0,000 |
| 79 | Europe | Italian Peninsula | W1023 | 0,000 | 1,000 | 0,000 | 0,000 |
| 80 | Europe | Karelian | W11 | 0,000 | 0,000 | 0,097 | 0,000 |
| 81 | Europe | Karelian | W14 | 0,000 | 0,000 | 0,000 | 0,000 |
| 82 | Europe | Italian Peninsula | W1551 | 0,000 | 1,000 | 0,000 | 0,000 |
| 83 | Europe | Italian Peninsula | W1593 | 0,016 | 0,876 | 0,000 | 0,000 |
| 84 | Europe | Karelian | W16 | 0,000 | 0,000 | 0,000 | 0,000 |
| 85 | Europe | Italian Peninsula | W2030 | 0,000 | 1,000 | 0,000 | 0,000 |
| 86 | Europe | Karelian | W21 | 0,000 | 0,000 | 0,019 | 0,000 |
| 87 | Europe | Italian Peninsula | W2124 | 0,000 | 1,000 | 0,000 | 0,000 |
| 88 | Europe | Italian Peninsula | W2185 | 0,000 | 1,000 | 0,000 | 0,000 |
| 89 | Europe | Italian Peninsula | W2193 | 0,000 | 1,000 | 0,000 | 0,000 |
| 90 | Europe | Italian Peninsula | W2265 | 0,000 | 1,000 | 0,000 | 0,000 |
| 91 | Europe | Italian Peninsula | W2280 | 0,000 | 1,000 | 0,000 | 0,000 |
| 92 | Europe | Italian Peninsula | W2291 | 0,000 | 1,000 | 0,000 | 0,000 |
| 93 | Europe | Italian Peninsula | W2371F | 0,000 | 1,000 | 0,000 | 0,000 |
| 94 | Europe | Italian Peninsula | W2430F | 0,000 | 1,000 | 0,000 | 0,000 |
| 95 | Europe | Italian Peninsula | W2487F | 0,000 | 0,993 | 0,000 | 0,008 |
| 96 | Europe | Italian Peninsula | W2594M | 0,000 | 1,000 | 0,000 | 0,000 |
| 97 | Europe | Karelian | W26 | 0,000 | 0,000 | 0,008 | 0,000 |
| 98 | Europe | Italian Peninsula | W2623M | 0,000 | 1,000 | 0,000 | 0,000 |
| 99 | Europe | Italian Peninsula | W2707M | 0,000 | 1,000 | 0,000 | 0,000 |

**Table S9.** ADMIXTURE results for K = 5 for European wolves.

| Number | Continent | Population | Sample | Admixture_k5 | | | |
|---|---|---|---|---|---|---|---|
| 100 | Europe | Italian Peninsula | W2878M | 0,000 | 1,000 | 0,000 | 0,000 |
| 101 | Europe | Italian Peninsula | W2883F | 0,000 | 1,000 | 0,000 | 0,000 |
| 102 | Europe | Italian Peninsula | W2886 | 0,000 | 0,918 | 0,005 | 0,000 |
| 103 | Europe | Italian Peninsula | W2910 | 0,000 | 1,000 | 0,000 | 0,000 |
| 104 | Europe | Italian Peninsula | W2951 | 0,000 | 1,000 | 0,000 | 0,000 |
| 105 | Europe | Karelian | W32 | 0,000 | 0,000 | 0,000 | 0,000 |
| 106 | Europe | Karelian | W4 | 0,000 | 0,000 | 0,000 | 0,000 |
| 107 | Europe | Karelian | W44 | 0,000 | 0,000 | 0,000 | 0,000 |
| 108 | Europe | Karelian | W45 | 0,000 | 0,000 | 0,004 | 0,000 |
| 109 | Europe | Italian Peninsula | W893 | 0,000 | 1,000 | 0,000 | 0,000 |
| 110 | Europe | Italian Peninsula | WIT11 | 0,000 | 1,000 | 0,000 | 0,000 |
| 111 | Europe | Italian Peninsula | WIT13 | 0,000 | 1,000 | 0,000 | 0,000 |
| 112 | Europe | Italian Peninsula | WIT14 | 0,000 | 1,000 | 0,000 | 0,000 |
| 113 | Europe | Italian Peninsula | WIT16 | 0,000 | 1,000 | 0,000 | 0,000 |
| 114 | Europe | Italian Peninsula | WIT17 | 0,000 | 1,000 | 0,000 | 0,000 |
| 115 | Europe | Italian Peninsula | WIT19 | 0,000 | 1,000 | 0,000 | 0,000 |
| 116 | Europe | Italian Peninsula | WIT20 | 0,000 | 1,000 | 0,000 | 0,000 |
| 117 | Europe | Italian Peninsula | WIT21 | 0,000 | 1,000 | 0,000 | 0,000 |
| 118 | Europe | Italian Peninsula | WIT22 | 0,000 | 1,000 | 0,000 | 0,000 |
| 119 | Europe | Italian Peninsula | WIT23 | 0,000 | 1,000 | 0,000 | 0,000 |
| 120 | Europe | NW Iberia | ptw | 0,000 | 0,000 | 0,000 | 1,000 |
| 121 | Europe | Italian Peninsula | w361703 | 0,000 | 1,000 | 0,000 | 0,000 |
| 122 | Europe | Italian Peninsula | w361756 | 0,000 | 0,933 | 0,000 | 0,005 |
| 123 | Europe | Italian Peninsula | w4926402 | 0,000 | 1,000 | 0,000 | 0,000 |

| |
|---|
| 0,000 |
| 0,000 |
| 0,000 |
| 0,000 |
| 0,000 |
| 0,000 |
| 0,000 |
| 0,000 |
| 0,000 |
| 0,000 |
| 0,000 |
| 0,000 |
| 0,000 |
| 0,000 |
| 0,935 |
| 0,779 |
| 0,766 |
| 0,992 |
| 0,990 |
| 0,000 |
| 0,000 |
| 0,000 |
| 0,000 |
| 0,000 |
| 0,000 |
| 0,237 |
| 0,215 |
| 1,000 |
| 0,000 |
| 0,236 |
| 0,000 |
| 0,000 |
| 0,000 |

| |
|---|
| 0,000 |
| 0,000 |
| 0,000 |
| 0,000 |
| 0,000 |
| 0,000 |
| 0,000 |
| 0,000 |
| 0,000 |
| 0,000 |
| 0,167 |
| 0,098 |
| 0,096 |
| 0,053 |
| 0,058 |
| 0,000 |
| 0,143 |
| 0,069 |
| 0,053 |
| 0,000 |
| 0,000 |
| 0,000 |
| 0,000 |
| 0,000 |
| 0,000 |
| 0,000 |
| 0,326 |
| 0,000 |
| 0,000 |
| 0,000 |
| 0,000 |
| 0,000 |
| 1,000 |

| |
|---|
| 1,000 |
| 1,000 |
| 1,000 |
| 0,955 |
| 1,000 |
| 1,000 |
| 0,909 |
| 0,874 |
| 1,000 |
| 0,999 |
| 1,000 |
| 1,000 |
| 0,000 |
| 0,904 |
| 1,000 |
| 0,000 |
| 0,108 |
| 1,000 |
| 0,000 |
| 0,981 |
| 0,000 |
| 0,000 |
| 0,000 |
| 0,000 |
| 0,000 |
| 0,000 |
| 0,000 |
| 0,000 |
| 0,000 |
| 0,000 |
| 0,992 |
| 0,000 |
| 0,000 |

| |
|---|
| 0,000 |
| 0,000 |
| 0,078 |
| 0,000 |
| 0,000 |
| 1,000 |
| 1,000 |
| 1,000 |
| 0,996 |
| 0,000 |
| 0,000 |
| 0,000 |
| 0,000 |
| 0,000 |
| 0,000 |
| 0,000 |
| 0,000 |
| 0,000 |
| 0,000 |
| 0,000 |
| 0,000 |
| 0,000 |
| 0,062 |
| 0,000 |

**Table S10**. Time of divergence (yrs, average) across comparisons of the different Euroepan wolf populations using TT-method and MisTI.

**MiSTI**

| | **Dinaric** | **Iberian** | **Italian** | **Karelian** | **Scandinavian** |
|---|---|---|---|---|---|
| **Dinaric** | | 28.202 | 12.869 | 28.665 | 38.782 |
| **Iberian** | 9.236 | | 13,67 | 21.078 | 19.804 |
| **Italian** | 12.147 | 5.873 | | 14.328 | 14.971 |
| **Karelian** | 12.602 | 18.916 | 21.918 | | 25.287 |
| **Scandinavian** | 25.894 | 37.172 | 36.717 | 2.286 | |

**TT-method**

TT-method: using only autosomes

**Table S11.** Statistical comparisons between the two methods to estimate divergence time (TT-method and MiSTI). Non-signinficant values (p >

| comparison | method1 | method2 | t_stat | t_pvalue | U_stat | U_pvalue |
|---|---|---|---|---|---|---|
| Dinaric – Iberian | TT | misti | -4,244 | 0,000 | 96 | 0,00 |
| **Dinaric – Italian** | **TT** | **misti** | **-0,374** | **0,712** | **269** | **0,40** |
| Dinaric – Karelian | TT | misti | -4,353 | 0,000 | 112 | 0,00 |
| **Dinaric – Scandinavian** | **TT** | **misti** | **-1,717** | **0,099** | **203** | **0,95** |
| Iberian – Italian | TT | misti | -4,473 | 0,000 | 121 | 0,00 |
| **Iberian – Karelian** | **TT** | **misti** | **-0,656** | **0,515** | **376** | **0,99** |
| Iberian – Scandinavian | TT | misti | 3,516 | 0,011 | 344 | 0,00 |
| Italian – Karelian | TT | misti | 4,032 | 0,000 | 627 | 0,00 |
| Italian – Scandinavian | TT | misti | 6,788 | 0,000 | 400 | 0,00 |
| Karelian – Scandinavian | TT | misti | -5,709 | 0,000 | 25 | 0,00 |

0.05 threshold) in bold.

**Table S12**. Nucleotide diversity in Europe and Asia. The Italian Peninsula and Scandinavian wolves show the lowest values.

| **Continent** | **Population** | **Mean $\pi$** | **CI 95%** |
|---|---|---|---|
| Europe | Dinaric-Balkan | 0.001421 | 0.001411 - 0.001430 |
| Europe | Italian Peninsula | 0.000992 | 0.000984 - 0.001000 |
| Europe | Karelian | 0.001446 | 0.001436 - 0.001456 |
| Europe | NW Iberia | 0.001226 | 0.001217 - 0.001234 |
| Europe | Scandinavia | 0.001068 | 0.001060 - 0.001076 |
| Asia | Western Asia | 0.001485 | 0.001475 - 0.001495 |
| Asia | Central Asia | 0.001386 | 0.001377 - 0.001396 |
| Asia | Eastern Asia | 0.001491 | 0.001481 - 0.001501 |
| Asia | Tibetan | 0.001236 | 0.001228 - 0.001244 |

**Table S13.** Individual-level heterozygosity and inbreeding coefficients for European wolf samples.

| Sample ID | Population | Continent | Observed homozygosity (SN | Expected homozygosity (SN | Total numl | *F* (Inbreeding coefficient) |
|---|---|---|---|---|---|---|
| AP_085T | Dinaric-Balkan | Europe | 13613875 | 12808311,2 | 16492388 | 0,219 |
| AP_086E | Dinaric-Balkan | Europe | 13319871 | 12810674,6 | 16495432 | 0,138 |
| AP_086F | Dinaric-Balkan | Europe | 13349193 | 12796869,1 | 16477902 | 0,150 |
| AP_087A | Dinaric-Balkan | Europe | 13553542 | 12795405,7 | 16476048 | 0,206 |
| AP_087E | Dinaric-Balkan | Europe | 13429900 | 12803217,6 | 16485736 | 0,170 |
| AP_087L | Dinaric-Balkan | Europe | 14031242 | 12806106,6 | 16489617 | 0,333 |
| AP_087U | Dinaric-Balkan | Europe | 13267464 | 12808471,7 | 16492633 | 0,125 |
| AP_0882 | Dinaric-Balkan | Europe | 14127418 | 12800390 | 16481949 | 0,360 |
| AP_088L | Dinaric-Balkan | Europe | 13574416 | 12807461,8 | 16491112 | 0,208 |
| AP_08A1 | Dinaric-Balkan | Europe | 13167030 | 12805602,3 | 16489000 | 0,098 |
| AP_08A3 | Dinaric-Balkan | Europe | 13402050 | 12798702,6 | 16479871 | 0,164 |
| AP_08A6 | Dinaric-Balkan | Europe | 13110888 | 12808168,3 | 16492137 | 0,082 |
| AP_08C3 | Dinaric-Balkan | Europe | 13386651 | 12804332,8 | 16487194 | 0,158 |
| AP_08C5 | Dinaric-Balkan | Europe | 13606402 | 12808001,6 | 16491938 | 0,217 |
| BiH_244 | Dinaric-Balkan | Europe | 13258884 | 12804943,4 | 16488012 | 0,123 |
| BiH_245 | Dinaric-Balkan | Europe | 13292206 | 12773595,6 | 16448481 | 0,141 |
| LUP006817 | Dinaric-Balkan | Europe | 13219961 | 12777422,9 | 16452994 | 0,120 |
| LUP006818 | Dinaric-Balkan | Europe | 13360119 | 12795301,2 | 16475471 | 0,153 |
| LUP006819 | Dinaric-Balkan | Europe | 13143331 | 12762961,3 | 16434625 | 0,104 |
| LUP006820 | Dinaric-Balkan | Europe | 13263787 | 12752995,7 | 16421908 | 0,139 |
| LUP006821 | Dinaric-Balkan | Europe | 13231609 | 12787007,1 | 16464844 | 0,121 |
| LUP006822 | Dinaric-Balkan | Europe | 13207666 | 12803149,6 | 16485658 | 0,110 |
| LUP006823 | Dinaric-Balkan | Europe | 13265512 | 12770081,6 | 16443692 | 0,135 |
| LUP006824 | Dinaric-Balkan | Europe | 13278581 | 12784137,7 | 16461042 | 0,134 |
| LUP006825 | Dinaric-Balkan | Europe | 13197987 | 12802477,3 | 16484785 | 0,107 |
| LUP006826 | Dinaric-Balkan | Europe | 13186974 | 12723912,9 | 16383648 | 0,127 |
| LUP006827 | Dinaric-Balkan | Europe | 13311527 | 12792195,8 | 16471940 | 0,141 |
| LUP006828 | Dinaric-Balkan | Europe | 13460908 | 12786868,5 | 16464626 | 0,183 |
| MSV2AC | Dinaric-Balkan | Europe | 13261671 | 12809414,4 | 16493795 | 0,123 |
| W1023 | Italian Peninsula | Europe | 14220785 | 12801930,9 | 16484095 | 0,385 |
| W1551 | Italian Peninsula | Europe | 12325058 | 11082011,9 | 14270799 | 0,390 |
| W1593 | Italian Peninsula | Europe | 13646660 | 12608323,1 | 16235527 | 0,286 |
| W2030 | Italian Peninsula | Europe | 14042990 | 12763253 | 16434969 | 0,349 |

**Table S13.** Individual-level heterozygosity and inbreeding coefficients for European wolf samples.

| Sample ID | Population | Continent | Observed homozygosity (SN | Expected homozygosity (SN | Total numl | *F* (Inbreeding coefficient) |
|---|---|---|---|---|---|---|
| W2124 | Italian Peninsula | Europe | 14137295 | 12773126,2 | 16447032 | 0,371 |
| W2185 | Italian Peninsula | Europe | 14283513 | 12769352 | 16443051 | 0,412 |
| W2193 | Italian Peninsula | Europe | 14077979 | 12798068,7 | 16479481 | 0,348 |
| W2265 | Italian Peninsula | Europe | 14217071 | 12787655,7 | 16466288 | 0,389 |
| W2280 | Italian Peninsula | Europe | 14013495 | 12807287,7 | 16491068 | 0,327 |
| W2291 | Italian Peninsula | Europe | 13901585 | 12780290,7 | 16456888 | 0,305 |
| W2371F | Italian Peninsula | Europe | 12535974 | 11175892,3 | 14382238 | 0,424 |
| W2430F | Italian Peninsula | Europe | 12043258 | 10890609,3 | 14018295 | 0,369 |
| W2487F | Italian Peninsula | Europe | 14206647 | 12558468,9 | 16170147 | 0,456 |
| W2594M | Italian Peninsula | Europe | 13954685 | 12744140 | 16410419 | 0,330 |
| W2623M | Italian Peninsula | Europe | 14045318 | 12782714,3 | 16459903 | 0,343 |
| W2707M | Italian Peninsula | Europe | 13630549 | 12373696,8 | 15932178 | 0,353 |
| W2878M | Italian Peninsula | Europe | 14116396 | 12756215,9 | 16425758 | 0,371 |
| W2883F | Italian Peninsula | Europe | 14062740 | 12747053,7 | 16413678 | 0,359 |
| W2886 | Italian Peninsula | Europe | 13718438 | 12680876 | 16327000 | 0,285 |
| W2910 | Italian Peninsula | Europe | 14272480 | 12803700,9 | 16486316 | 0,399 |
| W2951 | Italian Peninsula | Europe | 14101751 | 12804351,1 | 16487166 | 0,352 |
| w361703 | Italian Peninsula | Europe | 14355721 | 12806223,7 | 16489655 | 0,421 |
| w361756 | Italian Peninsula | Europe | 14010973 | 12807957,2 | 16491787 | 0,327 |
| w4926402 | Italian Peninsula | Europe | 13889289 | 12800950,1 | 16482763 | 0,296 |
| W893 | Italian Peninsula | Europe | 14323244 | 12791139,8 | 16470090 | 0,416 |
| WIT11 | Italian Peninsula | Europe | 14096507 | 12691652 | 16343816 | 0,385 |
| WIT13 | Italian Peninsula | Europe | 13800882 | 12286177,9 | 15822792 | 0,428 |
| WIT14 | Italian Peninsula | Europe | 14032080 | 12709135,6 | 16365892 | 0,362 |
| WIT16 | Italian Peninsula | Europe | 14240096 | 12620456,8 | 16252061 | 0,446 |
| WIT17 | Italian Peninsula | Europe | 13613248 | 12377660,2 | 15941905 | 0,347 |
| WIT19 | Italian Peninsula | Europe | 13706462 | 12488490,3 | 16083643 | 0,339 |
| WIT20 | Italian Peninsula | Europe | 14407563 | 12708354 | 16364520 | 0,465 |
| WIT21 | Italian Peninsula | Europe | 13951543 | 12584151,2 | 16206573 | 0,377 |
| WIT22 | Italian Peninsula | Europe | 13544142 | 12318155,6 | 15864974 | 0,346 |
| WIT23 | Italian Peninsula | Europe | 13958590 | 12701521,1 | 16356056 | 0,344 |
| CLUPRU000005 | Karelian | Europe | 13402967 | 12596827 | 16219511 | 0,223 |
| CLUPRU000006 | Karelian | Europe | 13000230 | 12653856,8 | 16292434 | 0,095 |

**Table S13.** Individual-level heterozygosity and inbreeding coefficients for European wolf samples.

| Sample ID | Population | Continent | Observed homozygosity (SN | Expected homozygosity (SN | Total numl | *F* (Inbreeding coefficient) |
|---|---|---|---|---|---|---|
| CLUPRU000007 | Karelian | Europe | 12973540 | 12657005 | 16296204 | 0,087 |
| CLUPRU000008 | Karelian | Europe | 12916841 | 12638033,2 | 16272310 | 0,077 |
| CLUPRU000019 | Karelian | Europe | 12569506 | 12243487,6 | 15764146 | 0,093 |
| CLUPRU000020 | Karelian | Europe | 12179799 | 11789288,2 | 15176466 | 0,115 |
| V113 | Karelian | Europe | 13370418 | 12810486,1 | 16495146 | 0,152 |
| V114 | Karelian | Europe | 13060991 | 12810926 | 16495757 | 0,068 |
| V115 | Karelian | Europe | 13034556 | 12810932,1 | 16495752 | 0,061 |
| V116 | Karelian | Europe | 13109791 | 12811117,8 | 16495991 | 0,081 |
| V117 | Karelian | Europe | 13180323 | 12809121,3 | 16493496 | 0,101 |
| V119 | Karelian | Europe | 13132868 | 12810715,4 | 16495446 | 0,087 |
| V120 | Karelian | Europe | 13089298 | 12811060,6 | 16495902 | 0,076 |
| V126 | Karelian | Europe | 13071392 | 12810928,7 | 16495744 | 0,071 |
| V132 | Karelian | Europe | 13156087 | 12811215,3 | 16496123 | 0,094 |
| V134 | Karelian | Europe | 13107934 | 12810662,7 | 16495391 | 0,081 |
| V136 | Karelian | Europe | 13218762 | 12808250,3 | 16492194 | 0,111 |
| V141 | Karelian | Europe | 13144134 | 12810937,4 | 16495752 | 0,090 |
| V143 | Karelian | Europe | 13209819 | 12810570 | 16495252 | 0,108 |
| W11 | Karelian | Europe | 13141328 | 12798779 | 16480258 | 0,093 |
| W14 | Karelian | Europe | 13236232 | 12809842,9 | 16494328 | 0,116 |
| W16 | Karelian | Europe | 13745173 | 12806838,8 | 16490464 | 0,255 |
| W21 | Karelian | Europe | 13186171 | 12809846,3 | 16494380 | 0,102 |
| W26 | Karelian | Europe | 13283349 | 12810674,6 | 16495427 | 0,128 |
| W29 | Karelian | Europe | 13366431 | 12808524,1 | 16492660 | 0,151 |
| W32 | Karelian | Europe | 13097553 | 12807434,5 | 16491183 | 0,079 |
| W4 | Karelian | Europe | 13704592 | 12807724,6 | 16491532 | 0,243 |
| W44 | Karelian | Europe | 13044184 | 12806795,2 | 16490527 | 0,064 |
| W45 | Karelian | Europe | 13142582 | 12810492,2 | 16495167 | 0,090 |
| JAL7480 | NW Iberia | Europe | 14743830 | 12775235,8 | 16449230 | 0,536 |
| JAL7486 | NW Iberia | Europe | 14279001 | 12804477,9 | 16487373 | 0,400 |
| JAL7491 | NW Iberia | Europe | 13598728 | 12807940,4 | 16491924 | 0,215 |
| JAL7492 | NW Iberia | Europe | 14309174 | 12803389,6 | 16485959 | 0,409 |
| L166 | NW Iberia | Europe | 13922247 | 12804599 | 16487546 | 0,303 |
| L253 | NW Iberia | Europe | 13639535 | 12715811,7 | 16373149 | 0,253 |

**Table S13.** Individual-level heterozygosity and inbreeding coefficients for European wolf samples.

| Sample ID | Population | Continent | Observed homozygosity (SN | Expected homozygosity (SN | Total numl | *F* (Inbreeding coefficient) |
|---|---|---|---|---|---|---|
| L409 | NW Iberia | Europe | 13787979 | 12743315,4 | 16407483 | 0,285 |
| L514 | NW Iberia | Europe | 13576098 | 12713753,3 | 16370413 | 0,236 |
| L547 | NW Iberia | Europe | 14041540 | 12741258,6 | 16404937 | 0,355 |
| L552 | NW Iberia | Europe | 13703988 | 12730069,7 | 16390580 | 0,266 |
| L588 | NW Iberia | Europe | 14086065 | 12741794,6 | 16406754 | 0,367 |
| L844 | NW Iberia | Europe | 14694354 | 12724048 | 16382152 | 0,539 |
| LUP006829 | NW Iberia | Europe | 14446007 | 12790721,4 | 16469583 | 0,450 |
| LUP006830 | NW Iberia | Europe | 13559684 | 12779378,5 | 16455701 | 0,212 |
| MW122 | NW Iberia | Europe | 13860348 | 12761608,6 | 16431669 | 0,299 |
| MW127 | NW Iberia | Europe | 14120921 | 12753324,3 | 16421038 | 0,373 |
| Penelope | NW Iberia | Europe | 13643003 | 12792904,8 | 16473012 | 0,231 |
| ptw | NW Iberia | Europe | 14119361 | 12708041,9 | 16363624 | 0,386 |
| D-00-12 | Scandinavia | Europe | 14049560 | 12768351,4 | 16441310 | 0,349 |
| D-06-14 | Scandinavia | Europe | 14072051 | 12800839 | 16482803 | 0,345 |
| D-07-16 | Scandinavia | Europe | 14080770 | 12208840,4 | 15713291 | 0,534 |
| D-08-10 | Scandinavia | Europe | 14553953 | 12610317,4 | 16236842 | 0,536 |
| D-10-68 | Scandinavia | Europe | 13879891 | 12614072,1 | 16243821 | 0,349 |
| D-11-58 | Scandinavia | Europe | 13134093 | 11956992,9 | 15391113 | 0,343 |
| D-85-02 | Scandinavia | Europe | 13096125 | 12808493,4 | 16492652 | 0,078 |
| G100-12 | Scandinavia | Europe | 13213210 | 12498623,9 | 16095097 | 0,199 |
| G100-14 | Scandinavia | Europe | 13777399 | 12808170 | 16492109 | 0,263 |
| G109-11 | Scandinavia | Europe | 13367086 | 12533877,4 | 16139838 | 0,231 |
| G126-13 | Scandinavia | Europe | 13552073 | 12303513,6 | 15844106 | 0,353 |
| G139-12 | Scandinavia | Europe | 13283745 | 12400427,6 | 15964873 | 0,248 |
| G31-13 | Scandinavia | Europe | 13443948 | 12577763,6 | 16194534 | 0,239 |
| G37-10 | Scandinavia | Europe | 13023646 | 12579873,3 | 16199127 | 0,123 |
| G50-12 | Scandinavia | Europe | 13998749 | 12398102,8 | 15965330 | 0,449 |
| M-01-06 | Scandinavia | Europe | 14309667 | 12802834,7 | 16485174 | 0,409 |
| M-06-03 | Scandinavia | Europe | 14245046 | 12786644,8 | 16464752 | 0,397 |
| M-09-05 | Scandinavia | Europe | 13745511 | 12489110,9 | 16082827 | 0,350 |
| M-10-04 | Scandinavia | Europe | 13384945 | 12781681,3 | 16457650 | 0,164 |
| M-98-08 | Scandinavia | Europe | 13495771 | 12809745,7 | 16494111 | 0,186 |
| CLUPKG000001 | Central Asia | Asia | 12821068 | 12303769 | 15839076 | 0,146 |

**Table S13.** Individual-level heterozygosity and inbreeding coefficients for European wolf samples.

| Sample ID | Population | Continent | Observed homozygosity (SN | Expected homozygosity (SN | Total numl | *F* (Inbreeding coefficient) |
|---|---|---|---|---|---|---|
| CLUPKZ000002 | Central Asia | Asia | 13565043 | 12710282,7 | 16365638 | 0,234 |
| CLUPRU000001 | Central Asia | Asia | 12936836 | 12781772,2 | 16458238 | 0,042 |
| CLUPRU000002 | Central Asia | Asia | 13041758 | 12774198,1 | 16448600 | 0,073 |
| CLUPRU000003 | Central Asia | Asia | 13222645 | 12796016,2 | 16476231 | 0,116 |
| CLUPTJ000001 | Central Asia | Asia | 12288174 | 12013553,1 | 15467094 | 0,080 |
| CLUPTJ000002 | Central Asia | Asia | 11777269 | 11488325,4 | 14788545 | 0,088 |
| CLUPTJ000003 | Central Asia | Asia | 13041853 | 12724711,6 | 16384589 | 0,087 |
| CLUPTJ000004 | Central Asia | Asia | 12915608 | 12483138,6 | 16071331 | 0,121 |
| CLUPTJ000005 | Central Asia | Asia | 12788058 | 12589354,4 | 16209398 | 0,055 |
| CLUPTJ000006 | Central Asia | Asia | 13381255 | 12686553,1 | 16335621 | 0,190 |
| CLUPTJ000007 | Central Asia | Asia | 12476823 | 12371788,4 | 15926384 | 0,030 |
| BGI-01505070006 | East Asia | Asia | 12856344 | 12681792,1 | 16330643 | 0,048 |
| BGI-01505070007 | East Asia | Asia | 12875659 | 12661717,8 | 16304600 | 0,059 |
| CLUPCN000001 | East Asia | Asia | 13104304 | 12789620,3 | 16468003 | 0,086 |
| CLUPCN000005 | East Asia | Asia | 13038546 | 12790987,5 | 16470129 | 0,067 |
| CLUPCN000006 | East Asia | Asia | 13006755 | 12791942,5 | 16471341 | 0,058 |
| CLUPCN000007 | East Asia | Asia | 12961410 | 12799604,9 | 16481013 | 0,044 |
| CLUPCN000008 | East Asia | Asia | 12698282 | 12784014,3 | 16461223 | -0,023 |
| CLUPCN000009 | East Asia | Asia | 12890299 | 12794692,4 | 16474512 | 0,026 |
| CLUPCN000010 | East Asia | Asia | 12867449 | 12795689,4 | 16475887 | 0,020 |
| CLUPRU000004 | East Asia | Asia | 13106228 | 12797818,7 | 16478609 | 0,084 |
| LUPWCHN00010 | East Asia | Asia | 12646170 | 12303696,9 | 15846800 | 0,097 |
| LUPZCHN00009 | East Asia | Asia | 12702450 | 12440487,1 | 16021017 | 0,073 |
| RKW13451 | East Asia | Asia | 13871890 | 12769646,5 | 16442775 | 0,300 |
| BGI-01505070001 | Tibetan | Asia | 14458800 | 12587929,1 | 16209646 | 0,517 |
| BGI-01505070002 | Tibetan | Asia | 13650485 | 12599659,7 | 16224899 | 0,290 |
| BGI-01505070003 | Tibetan | Asia | 13440622 | 12678477,9 | 16326113 | 0,209 |
| BGI-01505070004 | Tibetan | Asia | 12959366 | 12487977,6 | 16082932 | 0,131 |
| BGI-01505070005 | Tibetan | Asia | 13360782 | 12623824,7 | 16256475 | 0,203 |
| BGI-01505070008 | Tibetan | Asia | 14812697 | 12639708,3 | 16276097 | 0,598 |
| CLUPCN000003 | Tibetan | Asia | 13540879 | 12765477,7 | 16436598 | 0,211 |
| CLUPAZ000001 | West Asia | Asia | 13055678 | 12793179,4 | 16472948 | 0,071 |
| CLUPIR000001 | West Asia | Asia | 13117718 | 12764159,1 | 16435748 | 0,096 |

**Table S13.** Individual-level heterozygosity and inbreeding coefficients for European wolf samples.

| Sample ID | Population | Continent | Observed homozygosity (SN | Expected homozygosity (SN | Total numl | *F* (Inbreeding coefficient) |
|---|---|---|---|---|---|---|
| CLUPIR000003 | West Asia | Asia | 12977914 | 12747883 | 16414541 | 0,063 |
| CLUPIR000004 | West Asia | Asia | 12922718 | 12793335,8 | 16473047 | 0,035 |
| CLUPIR000005 | West Asia | Asia | 13008965 | 12789087,3 | 16467546 | 0,060 |
| irw | West Asia | Asia | 12927754 | 12703232,1 | 16357704 | 0,061 |
| WTR026 | West Asia | Asia | 13072755 | 12806853,2 | 16490476 | 0,072 |
| WTR028 | West Asia | Asia | 12999659 | 12781802,1 | 16458690 | 0,059 |
| WTR029 | West Asia | Asia | 13002670 | 12808296,4 | 16492274 | 0,053 |
| WTR030 | West Asia | Asia | 13016425 | 12808441,3 | 16492546 | 0,056 |
| WTR031 | West Asia | Asia | 13205016 | 12807973,9 | 16491971 | 0,108 |
| WTR032 | West Asia | Asia | 13024780 | 12803944,1 | 16486826 | 0,060 |
| WTR033 | West Asia | Asia | 13019119 | 12806921,2 | 16490446 | 0,058 |
| WTR034 | West Asia | Asia | 13331684 | 12808625,9 | 16492800 | 0,142 |
| WTR035 | West Asia | Asia | 12982552 | 12809301,5 | 16493659 | 0,047 |
| WTR037 | West Asia | Asia | 13068095 | 12809222,3 | 16493572 | 0,070 |
| WTR038 | West Asia | Asia | 13010770 | 12805737,4 | 16488941 | 0,056 |
| WTR040 | West Asia | Asia | 13031331 | 12803468,5 | 16486330 | 0,062 |
| WTR041 | West Asia | Asia | 13124203 | 12806773 | 16490521 | 0,086 |

| Observed heterozygosity (SN | Proportion of heterozygous sites |
|---|---|
| 2878513 | 0,175 |
| 3175561 | 0,193 |
| 3128709 | 0,190 |
| 2922506 | 0,177 |
| 3055836 | 0,185 |
| 2458375 | 0,149 |
| 3225169 | 0,196 |
| 2354531 | 0,143 |
| 2916696 | 0,177 |
| 3321970 | 0,201 |
| 3077821 | 0,187 |
| 3381249 | 0,205 |
| 3100543 | 0,188 |
| 2885536 | 0,175 |
| 3229128 | 0,196 |
| 3156275 | 0,192 |
| 3233033 | 0,197 |
| 3115352 | 0,189 |
| 3291294 | 0,200 |
| 3158121 | 0,192 |
| 3233235 | 0,196 |
| 3277992 | 0,199 |
| 3178180 | 0,193 |
| 3182461 | 0,193 |
| 3286798 | 0,199 |
| 3196674 | 0,195 |
| 3160413 | 0,192 |
| 3003718 | 0,182 |
| 3232124 | 0,196 |
| 2263310 | 0,137 |
| 1945741 | 0,136 |
| 2588867 | 0,159 |
| 2391979 | 0,146 |

| Observed heterozygosity (SN | Proportion of heterozygous sites |
|---|---|
| 2309737 | 0,140 |
| 2159538 | 0,131 |
| 2401502 | 0,146 |
| 2249217 | 0,137 |
| 2477573 | 0,150 |
| 2555303 | 0,155 |
| 1846264 | 0,128 |
| 1975037 | 0,141 |
| 1963500 | 0,121 |
| 2455734 | 0,150 |
| 2414585 | 0,147 |
| 2301629 | 0,144 |
| 2309362 | 0,141 |
| 2350938 | 0,143 |
| 2608562 | 0,160 |
| 2213836 | 0,134 |
| 2385415 | 0,145 |
| 2133934 | 0,129 |
| 2480814 | 0,150 |
| 2593474 | 0,157 |
| 2146846 | 0,130 |
| 2247309 | 0,138 |
| 2021910 | 0,128 |
| 2333812 | 0,143 |
| 2011965 | 0,124 |
| 2328657 | 0,146 |
| 2377181 | 0,148 |
| 1956957 | 0,120 |
| 2255030 | 0,139 |
| 2320832 | 0,146 |
| 2397466 | 0,147 |
| 2816544 | 0,174 |
| 3292204 | 0,202 |

| Observed heterozygosity (SN | Proportion of heterozygous sites |
|---|---|
| 3322664 | 0,204 |
| 3355469 | 0,206 |
| 3194640 | 0,203 |
| 2996667 | 0,197 |
| 3124728 | 0,189 |
| 3434766 | 0,208 |
| 3461196 | 0,210 |
| 3386200 | 0,205 |
| 3313173 | 0,201 |
| 3362578 | 0,204 |
| 3406604 | 0,207 |
| 3424352 | 0,208 |
| 3340036 | 0,202 |
| 3387457 | 0,205 |
| 3273432 | 0,198 |
| 3351618 | 0,203 |
| 3285433 | 0,199 |
| 3338930 | 0,203 |
| 3258096 | 0,198 |
| 2745291 | 0,166 |
| 3308209 | 0,201 |
| 3212078 | 0,195 |
| 3126229 | 0,190 |
| 3393630 | 0,206 |
| 2786940 | 0,169 |
| 3446343 | 0,209 |
| 3352585 | 0,203 |
| 1705400 | 0,104 |
| 2208372 | 0,134 |
| 2893196 | 0,175 |
| 2176785 | 0,132 |
| 2565299 | 0,156 |
| 2733614 | 0,167 |

| Observed heterozygosity (SN | Proportion of heterozygous sites |
|---|---|
| 2619504 | 0,160 |
| 2794315 | 0,171 |
| 2363397 | 0,144 |
| 2686592 | 0,164 |
| 2320689 | 0,141 |
| 1687798 | 0,103 |
| 2023576 | 0,123 |
| 2896017 | 0,176 |
| 2571321 | 0,156 |
| 2300117 | 0,140 |
| 2830009 | 0,172 |
| 2244263 | 0,137 |
| 2391750 | 0,145 |
| 2410752 | 0,146 |
| 1632521 | 0,104 |
| 1682889 | 0,104 |
| 2363930 | 0,146 |
| 2257020 | 0,147 |
| 3396527 | 0,206 |
| 2881887 | 0,179 |
| 2714710 | 0,165 |
| 2772752 | 0,172 |
| 2292033 | 0,145 |
| 2681128 | 0,168 |
| 2750586 | 0,170 |
| 3175481 | 0,196 |
| 1966581 | 0,123 |
| 2175507 | 0,132 |
| 2219706 | 0,135 |
| 2337316 | 0,145 |
| 3072705 | 0,187 |
| 2998340 | 0,182 |
| 3018008 | 0,191 |

| Observed heterozygosity (SN | Proportion of heterozygous sites |
|---|---|
| 2800595 | 0,171 |
| 3521402 | 0,214 |
| 3406842 | 0,207 |
| 3253586 | 0,197 |
| 3178920 | 0,206 |
| 3011276 | 0,204 |
| 3342736 | 0,204 |
| 3155723 | 0,196 |
| 3421340 | 0,211 |
| 2954366 | 0,181 |
| 3449561 | 0,217 |
| 3474299 | 0,213 |
| 3428941 | 0,210 |
| 3363699 | 0,204 |
| 3431583 | 0,208 |
| 3464586 | 0,210 |
| 3519603 | 0,214 |
| 3762941 | 0,229 |
| 3584213 | 0,218 |
| 3608438 | 0,219 |
| 3372381 | 0,205 |
| 3200630 | 0,202 |
| 3318567 | 0,207 |
| 2570885 | 0,156 |
| 1750846 | 0,108 |
| 2574414 | 0,159 |
| 2885491 | 0,177 |
| 3123566 | 0,194 |
| 2895693 | 0,178 |
| 1463400 | 0,090 |
| 2895719 | 0,176 |
| 3417270 | 0,207 |
| 3318030 | 0,202 |

| Observed heterozygosity (SN | Proportion of heterozygous sites |
| --- | --- |
| 3436627 | 0,209 |
| 3550329 | 0,216 |
| 3458581 | 0,210 |
| 3429950 | 0,210 |
| 3417721 | 0,207 |
| 3459031 | 0,210 |
| 3489604 | 0,212 |
| 3476121 | 0,211 |
| 3286955 | 0,199 |
| 3462046 | 0,210 |
| 3471327 | 0,211 |
| 3161116 | 0,192 |
| 3511107 | 0,213 |
| 3425477 | 0,208 |
| 3478171 | 0,211 |
| 3454999 | 0,210 |
| 3366318 | 0,204 |

**Table S14.** Population estimates inferred by GONE2 assuming population substructure (-x) and averaged across 20 independent replicat

| Population | Number of inferred subpopulations | Between subpopulations Fst | Between subpopulations migration rate |
|---|---|---|---|
| Dinaric-Balkan | 3 | 0,063 | 0,022 |
| Iberian | 9 | 0,137 | 0,059 |
| Italian_Peninsula | 6 | 0,034 | 0,45 |
| Karelian | 7 | 0,03 | 0,161 |
| Scandinavian | 2 | 0,029 | 0,241 |

tes.

| Contemporary Ne (generation 1) |
| --- |
| 238,24 |
| 216,01 |
| 105,9 |
| 248,09 |
| 17,64 |

**Table S15.** Sum of runs of homozygosity (SROH) per individual based on ROH length.

| **Individual** | **Population** | **ROH length** | **SROH (Gb)** |
|---|---|---|---|
| AP_085T | Dinaric-Balkan | 1Mb-2Mb | 0,06234 |
| AP_085T | Dinaric-Balkan | >2Mb | 0,33886 |
| AP_085T | Dinaric-Balkan | 100kb-1Mb | 0,16861 |
| AP_086E | Dinaric-Balkan | 1Mb-2Mb | 0,04803 |
| AP_086E | Dinaric-Balkan | >2Mb | 0,14447 |
| AP_086E | Dinaric-Balkan | 100kb-1Mb | 0,18366 |
| AP_087A | Dinaric-Balkan | 1Mb-2Mb | 0,07266 |
| AP_087A | Dinaric-Balkan | >2Mb | 0,27351 |
| AP_087A | Dinaric-Balkan | 100kb-1Mb | 0,17768 |
| AP_087E | Dinaric-Balkan | 1Mb-2Mb | 0,04893 |
| AP_087E | Dinaric-Balkan | >2Mb | 0,25344 |
| AP_087E | Dinaric-Balkan | 100kb-1Mb | 0,16625 |
| AP_087L | Dinaric-Balkan | 1Mb-2Mb | 0,05882 |
| AP_087L | Dinaric-Balkan | >2Mb | 0,58233 |
| AP_087L | Dinaric-Balkan | 100kb-1Mb | 0,14924 |
| AP_087U | Dinaric-Balkan | 1Mb-2Mb | 0,03494 |
| AP_087U | Dinaric-Balkan | >2Mb | 0,15538 |
| AP_087U | Dinaric-Balkan | 100kb-1Mb | 0,16807 |
| AP_0882 | Dinaric-Balkan | 1Mb-2Mb | 0,10750 |
| AP_0882 | Dinaric-Balkan | >2Mb | 0,55232 |
| AP_0882 | Dinaric-Balkan | 100kb-1Mb | 0,17942 |
| AP_088L | Dinaric-Balkan | 1Mb-2Mb | 0,06525 |
| AP_088L | Dinaric-Balkan | >2Mb | 0,26679 |
| AP_088L | Dinaric-Balkan | 100kb-1Mb | 0,17615 |
| AP_08A1 | Dinaric-Balkan | 1Mb-2Mb | 0,05351 |
| AP_08A1 | Dinaric-Balkan | >2Mb | 0,05758 |
| AP_08A1 | Dinaric-Balkan | 100kb-1Mb | 0,18404 |
| AP_08A3 | Dinaric-Balkan | 1Mb-2Mb | 0,04121 |
| AP_08A3 | Dinaric-Balkan | >2Mb | 0,20857 |
| AP_08A3 | Dinaric-Balkan | 100kb-1Mb | 0,17340 |

| | | | |
|---|---|---|---|
| AP_08A6 | Dinaric-Balkan | 1Mb-2Mb | 0,04703 |
| AP_08A6 | Dinaric-Balkan | >2Mb | 0,04080 |
| AP_08A6 | Dinaric-Balkan | 100kb-1Mb | 0,17201 |
| AP_08C3 | Dinaric-Balkan | 1Mb-2Mb | 0,06290 |
| AP_08C3 | Dinaric-Balkan | >2Mb | 0,17735 |
| AP_08C3 | Dinaric-Balkan | 100kb-1Mb | 0,18190 |
| BiH_244 | Dinaric-Balkan | 1Mb-2Mb | 0,04209 |
| BiH_244 | Dinaric-Balkan | >2Mb | 0,12756 |
| BiH_244 | Dinaric-Balkan | 100kb-1Mb | 0,16706 |
| BiH_245 | Dinaric-Balkan | 1Mb-2Mb | 0,05151 |
| BiH_245 | Dinaric-Balkan | >2Mb | 0,14495 |
| BiH_245 | Dinaric-Balkan | 100kb-1Mb | 0,17493 |
| LUP006817 | Dinaric-Balkan | 1Mb-2Mb | 0,05746 |
| LUP006817 | Dinaric-Balkan | >2Mb | 0,05158 |
| LUP006817 | Dinaric-Balkan | 100kb-1Mb | 0,19911 |
| LUP006818 | Dinaric-Balkan | 1Mb-2Mb | 0,06472 |
| LUP006818 | Dinaric-Balkan | >2Mb | 0,10887 |
| LUP006818 | Dinaric-Balkan | 100kb-1Mb | 0,20648 |
| LUP006819 | Dinaric-Balkan | 1Mb-2Mb | 0,04316 |
| LUP006819 | Dinaric-Balkan | >2Mb | 0,08781 |
| LUP006819 | Dinaric-Balkan | 100kb-1Mb | 0,18360 |
| LUP006820 | Dinaric-Balkan | 1Mb-2Mb | 0,04283 |
| LUP006820 | Dinaric-Balkan | >2Mb | 0,14110 |
| LUP006820 | Dinaric-Balkan | 100kb-1Mb | 0,18453 |
| LUP006821 | Dinaric-Balkan | 1Mb-2Mb | 0,07197 |
| LUP006821 | Dinaric-Balkan | >2Mb | 0,08617 |
| LUP006821 | Dinaric-Balkan | 100kb-1Mb | 0,19013 |
| LUP006822 | Dinaric-Balkan | 1Mb-2Mb | 0,06090 |
| LUP006822 | Dinaric-Balkan | >2Mb | 0,05058 |
| LUP006822 | Dinaric-Balkan | 100kb-1Mb | 0,19259 |
| LUP006823 | Dinaric-Balkan | 1Mb-2Mb | 0,05556 |

| | | | |
|---|---|---|---|
| LUP006823 | Dinaric-Balkan | >2Mb | 0,09195 |
| LUP006823 | Dinaric-Balkan | 100kb-1Mb | 0,20358 |
| LUP006824 | Dinaric-Balkan | 1Mb-2Mb | 0,07578 |
| LUP006824 | Dinaric-Balkan | >2Mb | 0,08943 |
| LUP006824 | Dinaric-Balkan | 100kb-1Mb | 0,21648 |
| LUP006825 | Dinaric-Balkan | 1Mb-2Mb | 0,04218 |
| LUP006825 | Dinaric-Balkan | >2Mb | 0,05564 |
| LUP006825 | Dinaric-Balkan | 100kb-1Mb | 0,19860 |
| LUP006826 | Dinaric-Balkan | 1Mb-2Mb | 0,05963 |
| LUP006826 | Dinaric-Balkan | >2Mb | 0,05055 |
| LUP006826 | Dinaric-Balkan | 100kb-1Mb | 0,22044 |
| LUP006827 | Dinaric-Balkan | 1Mb-2Mb | 0,08348 |
| LUP006827 | Dinaric-Balkan | >2Mb | 0,10056 |
| LUP006827 | Dinaric-Balkan | 100kb-1Mb | 0,19581 |
| LUP006828 | Dinaric-Balkan | 1Mb-2Mb | 0,08307 |
| LUP006828 | Dinaric-Balkan | >2Mb | 0,13074 |
| LUP006828 | Dinaric-Balkan | 100kb-1Mb | 0,23738 |
| MSV2AC | Dinaric-Balkan | 1Mb-2Mb | 0,04238 |
| MSV2AC | Dinaric-Balkan | >2Mb | 0,15202 |
| MSV2AC | Dinaric-Balkan | 100kb-1Mb | 0,17565 |
| JAL7480 | NW Iberia | 1Mb-2Mb | 0,11190 |
| JAL7480 | NW Iberia | >2Mb | 0,94658 |
| JAL7480 | NW Iberia | 100kb-1Mb | 0,15922 |
| JAL7486 | NW Iberia | 1Mb-2Mb | 0,10240 |
| JAL7486 | NW Iberia | >2Mb | 0,60916 |
| JAL7486 | NW Iberia | 100kb-1Mb | 0,18351 |
| JAL7491 | NW Iberia | 1Mb-2Mb | 0,07302 |
| JAL7491 | NW Iberia | >2Mb | 0,23992 |
| JAL7491 | NW Iberia | 100kb-1Mb | 0,21804 |
| JAL7492 | NW Iberia | 1Mb-2Mb | 0,06854 |
| JAL7492 | NW Iberia | >2Mb | 0,71486 |

| | | | |
|---|---|---|---|
| JAL7492 | NW Iberia | 100kb-1Mb | 0,18748 |
| L166 | NW Iberia | 1Mb-2Mb | 0,09714 |
| L166 | NW Iberia | >2Mb | 0,41620 |
| L166 | NW Iberia | 100kb-1Mb | 0,22022 |
| L253 | NW Iberia | 1Mb-2Mb | 0,10973 |
| L253 | NW Iberia | >2Mb | 0,20171 |
| L253 | NW Iberia | 100kb-1Mb | 0,27228 |
| L409 | NW Iberia | 1Mb-2Mb | 0,11016 |
| L409 | NW Iberia | >2Mb | 0,29167 |
| L409 | NW Iberia | 100kb-1Mb | 0,25846 |
| L514 | NW Iberia | 1Mb-2Mb | 0,10552 |
| L514 | NW Iberia | >2Mb | 0,18516 |
| L514 | NW Iberia | 100kb-1Mb | 0,25766 |
| L547 | NW Iberia | 1Mb-2Mb | 0,12309 |
| L547 | NW Iberia | >2Mb | 0,44814 |
| L547 | NW Iberia | 100kb-1Mb | 0,24068 |
| L552 | NW Iberia | 1Mb-2Mb | 0,11616 |
| L552 | NW Iberia | >2Mb | 0,17572 |
| L552 | NW Iberia | 100kb-1Mb | 0,28882 |
| L588 | NW Iberia | 1Mb-2Mb | 0,13686 |
| L588 | NW Iberia | >2Mb | 0,46877 |
| L588 | NW Iberia | 100kb-1Mb | 0,24169 |
| LUP006829 | NW Iberia | 1Mb-2Mb | 0,21757 |
| LUP006829 | NW Iberia | >2Mb | 0,45629 |
| LUP006829 | NW Iberia | 100kb-1Mb | 0,33171 |
| LUP006830 | NW Iberia | 1Mb-2Mb | 0,09531 |
| LUP006830 | NW Iberia | >2Mb | 0,12871 |
| LUP006830 | NW Iberia | 100kb-1Mb | 0,30012 |
| MW122 | NW Iberia | 1Mb-2Mb | 0,10788 |
| MW122 | NW Iberia | >2Mb | 0,30071 |
| MW122 | NW Iberia | 100kb-1Mb | 0,25650 |

| | | | |
|---|---|---|---|
| MW127 | NW Iberia | 1Mb-2Mb | 0,10604 |
| MW127 | NW Iberia | >2Mb | 0,52635 |
| MW127 | NW Iberia | 100kb-1Mb | 0,22814 |
| Penelope | NW Iberia | 1Mb-2Mb | 0,06845 |
| Penelope | NW Iberia | >2Mb | 0,25764 |
| Penelope | NW Iberia | 100kb-1Mb | 0,23219 |
| ptw | NW Iberia | 1Mb-2Mb | 0,08312 |
| ptw | NW Iberia | >2Mb | 0,59444 |
| ptw | NW Iberia | 100kb-1Mb | 0,20050 |
| W1023 | Italian Peninsula | 1Mb-2Mb | 0,10272 |
| W1023 | Italian Peninsula | >2Mb | 0,56406 |
| W1023 | Italian Peninsula | 100kb-1Mb | 0,24333 |
| W1551 | Italian Peninsula | 1Mb-2Mb | 0,12184 |
| W1551 | Italian Peninsula | >2Mb | 0,50337 |
| W1551 | Italian Peninsula | 100kb-1Mb | 0,26632 |
| W1593 | Italian Peninsula | 1Mb-2Mb | 0,10681 |
| W1593 | Italian Peninsula | >2Mb | 0,42429 |
| W1593 | Italian Peninsula | 100kb-1Mb | 0,18986 |
| W2030 | Italian Peninsula | 1Mb-2Mb | 0,07880 |
| W2030 | Italian Peninsula | >2Mb | 0,53547 |
| W2030 | Italian Peninsula | 100kb-1Mb | 0,21882 |
| W2124 | Italian Peninsula | 1Mb-2Mb | 0,08976 |
| W2124 | Italian Peninsula | >2Mb | 0,57369 |
| W2124 | Italian Peninsula | 100kb-1Mb | 0,20901 |
| W2185 | Italian Peninsula | 1Mb-2Mb | 0,10872 |
| W2185 | Italian Peninsula | >2Mb | 0,65717 |
| W2185 | Italian Peninsula | 100kb-1Mb | 0,21878 |
| W2193 | Italian Peninsula | 1Mb-2Mb | 0,13020 |
| W2193 | Italian Peninsula | >2Mb | 0,47868 |
| W2193 | Italian Peninsula | 100kb-1Mb | 0,23126 |
| W2265 | Italian Peninsula | 1Mb-2Mb | 0,12249 |

| | | | |
|---|---|---|---|
| W2265 | Italian Peninsula | >2Mb | 0,57814 |
| W2265 | Italian Peninsula | 100kb-1Mb | 0,22509 |
| W2280 | Italian Peninsula | 1Mb-2Mb | 0,09760 |
| W2280 | Italian Peninsula | >2Mb | 0,46085 |
| W2280 | Italian Peninsula | 100kb-1Mb | 0,22910 |
| W2291 | Italian Peninsula | 1Mb-2Mb | 0,11072 |
| W2291 | Italian Peninsula | >2Mb | 0,41305 |
| W2291 | Italian Peninsula | 100kb-1Mb | 0,22579 |
| W2371F | Italian Peninsula | 1Mb-2Mb | 0,20440 |
| W2371F | Italian Peninsula | >2Mb | 0,45827 |
| W2371F | Italian Peninsula | 100kb-1Mb | 0,28624 |
| W2430F | Italian Peninsula | 1Mb-2Mb | 0,12321 |
| W2430F | Italian Peninsula | >2Mb | 0,46771 |
| W2430F | Italian Peninsula | 100kb-1Mb | 0,23964 |
| W2487F | Italian Peninsula | 1Mb-2Mb | 0,15584 |
| W2487F | Italian Peninsula | >2Mb | 0,62533 |
| W2487F | Italian Peninsula | 100kb-1Mb | 0,24934 |
| W2594M | Italian Peninsula | 1Mb-2Mb | 0,13828 |
| W2594M | Italian Peninsula | >2Mb | 0,36727 |
| W2594M | Italian Peninsula | 100kb-1Mb | 0,28073 |
| W2623M | Italian Peninsula | 1Mb-2Mb | 0,11610 |
| W2623M | Italian Peninsula | >2Mb | 0,46728 |
| W2623M | Italian Peninsula | 100kb-1Mb | 0,23980 |
| W2707M | Italian Peninsula | 1Mb-2Mb | 0,19296 |
| W2707M | Italian Peninsula | >2Mb | 0,31195 |
| W2707M | Italian Peninsula | 100kb-1Mb | 0,30700 |
| W2878M | Italian Peninsula | 1Mb-2Mb | 0,11704 |
| W2878M | Italian Peninsula | >2Mb | 0,51450 |
| W2878M | Italian Peninsula | 100kb-1Mb | 0,25063 |
| W2883F | Italian Peninsula | 1Mb-2Mb | 0,12519 |
| W2883F | Italian Peninsula | >2Mb | 0,47317 |

| | | | |
|---|---|---|---|
| W2883F | Italian Peninsula | 100kb-1Mb | 0,26223 |
| W2886 | Italian Peninsula | 1Mb-2Mb | 0,14690 |
| W2886 | Italian Peninsula | >2Mb | 0,31238 |
| W2886 | Italian Peninsula | 100kb-1Mb | 0,24702 |
| W2910 | Italian Peninsula | 1Mb-2Mb | 0,11922 |
| W2910 | Italian Peninsula | >2Mb | 0,55907 |
| W2910 | Italian Peninsula | 100kb-1Mb | 0,25143 |
| W2951 | Italian Peninsula | 1Mb-2Mb | 0,13859 |
| W2951 | Italian Peninsula | >2Mb | 0,45094 |
| W2951 | Italian Peninsula | 100kb-1Mb | 0,25530 |
| W893 | Italian Peninsula | 1Mb-2Mb | 0,11457 |
| W893 | Italian Peninsula | >2Mb | 0,64372 |
| W893 | Italian Peninsula | 100kb-1Mb | 0,23733 |
| WIT11 | Italian Peninsula | 1Mb-2Mb | 0,23418 |
| WIT11 | Italian Peninsula | >2Mb | 0,28055 |
| WIT11 | Italian Peninsula | 100kb-1Mb | 0,40900 |
| WIT13 | Italian Peninsula | 1Mb-2Mb | 0,26952 |
| WIT13 | Italian Peninsula | >2Mb | 0,32779 |
| WIT13 | Italian Peninsula | 100kb-1Mb | 0,40039 |
| WIT14 | Italian Peninsula | 1Mb-2Mb | 0,21160 |
| WIT14 | Italian Peninsula | >2Mb | 0,24411 |
| WIT14 | Italian Peninsula | 100kb-1Mb | 0,39753 |
| WIT16 | Italian Peninsula | 1Mb-2Mb | 0,25051 |
| WIT16 | Italian Peninsula | >2Mb | 0,39309 |
| WIT16 | Italian Peninsula | 100kb-1Mb | 0,40034 |
| WIT17 | Italian Peninsula | 1Mb-2Mb | 0,21118 |
| WIT17 | Italian Peninsula | >2Mb | 0,17676 |
| WIT17 | Italian Peninsula | 100kb-1Mb | 0,42266 |
| WIT19 | Italian Peninsula | 1Mb-2Mb | 0,18825 |
| WIT19 | Italian Peninsula | >2Mb | 0,22272 |
| WIT19 | Italian Peninsula | 100kb-1Mb | 0,39987 |

| | | | |
|---|---|---|---|
| WIT20 | Italian Peninsula | 1Mb-2Mb | 0,27238 |
| WIT20 | Italian Peninsula | >2Mb | 0,40754 |
| WIT20 | Italian Peninsula | 100kb-1Mb | 0,40581 |
| WIT21 | Italian Peninsula | 1Mb-2Mb | 0,23247 |
| WIT21 | Italian Peninsula | >2Mb | 0,17904 |
| WIT21 | Italian Peninsula | 100kb-1Mb | 0,44630 |
| WIT22 | Italian Peninsula | 1Mb-2Mb | 0,20717 |
| WIT22 | Italian Peninsula | >2Mb | 0,12734 |
| WIT22 | Italian Peninsula | 100kb-1Mb | 0,45577 |
| WIT23 | Italian Peninsula | 1Mb-2Mb | 0,18233 |
| WIT23 | Italian Peninsula | >2Mb | 0,22035 |
| WIT23 | Italian Peninsula | 100kb-1Mb | 0,38041 |
| w361703 | Italian Peninsula | 1Mb-2Mb | 0,12472 |
| w361703 | Italian Peninsula | >2Mb | 0,64079 |
| w361703 | Italian Peninsula | 100kb-1Mb | 0,21028 |
| w361756 | Italian Peninsula | 1Mb-2Mb | 0,11858 |
| w361756 | Italian Peninsula | >2Mb | 0,47516 |
| w361756 | Italian Peninsula | 100kb-1Mb | 0,18379 |
| w4926402 | Italian Peninsula | 1Mb-2Mb | 0,10208 |
| w4926402 | Italian Peninsula | >2Mb | 0,41465 |
| w4926402 | Italian Peninsula | 100kb-1Mb | 0,20745 |
| CLUPRU000 | Karelian | 1Mb-2Mb | 0,08291 |
| CLUPRU000 | Karelian | >2Mb | 0,33725 |
| CLUPRU000 | Karelian | 100kb-1Mb | 0,15053 |
| CLUPRU000 | Karelian | 1Mb-2Mb | 0,04614 |
| CLUPRU000 | Karelian | >2Mb | 0,04616 |
| CLUPRU000 | Karelian | 100kb-1Mb | 0,14956 |
| CLUPRU000 | Karelian | 1Mb-2Mb | 0,03627 |
| CLUPRU000 | Karelian | >2Mb | 0,04764 |
| CLUPRU000 | Karelian | 100kb-1Mb | 0,15384 |
| CLUPRU000 | Karelian | 1Mb-2Mb | 0,04568 |

| | | | |
|---|---|---|---|
| CLUPRU000 | Karelian | >2Mb | 0,05425 |
| CLUPRU000 | Karelian | 100kb-1Mb | 0,14971 |
| CLUPRU000 | Karelian | 1Mb-2Mb | 0,05591 |
| CLUPRU000 | Karelian | >2Mb | 0,09329 |
| CLUPRU000 | Karelian | 100kb-1Mb | 0,14853 |
| V113 | Karelian | 1Mb-2Mb | 0,04724 |
| V113 | Karelian | >2Mb | 0,26697 |
| V113 | Karelian | 100kb-1Mb | 0,12134 |
| V114 | Karelian | 1Mb-2Mb | 0,03597 |
| V114 | Karelian | >2Mb | 0,06315 |
| V114 | Karelian | 100kb-1Mb | 0,12273 |
| V115 | Karelian | 1Mb-2Mb | 0,02607 |
| V115 | Karelian | >2Mb | 0,05599 |
| V115 | Karelian | 100kb-1Mb | 0,13743 |
| V116 | Karelian | 1Mb-2Mb | 0,02508 |
| V116 | Karelian | >2Mb | 0,13455 |
| V116 | Karelian | 100kb-1Mb | 0,11501 |
| V117 | Karelian | 1Mb-2Mb | 0,04450 |
| V117 | Karelian | >2Mb | 0,13362 |
| V117 | Karelian | 100kb-1Mb | 0,13214 |
| V119 | Karelian | 1Mb-2Mb | 0,04273 |
| V119 | Karelian | >2Mb | 0,11652 |
| V119 | Karelian | 100kb-1Mb | 0,13729 |
| V120 | Karelian | 1Mb-2Mb | 0,03998 |
| V120 | Karelian | >2Mb | 0,06067 |
| V120 | Karelian | 100kb-1Mb | 0,14322 |
| V126 | Karelian | 1Mb-2Mb | 0,03811 |
| V126 | Karelian | >2Mb | 0,06293 |
| V126 | Karelian | 100kb-1Mb | 0,14425 |
| V132 | Karelian | 1Mb-2Mb | 0,04111 |
| V132 | Karelian | >2Mb | 0,08692 |

| | | | |
|---|---|---|---|
| V132 | Karelian | 100kb-1Mb | 0,14691 |
| V134 | Karelian | 1Mb-2Mb | 0,02577 |
| V134 | Karelian | >2Mb | 0,13609 |
| V134 | Karelian | 100kb-1Mb | 0,12500 |
| V136 | Karelian | 1Mb-2Mb | 0,03838 |
| V136 | Karelian | >2Mb | 0,17200 |
| V136 | Karelian | 100kb-1Mb | 0,12793 |
| V141 | Karelian | 1Mb-2Mb | 0,02057 |
| V141 | Karelian | >2Mb | 0,13449 |
| V141 | Karelian | 100kb-1Mb | 0,13805 |
| V143 | Karelian | 1Mb-2Mb | 0,02900 |
| V143 | Karelian | >2Mb | 0,16831 |
| V143 | Karelian | 100kb-1Mb | 0,12757 |
| W11 | Karelian | 1Mb-2Mb | 0,03526 |
| W11 | Karelian | >2Mb | 0,11279 |
| W11 | Karelian | 100kb-1Mb | 0,13936 |
| W14 | Karelian | 1Mb-2Mb | 0,02430 |
| W14 | Karelian | >2Mb | 0,15307 |
| W14 | Karelian | 100kb-1Mb | 0,14039 |
| W16 | Karelian | 1Mb-2Mb | 0,04860 |
| W16 | Karelian | >2Mb | 0,46076 |
| W16 | Karelian | 100kb-1Mb | 0,11602 |
| W21 | Karelian | 1Mb-2Mb | 0,04315 |
| W21 | Karelian | >2Mb | 0,13567 |
| W21 | Karelian | 100kb-1Mb | 0,13343 |
| W26 | Karelian | 1Mb-2Mb | 0,04405 |
| W26 | Karelian | >2Mb | 0,22826 |
| W26 | Karelian | 100kb-1Mb | 0,11866 |
| W32 | Karelian | 1Mb-2Mb | 0,03328 |
| W32 | Karelian | >2Mb | 0,11315 |
| W32 | Karelian | 100kb-1Mb | 0,14189 |

| | | | |
|---|---|---|---|
| W4 | Karelian | 1Mb-2Mb | 0,05298 |
| W4 | Karelian | >2Mb | 0,48598 |
| W4 | Karelian | 100kb-1Mb | 0,12238 |
| W44 | Karelian | 1Mb-2Mb | 0,02597 |
| W44 | Karelian | >2Mb | 0,06492 |
| W44 | Karelian | 100kb-1Mb | 0,13411 |
| W45 | Karelian | 1Mb-2Mb | 0,04817 |
| W45 | Karelian | >2Mb | 0,11267 |
| W45 | Karelian | 100kb-1Mb | 0,12868 |
| D-00-12 | Scandinavian | 1Mb-2Mb | 0,06160 |
| D-00-12 | Scandinavian | >2Mb | 0,67606 |
| D-00-12 | Scandinavian | 100kb-1Mb | 0,10881 |
| D-06-14 | Scandinavian | 1Mb-2Mb | 0,05846 |
| D-06-14 | Scandinavian | >2Mb | 0,71720 |
| D-06-14 | Scandinavian | 100kb-1Mb | 0,10634 |
| D-07-16 | Scandinavian | 1Mb-2Mb | 0,29908 |
| D-07-16 | Scandinavian | >2Mb | 0,16150 |
| D-07-16 | Scandinavian | 100kb-1Mb | 0,77089 |
| D-08-10 | Scandinavian | 1Mb-2Mb | 0,20610 |
| D-08-10 | Scandinavian | >2Mb | 0,85836 |
| D-08-10 | Scandinavian | 100kb-1Mb | 0,17350 |
| D-10-68 | Scandinavian | 1Mb-2Mb | 0,18100 |
| D-10-68 | Scandinavian | >2Mb | 0,51697 |
| D-10-68 | Scandinavian | 100kb-1Mb | 0,16555 |
| D-11-58 | Scandinavian | 1Mb-2Mb | 0,18972 |
| D-11-58 | Scandinavian | >2Mb | 0,26316 |
| D-11-58 | Scandinavian | 100kb-1Mb | 0,31353 |
| D-85-02 | Scandinavian | 1Mb-2Mb | 0,03623 |
| D-85-02 | Scandinavian | >2Mb | 0,10442 |
| D-85-02 | Scandinavian | 100kb-1Mb | 0,14291 |
| G100-12 | Scandinavian | 1Mb-2Mb | 0,14485 |

| | | | |
|---|---|---|---|
| G100-12 | Scandinavian | >2Mb | 0,13087 |
| G100-12 | Scandinavian | 100kb-1Mb | 0,23126 |
| G100-14 | Scandinavian | 1Mb-2Mb | 0,04853 |
| G100-14 | Scandinavian | >2Mb | 0,46705 |
| G100-14 | Scandinavian | 100kb-1Mb | 0,12305 |
| G126-13 | Scandinavian | 1Mb-2Mb | 0,22122 |
| G126-13 | Scandinavian | >2Mb | 0,28628 |
| G126-13 | Scandinavian | 100kb-1Mb | 0,29937 |
| G139-12 | Scandinavian | 1Mb-2Mb | 0,13666 |
| G139-12 | Scandinavian | >2Mb | 0,16532 |
| G139-12 | Scandinavian | 100kb-1Mb | 0,26234 |
| G50-12 | Scandinavian | 1Mb-2Mb | 0,30097 |
| G50-12 | Scandinavian | >2Mb | 0,41448 |
| G50-12 | Scandinavian | 100kb-1Mb | 0,32063 |
| M-01-06 | Scandinavian | 1Mb-2Mb | 0,05962 |
| M-01-06 | Scandinavian | >2Mb | 0,84102 |
| M-01-06 | Scandinavian | 100kb-1Mb | 0,09532 |
| M-06-03 | Scandinavian | 1Mb-2Mb | 0,05582 |
| M-06-03 | Scandinavian | >2Mb | 0,83235 |
| M-06-03 | Scandinavian | 100kb-1Mb | 0,10978 |
| M-09-05 | Scandinavian | 1Mb-2Mb | 0,18604 |
| M-09-05 | Scandinavian | >2Mb | 0,37637 |
| M-09-05 | Scandinavian | 100kb-1Mb | 0,25562 |
| M-10-04 | Scandinavian | 1Mb-2Mb | 0,06699 |
| M-10-04 | Scandinavian | >2Mb | 0,25512 |
| M-10-04 | Scandinavian | 100kb-1Mb | 0,13350 |
| M-98-08 | Scandinavian | 1Mb-2Mb | 0,05016 |
| M-98-08 | Scandinavian | >2Mb | 0,33037 |
| M-98-08 | Scandinavian | 100kb-1Mb | 0,12630 |

**Table S16.** Genetic load counts per individual based on different functional cagories.

| Individual | Population | LOW Realized load | LOW Masked load | MOD_TOL Realized load | MOD_TOL Masked load |
|---|---|---|---|---|---|
| AP_085T | Dinaric-Balkan | 3868 | 2615 | 1142 | 814 |
| AP_085Y | Dinaric-Balkan | 3041 | 3878 | 950 | 1091 |
| AP_086E | Dinaric-Balkan | 3714 | 2924 | 1133 | 851 |
| AP_087A | Dinaric-Balkan | 3840 | 2750 | 1136 | 833 |
| AP_087E | Dinaric-Balkan | 3813 | 2770 | 1111 | 818 |
| AP_087L | Dinaric-Balkan | 3950 | 2498 | 1153 | 758 |
| AP_087U | Dinaric-Balkan | 3749 | 2906 | 1095 | 877 |
| AP_0882 | Dinaric-Balkan | 4070 | 2156 | 1198 | 670 |
| AP_088L | Dinaric-Balkan | 3832 | 2694 | 1163 | 831 |
| AP_08A1 | Dinaric-Balkan | 3644 | 3086 | 1065 | 924 |
| AP_08A3 | Dinaric-Balkan | 3728 | 2871 | 1101 | 848 |
| AP_08A6 | Dinaric-Balkan | 3648 | 3038 | 1042 | 938 |
| AP_08C3 | Dinaric-Balkan | 3752 | 2827 | 1083 | 874 |
| BiH_244 | Dinaric-Balkan | 3687 | 2970 | 1118 | 865 |
| BiH_245 | Dinaric-Balkan | 3746 | 2868 | 1131 | 853 |
| LUP006817 | Dinaric-Balkan | 3680 | 2914 | 1065 | 873 |
| LUP006818 | Dinaric-Balkan | 3694 | 2881 | 1076 | 882 |
| LUP006819 | Dinaric-Balkan | 3652 | 3096 | 1038 | 937 |
| LUP006820 | Dinaric-Balkan | 3609 | 2894 | 1057 | 873 |
| LUP006821 | Dinaric-Balkan | 3707 | 3053 | 1090 | 894 |
| LUP006822 | Dinaric-Balkan | 3667 | 2921 | 1078 | 884 |
| LUP006823 | Dinaric-Balkan | 3794 | 2769 | 1075 | 832 |
| LUP006824 | Dinaric-Balkan | 3697 | 2848 | 1091 | 879 |
| LUP006825 | Dinaric-Balkan | 3676 | 2974 | 1073 | 907 |
| LUP006826 | Dinaric-Balkan | 3713 | 2898 | 1139 | 795 |
| LUP006827 | Dinaric-Balkan | 3798 | 2807 | 1101 | 815 |
| LUP006828 | Dinaric-Balkan | 3818 | 2716 | 1061 | 851 |
| MSV2AC | Dinaric-Balkan | 3823 | 2756 | 1118 | 819 |
| JAL7480 | NW Iberia | 4268 | 1735 | 1251 | 569 |
| JAL7486 | NW Iberia | 4201 | 1976 | 1243 | 644 |

| | | | | | |
|---|---|---|---|---|---|
| JAL7491 | NW Iberia | 3818 | 2602 | 1108 | 783 |
| JAL7492 | NW Iberia | 4129 | 1974 | 1203 | 631 |
| L166 | NW Iberia | 3887 | 2437 | 1139 | 751 |
| L253 | NW Iberia | 3914 | 2500 | 1165 | 760 |
| L409 | NW Iberia | 3977 | 2351 | 1170 | 718 |
| L514 | NW Iberia | 3852 | 2561 | 1151 | 742 |
| L547 | NW Iberia | 4036 | 2167 | 1224 | 671 |
| L552 | NW Iberia | 3869 | 2508 | 1130 | 779 |
| L588 | NW Iberia | 3997 | 2238 | 1222 | 683 |
| LUP006829 | NW Iberia | 4243 | 1868 | 1203 | 606 |
| LUP006830 | NW Iberia | 3837 | 2675 | 1113 | 823 |
| MW122 | NW Iberia | 3861 | 2486 | 1170 | 730 |
| MW127 | NW Iberia | 4093 | 2021 | 1224 | 635 |
| Penelope | NW Iberia | 3922 | 2427 | 1138 | 779 |
| ptw | NW Iberia | 4045 | 2151 | 1159 | 695 |
| W1023 | Italian Peninsula | 4156 | 2100 | 1205 | 641 |
| W1551 | Italian Peninsula | 4023 | 2239 | 1176 | 678 |
| W1593 | Italian Peninsula | 3792 | 2610 | 1087 | 828 |
| W2030 | Italian Peninsula | 3966 | 2344 | 1154 | 760 |
| W2124 | Italian Peninsula | 3982 | 2304 | 1130 | 721 |
| W2185 | Italian Peninsula | 4023 | 2182 | 1147 | 678 |
| W2193 | Italian Peninsula | 3796 | 2492 | 1115 | 759 |
| W2265 | Italian Peninsula | 3933 | 2367 | 1135 | 753 |
| W2280 | Italian Peninsula | 3848 | 2524 | 1140 | 792 |
| W2291 | Italian Peninsula | 3952 | 2347 | 1147 | 702 |
| W2371F | Italian Peninsula | 4156 | 2029 | 1180 | 687 |
| W2430F | Italian Peninsula | 4010 | 2287 | 1151 | 730 |
| W2487F | Italian Peninsula | 4205 | 2029 | 1212 | 647 |
| W2594M | Italian Peninsula | 4062 | 2268 | 1176 | 709 |
| W2623M | Italian Peninsula | 4020 | 2322 | 1196 | 712 |
| W2707M | Italian Peninsula | 3999 | 2321 | 1161 | 722 |

| | | | | | |
|---|---|---|---|---|---|
| W2878M | Italian Peninsula | 3974 | 2283 | 1186 | 692 |
| W2883F | Italian Peninsula | 4116 | 2211 | 1190 | 686 |
| W2886 | Italian Peninsula | 3936 | 2403 | 1128 | 751 |
| W2910 | Italian Peninsula | 4005 | 2287 | 1134 | 729 |
| W2951 | Italian Peninsula | 3953 | 2367 | 1178 | 739 |
| W893 | Italian Peninsula | 4116 | 2216 | 1187 | 718 |
| WIT11 | Italian Peninsula | 4038 | 2206 | 1189 | 656 |
| WIT13 | Italian Peninsula | 4205 | 1968 | 1252 | 586 |
| WIT14 | Italian Peninsula | 4024 | 2245 | 1183 | 719 |
| WIT16 | Italian Peninsula | 4183 | 2078 | 1217 | 645 |
| WIT17 | Italian Peninsula | 3911 | 2389 | 1147 | 723 |
| WIT19 | Italian Peninsula | 3934 | 2295 | 1146 | 724 |
| WIT20 | Italian Peninsula | 4261 | 1894 | 1240 | 620 |
| WIT21 | Italian Peninsula | 4054 | 2162 | 1154 | 683 |
| WIT22 | Italian Peninsula | 4046 | 2329 | 1197 | 685 |
| WIT23 | Italian Peninsula | 3997 | 2460 | 1179 | 764 |
| w361703 | Italian Peninsula | 3970 | 2265 | 1159 | 723 |
| w361756 | Italian Peninsula | 3836 | 2423 | 1109 | 761 |
| w4926402 | Italian Peninsula | 3701 | 2672 | 1047 | 849 |
| CLUPRU000005 | Karelian | 3798 | 2798 | 1148 | 816 |
| CLUPRU000007 | Karelian | 3611 | 3037 | 1054 | 971 |
| CLUPRU000008 | Karelian | 3626 | 3128 | 1093 | 892 |
| CLUPRU000019 | Karelian | 3686 | 3067 | 1112 | 875 |
| CLUPRU000020 | Karelian | 3730 | 2918 | 1093 | 910 |
| V113 | Karelian | 3813 | 2752 | 1098 | 837 |
| V114 | Karelian | 3497 | 3181 | 1049 | 957 |
| V115 | Karelian | 3645 | 3108 | 1055 | 911 |
| V116 | Karelian | 3710 | 2991 | 1078 | 918 |
| V117 | Karelian | 3728 | 3022 | 1080 | 924 |
| V119 | Karelian | 3713 | 3061 | 1065 | 979 |
| V120 | Karelian | 3638 | 3081 | 1065 | 929 |

| | | | | | |
|---|---|---|---|---|---|
| V126 | Karelian | 3712 | 2933 | 1101 | 903 |
| V132 | Karelian | 3650 | 3047 | 1077 | 917 |
| V134 | Karelian | 3645 | 3109 | 1069 | 881 |
| V136 | Karelian | 3725 | 2914 | 1070 | 909 |
| V141 | Karelian | 3677 | 3020 | 1057 | 895 |
| V143 | Karelian | 3665 | 2960 | 1093 | 902 |
| W11 | Karelian | 3742 | 2941 | 1085 | 939 |
| W14 | Karelian | 3720 | 2951 | 1059 | 925 |
| W16 | Karelian | 3899 | 2626 | 1102 | 833 |
| W21 | Karelian | 3658 | 2998 | 1031 | 923 |
| W26 | Karelian | 3851 | 2853 | 1094 | 860 |
| W32 | Karelian | 3693 | 3004 | 1076 | 942 |
| W4 | Karelian | 4059 | 2311 | 1180 | 714 |
| W44 | Karelian | 3669 | 3093 | 1027 | 967 |
| W45 | Karelian | 3610 | 3049 | 1054 | 931 |
| D-00-12 | Scandinavian | 4105 | 2209 | 1157 | 713 |
| D-06-14 | Scandinavian | 4151 | 2143 | 1167 | 649 |
| D-07-16 | Scandinavian | 4377 | 1593 | 1199 | 554 |
| D-08-10 | Scandinavian | 4326 | 1728 | 1220 | 587 |
| D-10-68 | Scandinavian | 4077 | 2238 | 1156 | 682 |
| D-11-58 | Scandinavian | 4020 | 2243 | 1138 | 711 |
| D-85-02 | Scandinavian | 3659 | 3030 | 1054 | 913 |
| G100-12 | Scandinavian | 3921 | 2556 | 1105 | 785 |
| G100-14 | Scandinavian | 3954 | 2400 | 1161 | 760 |
| G126-13 | Scandinavian | 4071 | 2241 | 1133 | 772 |
| G139-12 | Scandinavian | 3883 | 2577 | 1113 | 780 |
| G50-12 | Scandinavian | 4271 | 1804 | 1199 | 570 |
| M-01-06 | Scandinavian | 4100 | 2123 | 1157 | 663 |
| M-06-03 | Scandinavian | 4299 | 1879 | 1213 | 598 |
| M-09-05 | Scandinavian | 4137 | 2121 | 1168 | 635 |
| M-10-04 | Scandinavian | 3816 | 2746 | 1057 | 890 |

| | | | | | |
|---|---|---|---|---|---|
| M-98-08 | Scandinavian | 3781 | 2874 | 1065 | 890 |

| MOD_DEL Realized load | MOD_DEL Masked load | HIGH Realized load | HIGH Masked load |
|---|---|---|---|
| 74 | 76 | 77 | 60 |
| 59 | 98 | 67 | 74 |
| 69 | 78 | 78 | 58 |
| 72 | 79 | 82 | 53 |
| 65 | 84 | 77 | 57 |
| 73 | 72 | 75 | 58 |
| 68 | 80 | 73 | 60 |
| 76 | 69 | 92 | 43 |
| 72 | 78 | 83 | 52 |
| 72 | 83 | 76 | 66 |
| 72 | 80 | 77 | 60 |
| 68 | 85 | 71 | 67 |
| 72 | 74 | 82 | 63 |
| 65 | 86 | 69 | 69 |
| 73 | 79 | 77 | 63 |
| 64 | 85 | 73 | 57 |
| 60 | 88 | 75 | 53 |
| 64 | 90 | 72 | 74 |
| 67 | 73 | 74 | 61 |
| 53 | 91 | 73 | 66 |
| 59 | 95 | 74 | 62 |
| 66 | 93 | 74 | 67 |
| 71 | 76 | 67 | 67 |
| 64 | 82 | 71 | 69 |
| 67 | 88 | 75 | 57 |
| 62 | 90 | 72 | 60 |
| 65 | 82 | 69 | 72 |
| 69 | 82 | 77 | 68 |
| 72 | 63 | 87 | 37 |
| 73 | 72 | 88 | 50 |

| | | | |
|---|---|---|---|
| 60 | 81 | 72 | 66 |
| 70 | 69 | 73 | 54 |
| 67 | 72 | 83 | 56 |
| 70 | 72 | 80 | 58 |
| 66 | 73 | 77 | 50 |
| 66 | 80 | 76 | 62 |
| 73 | 68 | 86 | 42 |
| 56 | 86 | 70 | 71 |
| 67 | 77 | 77 | 58 |
| 71 | 67 | 79 | 46 |
| 64 | 85 | 76 | 59 |
| 66 | 76 | 87 | 56 |
| 63 | 76 | 77 | 53 |
| 57 | 84 | 73 | 68 |
| 59 | 81 | 85 | 43 |
| 63 | 89 | 88 | 42 |
| 74 | 78 | 86 | 39 |
| 71 | 81 | 77 | 62 |
| 79 | 70 | 73 | 53 |
| 71 | 77 | 80 | 45 |
| 67 | 72 | 80 | 45 |
| 64 | 89 | 73 | 57 |
| 70 | 82 | 78 | 51 |
| 74 | 78 | 78 | 55 |
| 61 | 86 | 79 | 54 |
| 73 | 73 | 89 | 40 |
| 71 | 81 | 79 | 53 |
| 70 | 79 | 81 | 51 |
| 61 | 86 | 77 | 51 |
| 65 | 86 | 81 | 49 |
| 75 | 80 | 73 | 62 |

| | | | |
|---|---|---|---|
| 66 | 91 | 73 | 60 |
| 64 | 88 | 75 | 56 |
| 72 | 77 | 72 | 57 |
| 59 | 89 | 71 | 58 |
| 71 | 76 | 76 | 51 |
| 73 | 84 | 82 | 53 |
| 83 | 67 | 86 | 47 |
| 73 | 75 | 95 | 35 |
| 72 | 80 | 80 | 52 |
| 81 | 67 | 77 | 54 |
| 63 | 88 | 72 | 57 |
| 68 | 86 | 71 | 54 |
| 81 | 63 | 88 | 39 |
| 70 | 73 | 75 | 48 |
| 73 | 73 | 84 | 47 |
| 77 | 74 | 81 | 59 |
| 62 | 82 | 84 | 45 |
| 63 | 79 | 71 | 61 |
| 62 | 90 | 67 | 62 |
| 62 | 91 | 83 | 52 |
| 70 | 88 | 78 | 68 |
| 67 | 85 | 72 | 68 |
| 60 | 85 | 65 | 73 |
| 61 | 91 | 77 | 60 |
| 69 | 81 | 68 | 71 |
| 61 | 90 | 74 | 74 |
| 61 | 91 | 69 | 68 |
| 57 | 102 | 69 | 72 |
| 66 | 85 | 65 | 81 |
| 59 | 95 | 73 | 68 |
| 69 | 86 | 73 | 68 |

| | | | |
|---|---|---|---|
| 50 | 99 | 74 | 74 |
| 67 | 85 | 68 | 71 |
| 57 | 88 | 73 | 68 |
| 66 | 78 | 76 | 63 |
| 63 | 87 | 81 | 57 |
| 61 | 89 | 69 | 70 |
| 58 | 105 | 80 | 57 |
| 74 | 77 | 71 | 73 |
| 66 | 81 | 76 | 63 |
| 55 | 90 | 72 | 66 |
| 62 | 83 | 74 | 72 |
| 50 | 101 | 77 | 67 |
| 69 | 75 | 77 | 64 |
| 52 | 102 | 70 | 73 |
| 50 | 100 | 66 | 71 |
| 58 | 98 | 79 | 54 |
| 72 | 71 | 80 | 57 |
| 80 | 65 | 92 | 44 |
| 65 | 81 | 88 | 42 |
| 63 | 83 | 86 | 55 |
| 67 | 82 | 89 | 45 |
| 58 | 95 | 68 | 70 |
| 64 | 77 | 70 | 62 |
| 61 | 77 | 83 | 60 |
| 63 | 91 | 77 | 58 |
| 71 | 76 | 76 | 64 |
| 73 | 78 | 77 | 48 |
| 67 | 75 | 89 | 43 |
| 72 | 74 | 86 | 56 |
| 63 | 87 | 77 | 65 |
| 65 | 82 | 70 | 73 |

| | | | |
|---|---|---|---|
| 59 | 95 | 73 | 70 |

**Table S17.** Statistical comparisons between genetic load counts among European wolf population pairs, focusing on deleterious variants.

| Comparison | Load | Category | median_pop1 | median_pop2 | p_value | p_adj_FDR |
|---|---|---|---|---|---|---|
| Dinaric-Balkan vs Iberian | Masked | HIGH | 61,5 | 56 | 0,007525290018 | 0,01254215003 |
| Dinaric-Balkan vs Iberian | Masked | MOD_DEL | 82 | 76 | 0,003386830704 | 0,01128943568 |
| Dinaric-Balkan vs Italian Peninsula | Masked | HIGH | 61,5 | 53 | 1,59E-06 | 7,95E-06 |
| Dinaric-Balkan vs Italian Peninsula | Masked | MOD_DEL | 82 | 80 | 0,1660408747 | 0,2075510934 |
| Dinaric-Balkan vs Karelian | Masked | HIGH | 61,5 | 68 | 0,002585516347 | 0,005171032694 |
| Dinaric-Balkan vs Karelian | Masked | MOD_DEL | 82 | 88 | 0,004740791148 | 0,01185197787 |
| Dinaric-Balkan vs Scandinavian | Masked | HIGH | 61,5 | 57 | 0,08697925063 | 0,1087240633 |
| Dinaric-Balkan vs Scandinavian | Masked | MOD_DEL | 82 | 81 | 0,5499787772 | 0,5844865974 |
| Iberian vs Italian Peninsula | Masked | HIGH | 56 | 53 | 0,2531806321 | 0,2813118134 |
| Iberian vs Italian Peninsula | Masked | MOD_DEL | 76 | 80 | 0,04214917139 | 0,07024861898 |
| Iberian vs Karelian | Masked | HIGH | 56 | 68 | 2,34E-05 | 7,80E-05 |
| Iberian vs Karelian | Masked | MOD_DEL | 76 | 88 | 4,50E-06 | 4,50E-05 |
| Iberian vs Scandinavian | Masked | HIGH | 56 | 57 | 0,5348314309 | 0,5348314309 |
| Iberian vs Scandinavian | Masked | MOD_DEL | 76 | 81 | 0,04930015807 | 0,07042879724 |
| Italian Peninsula vs Karelian | Masked | HIGH | 53 | 68 | 8,22E-10 | 8,22E-09 |
| Italian Peninsula vs Karelian | Masked | MOD_DEL | 80 | 88 | 6,69E-05 | 0,0003346383197 |
| Italian Peninsula vs Scandinavian | Masked | HIGH | 53 | 57 | 0,05812623053 | 0,08303747219 |
| Italian Peninsula vs Scandinavian | Masked | MOD_DEL | 80 | 81 | 0,5844865974 | 0,5844865974 |
| Karelian vs Scandinavian | Masked | HIGH | 68 | 57 | 0,0004018202087 | 0,001004550522 |
| Karelian vs Scandinavian | Masked | MOD_DEL | 88 | 81 | 0,00764893601 | 0,01529787202 |
| Dinaric-Balkan vs Iberian | Realized | HIGH | 76,5 | 78 | 0,1693727518 | 0,2037057276 |
| Dinaric-Balkan vs Iberian | Realized | MOD_DEL | 51 | 60 | 9,00E-07 | 3,00E-06 |
| Dinaric-Balkan vs Italian Peninsula | Realized | HIGH | 76,5 | 83 | 1,42E-07 | 7,10E-07 |
| Dinaric-Balkan vs Italian Peninsula | Realized | MOD_DEL | 51 | 51 | 0,8401981243 | 0,8401981243 |
| Dinaric-Balkan vs Karelian | Realized | HIGH | 76,5 | 73 | 0,002219435265 | 0,00443887053 |
| Dinaric-Balkan vs Karelian | Realized | MOD_DEL | 51 | 50 | 0,6002948261 | 0,6669942512 |
| Dinaric-Balkan vs Scandinavian | Realized | HIGH | 76,5 | 76 | 0,953215676 | 0,953215676 |
| Dinaric-Balkan vs Scandinavian | Realized | MOD_DEL | 51 | 54 | 0,1241991238 | 0,2069985396 |
| Iberian vs Italian Peninsula | Realized | HIGH | 78 | 83 | 0,07981958604 | 0,1140279801 |
| Iberian vs Italian Peninsula | Realized | MOD_DEL | 60 | 51 | 5,32E-07 | 2,66E-06 |
| Iberian vs Karelian | Realized | HIGH | 78 | 73 | 0,0007499876633 | 0,001874969158 |

| | | | | | | |
|---|---|---|---|---|---|---|
| Iberian vs Karelian | Realized | MOD_DEL | 60 | 50 | 3,64E-07 | 2,66E-06 |
| Iberian vs Scandinavian | Realized | HIGH | 78 | 76 | 0,1833351548 | 0,2037057276 |
| Iberian vs Scandinavian | Realized | MOD_DEL | 60 | 54 | 0,0004815442576 | 0,001203860644 |
| Italian Peninsula vs Karelian | Realized | HIGH | 83 | 73 | 2,01E-10 | 2,01E-09 |
| Italian Peninsula vs Karelian | Realized | MOD_DEL | 51 | 50 | 0,3913191926 | 0,4891489907 |
| Italian Peninsula vs Scandinavian | Realized | HIGH | 83 | 76 | 4,47E-05 | 0,0001490044618 |
| Italian Peninsula vs Scandinavian | Realized | MOD_DEL | 51 | 54 | 0,1526764918 | 0,2181092741 |
| Karelian vs Scandinavian | Realized | HIGH | 73 | 76 | 0,02685183548 | 0,04475305913 |
| Karelian vs Scandinavian | Realized | MOD_DEL | 50 | 54 | 0,07355084865 | 0,1471016973 |
